# Conserved influenza A epitope candidate regions and a benchmark of ESM-2 sequence features

**DOI:** 10.64898/2026.08.16.745106

**Authors:** Qingxiu Li, Zhenjun Li

## Abstract

Influenza A virus antigenic drift forces annual vaccine reformulation, motivating the search for conserved epitope candidates that could support broadly protective vaccines. We systematically screened influenza A virus sequences (H1N1, H3N2, H5N1; nine viral proteins) to define 98 conserved candidate regions, 38 of which were identical across the H1N1, H3N2, and H5N1 consensus sequences — all in the polymerase complex and nucleoprotein (PB2, PB1, PA, NP) — whereas the ten surface-glycoprotein (HA/NA) candidates were subtype-specific. We then benchmarked two protein-language-model (ESM-2) features against alignment conservation. Group-masked log-probability correlated moderately with MSA conservation (Spearman ρ = 0.25–0.39 for HA) but provided no incremental value for T-cell epitope discrimination (ΔAUROC +0.004, p = 0.46); attention-derived contact-density was not a valid solvent-accessibility proxy. A curated antibody-epitope benchmark (22 clusters, 5 neutralization-supported) was underpowered for a high-confidence B-cell test. We document data-quality and reproducibility pitfalls (length heterogeneity, coordinate mapping, and pseudoreplication) and release the auditable benchmark. These results provide an auditable candidate resource and show that, in the evaluated benchmarks, ESM-2 sequence scores did not improve epitope prioritization beyond alignment-derived conservation.

## Introduction

IAV remains a major health burden, with seasonal epidemics causing substantial morbidity and mortality (Iuliano et al., 2018). Because hemagglutinin (HA) and neuraminidase (NA) undergo continuous antigenic drift, strain-specific vaccines must be reformulated annually, motivating a long-standing goal of broadly protective (“universal”) vaccines against conserved viral determinants (Krammer and Palese, 2015; Wei et al., 2020; Erbelding et al., 2018).

Conserved regions of the viral proteome are the natural substrate for such vaccines. Systematic computational screens identify them by combining multiple-sequence-alignment (MSA) conservation with B-cell and T-cell epitope prediction, then prioritizing candidates by structural and functional criteria. A central limitation is that positional conservation alone ignores co-evolutionary and structural context, and does not distinguish surface-exposed from buried residues.

Protein language models (pLMs) such as ESM-2 are trained with a masked-language objective on hundreds of millions of sequences and learn representations that implicitly encode evolutionary and structural information (Rives et al., 2021; Lin et al., 2023; Rao et al., 2019). Their masked-language log-likelihood has been used as a proxy for mutational constraint (Meier et al., 2021) and to study evolutionary dynamics, including viral immune escape (Hie et al., 2022). Whether these features provide incremental value — beyond cheap alignment conservation — for the specific task of epitope prioritization remains an open, largely untested question.

Here we address two aims. First, we report a systematic screen for conserved epitope candidates across nine IAV proteins. Second, we critically evaluate whether two ESM-2 features (group-masked residue log-probability and attention-derived contact-density) improve prioritization over conservation, using matched benchmarks and, where possible, experimentally derived antibody epitopes. We emphasize honesty about negative findings and about the statistical power of the available validation data.

## Results

### 1. Systematic screen for conserved epitope candidates

We built majority-rule consensus amino-acid sequences for nine IAV proteins (HA, NA, M1, NP, NEP, NS1, PA, PB1, PB2) within each subtype (H1N1, H3N2, H5N1), after retaining mode-length translated sequences (Table 1). Raw per-protein files were heterogeneous in length (40–49 distinct nucleotide lengths per HA file; Supplementary Note 1), so mode-length filtering was necessary to obtain full-length, in-frame sequences before positional conservation was computed. The overall screening workflow is summarized in Fig. 1a.

**Fig 1.**
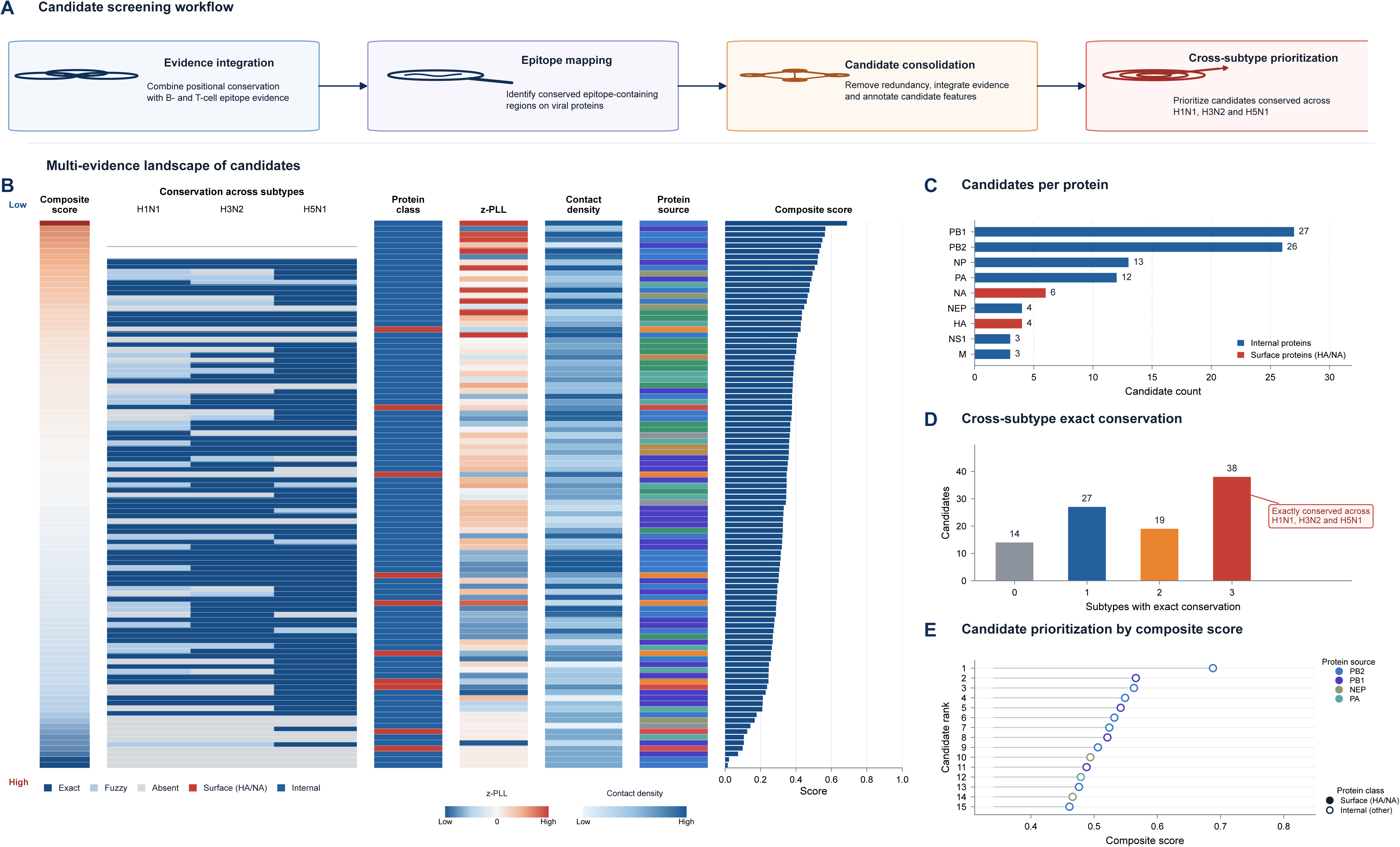
Systematic screening and multi-evidence prioritization of conserved influenza A epitope candidates. (a) Candidate screening workflow, from evidence integration through epitope mapping and redundancy removal to cross-subtype prioritization. (b) Multi-evidence landscape of the 98 candidate regions, ranked by composite score: composite-score colour strip, conservation status across H1N1/H3N2/H5N1 (exact/fuzzy/absent), protein class (surface = HA/NA, red; internal, blue), ESM-2 z-PLL, contact density, protein source, and composite score. (c) Number of candidate regions per viral protein. (d) Distribution of candidates by the number of subtypes in which they are exactly conserved; 38 candidates are identical across the three subtype consensus sequences. (e) Composite scores of the top 15 candidates, coloured by protein source; filled and open circles denote surface (HA/NA) and internal proteins, respectively.

**Table 1.** Sequence and consensus statistics. Raw and mode-length-filtered sequence counts per subtype × protein (H1N1: 7,110 raw / 4,654 filtered; H3N2: 5,899 / 3,192; H5N1: 2,320 / 811; full table in Supplementary Table S1).

Combining positional conservation (five thresholds: 0.5, 0.6, 0.7, 0.8, 0.9), 16-mer linear B-cell epitopes, and netMHCpan-4.1 T-cell epitopes (14 HLA-I alleles, %Rank_EL < 2% and affinity < 500 nM), we obtained 105 epitope-containing regions, which collapsed to 98 unique candidate regions after sequence de-duplication (Table 3; Supplementary Table S1). These candidate regions span the surface glycoproteins HA and NA as well as internal proteins (NP, M1, NEP, PA, PB1, PB2; Fig. 1c).

### 2. Cross-subtype and temporal conservation

Of the 98 candidate regions, 38 were identical across the H1N1, H3N2, and H5N1 consensus sequences (Fig. 1b, Fig. 1d), but all 38 map to the polymerase complex and nucleoprotein (PB2, PB1, PA, NP); no surface glycoprotein candidate was conserved across all three subtypes. These 38 candidates were also highly conserved at the isolate level: on average, 99.1%, 97.3%, and 98.0% of H1N1, H3N2, and H5N1 isolates retained ≥90% identity, and 85.8%, 78.7%, and 83.2% matched exactly (Table 3). The ten HA/NA candidates were instead subtype-specific: the HA candidate CKLRGVAPLHLGKCNIAGW is H1N1-specific (absent from the H3N2 and H5N1 consensus sequences), five NA candidates are H5N1-specific, and one (RYPGVRCVC, at the NA enzyme active site) is H3N2-specific. Temporal analysis within the native subtype showed that the top HA candidate was maintained at 53–100% sequence identity across 11 consecutive years (2007–2017) in H1N1, and the H3N2-native NA candidate RYPGVRCVC was conserved at 99– 100% across 2010–2025 (Table 2), with no clear monotonic decline observed. These regions are therefore conserved within, but not across, subtypes — a distinction that constrains their utility as broadly cross-subtype immunogens.

**Table 2.** Temporal and lineage conservation of the selected HA candidate (HA-34; CKLRGVAPLHLGKCNIAGW). (a) Temporal conservation (H1N1 HA, 2006–2017); (b) lineage conservation (H1N1, 2015–2025); (c) stability summary. Conserved fraction = proportion of sequences with ≥90% identity. In panel (c), cross-subtype support is the conserved fraction in the H3N2 consensus, which is 0 because this candidate is H1N1-specific.

**Table 3.** The 98 candidate regions. (protein, sequence, consensus coordinates, cross-subtype conservation with isolate-level exact and ≥90% identity fractions per subtype, B/T score, PLL, contact-density).

To understand the selective forces underlying this conservation, we estimated per-codon dN/dS (Nei–Gojobori) on 1,373 in-frame H1N1 HA nucleotide sequences. Whole-HA dN/dS was 0.645, consistent with predominant purifying selection (Fig. 4d). The candidate regions avoided the ten high-frequency (>90%) mutation positions identified in the alignment, indicating that they do not coincide with the fastest-drifting residues. Site-level dN/dS estimates were unstable and were not treated as formal evidence of positive selection; as shown above, this constraint does not extend across subtypes.

For post-discovery temporal validation, we queried a companion 2015–2025 H1N1 HA dataset (2,219 sequences with clade annotations); the 2018–2025 subset constitutes the post-discovery validation, whereas 2015–2017 overlaps the discovery window and is shown only as part of the full temporal trend. The top HA candidate CKLRGVAPLHLGKCNIAGW was conserved at 96–100% identity across all six major recent lineages (6B.1A.5a.2a, 6B.1A, 6B.1, 6B.1A.5a, 6B.2, 6B.1A.5a.1), with minimal between-lineage variation (0.013; Table 2). This cross-lineage conservation within H1N1 provides forward-looking support that the candidate region remains functionally constrained in currently circulating strains. To test whether this conservation extends across subtypes, we queried a later-downloaded external-source validation dataset of 2010–2025 H3N2 sequences (4,265 HA and 4,264 NA, with clade annotations). The H1N1-specific HA candidate CKLRGVAPLHLGKCNIAGW was absent (0% at ≥90% identity) from all H3N2 clades and years, and the five H5N1-derived NA candidates were likewise absent; only the H3N2-native NA candidate RYPGVRCVC was conserved (99–100%). Surface glycoprotein candidates are therefore conserved across lineages within their native subtype but not across subtypes (Supplementary Fig. S2).

### 3. Prioritization and illustrative assemblies

Candidates were ranked using a heuristic composite score = 0.4×normalized B/T score + 0.3×normalized PLL + 0.2×normalized contact-density + 0.1×normalized match count. Two caveats apply. First, the weights are not trained or validated, and the top-10 ranking had only moderate stability across alternative weight schemes (Jaccard similarity 0.43–0.54), so the exact ordering is exploratory (Fig. 1e). Second, and more importantly, the ESM-2 components of this score (PLL and contact-density) are shown below to provide no incremental value over conservation (§4). Accordingly, conservation and epitope evidence constitute the primary prioritization signal, and the ESM-2 features are reported separately as a post hoc benchmark rather than as components of a validated ranking. From the top-ranked regions we assembled illustrative multi-epitope assemblies (HA and NA, excluding transmembrane segments). AlphaFold2 predictions of these fragments showed modest confidence (pLDDT 55.9–64.3; pTM 0.28–0.42), insufficient to claim structural feasibility; the assemblies are schematic illustrations rather than validated immunogen designs.

### 4. Critical benchmark of ESM-2 sequence features

To test whether ESM-2 adds value over conservation, we computed two zero-shot features per residue: a group-masked residue log-probability (PLL) and an attention-derived contact-density score.

#### Group-masked log-probability overlaps partially with conservation and localizes to functional regions

PLL correlated moderately with MSA positional identity (Spearman ρ = 0.25–0.39 for HA; 0.11–0.16 for NA) and negatively with Shannon entropy (ρ = −0.25 to −0.37), consistent with it acting largely as a conservation proxy (Fig. 2a). The correlation is far from 1.0, so PLL does contain orthogonal information, but its dominant signal is conservation. As a positive control for the zero-shot feature, the position-resolved PLL profile recapitulated the functional anatomy of HA without task-specific fine-tuning (Fig. 2b).

**Fig 2.**
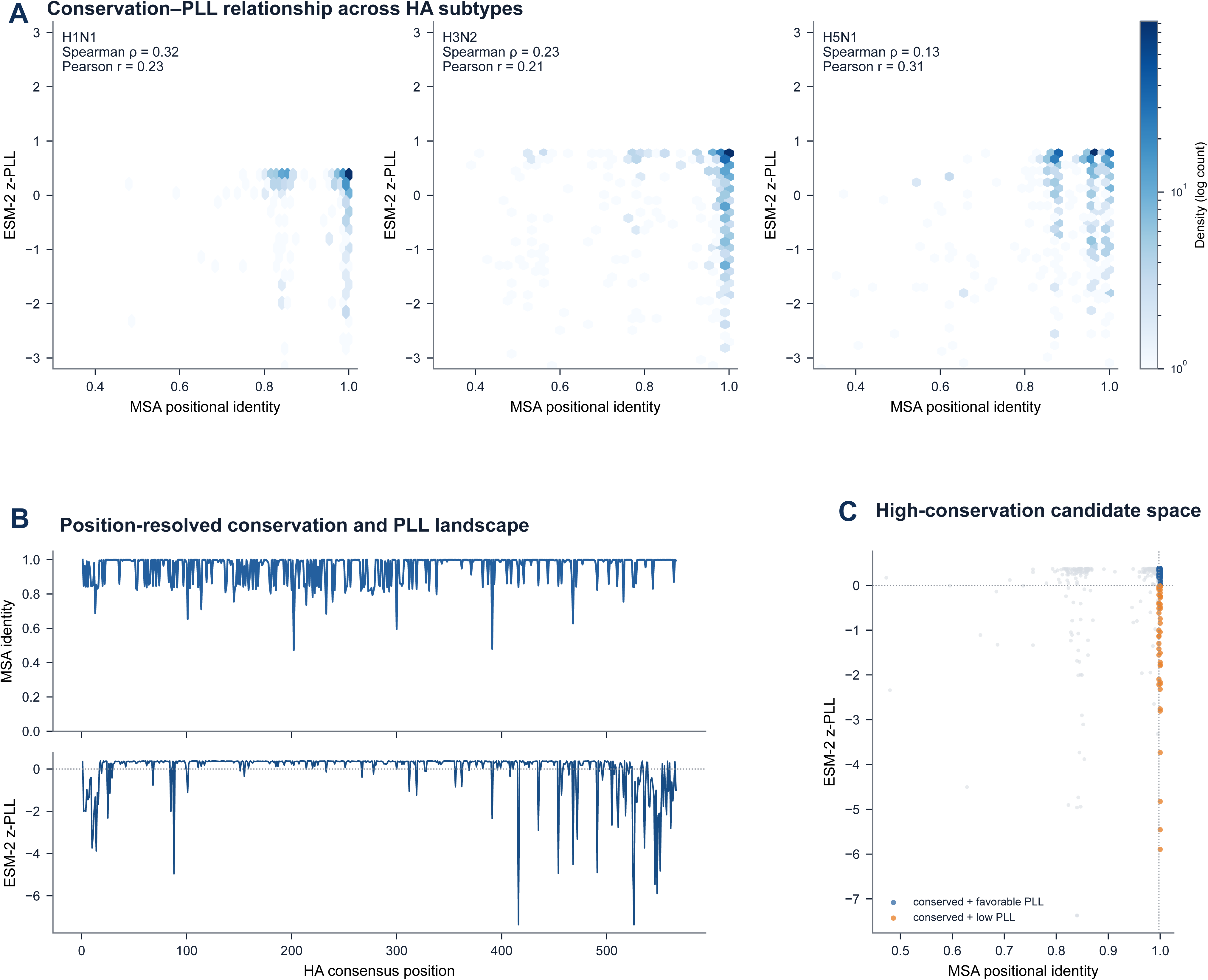
Conservation and ESM-2 PLL relationship across HA subtypes. (a) Hexbin density of ESM-2 z-PLL against MSA positional identity for H1N1, H3N2, and H5N1 HA, with Spearman and Pearson correlations annotated. (b) Position-resolved MSA conservation (upper track) and ESM-2 z-PLL (lower track) along the H1N1 HA consensus. (c) Quadrant view of the H1N1 HA conservation–PLL space, highlighting conserved positions with favourable versus low PLL.

#### PLL provides no incremental value for T-cell epitopes

In a matched benchmark of 101 experimentally characterized IEDB T-cell epitopes against 101 conservation-matched background windows, conservation alone achieved AUROC 0.526, and adding PLL changed it to 0.529 (ΔAUROC +0.004, bootstrap 95% CI −0.07 to +0.08, p = 0.46); adding contact-density gave 0.559 (Δ +0.031, p = 0.12) (Fig. 3a). AUPRC and precision metrics followed the same pattern (Fig. 3b). In this matched dataset, conservation showed little discrimination, so neither it nor group-masked log-probability discriminated T-cell epitopes well.

**Fig 3.**
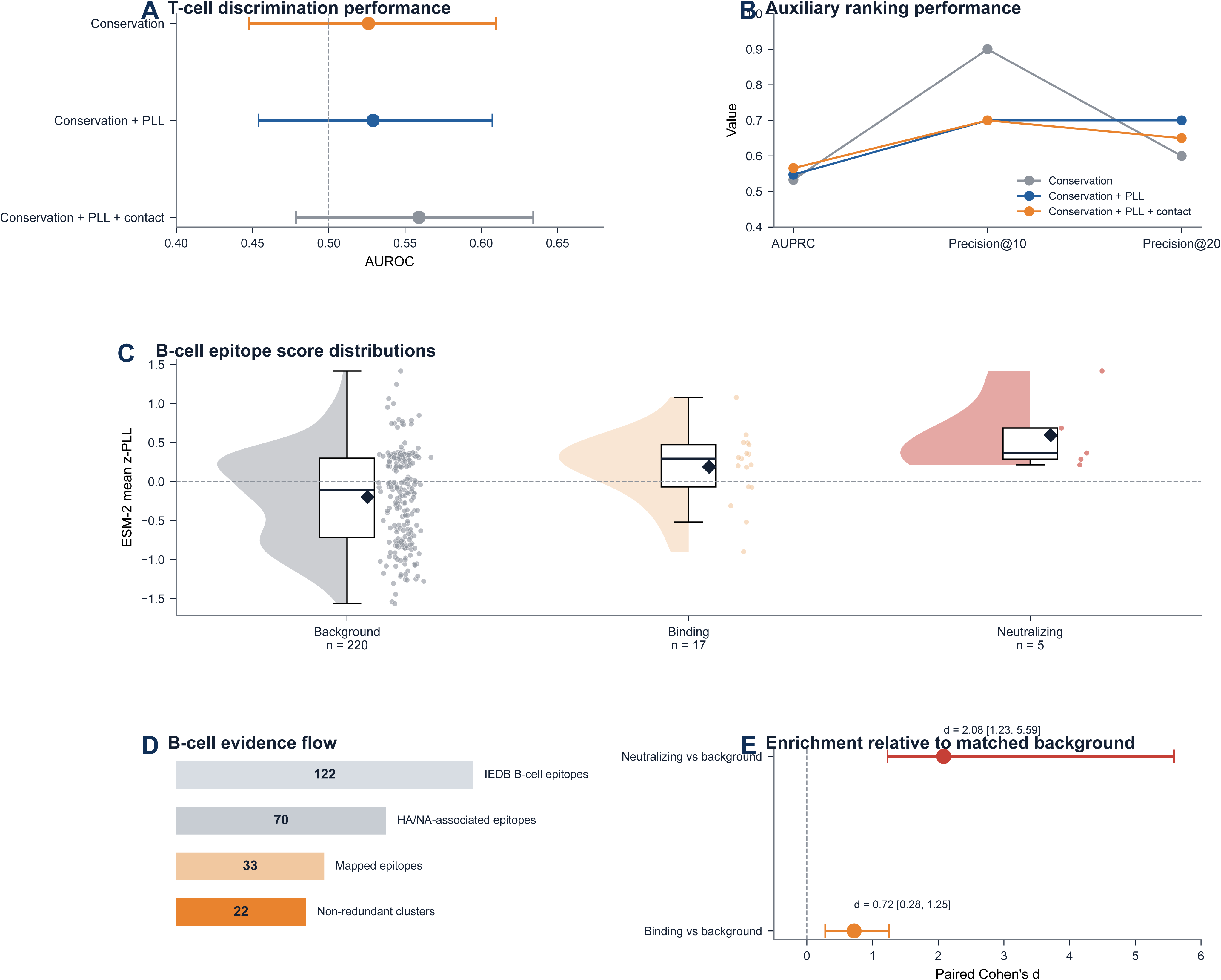
T-cell and B-cell benchmark validation. (a) T-cell discrimination performance: AUROC with 95% CI for conservation, conservation + PLL, and conservation + PLL + contact-density models. (b) Auxiliary ranking metrics (AUPRC, Precision@10, Precision@20) for the same three models. (c) B-cell epitope score distributions (mean z-PLL) for background, binding, and neutralizing epitopes, shown as half-violin with box and raw points. (d) B-cell evidence flow (122 → 70 → 33 → 22). (e) Enrichment relative to matched background (paired Cohen’s d with cluster-level bootstrap 95% CI) for binding and neutralizing epitopes.

#### Contact-density is not a solvent-accessibility proxy

Against FreeSASA-derived relative solvent-accessible surface area (RSA) on experimental structures, contact-density correlated +0.147 (HA, PDB 1RU7) but −0.132 (NA, PDB 2HTY), with inconsistent direction, and barely separated surface from buried residues (Fig. 4a). Contact-density should therefore not be used as an accessibility feature in prioritization.

**Fig 4.**
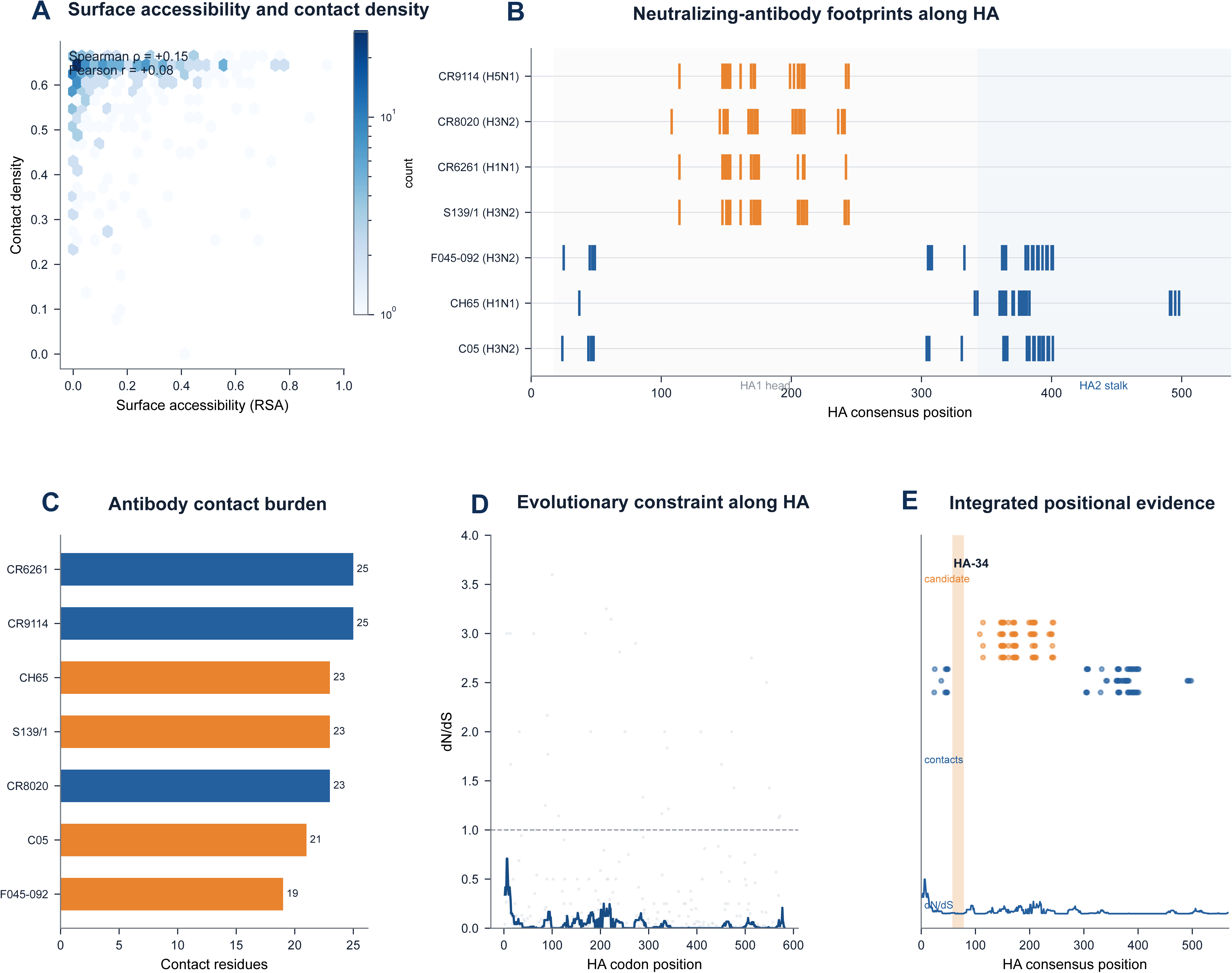
Structural, antibody, and evolutionary context. (a) Surface accessibility (FreeSASA RSA) versus ESM-2 contact density, with Spearman and Pearson correlations annotated. (b) Neutralizing-antibody contact footprints mapped to the HA consensus, coloured by targeting region (head vs stalk). (c) Contact-residue burden per antibody. (d) Per-codon dN/dS along H1N1 HA (raw points and sliding median). (e) Integrated positional evidence: candidate region, antibody contacts, and dN/dS along the HA consensus.

#### Antibody/neutralizing benchmark is underpowered

A curated, non-redundant antibody-epitope benchmark (70 IEDB HA/NA epitopes → 22 clusters after subtype/coordinate mapping and overlap clustering; 5 neutralization-supported; Fig. 3d) is too small for a high-confidence B-cell test (Table 4). Descriptively, group-masked log-probability followed a clear hierarchy — neutralization-supported (+0.59), antibody-binding (+0.19), and conservation-matched background (−0.20) (Fig. 3c). At the cluster level, comparing each epitope with its matched background via a paired Wilcoxon test, epitopes were enriched relative to background overall (p = 2.1×10⁻⁴), driven by the binding set (n = 17, p = 0.004, paired Cohen’s d = 0.72); the neutralization-supported set (n = 5) showed a large but not statistically significant enrichment (paired Cohen’s d = 2.08, p = 0.06), consistent with its small sample size (Fig. 3e). Because neutralization-supported epitopes are conserved by construction, this hierarchy may be driven by conservation rather than by group-masked log-probability-specific signal. As an independent, structurally defined positive set, we mapped the contact footprints of three broadly neutralizing stalk antibodies (CR6261, CR8020, CR9114) to the HA2 stalk, four receptor-binding-site (RBS) antibodies (CH65, C05, S139/1, F045-092) to the HA1 globular head, and one neuraminidase antibody (CD6) to the NA enzyme active site, each mapped to its native subtype consensus (Fig. 4b, Fig. 4c). The three H3N2 RBS antibodies yielded highly overlapping consensus footprints (a positive control for the mapping), whereas none of the ten surface candidate epitopes overlapped with any of the eight neutralizing footprints, so these candidates cannot currently be interpreted as known neutralizing-antibody epitopes. Because these neutralizing epitopes are, by construction, the most conserved regions (Jiao et al., 2023), conservation alone identifies them, leaving little room for PLL to add value.

**Table 4.** Antibody-epitope benchmark statistics. (70 IEDB epitopes → 22 clusters; 5 neutralization-supported; 83 PMIDs).

### 5. Data-quality and reproducibility observations

Several pitfalls emerged that we document for the community: (i) raw per-protein files are length-heterogeneous (partial/frameshifted sequences), so mode-length filtering is essential; (ii) the EBI linear-epitope endpoint misses conformational antibody epitopes, which led an earlier iteration to underestimate available B-cell data (n = 4) by ∼17-fold; (iii) a match-count field conflates “region × subtype × threshold” hits with subtype conservation; and (iv) 33 mapped epitopes collapsed to 22 non-redundant clusters, indicating substantial pseudoreplication that must be handled by cluster-level resampling.

## Discussion

We report two complementary findings. First, a systematic set of conserved influenza epitope candidates — cross-subtype conserved for the internal proteins, and subtype-specific but temporally and lineage-stable for the surface glycoproteins — provided as a computational resource with explicit caveats (no experimental validation). Second, a critical benchmark showing that ESM-2 masked-language likelihood largely overlaps with alignment conservation and provides no measurable incremental value for epitope prioritization, while attention-derived contact-density is not a valid accessibility proxy.

The negative ESM-2 finding is informative: it delineates a boundary of pLM utility. Masked-language likelihood encodes sequence conservation, which is cheaply available from alignments; for conservation-dependent tasks such as neutralizing-epitope identification, alignment conservation is already sufficient. For T-cell epitopes, neither conservation nor likelihood was predictive in the evaluated benchmark. This distinction remains important because cross-reactive T-cell responses to conserved influenza proteins have been associated with reduced symptomatic disease and are a central rationale for broadly protective vaccine strategies (Sridhar et al., 2013; La Gruta and Turner, 2014). We therefore caution against treating pLM likelihood as a universal “fitness” or “epitope” score without task-specific validation.

Limitations include: the benchmark’s small B-cell sample (22 clusters, 5 neutralization-supported), the 2002–2017 discovery window, and the absence of experimental validation. Future work should expand the antibody-epitope benchmark with additional structurally defined footprints and test the same question across other pathogens and pLMs.

In conclusion, the conserved candidate list is a candidate resource for subsequent experimental evaluation, and our benchmark provides an honest, reusable assessment that ESM-2 sequence features do not currently improve epitope prioritization beyond alignment conservation.

## Methods

### Data and consensus

The discovery dataset comprised influenza A virus sequences downloaded from NCBI/GenBank (Sayers et al., 2025) (full-length genomes; collection years 2002–2017), stratified by subtype (H1N1, H3N2, H5N1) and segmented into nine proteins. Sequences were translated in the standard genetic code; entries with incomplete coding frames were excluded. Within each subtype × protein, the modal translated length was identified, and only sequences of that length with ≤5% ambiguous residues were retained, removing partial, truncated, and frameshifted entries as well as duplicate sequences. Majority-vote consensus sequences were then built per subtype × protein by direct per-position voting on this mode-length set, without a separate alignment step, so consensus coordinates can be offset by insertions/deletions relative to any single sequence. For alignment-aware conservation (positional identity, Shannon entropy), the mode-length sequences were aligned with MAFFT. Two later-downloaded GISAID holdout datasets were used for temporal validation: a 2015–2025 H1N1 HA set (2,219 sequences with clade annotations) and a 2010–2025 H3N2 HA/NA set (4,265 HA and 4,264 NA sequences). These are temporal holdouts rather than independent external cohorts, because they derive from a different source (GISAID) and overlap the discovery window in their earliest years.

### Selection pressure (dN/dS)

Per-codon dN/dS was estimated by the Nei–Gojobori method (Nei and Gojobori, 1986) on 1,373 in-frame, mode-length H1N1 HA nucleotide sequences relative to their majority consensus. Codons with dS = 0 were treated as undefined; gaps and ambiguous bases were excluded from the counting. A sliding median (window = 15 codons) was applied for visualization only, and site-level estimates were not treated as formal evidence of positive selection. High-frequency mutations were identified as positions with a mutation frequency >90% in a companion alignment.

### Epitope prediction

B-cell linear epitopes predicted with ABCpred (Saha and Raghava, 2006) and the IEDB online B-cell prediction tool (16-mer sliding window). netMHCpan-4.1 across 14 HLA-I alleles (HLA-A*01:01, A*02:01, A*11:01, A*24:02, B*07:02, B*15:01, B*15:02, B*44:02, B*46:01, B*51:01, B*58:01, C*03:04, C*07:01, C*07:02; 8–14-mers, %Rank_EL < 2% and Aff < 500 nM) (Reynisson et al., 2020); netMHCIIpan-4.3 across six HLA-DRB1 alleles (DRB1*01:01, 03:01, 04:01, 07:01, 11:01, 15:01) for CD4 epitopes (Nilsson et al., 2023). The screening HLA-I and HLA-DRB1 alleles cover approximately 94.4% and 73.0% of the global population, respectively (estimated with the IEDB Population Coverage tool).

### ESM-2 features

esm2_t33_650M_UR50D, zero-shot. Group-masked log-probability approximation: residues were partitioned into eight groups (∼12.5%), masked, and scored in eight forward passes; this approximation closely matched single-residue masking in the evaluated subset (Spearman ρ = 0.93). Contact-density: 1 − normalized summed contact probability from the attention contact-regression head.

### T-cell benchmark

IEDB T-cell epitopes (n = 103) were located on the subtype consensus sequences, of which 101 could be unambiguously located. For each positive, a single matched non-epitope window of the same length and protein was sampled with positional conservation within ±0.05 of the positive, excluding any known epitope. Three score combinations were compared — conservation alone, conservation + PLL, and conservation + PLL + contact-density — with each feature min-max normalized before summation. AUROC and AUPRC were computed directly on these scores (no model training or cross-validation), and 95% confidence intervals and incremental ΔAUROC were obtained by bootstrap resampling of samples (n = 1000, seed = 42) (Efron, 1979; Efron and Tibshirani, 1994).

### B-cell benchmark

IEDB B-cell epitopes were retrieved via the IEDB query API (source organism Influenza A virus) (Vita et al., 2025) on 2026-08-13. Subtype was assigned from strain names or sequence mapping; consensus coordinates were determined by sequence alignment (exact = PASS, 1-mismatch = AMBIGUOUS, else FAIL). Overlapping epitopes (≥50% consensus-interval overlap) were clustered, collapsing 70 HA/NA epitopes to 22 non-redundant clusters (5 neutralization-supported). Matched background windows were sampled per positive (same protein, length ±20%, conservation ±0.05, 10 per positive, with a 10-residue buffer). For statistical inference, the cluster was the unit of analysis: each cluster was compared with the mean of its matched background windows using a paired Wilcoxon signed-rank test, and effect sizes were reported as paired Cohen’s d with cluster-level bootstrap 95% confidence intervals, so that the non-independent matched background windows did not inflate significance.

### Structure

FreeSASA RSA on experimental HA (PDB 1RU7) and NA (PDB 2HTY); antibody-contact footprints from antibody–antigen complexes (stalk: PDB 3GBN, 3SDY, 4FQI; RBS: PDB 3SM5, 4FP8, 4GMS, 4O58; NA: PDB 4QNP) using a 4.5 Å distance cutoff. Structures were obtained from the Protein Data Bank (Berman et al., 2000); AlphaFold2 was used for illustrative assemblies (Jumper et al., 2021).

### Software

Analyses were performed in Python 3.13 with Biopython, numpy, pandas, scipy, matplotlib, and scikit-learn (Cock et al., 2009; Harris et al., 2020; McKinney, 2010; Virtanen et al., 2020; Hunter, 2007; Pedregosa et al., 2011). ESM-2 (esm2_t33_650M_UR50D) was used zero-shot via the ESM framework; MAFFT v7 for alignment; FreeSASA for solvent-accessibility computation.

## Data and code availability

Discovery sequences are available from NCBI/GenBank. The GISAID holdout sequences are available from GISAID (Shu and McCauley, 2017) under its access policy; their complete accession list (6,483 isolates) is provided as Supplementary Data (gisaid_accessions.tsv). The benchmark datasets (IEDB epitope mappings, consensus sequences, matched backgrounds) are provided in the Supplementary Data; analysis scripts are available from the corresponding author upon reasonable request. ESM-2 is available from https://github.com/facebookresearch/esm.

## Acknowledgements

We gratefully acknowledge all data contributors, i.e., the authors and their originating laboratories responsible for obtaining the specimens, and their submitting laboratories for generating the genetic sequences and metadata and sharing them via the GISAID Initiative, on which part of this research is based.

## Author contributions

Q.L. performed all data analysis and wrote the manuscript. Z.L. conceived and supervised the study, and reviewed and revised the manuscript.

## Supplementary figure legends

**Supplementary Fig. S1.** Complete multi-evidence landscape of all 98 candidate regions (rank, composite score, combined B/T score, cross-subtype conservation, z-PLL, contact density, protein class, and protein source).

**Supplementary Fig. S2.** Temporal and lineage conservation validation: yearly conservation of the H1N1 HA candidate (2006–2017, point size reflecting sequence count), conservation across six H1N1 clades, H3N2 candidate × clade conservation heatmap, and H3N2 clade sample sizes.

**Supplementary Fig. S3.** Model and external benchmark validation: conservation– PLL relationship for three HA subtypes, T-cell benchmark, B-cell score distributions, and B-cell evidence flow.

## Supplementary Note 1

Raw per-protein files contain 29–49 distinct nucleotide lengths (e.g., H1N1 HA: 40 lengths; H5N1 HA: 49 lengths), reflecting partial, truncated, or frameshifted entries. Mode-length filtering (retaining the modal translated length) was therefore applied before alignment and conservation computation.

