## Supplementary material for "Conserved influenza A epitope candidate regions and a benchmark of ESM-2 sequence features": Li_et_al_Supplementary_Figures

### Figure S1 | Complete evidence for all 98 candidate regions (ranked by final score)

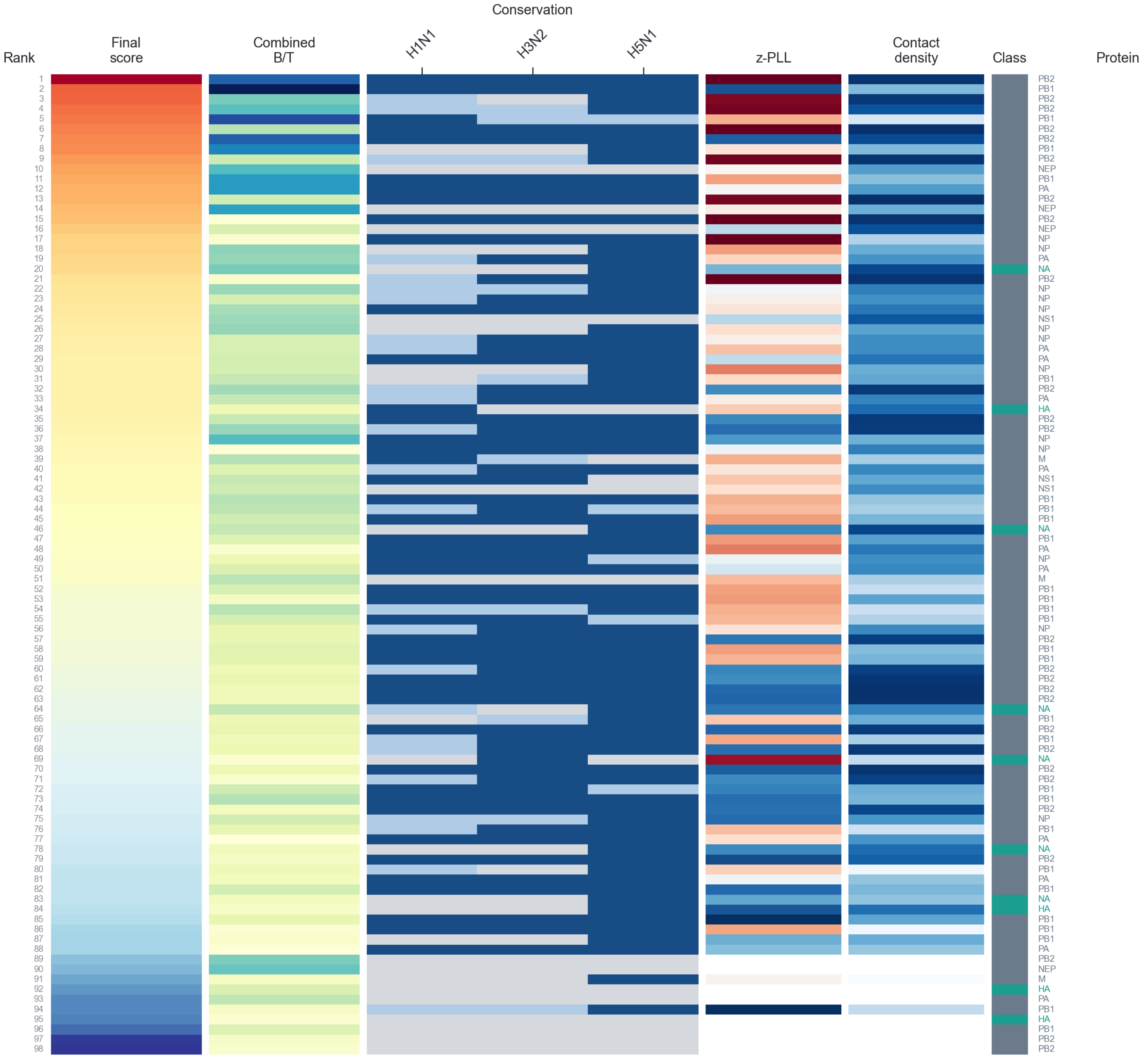

**Figure S2 | Temporal and lineage conservation validation**

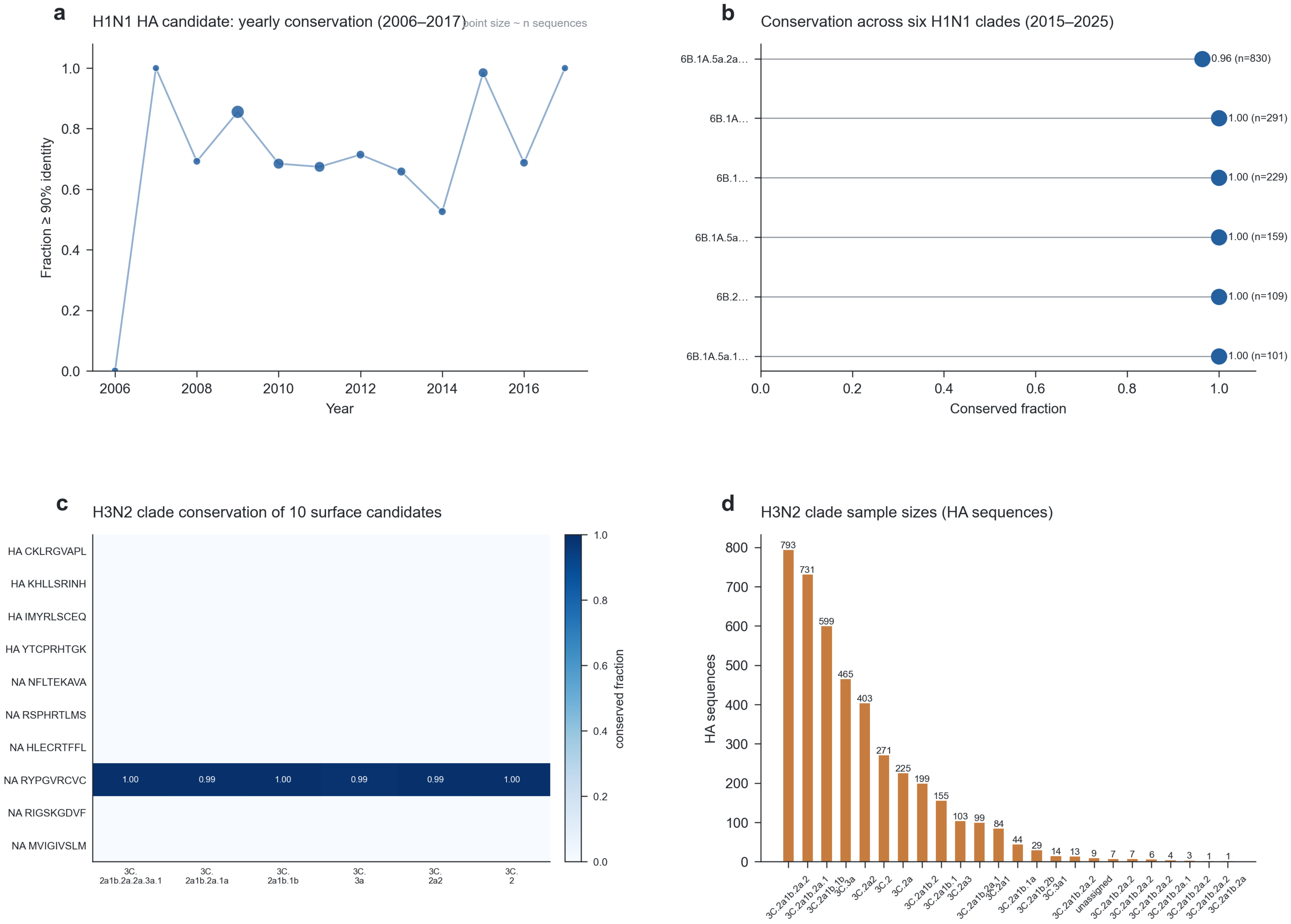

**Figure S3 | Model and external benchmark validation**

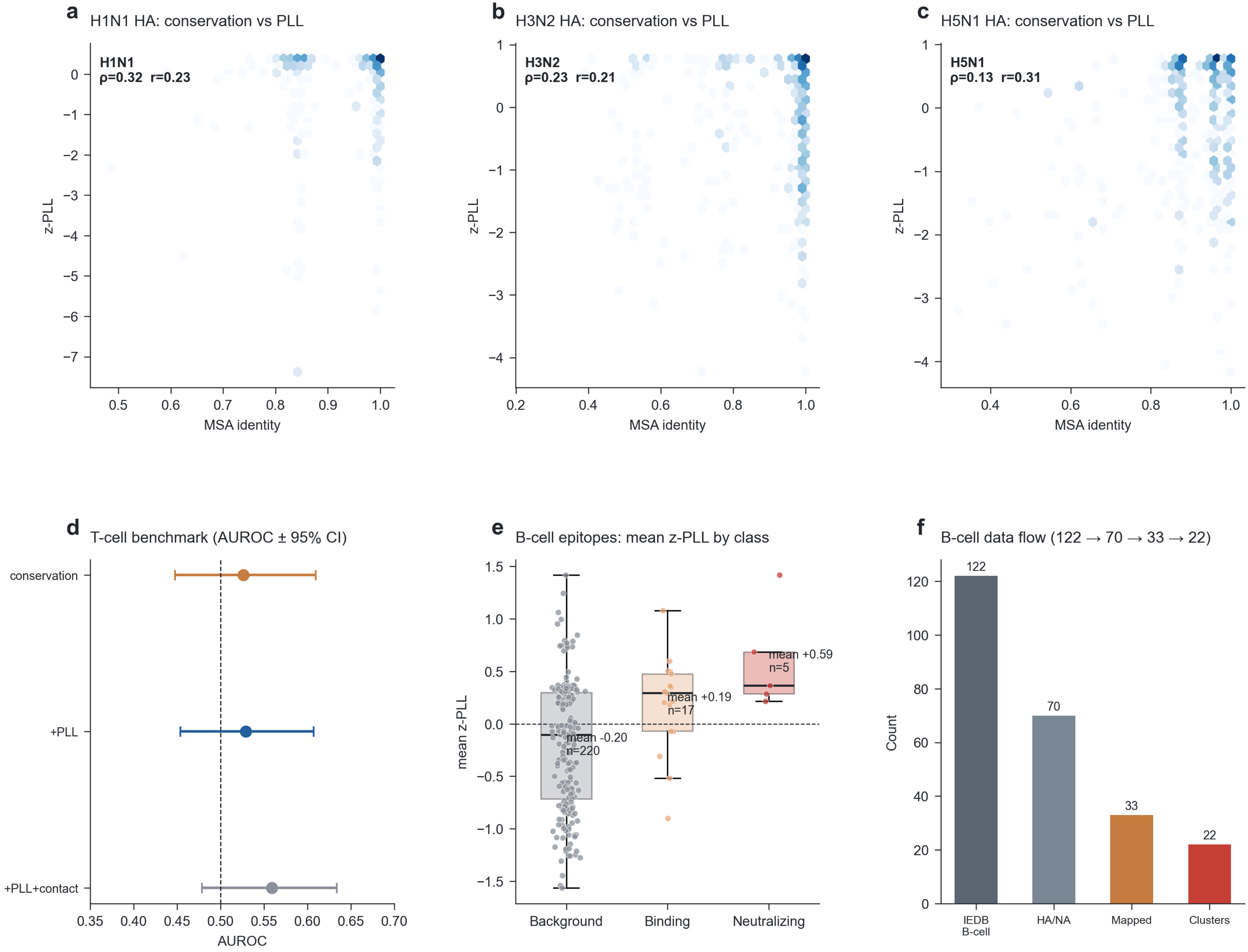
