## Supplementary material for "Conserved influenza A epitope candidate regions and a benchmark of ESM-2 sequence features": Li_et_al_Supplementary_Data: 00_BENCHMARK_AUDIT.md

# B-cell benchmark 数据审计 — 为什么之前 n=4

生成日期：2026-08-13

## 问题

之前的 IEDB benchmark 只定位到 n=4 个 B-cell 表位，导致 B-cell benchmark 无法做。

## 根因（结论：B — 检索/过滤丢失了大量记录，非数据缺乏）

### 1. EBI `epitope` 端点只返回线性表位

之前用 EBI Proteins API 的 `/epitope` 端点（`09_iedb_enrichment.py`）。实测：
- P03452（H1N1 HA）返回 22 个 epitope，**全部**描述为 "linear peptidic epitope"。
- **0 个构象/不连续表位**（antibody 表位大多是构象性的，被完全漏掉）。

### 2. 分类过严

之前的分类逻辑：描述含 "B cell" 且不含 "T cell" → B。这排除了大量"B+T 混合"的 antibody 表位。

### 3. 部分 UniProt accession 已废弃

P03474（NA）等 accession 返回 0 个 epitope（已废弃）。

### 4. IEDB query-api 之前被网络重置

早期 `query-api.iedb.org` 连接被重置（无法直查 IEDB），只能走 EBI 线性端点。**现在已可达（HTTP 200）**。

## 修复后：直接查 IEDB

用 `query-api.iedb.org/epitope_search` + `bcell_ids=not.is.null` 过滤：

| 指标 | 数量 |
|---|---|
| 流感 A 总 B-cell 表位 | **122** |
| 其中 HA/NA（表面糖蛋白） | **70** |
| 构象（不连续）表位 | 13 |

## 结论

之前 n=4 是**检索程序丢失**（线性-only 端点 + 过严分类 + API 被阻），不是 IEDB 数据缺乏。修复后获得 70 个 HA/NA B-cell 表位，足以做 benchmark。

## 数据来源与可审计性

- 来源：IEDB（Immune Epitope Database）query API，`query-api.iedb.org`。
- 检索日期：2026-08-13。
- 过滤条件：`source_organism_names=cs.{Influenza A virus}` + `bcell_ids=not.is.null`。
- 每条保留 IEDB ID、PMID、protein、position、sequence、evidence type、assay names、isotype。
- 原始 JSON：`/tmp/iedb_bcell_raw.json`（122 条完整记录）。
