## Supplementary material for "Conserved influenza A epitope candidate regions and a benchmark of ESM-2 sequence features": Li_et_al_Supplementary_Data: 02_DATASET_SUMMARY.md

# B-cell / neutralizing-antibody benchmark 数据集摘要

生成日期：2026-08-13
来源：IEDB query API（query-api.iedb.org），过滤 `source_organism_names=cs.{Influenza A virus}` + `bcell_ids=not.is.null`。

## 数据量（诚实统计）

| 指标 | 数量 |
|---|---|
| 流感 A 总 B-cell 表位 | 122 |
| **HA/NA 表面糖蛋白表位** | **70**（HA 64 + NA 6） |
| — neutralizing（真病毒中和 assay） | **16** |
| — antibody-binding（结合 assay） | 51 |
| — B-cell-other | 3 |
| 线性表位（peptide-level 可用） | 58 |
| 构象/不连续表位（需 residue-level） | 12 |
| **独立文献（PMID）总数** | **154** |
| 独立文献（中和表位） | 27 |

## 结构类型分布

| structure_type | 数量 | 说明 |
|---|---|---|
| Linear peptide | 58 | 有连续序列 → peptide-level benchmark |
| Discontinuous peptide | 7 | 仅残基位置 → residue-level benchmark |
| Discontinuous peptide on multi chain | 5 | 多链构象表位 → residue-level |

## 中和表位结构分布

- 9 个线性（有序列）。
- 7 个构象/多链（仅残基位置）。

## 未完成项（下一步）

1. **亚型映射**：数据里 `source_organism` 只是 "Influenza A virus"，具体 H1/H3/H5 需从抗原 accession/株名映射到我的共识序列。
2. **位置标准化**：`position` 字段是 IEDB 的抗原坐标（不同株、不同编号体系），需映射到我的 consensus 坐标。
3. **residue-level 表位**：构象表位的 `structure_descriptions` 是残基列表（非连续肽），需转换为 residue-level positive。
4. **negative 采样**：需对 HA/NA 做 matched negative（同蛋白、同长度/结构区、同保守度、非已知表位）。

## 关键结论

**之前 n=4 是检索程序丢失，不是数据缺乏。** IEDB 实际有 70 个 HA/NA B-cell 表位（16 个中和），足以做 peptide-level（58 个线性）和 residue-level（12 个构象）benchmark。

数据文件：`01_BCELL_DATASET.tsv`（70 条，含 IEDB ID / PMID / protein / position / sequence / evidence / assay names / isotype / reference titles）。
