## Supplementary material for "Conserved influenza A epitope candidate regions and a benchmark of ESM-2 sequence features": Li_et_al_Supplementary_Data: 03_subtype_mapping.py

```
#!/usr/bin/env python3
"""
Subtype/strain mapping for the 70 HA/NA B-cell epitopes.

Rules (fixed before any performance look):
  1. Parse subtype HxNy from the source-organism / antigen-name fields.
  2. If exactly one subtype across all antigens -> assign, evidence="strain_name", confidence=high.
  3. If multiple subtypes -> "multiple", confidence=low.
  4. If none -> map linear_sequence to the project consensus (H1N1/H3N2/H5N1 HA+NA);
     best-match subtype -> assign, evidence="sequence_mapping".
  5. Still unresolved -> "unresolved" (do NOT force-assign).
"""
import json, re, csv
from Bio import SeqIO

BASE = "/media/qxl/My_Passport/流行病学数据/广谱流感疫苗候选课题"
CONS = f"{BASE}/08_ESM2_AI分析与论文/data/consensus"
OUT = f"{BASE}/07_MANUSCRIPT_STRENGTHENING/08_BCELL_EXTERNAL_BENCHMARK"

rows = json.load(open("/tmp/iedb_bcell_raw.json"))

def is_ha_na(r):
    return any(("emagglutinin" in a.get("name","").lower()) or
               ("euraminidase" in a.get("name","").lower())
               for a in r.get("curated_source_antigens", []))

def protein_of(r):
    for a in r.get("curated_source_antigens", []):
        n = a.get("name","").lower()
        if "emagglutinin" in n: return "HA"
        if "euraminidase" in n: return "NA"
    return "other"

def parse_subtypes(r):
    subs = set()
    for a in r.get("curated_source_antigens", []):
        org = a.get("source_organism_name","") or ""
        name = a.get("name","") or ""
        for m in re.findall(r"H(\d)N(\d)", org + " " + name):
            subs.add(f"H{m[0]}N{m[1]}")
    return subs

# load consensus
cons = {}
for sub in ["H1N1","H3N2","H5N1"]:
    for prot in ["HA","NA"]:
        p = f"{CONS}/{sub}_{prot}.fasta"
        cons[(sub,prot)] = str(list(SeqIO.parse(p,"fasta"))[0].seq)

def seq_best_subtype(seq, prot):
    if not seq: return None, 0.0
    best = (None, 0.0)
    for sub in ["H1N1","H3N2","H5N1"]:
        s = cons[(sub,prot)]
        for i in range(len(s)-len(seq)+1):
            ident = sum(a==b for a,b in zip(seq, s[i:i+len(seq)]))/len(seq)
            if ident > best[1]: best = (sub, ident)
    return best

ha_na = [r for r in rows if is_ha_na(r)]
out = []
for r in ha_na:
    prot = protein_of(r)
    seq = r.get("linear_sequence") or ""
    subs = parse_subtypes(r)
    accessions = ";".join(sorted(set(a.get("accession","") for a in r.get("curated_source_antigens",[]))))
    strains = ";".join(sorted(set(a.get("source_organism_name","") for a in r.get("curated_source_antigens",[]))))
    if len(subs) == 1:
        subtype = list(subs)[0]; ev = "strain_name"; conf = "high"
    elif len(subs) > 1:
        subtype = ";".join(sorted(subs)); ev = "strain_name_conflict"; conf = "low"
    else:
        st, ident = seq_best_subtype(seq, prot)
        if st and ident >= 0.85:
            subtype = st; ev = "sequence_mapping"; conf = "medium"
        else:
            subtype = "unresolved"; ev = "none"; conf = "low"
    out.append(dict(
        iedb_id=r.get("structure_id"), protein=prot,
        epitope_sequence=seq, source_accession=accessions, strain=strains[:120],
        subtype=subtype, mapping_evidence=ev, mapping_confidence=conf,
        reference_pmid=";".join(r.get("pubmed_ids") or [])))

with open(f"{OUT}/03_EPITOPE_SUBTYPE_MAPPING.tsv","w",newline="") as f:
    w = csv.DictWriter(f, fieldnames=list(out[0].keys()), delimiter="\t")
    w.writeheader(); w.writerows(out)

from collections import Counter
print(f"共 {len(out)} 条 HA/NA 表位")
print("subtype:", dict(Counter(r["subtype"] for r in out)))
print("evidence:", dict(Counter(r["mapping_evidence"] for r in out)))
print("confidence:", dict(Counter(r["mapping_confidence"] for r in out)))
print(f"saved {OUT}/03_EPITOPE_SUBTYPE_MAPPING.tsv")
```
