## Supplementary material for "Conserved influenza A epitope candidate regions and a benchmark of ESM-2 sequence features": Li_et_al_Supplementary_Data: 04_consensus_coordinate_mapping.py

```
#!/usr/bin/env python3
"""
Consensus-coordinate mapping for linear HA/NA B-cell epitopes.

Rules (fixed before any performance look):
  - For each linear peptide of length L, map to the assigned subtype's consensus.
  - unique exact match (identity 1.0)          -> PASS
  - exact match at >=2 positions                -> AMBIGUOUS
  - 1-mismatch match                            -> AMBIGUOUS
  - >=2 mismatches or no match (best < 0.9)     -> FAIL
  - subtype unresolved / H2N2 (no consensus)    -> FAIL (cannot map)
"""
import json, csv
from Bio import SeqIO

BASE = "/media/qxl/My_Passport/流行病学数据/广谱流感疫苗候选课题"
CONS = f"{BASE}/08_ESM2_AI分析与论文/data/consensus"
OUT = f"{BASE}/07_MANUSCRIPT_STRENGTHENING/08_BCELL_EXTERNAL_BENCHMARK"

submap = list(csv.DictReader(open(f"{OUT}/03_EPITOPE_SUBTYPE_MAPPING.tsv"), delimiter="\t"))
cons = {}
for sub in ["H1N1","H3N2","H5N1"]:
    for prot in ["HA","NA"]:
        cons[(sub,prot)] = str(list(SeqIO.parse(f"{CONS}/{sub}_{prot}.fasta","fasta"))[0].seq)

def map_linear(seq, sub, prot):
    """Return (status, start, end, identity, n_positions)."""
    if sub not in ("H1N1","H3N2","H5N1"):
        return "FAIL", None, None, 0.0, 0, "no_consensus"
    s = cons[(sub,prot)]
    L = len(seq)
    exact_positions = []
    best = (None, 0.0)
    for i in range(len(s)-L+1):
        mm = sum(a!=b for a,b in zip(seq, s[i:i+L]))
        ident = 1 - mm/L
        if mm == 0:
            exact_positions.append(i)
        if ident > best[1]:
            best = (i, ident)
    if len(exact_positions) == 1:
        i = exact_positions[0]
        return "PASS", i+1, i+L, 1.0, len(exact_positions), "exact"
    elif len(exact_positions) > 1:
        i = exact_positions[0]
        return "AMBIGUOUS", i+1, i+L, 1.0, len(exact_positions), "multiple_exact"
    # no exact -> check 1 mismatch
    i, ident = best
    if i is None:
        return "FAIL", None, None, 0.0, 0, "no_match"
    mm = round((1-ident)*L)
    if mm == 1:
        return "AMBIGUOUS", i+1, i+L, ident, 1, "1_mismatch"
    else:
        return "FAIL", i+1, i+L, ident, 0, f"{mm}_mismatch"

out = []
for r in submap:
    prot = r["protein"]; sub = r["subtype"]; seq = r["epitope_sequence"]
    if not seq:
        status, st, en, ident, np_, note = "FAIL", None, None, 0.0, 0, "no_sequence"
    else:
        status, st, en, ident, np_, note = map_linear(seq, sub, prot)
    out.append(dict(
        iedb_id=r["iedb_id"], protein=prot, subtype=sub, epitope_sequence=seq,
        mapped_start=st if st else "", mapped_end=en if en else "",
        identity=round(ident,3), n_mapping_positions=np_,
        mapping_status=status, mapping_note=note))

with open(f"{OUT}/04_CONSENSUS_COORDINATE_MAPPING.tsv","w",newline="") as f:
    w = csv.DictWriter(f, fieldnames=list(out[0].keys()), delimiter="\t")
    w.writeheader(); w.writerows(out)

from collections import Counter
print(f"共 {len(out)} 条")
print("mapping_status:", dict(Counter(r["mapping_status"] for r in out)))
print("mapping_note:", dict(Counter(r["mapping_note"] for r in out)))
print(f"saved {OUT}/04_CONSENSUS_COORDINATE_MAPPING.tsv")
```
