## Supplementary material for "Conserved influenza A epitope candidate regions and a benchmark of ESM-2 sequence features": Li_et_al_Supplementary_Data: 05_nonredundant_clusters.py

```
#!/usr/bin/env python3
"""
Remove pseudoreplication: cluster highly-overlapping epitopes into non-redundant
epitope clusters (union-find on consensus-coordinate overlap >= 50%).

Rule (fixed): within same protein + subtype, two epitopes belong to the same
cluster if their mapped consensus residue intervals overlap by >=50% of the
shorter interval.
"""
import csv
from collections import defaultdict

BASE = "/media/qxl/My_Passport/流行病学数据/广谱流感疫苗候选课题"
OUT = f"{BASE}/07_MANUSCRIPT_STRENGTHENING/08_BCELL_EXTERNAL_BENCHMARK"

submap = list(csv.DictReader(open(f"{OUT}/03_EPITOPE_SUBTYPE_MAPPING.tsv"), delimiter="\t"))
coord = list(csv.DictReader(open(f"{OUT}/04_CONSENSUS_COORDINATE_MAPPING.tsv"), delimiter="\t"))
coord_map = {r["iedb_id"]: r for r in coord}

# keep mapped linear epitopes (PASS/AMBIGUOUS) only
rows = []
for r in submap:
    c = coord_map.get(r["iedb_id"])
    if c and c["mapping_status"] in ("PASS", "AMBIGUOUS"):
        rows.append(dict(
            iedb_id=r["iedb_id"], protein=r["protein"], subtype=r["subtype"],
            seq=r["epitope_sequence"], start=int(c["mapped_start"]), end=int(c["mapped_end"]),
            evidence=r.get("mapping_evidence",""), ref_pmid=r["reference_pmid"]))

# union-find on overlap
parent = {i: i for i in range(len(rows))}
def find(x):
    while parent[x] != x:
        parent[x] = parent[parent[x]]; x = parent[x]
    return x
def union(a, b):
    ra, rb = find(a), find(b)
    if ra != rb: parent[ra] = rb

for i in range(len(rows)):
    for j in range(i+1, len(rows)):
        a, b = rows[i], rows[j]
        if a["protein"] != b["protein"] or a["subtype"] != b["subtype"]:
            continue
        # residue sets
        sa = set(range(a["start"], a["end"]+1))
        sb = set(range(b["start"], b["end"]+1))
        inter = sa & sb
        if inter:
            frac = len(inter) / min(len(sa), len(sb))
            if frac >= 0.5:
                union(i, j)

groups = defaultdict(list)
for i in range(len(rows)):
    groups[find(i)].append(i)

clusters = []
for g in groups.values():
    members = [rows[i] for i in g]
    # representative = longest seq
    rep = max(members, key=lambda m: len(m["seq"]))
    pmids = set()
    has_binding = False; has_neut = False
    for m in members:
        for p in m["ref_pmid"].split(";"):
            if p: pmids.add(p)
        # evidence type from raw (use subtype mapping evidence + a placeholder; actual evidence from 01)
        # NOTE: bind/neut evidence will be joined from 01_BCELL_DATASET in QC step
    clusters.append(dict(
        cluster_id=f"E{len(clusters)+1:03d}",
        protein=rep["protein"], subtype=rep["subtype"],
        representative_seq=rep["seq"],
        consensus_start=rep["start"], consensus_end=rep["end"],
        n_IEDB_records=len(members),
        n_PMIDs=len(pmids),
        member_iedb_ids=";".join(str(m["iedb_id"]) for m in members),
        member_sequences=";".join(m["seq"] for m in members),
        pmid_list=";".join(sorted(pmids))))

clusters.sort(key=lambda c: -c["n_IEDB_records"])
with open(f"{OUT}/05_NONREDUNDANT_EPITOPE_CLUSTERS.tsv","w",newline="") as f:
    w = csv.DictWriter(f, fieldnames=list(clusters[0].keys()), delimiter="\t")
    w.writeheader(); w.writerows(clusters)

print(f"原始可映射表位: {len(rows)}")
print(f"非冗余 epitope cluster: {len(clusters)}")
print("protein:", dict(__import__('collections').Counter(c['protein'] for c in clusters)))
print("subtype:", dict(__import__('collections').Counter(c['subtype'] for c in clusters)))
print("n_records 分布:", dict(__import__('collections').Counter(c['n_IEDB_records'] for c in clusters)))
print(f"saved {OUT}/05_NONREDUNDANT_EPITOPE_CLUSTERS.tsv")
```
