## Supplementary material for "Conserved influenza A epitope candidate regions and a benchmark of ESM-2 sequence features": Li_et_al_Supplementary_Data: 06_benchmark_ready.py

```
#!/usr/bin/env python3
"""
Join evidence (binding / neutralization) to non-redundant clusters, assign
hierarchical labels (Level 1/2/3), and emit the benchmark-ready dataset.

Labels:
  Level 1 = all experimentally supported antibody epitopes (all clusters)
  Level 2 = antibody-binding supported (has_binding_evidence)
  Level 3 = neutralization-supported (has_neutralization_evidence)
"""
import csv

BASE = "/media/qxl/My_Passport/流行病学数据/广谱流感疫苗候选课题"
OUT = f"{BASE}/07_MANUSCRIPT_STRENGTHENING/08_BCELL_EXTERNAL_BENCHMARK"

# evidence per IEDB record (from 01)
ev = {}
for r in csv.DictReader(open(f"{OUT}/01_BCELL_DATASET.tsv"), delimiter="\t"):
    ev[r["iedb_id"]] = r["evidence"]

clusters = list(csv.DictReader(open(f"{OUT}/05_NONREDUNDANT_EPITOPE_CLUSTERS.tsv"), delimiter="\t"))

for c in clusters:
    ids = c["member_iedb_ids"].split(";")
    evidences = [ev.get(i, "unknown") for i in ids]
    c["has_binding_evidence"] = any(e in ("antibody-binding", "neutralization") for e in evidences)
    c["has_neutralization_evidence"] = any(e == "neutralization" for e in evidences)
    c["evidence_breakdown"] = ";".join(sorted(set(evidences)))
    # label
    if c["has_neutralization_evidence"]:
        c["label_level"] = "L3_neutralization"
    elif c["has_binding_evidence"]:
        c["label_level"] = "L2_binding"
    else:
        c["label_level"] = "L1_antibody"

with open(f"{OUT}/06_BENCHMARK_READY_DATASET.tsv","w",newline="") as f:
    w = csv.DictWriter(f, fieldnames=list(clusters[0].keys()), delimiter="\t")
    w.writeheader(); w.writerows(clusters)

from collections import Counter
print(f"非冗余 cluster 数: {len(clusters)}")
print("label_level:", dict(Counter(c["label_level"] for c in clusters)))
print("has_neutralization:", sum(1 for c in clusters if c["has_neutralization_evidence"]=="True"))
print("has_binding:", sum(1 for c in clusters if c["has_binding_evidence"]=="True"))
print(f"saved {OUT}/06_BENCHMARK_READY_DATASET.tsv")
```
