## Supplementary material for "Conserved influenza A epitope candidate regions and a benchmark of ESM-2 sequence features": Li_et_al_Supplementary_Data: 07_matched_background.py

```
#!/usr/bin/env python3
"""
Matched background controls (pseudo-negatives), NOT true negatives.

For each positive epitope cluster, sample background peptides from the same
protein + subtype, matching on:
  - length (within +-20%)
  - conservation (positional identity within +-0.05)
while EXCLUDING any window overlapping a known positive region + buffer.

Buffer = 10 residues on each side of every positive (rationale: adjacent
residues are structurally coupled and not independent background).
10 background peptides per positive (sampled; does NOT inflate statistical n).
"""
import csv, os
from collections import Counter
import numpy as np
from Bio import SeqIO

BASE = "/media/qxl/My_Passport/流行病学数据/广谱流感疫苗候选课题"
RAW = f"{BASE}/01_原始序列与分层数据/序列与地区分层/全球/序列/综合"
CONS = f"{BASE}/08_ESM2_AI分析与论文/data/consensus"
OUT = f"{BASE}/07_MANUSCRIPT_STRENGTHENING/08_BCELL_EXTERNAL_BENCHMARK"
RNG = np.random.default_rng(42)
BUFFER = 10
N_BG_PER_POS = 10

def mode_seqs(sub, prot):
    p = f"{RAW}/分类序列_{sub}/{prot}_sequences.fasta"
    recs = list(SeqIO.parse(p, "fasta"))
    aa = [str(r.seq[:(len(r.seq)//3)*3].translate()) for r in recs]
    ln = Counter(len(s) for s in aa).most_common(1)[0][0]
    return [s for s in aa if len(s) == ln and s.count("X") <= 0.05*len(s)]

def cons_identity(sub, prot):
    seqs = mode_seqs(sub, prot)
    L = len(seqs[0])
    cons = [Counter(s[i] for s in seqs).most_common(1)[0][0] for i in range(L)]
    return np.array([sum(1 for s in seqs if s[i]==cons[i])/len(seqs) for i in range(L)])

cons_seq = {}
for sub in ["H1N1","H3N2","H5N1"]:
    for prot in ["HA","NA"]:
        cons_seq[(sub,prot)] = str(list(SeqIO.parse(f"{CONS}/{sub}_{prot}.fasta","fasta"))[0].seq)

clusters = list(csv.DictReader(open(f"{OUT}/06_BENCHMARK_READY_DATASET.tsv"), delimiter="\t"))

# build exclusion mask per (subtype, protein): all positive regions + buffer
excl = {}
for sub in ["H1N1","H3N2","H5N1"]:
    for prot in ["HA","NA"]:
        excl[(sub,prot)] = set()
for c in clusters:
    key = (c["subtype"], c["protein"])
    s, e = int(c["consensus_start"]), int(c["consensus_end"])
    L = len(cons_seq.get(key, ""))
    for p in range(max(0, s-BUFFER-1), min(L, e+BUFFER)):
        excl[key].add(p)

out = []
for c in clusters:
    key = (c["subtype"], c["protein"])
    s, e = int(c["consensus_start"]), int(c["consensus_end"])
    L = len(cons_seq[key])
    plen = e - s + 1
    ci = cons_identity(c["subtype"], c["protein"])
    pos_cons = np.mean(ci[s-1:e])
    # scan candidate windows of length ~plen
    candidates = []
    for i in range(0, L - plen + 1):
        # exclusion: any position in the window in excl set
        if any(p in excl[key] for p in range(i, i+plen)):
            continue
        wcons = np.mean(ci[i:i+plen])
        if abs(wcons - pos_cons) <= 0.05:
            candidates.append((i, wcons))
    # sample N_BG_PER_POS
    if candidates:
        sample = RNG.choice(len(candidates), size=min(N_BG_PER_POS, len(candidates)), replace=False)
        for k in sample:
            i, wcons = candidates[k]
            out.append(dict(
                cluster_id=c["cluster_id"], protein=c["protein"], subtype=c["subtype"],
                positive_seq=c["representative_seq"],
                bg_start=i+1, bg_end=i+plen,
                bg_sequence=cons_seq[key][i:i+plen],
                bg_conservation=round(wcons,3),
                positive_conservation=round(pos_cons,3),
                positive_label=c["label_level"]))

with open(f"{OUT}/07_MATCHED_BACKGROUND.tsv","w",newline="") as f:
    w = csv.DictWriter(f, fieldnames=list(out[0].keys()), delimiter="\t")
    w.writeheader(); w.writerows(out)

print(f"生成 {len(out)} 个 matched background peptides")
print(f"覆盖 {len(set(r['cluster_id'] for r in out))} 个 positive clusters")
bg_per = Counter(r['cluster_id'] for r in out)
print("每个 positive 的背景数分布:", dict(Counter(bg_per.values())))
print(f"saved {OUT}/07_MATCHED_BACKGROUND.tsv")
```
