## Supplementary material for "Conserved influenza A epitope candidate regions and a benchmark of ESM-2 sequence features": Li_et_al_Supplementary_Data: 08_BENCHMARK_QC_REPORT.md

# B-cell benchmark QC 报告

生成日期：2026-08-13
原则：所有 inclusion/exclusion/matching 规则在查看任何 PLL 性能前固定；不伪造；不调规则。

---

## 数据流水线追踪（70 → 22）

| 步骤 | 数量 |
|---|---|
| IEDB 流感 A B-cell 表位 | 122 |
| 其中 HA/NA（表面糖蛋白） | **70**（58 linear + 12 discontinuous） |
| 亚型映射成功（H1N1/H3N2/H5N1） | 43（H1N1=20, H3N2=23） |
| 亚型无法判定（H2N2 / unresolved / multiple） | 27（H2N2=3, unresolved=20, multiple=4） |
| 共识坐标 PASS（exact unique） | 26 |
| 共识坐标 AMBIGUOUS（1-mismatch） | 7 |
| 共识坐标 FAIL | 37（12 构象无序列 + 15 无共识 + 10 序列漂移） |
| **可映射线性表位** | **33** |
| **非冗余 epitope cluster（去伪重复后）** | **22** |

## 非冗余 cluster 分布

| 维度 | 分布 |
|---|---|
| protein | HA 18 / NA 4 |
| subtype | H1N1 12 / H3N2 10 |
| label | L3_neutralization **5** / L2_binding 17 |
| n_records | 17 个 singleton + 5 个 cluster（含 2–5 条记录） |

## 独立文献

| 指标 | 数量 |
|---|---|
| 独立 PMID（全部 70） | 154 |
| 独立 PMID（中和表位） | 27 |
| 独立 PMID（22 cluster） | 83 |

## 排除/合并记录（透明）

- **12 条 discontinuous**（构象表位）：无连续序列，转入 residue-level 轨道（未在本阶段处理）。
- **15 条无共识**：H2N2（3，项目无 H2N2 共识）+ unresolved（20，序列映射 <85% identity）+ multiple（4，多亚型冲突）。
- **10 条序列漂移**：有亚型但序列与 consensus ≥2 错配（不放松映射）。
- **11 条合并**：33 可映射 → 22 cluster（同蛋白同亚型、共识坐标重叠 ≥50% 的合并）。

## 最终 independent units

**22 个非冗余 epitope cluster**（不是 70，也不是 33）。后续 bootstrap / permutation 必须以 cluster 为 resampling unit，不能以单条 peptide record 为单位。

---

## 统计功效判断（诚实）

**A. peptide-level benchmark（22 cluster）**
- 22 个 positive（18 HA + 4 NA）。
- 220 个 matched background（10/positive）。
- **功效不足**：22 个 positive 对稳定 AUROC/AUPRC 而言偏少（通常需 ≥30–50）。可做描述性 / 效应量分析，但 AUROC 的置信区间会较宽。**不应把"10 background/positive"当作样本量扩大 10 倍。**

**B. residue-level benchmark（12 discontinuous epitope）**
- 12 个构象表位，尚未做 residue 映射（需源抗原序列 + 逐残基转换）。
- **功效严重不足**：即使映射完成，12 个 epitope 不足以做稳健统计。

**C. neutralization-only benchmark（5 cluster）**
- 仅 5 个中和 cluster。
- **功效严重不足**：无法单独做 AUROC。只能作为描述性案例，与 binding（17）分开报告，不得合并。

---

## 结论

1. 数据集从 n=4 提升到 **22 个非冗余 HA/NA B-cell epitope cluster（含 5 个中和 + 17 个结合）**，可审计、有 PMID、有亚型/坐标映射。
2. **但即使是 22 个 cluster，对正式 AUROC/AUPRC 仍偏少**，尤其是中和-only（5）和 residue-level（12）。
3. 诚实的定位应是：**"small, curated benchmark"**，报告效应量 + 置信区间 + 描述性趋势，而非声称高功效统计检验。
4. 所有规则（亚型映射、坐标 PASS/AMBIGUOUS/FAIL、聚类 50% 重叠、background 匹配 ±0.05 保守度、buffer=10、10 bg/positive）已在查看 PLL 前固定。

## 数据文件

- 01_BCELL_DATASET.tsv（70 原始 HA/NA 表位）
- 03_EPITOPE_SUBTYPE_MAPPING.tsv（亚型/株系映射）
- 04_CONSENSUS_COORDINATE_MAPPING.tsv（坐标映射 PASS/AMBIGUOUS/FAIL）
- 05_NONREDUNDANT_EPITOPE_CLUSTERS.tsv（22 cluster）
- 06_BENCHMARK_READY_DATASET.tsv（含 label + 证据）
- 07_MATCHED_BACKGROUND.tsv（220 matched background）
