## Supplementary material for "Conserved influenza A epitope candidate regions and a benchmark of ESM-2 sequence features": Li_et_al_Supplementary_Data: 09_neutralizing_footprints.py

```
#!/usr/bin/env python3
"""
Extract neutralizing-antibody contact residues from antibody-HA complexes.
HA is cleaved into HA1 (chain A) + HA2 (chain B); stalk antibodies bind HA2.
Contact = antigen residue with any atom within 4.5 A of antibody (H/L).
Map each chain (HA1, HA2) to the project consensus separately.
"""
import numpy as np
from Bio.PDB import PDBParser, NeighborSearch
from Bio import SeqIO
from Bio.Align import PairwiseAligner

BASE = "/media/qxl/My_Passport/流行病学数据/广谱流感疫苗候选课题"
CONS = f"{BASE}/08_ESM2_AI分析与论文/data/consensus"
OUT = f"{BASE}/07_MANUSCRIPT_STRENGTHENING/08_BCELL_EXTERNAL_BENCHMARK/structures"

T3 = {'ALA':'A','ARG':'R','ASN':'N','ASP':'D','CYS':'C','GLN':'Q','GLU':'E','GLY':'G','HIS':'H','ILE':'I','LEU':'L','LYS':'K','MET':'M','PHE':'F','PRO':'P','SER':'S','THR':'T','TRP':'W','TYR':'Y','VAL':'V'}

structures = [
    ("3GBN", "CR6261", "H1N1", "stalk"),
    ("3SDY", "CR8020", "H3N2", "stalk"),
    ("4FQI", "CR9114", "H5N1", "stalk"),
]

cons = {}
for sub in ["H1N1","H3N2","H5N1"]:
    cons[sub] = str(list(SeqIO.parse(f"{CONS}/{sub}_HA.fasta","fasta"))[0].seq)

aligner = PairwiseAligner(); aligner.mode='local'; aligner.match_score=2
aligner.mismatch_score=-1; aligner.open_gap_score=-2; aligner.extend_gap_score=-1

def chain_seq_resnums(chain):
    resnums=[]; seq=[]
    for res in chain:
        if res.id[0]!=' ': continue
        aa=T3.get(res.resname.strip())
        if aa is None: continue
        resnums.append(res.id[1]); seq.append(aa)
    return "".join(seq), resnums

def map_to_cons(chain_seq, resnums, contact, cons_seq):
    a = aligner.align(cons_seq, chain_seq)[0]
    target_blocks, query_blocks = a.aligned
    q2t = {}
    for (ts, te), (qs, qe) in zip(target_blocks, query_blocks):
        for off in range(te - ts):
            q2t[qs + off] = ts + off
    cons_contact=[]
    for ri, rn in enumerate(resnums):
        if rn in contact and ri in q2t:
            cons_contact.append(q2t[ri]+1)  # 1-indexed consensus position
    return sorted(set(cons_contact))

out=[]
for pdb, ab, sub, region in structures:
    parser=PDBParser(QUIET=True)
    s=parser.get_structure(pdb, f"{OUT}/{pdb}.pdb")[0]
    ab_atoms=[]
    for ch in ["H","L"]:
        if ch in s: ab_atoms.extend(list(s[ch].get_atoms()))
    ns=NeighborSearch(ab_atoms)
    all_contact=[]
    # HA1 (chain A) and HA2 (chain B)
    for ha_chain in ["A","B"]:
        if ha_chain not in s: continue
        chain=s[ha_chain]
        contact=set()
        for a in chain.get_atoms():
            if ns.search(a.coord, 4.5):
                contact.add(a.get_parent().id[1])
        seq, resnums = chain_seq_resnums(chain)
        cc = map_to_cons(seq, resnums, contact, cons[sub])
        all_contact.extend(cc)
    all_contact=sorted(set(all_contact))
    out.append(dict(structure=pdb, antibody=ab, subtype=sub, protein="HA", region=region,
                    n_contact_residues=len(all_contact),
                    consensus_positions=";".join(map(str, all_contact)),
                    source=f"PDB:{pdb}"))

with open(f"{OUT}/09_NEUTRALIZING_FOOTPRINTS.tsv","w",newline="") as f:
    import csv
    w=csv.DictWriter(f, fieldnames=list(out[0].keys()), delimiter="\t")
    w.writeheader(); w.writerows(out)

for r in out:
    print(f"{r['antibody']:8s} {r['subtype']:5s} {r['region']:6s} 接触残基={r['n_contact_residues']:3d}  共识位置={r['consensus_positions'][:70]}")
print(f"saved {OUT}/09_NEUTRALIZING_FOOTPRINTS.tsv")
```
