## Supplementary material for "Conserved influenza A epitope candidate regions and a benchmark of ESM-2 sequence features": Li_et_al_Supplementary_Data: 10_MANUSCRIPT_OUTLINE.md

# 新框架 manuscript 大纲（可投稿定位）

定位：**benchmark + 诚实负面结果 + 方法学警示**，目标 Bioinformatics Advances / BMC Bioinformatics。

---

## 标题（建议）

**Benchmarking the utility and limitations of protein-language-model sequence features for influenza A epitope prioritization**

（备选）*Does ESM-2 add value over alignment conservation for influenza epitope prioritization? A curated, non-redundant benchmark*

---

## 摘要（核心信息）

1. 我们构建了一个**可审计、非冗余**的甲流表位优先排序 benchmark：70 个 HA/NA B-cell 表位 → 22 个非冗余 cluster（5 中和 + 17 结合），103 个 T-cell 表位，全部映射到共识坐标，去伪重复、matched background。
2. 评估 ESM-2 零样本特征（group-masked PLL、attention contact-density）是否在比对保守性之外提供增量。
3. **发现**：
   - PLL 与 MSA 保守度**中等相关**（ρ=0.25–0.39），本质上是"保守性代理"。
   - PLL 对 **T-cell 表位识别无增量**（ΔAUROC +0.004, p=0.455，接近随机）。
   - **contact-density 不是可及性代理**（HA ρ=+0.147 / NA ρ=−0.132，方向不一致）。
   - B-cell/中和 benchmark **功效不足**（22 cluster，5 中和），无法高功效检验。
4. 附加方法学警示：GISAID 序列异质、IEDB 线性端点漏构象表位、坐标映射、伪重复。
5. **结论**：对保守性依赖的表位任务，**alignment conservation 已经足够，ESM-2 似然未提供可测的增量**。

---

## 引言（两段）

- 段1：广谱流感疫苗需要保守表位；传统流程用 MSA 保守性 + B/T 表位预测。
- 段2：蛋白质语言模型（ESM-2）被寄予厚望，声称零样本似然能捕捉"进化约束 + 结构信息"。**问题**：这些特征是否真的在传统保守性之外提供增量？本文用严格的 benchmark 回答。

---

## 结果（6 小节）

### 1. 一个可审计、非冗余的 benchmark（数据集本身是贡献）
- IEDB 检索 → 70 HA/NA B-cell 表位（含构象表位）→ 亚型/株系映射 → 共识坐标映射（PASS/AMBIGUOUS/FAIL）→ 去伪重复（22 cluster）→ matched background。
- 报告数据流水线（70→33→22）和 QC 表。

### 2. ESM-2 PLL ≈ 保守性代理（ρ=0.25–0.39）
- MSA 相关（Shannon entropy 负相关），中等、远非 1.0。
- 数据质量：原始 GISAID 序列 40–49 种长度，mode-length 过滤必要。

### 3. PLL 对 T-cell 表位无增量（统计检验）
- Model A（conservation）vs C（+PLL）vs D（+contact）：AUROC 0.526/0.529/0.559，增量 +0.004（p=0.455）。
- PLL 与 conservation 在匹配集内正交（ρ=−0.01）。

### 4. contact-density 不是可及性（RSA 验证）
- FreeSASA 真实 RSA：HA +0.147 / NA −0.132，方向不一致。
- 结论：contact-density 不应作为排序特征。

### 5. B-cell / 中和 benchmark：功效不足（诚实）
- 22 cluster（5 中和）。中和表位本质是保守区（stalk/活性区），conservation alone 即可识别，PLL 无增量空间。
- 补充：3 个 stalk 广谱中和抗体（CR6261/CR8020/CR9114）的晶体结构足迹映射到共识 HA2 stalk（残基 360–390）。

### 6. 方法学警示（对领域的贡献）
- GISAID 序列异质 → 必须 mode-length 过滤。
- IEDB 线性端点漏构象表位（n=4 是检索丢失，非数据缺乏）。
- match_count ≠ 亚型数；netMHCpan 阈值笔误；contact-density ≠ accessibility。
- 伪重复：33 表位 → 22 cluster，必须以 cluster 为 resampling unit。

---

## 讨论

- 诚实的核心结论：**对"保守性依赖型"表位任务，ESM-2 掩码语言似然 ≈ 序列保守性，未提供可测增量**。
- 这一"负面结果"的价值：防止后人重复踩坑；明确 ESM-2 用途边界（它擅长结构/功能预测，但不擅长"超越保守性的表位排序"）。
- 局限：B-cell benchmark 小样本；数据只到 2017；B-cell 工具名缺失。

---

## 方法（摘要）

- benchmark 构建：IEDB query API（过滤 + bcell_ids），亚型映射（株名 + 序列），共识坐标映射（exact/1-mismatch），聚类（50% 重叠），matched background（±0.05 保守度、buffer=10）。
- ESM-2：group-masked PLL（8 组、单点掩码敏感性 ρ=0.93）。
- MSA：MAFFT（mode-length 过滤后），positional identity + Shannon entropy。
- 结构：FreeSASA（1RU7 HA、2HTY NA）+ 抗体复合物足迹（3GBN/3SDY/4FQI）。
- 统计：AUROC/AUPRC/bootstrap，以 cluster 为 resampling unit，报告效应量 + CI，明确功效不足。

---

## 表/图清单

- **Table 1**：数据流水线（70→22，各步 N）。
- **Table 2**：22 个非冗余 cluster（蛋白/亚型/序列/坐标/证据/PMID）。
- **Table 3**：T-cell benchmark 模型对比（AUROC/CI/增量）。
- **Fig 1**：数据流水线示意图。
- **Fig 2**：PLL vs MSA 保守度散点（ρ 标注）。
- **Fig 3**：T-cell benchmark AUROC（A/C/D 对比）。
- **Fig 4**：contact-density vs 真实 RSA（HA/NA）。
- **Fig 5**：中和抗体足迹（HA2 stalk 映射）。

---

## 数据可用性

- benchmark 数据集 + 脚本（本目录，可发布到 Zenodo）。
- 不含受限 GISAID 序列（只用公开 IEDB 表位 + 共识序列）。

---

## 与旧稿的关系

- 旧稿"ESM-2 改进广谱疫苗候选优先排序"→ **废止**。
- 新稿"benchmark + 负面结果"→ 诚实、可辩护、可投稿。
- 所有旧分析（PLL、MSA、RSA、IEDB、temporal）作为支撑材料保留，但**结论改写**。
