## Supplementary material for "Conserved influenza A epitope candidate regions and a benchmark of ESM-2 sequence features": Li_et_al_Supplementary_Data: 10_neutralizing_footprints_extended.py

```
#!/usr/bin/env python3
"""
扩展中和抗体足迹：在既有 3 个 stalk 抗体（CR6261/CR8020/CR9114）基础上，
新增 4 个 RBD（HA1 球状头部）抗体 + 1 个 NA 抗体。

方法（与 09_neutralizing_footprints.py 一致）：
- 接触 = 抗原残基任一原子与抗体（重链+轻链）任一原子距离 ≤ 4.5 Å。
- HA 结构：HA1/HA2 分别用局部比对映射到全长 HA 共识；NA 结构：NA 链映射到 NA 共识。
- 每个结构用第一拷贝（HA 结构）或全部单体（CD6 二聚体表位）。

新增结构（PDB 已下载并核验链组成）：
- CH65     3SM5  H1N1  RBD  (HA1 A,C,E / HA2 B,D,F / 重 H,I,J / 轻 L,M,N)
- C05      4FP8  H3N2  RBD  (HA1 A,B,C,D / 重 H,I,J,K / 轻 L,M,N,O; 无 HA2)
- S139/1   4GMS  H3N2  RBD  (同 3SM5 结构)
- F045-092 4O58  H1N1  RBD  (HA1 A / HA2 B / 重 H / 轻 L)
- CD6      4QNP  H1N1  NA   (NA A,B / 重 E,H / 轻 F,L; 二聚体表位)
"""
import csv
import numpy as np
from Bio.PDB import PDBParser, NeighborSearch
from Bio import SeqIO
from Bio.Align import PairwiseAligner

BASE = "/media/qxl/My_Passport/流行病学数据/广谱流感疫苗候选课题"
CONS = f"{BASE}/08_ESM2_AI分析与论文/data/consensus"
OUT = f"{BASE}/07_MANUSCRIPT_STRENGTHENING/08_BCELL_EXTERNAL_BENCHMARK/structures"

T3 = {'ALA':'A','ARG':'R','ASN':'N','ASP':'D','CYS':'C','GLN':'Q','GLU':'E','GLY':'G',
      'HIS':'H','ILE':'I','LEU':'L','LYS':'K','MET':'M','PHE':'F','PRO':'P','SER':'S',
      'THR':'T','TRP':'W','TYR':'Y','VAL':'V'}

# (pdb, antibody, subtype, protein, region, antigen_chains, antibody_chains)
structures = [
    # ---- 已有 stalk（HA2 茎部）----
    ("3GBN", "CR6261",  "H1N1", "HA", "stalk",              ["A", "B"], ["H", "L"]),
    ("3SDY", "CR8020",  "H3N2", "HA", "stalk",              ["A", "B"], ["H", "L"]),
    ("4FQI", "CR9114",  "H5N1", "HA", "stalk",              ["A", "B"], ["H", "L"]),
    # ---- 新增 RBD（HA1 受体结合位点）----
    ("3SM5", "CH65",    "H1N1", "HA", "RBD (receptor-binding site)", ["A", "B"], ["H", "L"]),
    ("4FP8", "C05",     "H3N2", "HA", "RBD (receptor-binding site)", ["A"],      ["H", "L"]),
    ("4GMS", "S139/1",  "H3N2", "HA", "RBD (receptor-binding site)", ["A", "B"], ["H", "L"]),
    ("4O58", "F045-092","H3N2", "HA", "RBD (receptor-binding site)", ["A", "B"], ["H", "L"]),
    # ---- 新增 NA（酶活性区）----
    ("4QNP", "CD6",     "H1N1", "NA", "enzyme active site",           ["A", "B"], ["E", "F", "H", "L"]),
]

# 共识序列：HA 与 NA
cons = {}
for sub in ["H1N1", "H3N2", "H5N1"]:
    cons[(sub, "HA")] = str(list(SeqIO.parse(f"{CONS}/{sub}_HA.fasta", "fasta"))[0].seq)
    cons[(sub, "NA")] = str(list(SeqIO.parse(f"{CONS}/{sub}_NA.fasta", "fasta"))[0].seq)

aligner = PairwiseAligner()
aligner.mode = 'local'; aligner.match_score = 2
aligner.mismatch_score = -1; aligner.open_gap_score = -2; aligner.extend_gap_score = -1

def chain_seq_resnums(chain):
    resnums = []; seq = []
    for res in chain:
        if res.id[0] != ' ':
            continue
        aa = T3.get(res.resname.strip())
        if aa is None:
            continue
        resnums.append(res.id[1]); seq.append(aa)
    return "".join(seq), resnums

def map_to_cons(chain_seq, resnums, contact, cons_seq):
    a = aligner.align(cons_seq, chain_seq)[0]
    target_blocks, query_blocks = a.aligned
    q2t = {}
    for (ts, te), (qs, qe) in zip(target_blocks, query_blocks):
        for off in range(te - ts):
            q2t[qs + off] = ts + off
    cons_contact = []
    for ri, rn in enumerate(resnums):
        if rn in contact and ri in q2t:
            cons_contact.append(q2t[ri] + 1)  # 1-indexed
    return sorted(set(cons_contact))

out = []
for pdb, ab, sub, prot, region, ag_chains, ab_chains in structures:
    parser = PDBParser(QUIET=True)
    s = parser.get_structure(pdb, f"{OUT}/{pdb}.pdb")[0]
    # 抗体原子（重链+轻链，全部指定链）
    ab_atoms = []
    for ch in ab_chains:
        if ch in s:
            ab_atoms.extend(list(s[ch].get_atoms()))
    ns = NeighborSearch(ab_atoms)
    all_contact = []
    for ag in ag_chains:
        if ag not in s:
            continue
        chain = s[ag]
        contact = set()
        for atom in chain.get_atoms():
            if ns.search(atom.coord, 4.5):
                contact.add(atom.get_parent().id[1])
        seq, resnums = chain_seq_resnums(chain)
        cc = map_to_cons(seq, resnums, contact, cons[(sub, prot)])
        all_contact.extend(cc)
    all_contact = sorted(set(all_contact))
    out.append(dict(structure=pdb, antibody=ab, subtype=sub, protein=prot, region=region,
                    n_contact_residues=len(all_contact),
                    consensus_positions=";".join(map(str, all_contact)),
                    source=f"PDB:{pdb}"))

with open(f"{OUT}/09_NEUTRALIZING_FOOTPRINTS_EXTENDED.tsv", "w", newline="") as f:
    w = csv.DictWriter(f, fieldnames=list(out[0].keys()), delimiter="\t")
    w.writeheader(); w.writerows(out)

print(f"{'antibody':9s} {'subtype':5s} {'protein':3s} {'region':28s} {'n_contact':>8s}")
for r in out:
    print(f"{r['antibody']:9s} {r['subtype']:5s} {r['protein']:3s} {r['region']:28s} {r['n_contact_residues']:8d}")
print(f"\nsaved {OUT}/09_NEUTRALIZING_FOOTPRINTS_EXTENDED.tsv")
```
