## Supplementary material for "Conserved influenza A epitope candidate regions and a benchmark of ESM-2 sequence features": Li_et_al_Supplementary_Data: 11_EXTENDED_FOOTPRINTS_SUMMARY.md

# 中和抗体足迹扩展（RBD + NA）—— 结果摘要

生成日期：2026-08-13

## 一、新增足迹（PDB 已下载并核验链组成与菌株来源）

方法：接触 = 抗原残基任一原子与抗体（重链+轻链）任一原子距离 ≤ 4.5 Å；HA1/HA2 分别局部比对映射到全长 HA 共识，NA 链映射到 NA 共识。

| 抗体 | PDB | 结合菌株 | 亚型 | 蛋白 | 区域 | 接触残基数 |
|---|---|---|---|---|---|---|
| CH65 | 3SM5 | A/Solomon Islands/3/2006 | H1N1 | HA | RBD | 23 |
| C05 | 4FP8 | A/Hong Kong/1/1968 | H3N2 | HA | RBD | 21 |
| S139/1 | 4GMS | A/Victoria/3/1975 | H3N2 | HA | RBD | 23 |
| F045-092 | 4O58 | A/Victoria/3/1975 | H3N2 | HA | RBD | 19 |
| CD6 | 4QNP | A/California/07/2009 (pdm09) | H1N1 | NA | 酶活性区 | 30 |

（原有 3 个 stalk 抗体 CR6261/3GBN、CR8020/3SDY、CR9114/4FQI 保持不变，接触数 25/23/25。）

**RBD 足迹一致性（正面质控）**：C05、S139/1、F045-092 三个 H3N2 RBD 抗体在共识上的接触位点高度重叠（147–153、169–175、205–210、242–244），符合"都结合受体结合位点"的预期，说明映射可靠。

## 二、关键发现：表面候选与中和足迹**无重叠**（诚实负面）

把 10 个表面候选（4 HA + 6 NA）定位到各自原生亚型共识后，与 8 个中和抗体足迹做重叠检验：

| 候选 | 原生亚型 | 共识位置 | 与中和足迹 |
|---|---|---|---|
| HA CKLRGVAPLHLGKCNIAGW | H1N1 | 59–77 | **无重叠**（且不在 RBD 108–241 内） |
| HA KHLLSRINHFEKI | H5N1 | 118–130 | 无重叠 |
| NA RYPGVRCVC | H3N2 | 283–291 | 无重叠* |
| 其余 NA 5 个 | H5N1 | — | 无重叠 |

\* 注意：当前 NA 足迹只有 H1N1（CD6），无 H3N2 NA 足迹，故 RYPGVRCVC 的"无重叠"只是亚型不匹配，非真阴性。

**重要提醒（供 manuscript 修正）**：
1. **明星 HA 候选 `CKLRGVAPLHLGKCNIAGW` 被误标为"受体结合位点（RBD）"**。实测其在 H1N1 共识位置为 59–77（成熟 HA1 约 42–60，位于头部 N 端/vestigial esterase 子域），**不在 RBD**（RBD 足迹在共识 108–241，受体口袋约 190–220）。ai_ranking.csv 里该候选的 domain 字段还标"330-400区域"，同样与实际位置不符——原管线的编号体系与全球共识不一致（与审计冲突 2 同源）。
2. 候选表位与中和抗体足迹**不重合**，意味着"候选=已知中和表位"的隐含联系不成立，manuscript 不应暗示候选命中已知中和位点。

## 三、需用户确认的引用（抗体名 + PDB 已核验，文献页码待定）

- CH65：Whittle JR et al. *PNAS* 2011;108:14216（3SM5）
- C05：Ekiert DC et al. *Nature* 2012;489:526（4FP8）
- S139/1：Lee PS et al. *PNAS* 2012;109:17040（4GMS）
- F045-092：Lee PS et al. *PNAS* 2014;111:E1046（4O58）
- CD6：Wan H et al. *Nat Commun* 2015;6:6114（4QNP，PMID 25668439，已核验）

## 四、产物

- `structures/09_NEUTRALIZING_FOOTPRINTS_EXTENDED.tsv`（8 个抗体足迹）
- `10_neutralizing_footprints_extended.py`（复现脚本）
- `structures/3SM5.pdb / 4FP8.pdb / 4GMS.pdb / 4O58.pdb / 4QNP.pdb`（新下载结构）
