## Supplementary material for "Conserved influenza A epitope candidate regions and a benchmark of ESM-2 sequence features": Li_et_al_Supplementary_Data: 11_MANUSCRIPT_OUTLINE_FINAL.md

# Manuscript 大纲（最终定位）

定位：**计算候选筛选为主体 + ESM-2 批判性评估为辅**，诚实、有实质、可投稿。
目标期刊：BMC Bioinformatics / PeerJ / Virology Journal 级。

---

## 标题

**Systematic identification of cross-subtype conserved influenza A epitope candidates, with a critical benchmark of protein-language-model sequence features**

---

## 摘要（四句话）

1. 我们系统筛选了 1544 条 GISAID 甲流序列（H1N1/H3N2/H5N1，9 个蛋白）中**跨亚型保守的表位候选**：98 个候选区域，其中 38 个在三个亚型中精确保守，且 2005–2017 保持时间稳定。
2. 用保守性 + B/T 表位 + 结构 + 时间稳定性对候选分层排序，产出优先清单与示意性组装。
3. 我们**额外批判性评估了 ESM-2 特征**是否增强排序：PLL 与保守度中等相关（ρ=0.25–0.39，本质是保守性代理），对 T-cell 表位无增量（ΔAUROC +0.004, p=0.455），contact-density 非可及性代理。
4. 我们记录了数据质量与可复现性问题（序列异质、坐标映射、伪重复），并提供可审计的 benchmark 数据集。

---

## 引言（3 段）

- **段1**：甲流抗原漂移迫使疫苗年度更新；广谱疫苗需保守表位；传统流程 = MSA 保守性 + B/T 表位预测。
- **段2**：蛋白质语言模型（ESM-2）被广泛用于蛋白分析，其零样本似然被寄望于捕捉"进化约束"。**开放问题**：对表位优先排序，它是否在传统保守性之外提供增量？
- **段3**：本文两件事——①系统筛选跨亚型保守表位候选；②用严格 benchmark 回答 ESM-2 是否有增量（诚实报告）。

---

## 结果（5 节）

### 1. 跨亚型保守表位候选的系统筛选
- 数据：1544 序列、9 蛋白、3 亚型；共识序列构建（mode-length 过滤，因原始序列长度异质 40–49 种）。
- 保守性（5 阈值）+ B-cell（16-mer）+ T-cell（netMHCpan-4.1, 14 HLA-I）。
- 产出：105 表位集中区 → 98 去重候选区域。
- **Table 1**：数据流水线；**Table 2**：候选清单。

### 2. 跨亚型与时间保守性
- 38/98 候选在 H1N1/H3N2/H5N1 三共识中精确保守。
- 时间：HA RBD 候选 2007–2017 连续 11 年 53–100% 一致度；NA 活性区候选同样稳定。
- **Fig 1**：跨亚型保守性热图；**Fig 2**：时间稳定性。

### 3. 候选分层排序与结构
- 分层（表面糖蛋白 vs 内部蛋白）排序；权重为启发式，敏感性分析（top-10 Jaccard 0.43–0.54）。
- 示意性多表位组装（AlphaFold pLDDT 55.9–64.3，诚实说明不足以支持结构可行性）。
- **Fig 3**：排序；**Fig 4**：示意组装。

### 4. ESM-2 特征的批判性评估（诚实负面）
- PLL 与 MSA 保守度中等相关（ρ=0.25–0.39），与 Shannon 熵负相关 → 本质是保守性代理。
- T-cell benchmark：PLL 无增量（ΔAUROC +0.004, p=0.455）。
- contact-density vs 真实 RSA：HA +0.147 / NA −0.132，方向不一致 → 非可及性代理。
- B-cell/中和 benchmark：22 非冗余 cluster（5 中和），功效不足；补充 3 个 stalk 广谱中和抗体（CR6261/CR8020/CR9114）晶体足迹（HA2 stalk）。
- **Fig 5**：PLL vs 保守度散点；**Fig 6**：T-cell AUROC；**Fig 7**：RSA 验证。

### 5. 数据质量与可复现性（对领域的贡献）
- GISAID 序列异质 → 必须 mode-length 过滤。
- IEDB 线性端点漏构象表位（此前 n=4 是检索丢失，非数据缺乏）。
- match_count ≠ 亚型数；netMHCpan 阈值笔误（%Rank<2%）。
- 伪重复：33 表位 → 22 cluster，须以 cluster 为 resampling unit。
- **Table 3**：benchmark 数据集统计。

---

## 讨论

- 主体结论：产出一份**跨亚型、时间稳定**的甲流保守表位候选清单（纯计算，未验证）。
- 批判性结论：**对保守性依赖型表位任务，ESM-2 掩码似然 ≈ 序列保守性，未提供可测增量**——明确其用途边界，防止后人重复踩坑。
- 局限：B-cell benchmark 小样本；数据只到 2017；纯计算无实验验证；B-cell 工具名缺失。

---

## 方法（摘要）

- 数据：GISAID（2005–2017），9 蛋白 × 3 亚型，mode-length 过滤 + 多数投票共识。
- 保守性：MAFFT（过滤后）+ positional identity + Shannon entropy。
- 表位：netMHCpan-4.1（14 HLA-I, %Rank_EL<2% & Aff<500nM）+ 16-mer B-cell + netMHCIIpan-4.3（6 DRB1）。
- ESM-2：group-masked PLL（8 组，单点敏感性 ρ=0.93）+ attention contact-density。
- 结构：FreeSASA（1RU7 HA, 2HTY NA）+ 抗体复合物足迹（3GBN/3SDY/4FQI）。
- benchmark：IEDB query API，亚型/坐标映射，聚类（50% 重叠），matched background（±0.05 保守度、buffer=10），cluster 为 resampling unit。

---

## 表/图清单

- Table 1：数据流水线（各步 N）
- Table 2：98 候选清单（蛋白/序列/坐标/保守度/跨亚型/B/T）
- Table 3：benchmark 统计（70→22 cluster、5 中和、证据分布）
- Fig 1：跨亚型保守性热图
- Fig 2：时间稳定性
- Fig 3：候选排序
- Fig 4：示意组装
- Fig 5：PLL vs 保守度散点
- Fig 6：T-cell AUROC（A/C/D）
- Fig 7：contact-density vs RSA

---

## 数据可用性

- benchmark 数据集 + 脚本（可 Zenodo）。
- 不含受限 GISAID 序列（用公开 IEDB 表位 + 共识）。

---

## 与旧稿的关系

- 旧稿"ESM-2 改进广谱疫苗候选"→ **废止**。
- 新稿：**候选筛选为主体（有实质）+ ESM-2 批判性评估（诚实）**。
- 所有真实分析保留，但 ESM-2 从"创新点"降为"被评估的特征"。
