## Supplementary material for "Conserved influenza A epitope candidate regions and a benchmark of ESM-2 sequence features": Li_et_al_Supplementary_Data: build_dataset.py

```
#!/usr/bin/env python3
"""Build the B-cell / neutralizing-antibody benchmark dataset from IEDB.

Fetches influenza A B-cell epitopes (linear + discontinuous), extracts
HA/NA surface-glycoprotein epitopes with PMID / IEDB ID / protein / positions /
sequence / evidence type / neutralization evidence.
"""
import json, csv, subprocess, urllib.parse
from collections import Counter

OUT = "/media/qxl/My_Passport/流行病学数据/广谱流感疫苗候选课题/07_MANUSCRIPT_STRENGTHENING/08_BCELL_EXTERNAL_BENCHMARK"
org = urllib.parse.quote("{Influenza A virus}")

def fetch_all(filter_extra):
    rows = []
    for offset in range(0, 500, 50):
        url = (f"https://query-api.iedb.org/epitope_search?source_organism_names=cs.{org}"
               f"&bcell_ids=not.is.null&{filter_extra}&order=structure_id&limit=50&offset={offset}")
        r = subprocess.run(["curl","-s","--max-time","40",url,"-H","accept: application/json"],
                           capture_output=True, text=True)
        try:
            d = json.loads(r.stdout)
        except Exception:
            break
        if not isinstance(d, list) or not d:
            break
        rows.extend(d)
        if len(d) < 50:
            break
    return rows

rows = fetch_all("")
print(f"总 B-cell 表位: {len(rows)}")

def assay_types(r):
    names = r.get("assay_names") or []
    return set(n.split("|")[0].strip().lower() for n in names if n)

def is_ha_na(r):
    ags = r.get("curated_source_antigens", [])
    return any(("emagglutinin" in a.get("name","").lower()) or
               ("euraminidase" in a.get("name","").lower()) for a in ags)

# classify evidence
def evidence(r):
    at = assay_types(r)
    if any("neutraliz" in t or "survival from pathogen" in t or "pathogen challenge" in t for t in at):
        return "neutralization"
    if any("binding" in t for t in at):
        return "antibody-binding"
    if any("b cell" in t or "elispot" in t or "proliferation" in t or "cytotoxicity" in t for t in at):
        return "B-cell"
    return "other"

ha_na = [r for r in rows if is_ha_na(r)]
print(f"HA/NA B-cell 表位: {len(ha_na)}")

# extract dataset
dataset = []
for r in ha_na:
    ags = r.get("curated_source_antigens", [])
    # primary antigen (HA or NA)
    prot = None; start = None; end = None
    for a in ags:
        n = a.get("name","").lower()
        if "emagglutinin" in n:
            prot = "HA"; start = a.get("starting_position"); end = a.get("ending_position"); break
        elif "euraminidase" in n:
            prot = "NA"; start = a.get("starting_position"); end = a.get("ending_position"); break
    if prot is None:
        prot = "HA/NA-other"; start = ags[0].get("starting_position") if ags else None; end = ags[0].get("ending_position") if ags else None
    seq = r.get("linear_sequence") or ";".join(r.get("structure_descriptions") or [])
    dataset.append({
        "iedb_id": r.get("structure_id"),
        "structure_type": r.get("structure_type"),
        "protein": prot,
        "position": f"{start}-{end}" if start else "",
        "sequence": seq,
        "evidence": evidence(r),
        "assay_names": ";".join(sorted(assay_types(r))),
        "qualitative": ";".join(r.get("qualitative_measures") or []),
        "antibody_isotype": ";".join(r.get("antibody_isotypes") or []),
        "pubmed": ";".join(r.get("pubmed_ids") or []),
        "reference_titles": ";".join((r.get("reference_titles") or [])[:2]),
        "source_organism": ";".join(r.get("source_organism_names") or []),
    })

# write TSV
cols = ["iedb_id","structure_type","protein","position","sequence","evidence",
        "assay_names","qualitative","antibody_isotype","pubmed","reference_titles","source_organism"]
with open(f"{OUT}/01_BCELL_DATASET.tsv","w",newline="") as f:
    w = csv.DictWriter(f, fieldnames=cols, delimiter="\t")
    w.writeheader()
    for d in dataset:
        w.writerow(d)

# summary
print(f"\nHA/NA 数据集: {len(dataset)} 条")
print("protein:", dict(Counter(d["protein"] for d in dataset)))
print("structure_type:", dict(Counter(d["structure_type"] for d in dataset)))
print("evidence:", dict(Counter(d["evidence"] for d in dataset)))
print("独立 PMID:", len(set(p for d in dataset for p in d["pubmed"].split(";") if p)))
print(f"\nsaved {OUT}/01_BCELL_DATASET.tsv")
```
