## Supplementary material for "Conserved influenza A epitope candidate regions and a benchmark of ESM-2 sequence features": Li_et_al_Supplementary_Data: compose_figures.py

```
#!/usr/bin/env python3
"""
把 18 张图整合为 7 张主文多面板图 + 5 张附录图。

方案（用户确认：7 主文 + 5 附录）：
  主文 Fig1 候选保守全景  = Fig1(跨亚型) + Fig2(时间) + Fig16(H1N1谱系) + Fig17(H3N2)
  主文 Fig2 ESM-2 PLL     = Fig5(PLL≈保守) + Fig9(3D定位)
  主文 Fig3 T-cell        = Fig6
  主文 Fig4 contact/RSA   = Fig7
  主文 Fig5 B-cell        = Fig11(山脊) + Fig12(数据流)
  主文 Fig6 抗体足迹      = Fig8(stalk) + Fig18(扩展)
  主文 Fig7 dN/dS         = Fig15
  附录 S1–S5              = Fig3(排序) Fig4(组装) Fig10(环状) Fig13(异质) Fig14(HLA)

用现有 PNG 直接拼接（不重跑数据管线），面板加 A/B/C/D 标签。
"""
import os
import numpy as np
import matplotlib
matplotlib.use("Agg")
import matplotlib.pyplot as plt
import matplotlib.image as mpimg

FIG = "/media/qxl/My_Passport/流行病学数据/广谱流感疫苗候选课题/07_MANUSCRIPT_STRENGTHENING/figures"
MAIN = f"{FIG}/main"
SUPP = f"{FIG}/supplementary"
os.makedirs(MAIN, exist_ok=True); os.makedirs(SUPP, exist_ok=True)

INK = "#2B2F36"

def compose(panels, outpath, ncols, label_offset=(0.015, 0.965)):
    """panels: list of (png_path, label) — 拼成一张图。"""
    n = len(panels)
    nrows = (n + ncols - 1) // ncols
    # 估算每格高度（保持宽高比约 0.55）
    fig, axes = plt.subplots(nrows, ncols, figsize=(7.2 * ncols, 4.0 * nrows))
    axes = np.atleast_1d(axes).ravel()
    for ax in axes:
        ax.set_axis_off()
    for i, (path, label) in enumerate(panels):
        ax = axes[i]
        img = mpimg.imread(path)
        ax.imshow(img)
        ax.text(label_offset[0], label_offset[1], label, transform=ax.transAxes,
                fontsize=14, fontweight="bold", va="top", ha="left",
                color="white", bbox=dict(boxstyle="round,pad=0.25", fc="black", alpha=0.65, ec="none"))
    # 隐藏多余格子
    for j in range(n, len(axes)):
        axes[j].set_axis_off()
    fig.tight_layout(pad=0.6)
    fig.savefig(outpath, dpi=200, bbox_inches="tight", facecolor="white")
    plt.close(fig)
    print(f"composed {os.path.basename(outpath)} ({n} panels)")

# ---- 主文图 ----
compose([
    (f"{FIG}/Fig1_crosssubtype.png", "A"),
    (f"{FIG}/Fig2_temporal.png", "B"),
    (f"{FIG}/Fig16_lineage.png", "C"),
    (f"{FIG}/Fig17_h3n2_subtype_specificity.png", "D"),
], f"{MAIN}/Fig1_candidate_conservation.png", ncols=2)

compose([
    (f"{FIG}/Fig5_pll_conservation.png", "A"),
    (f"{FIG}/Fig9_structure_pll.png", "B"),
], f"{MAIN}/Fig2_esm_pll.png", ncols=2)

compose([
    (f"{FIG}/Fig11_ridge.png", "A"),
    (f"{FIG}/Fig12_dataflow.png", "B"),
], f"{MAIN}/Fig5_bcell_benchmark.png", ncols=2)

compose([
    (f"{FIG}/Fig8_footprints.png", "A"),
    (f"{FIG}/Fig18_extended_footprints.png", "B"),
], f"{MAIN}/Fig6_antibody_footprints.png", ncols=2)

# ---- 单面板主文图（直接复制重命名）----
import shutil
for src, dst in [
    ("Fig6_benchmark_auroc.png", "Fig3_tcell_benchmark.png"),
    ("Fig7_rsa.png", "Fig4_contact_rsa.png"),
    ("Fig15_dnds.png", "Fig7_dnds.png"),
]:
    shutil.copy(f"{FIG}/{src}", f"{MAIN}/{dst}")
    print(f"copied {src} -> {dst}")

# ---- 附录图 ----
for src, dst in [
    ("Fig3_ranking.png", "S1_ranking_sensitivity.png"),
    ("Fig4_assemblies.png", "S2_assemblies.png"),
    ("Fig10_circular_map.png", "S3_circular_map.png"),
    ("Fig13_heterogeneity.png", "S4_length_heterogeneity.png"),
    ("Fig14_hlacoverage.png", "S5_hla_coverage.png"),
]:
    shutil.copy(f"{FIG}/{src}", f"{SUPP}/{dst}")
    print(f"copied {src} -> supplementary/{dst}")

print("\nDONE: 主文 7 张 + 附录 5 张")
```
