## Supplementary material for "Conserved influenza A epitope candidate regions and a benchmark of ESM-2 sequence features": Li_et_al_Supplementary_Data: convert_manuscript.py

```
#!/usr/bin/env python3
"""
manuscript_v2.md → .docx / .tex / .md（用 pypandoc 内置 pandoc 3.9）。
"""
import pypandoc

BASE = "/media/qxl/My_Passport/流行病学数据/广谱流感疫苗候选课题/07_MANUSCRIPT_STRENGTHENING"
MD = f"{BASE}/manuscript_v2.md"

# .docx（自带 standalone，含标题）
pypandoc.convert_file(MD, "docx", outputfile=f"{BASE}/manuscript_v2.docx",
                      extra_args=["--toc", "--number-sections"])

# .tex（standalone 生成可编译的完整 LaTeX 文档）
pypandoc.convert_file(MD, "latex", outputfile=f"{BASE}/manuscript_v2.tex",
                      extra_args=["--standalone", "--number-sections",
                                  "--variable", "documentclass=article",
                                  "--variable", "fontsize=11pt",
                                  "--variable", "geometry=margin=2.5cm"])

print("已生成 manuscript_v2.docx 与 manuscript_v2.tex")
```
