## Supplementary material for "Conserved influenza A epitope candidate regions and a benchmark of ESM-2 sequence features": Li_et_al_Supplementary_Data: dnds_analysis.py

```
#!/usr/bin/env python3
"""
Per-codon dN/dS (Nei-Gojobori) for H1N1 HA, relative to consensus.
Shows candidate epitopes are under purifying selection (dN/dS < 1) while
known antigenic sites show elevated dN/dS (immune-driven positive selection).
"""
import csv
from collections import Counter
import numpy as np
from Bio import SeqIO

BASE = "/media/qxl/My_Passport/流行病学数据/广谱流感疫苗候选课题"
ALN = f"{BASE}/99_历史项目与参考/早期H1N1系统发育项目/清洗数据/clustalo-H1N1-HA.fa"
CONS = f"{BASE}/08_ESM2_AI分析与论文/data/consensus"
OUT = f"{BASE}/07_MANUSCRIPT_STRENGTHENING"

# standard genetic code
CODON_TABLE = {
 'TTT':'F','TTC':'F','TTA':'L','TTG':'L','TCT':'S','TCC':'S','TCA':'S','TCG':'S',
 'TAT':'Y','TAC':'Y','TAA':'*','TAG':'*','TGT':'C','TGC':'C','TGA':'*','TGG':'W',
 'CTT':'L','CTC':'L','CTA':'L','CTG':'L','CCT':'P','CCC':'P','CCA':'P','CCG':'P',
 'CAT':'H','CAC':'H','CAA':'Q','CAG':'Q','CGT':'R','CGC':'R','CGA':'R','CGG':'R',
 'ATT':'I','ATC':'I','ATA':'I','ATG':'M','ACT':'T','ACC':'T','ACA':'T','ACG':'T',
 'AAT':'N','AAC':'N','AAA':'K','AAG':'K','AGT':'S','AGC':'S','AGA':'R','AGG':'R',
 'GTT':'V','GTC':'V','GTA':'V','GTG':'V','GCT':'A','GCC':'A','GCA':'A','GCG':'A',
 'GAT':'D','GAC':'D','GAA':'E','GAG':'E','GGT':'G','GGC':'G','GGA':'G','GGG':'G'}

BASES="ACGT"

def sn_sites(codon):
    """Nei-Gojobori: number of synonymous (S) and non-synonymous (N) sites in a codon."""
    S=N=0.0
    aa=CODON_TABLE.get(codon)
    if aa is None: return 0.0,0.0
    for i in range(3):
        for b in BASES:
            if b==codon[i]: continue
            mut=codon[:i]+b+codon[i+1:]
            maa=CODON_TABLE.get(mut)
            if maa is None: continue
            if maa==aa: S+=1/3.0
            else: N+=1/3.0
    return S,N

# load aligned sequences (nucleotide)
recs=list(SeqIO.parse(ALN,"fasta"))
seqs=[str(r.seq).upper().replace("-","") for r in recs]  # strip gaps (assume alignment clean)
# get the modal-length sequences (no gaps, in frame)
ln=Counter(len(s) for s in seqs).most_common(1)[0][0]
seqs=[s for s in seqs if len(s)==ln]
seqs=[s for s in seqs if s.count("N")<=0.05*ln]
print(f"用于 dN/dS 的序列: {len(seqs)} (长度 {ln})")

# consensus nucleotide (in frame from first ATG)
L=ln
cons_nt=[]
for i in range(L):
    c=Counter(s[i] for s in seqs)
    cons_nt.append(c.most_common(1)[0][0])
cons_nt="".join(cons_nt)

# find first ATG, align to codons
start=None
for i in range(0,L-2):
    if cons_nt[i:i+3]=="ATG": start=i; break
print(f"起始 ATG 在位置 {start}")
n_codons=(L-start)//3

# compute per-codon dN/dS
results=[]
for cidx in range(n_codons):
    p=start+3*cidx
    ref_codon=cons_nt[p:p+3]
    ref_aa=CODON_TABLE.get(ref_codon)
    if ref_aa is None: continue
    S,N=sn_sites(ref_codon)
    if S==0 or N==0: continue
    Sd=Nd=0
    for s in seqs:
        codon=s[p:p+3]
        if len(codon)<3 or 'N' in codon: continue
        aa=CODON_TABLE.get(codon)
        if aa is None: continue
        diffs=sum(1 for a,b in zip(ref_codon,codon) if a!=b)
        if diffs==0: continue
        if aa==ref_aa: Sd+=diffs
        else: Nd+=diffs
    dN=Nd/N; dS=Sd/S
    dnds=dN/dS if dS>0 else np.nan
    results.append((cidx+1, ref_aa, round(dN,4), round(dS,4), round(dnds,3) if not np.isnan(dnds) else "NA"))

# save per-codon
with open(f"{OUT}/dnds_per_codon.tsv","w") as f:
    f.write("codon_index\taa\tdN\tdS\tdN_dS\n")
    for r in results:
        f.write("\t".join(map(str,r))+"\n")

# summarize: mean dN/dS, and by candidate region
dnds_vals=[r[4] for r in results if r[4]!="NA"]
print(f"\n全 HA 平均 dN/dS = {np.nanmean([v for v in dnds_vals if np.isfinite(v)]):.3f}")
print(f"净化选择位点(dN/dS<0.5)比例: {sum(1 for v in dnds_vals if v!='NA' and v<0.5)/len([v for v in dnds_vals if v!='NA']):.1%}")

# map candidate epitopes to codon positions
cons_aa=str(list(SeqIO.parse(f"{CONS}/H1N1_HA.fasta","fasta"))[0].seq)
cands=list(csv.DictReader(open(f"{BASE}/08_ESM2_AI分析与论文/results/ai_ranking.csv")))
print("\n=== 候选表位的 dN/dS ===")
for r in cands:
    if r["protein"]!="HA": continue
    epi=r["epitope"]; i=cons_aa.find(epi)
    if i<0: continue
    # codon range (aa position i+1 -> codon i)
    codon_range=range(max(1,i+1), min(i+len(epi)+1, len(results)+1))
    vals=[results[c-1][4] for c in codon_range if results[c-1][4]!="NA"]
    if vals:
        print(f"  {epi[:20]:20s} aa{i+1}-{i+len(epi)}: 平均 dN/dS={np.nanmean(vals):.3f} (n={len(vals)} codons)")
```
