## Supplementary material for "Conserved influenza A epitope candidate regions and a benchmark of ESM-2 sequence features": Li_et_al_Supplementary_Data: export_supplementary_tables.py

```
#!/usr/bin/env python3
"""
导出 4 张补充表（Table S1–S4）为投稿用 TSV。
数据源：NC_main_figures_plotting_data.xlsx（12 个数据 Sheet，缺失值统一 NA）。
na_filter=False 保留 protein 列的 "NA"（神经氨酸酶）。
"""
from pathlib import Path
import pandas as pd

ROOT = Path("/media/qxl/My_Passport/流行病学数据/广谱流感疫苗候选课题")
XLSX = ROOT / "NC_main_figures_plotting_data.xlsx"
OUT = ROOT / "07_MANUSCRIPT_STRENGTHENING" / "supplementary_tables"
OUT.mkdir(exist_ok=True)

book = pd.read_excel(XLSX, sheet_name=None, na_filter=False)

# 各 Sheet 列顺序整理（保持原 Excel 顺序即可）
def write(tsv_name, sheet):
    df = book[sheet]
    df.to_csv(OUT / tsv_name, sep="\t", index=False, na_rep="NA")
    print(f"{tsv_name:36s} <- {sheet} ({len(df)} rows)")

# Table S1：98 候选完整信息
write("Table_S1_candidates.tsv", "Fig1_candidate_conservation")

# Table S2：时间与 clade 保守性（3 块）
write("Table_S2_H1N1_temporal.tsv", "Fig1_temporal")
write("Table_S2_H1_clades.tsv", "Fig1_H1_clades")
write("Table_S2_H3_clades.tsv", "Fig1_H3_clades")

# Table S3：T/B 细胞 benchmark（3 块）
write("Table_S3_Tcell_benchmark.tsv", "Fig3_Tcell_benchmark")
write("Table_S3_Bcell_PLL.tsv", "Fig5_Bcell_PLL")
write("Table_S3_dataflow.tsv", "Fig5_dataflow")

# Table S4：足迹/RSA/dN/dS（3 块）
write("Table_S4_antibody_footprints.tsv", "Fig6_antibody_footprints")
write("Table_S4_RSA_contact.tsv", "Fig4_RSA_contact")
write("Table_S4_dNdS.tsv", "Fig7_dNdS")

print("\n导出完成 ->", OUT)
```
