## Supplementary material for "Conserved influenza A epitope candidate regions and a benchmark of ESM-2 sequence features": Li_et_al_Supplementary_Data: fig_style.py

```
#!/usr/bin/env python3
"""共享绘图样式（提取自 figure1_redesign.py，供 F1–F5 统一视觉体系）。"""
import matplotlib as mpl
import matplotlib.pyplot as plt
from matplotlib.colors import LinearSegmentedColormap

# ---- 颜色体系 ----
NAVY = "#102F57"
BLUE = "#245F9E"
BLUE_DARK = "#174C84"
BLUE_MID = "#4D86BE"
BLUE_LIGHT = "#AFCBE6"
RED = "#C63F36"
RED_DARK = "#9F2F29"
ORANGE = "#E9832E"
ORANGE_LIGHT = "#F1C8A0"
GRAY = "#8D939C"
GRAY_LIGHT = "#D8DDE3"
GRID = "#D9DDE3"
TEXT = "#142033"
WHITE = "#FFFFFF"

# 蛋白配色（panel E / protein strip）
PROTEIN_COLORS = {
    "PB1": "#4A3FC0", "PB2": "#3B78C8", "NP": "#3C9270", "PA": "#5AA9A1",
    "HA": "#D04C4C", "NA": "#E98636", "NEP": "#909A70", "NS1": "#B58B50", "M": "#8D939C",
}
OTHER_PROTEIN_COLOR = "#777D87"

# 连续 colormap
ZPLL_CMAP = LinearSegmentedColormap.from_list(
    "zpll", ["#1E5A94", "#B8D4E9", "#F7F7F7", "#F1B08D", "#C63F36"])
CONTACT_CMAP = LinearSegmentedColormap.from_list(
    "contact", ["#EAF2F9", "#96C0DF", "#1C5A96"])


def setup_style():
    mpl.rcParams.update({
        "font.family": ["Arial", "DejaVu Sans"],
        "font.size": 8.4,
        "axes.titlesize": 10.0,
        "axes.labelsize": 8.2,
        "xtick.labelsize": 7.2,
        "ytick.labelsize": 7.2,
        "axes.linewidth": 0.65,
        "axes.edgecolor": "#6F7680",
        "text.color": TEXT,
        "xtick.color": TEXT,
        "ytick.color": TEXT,
        "xtick.direction": "out",
        "ytick.direction": "out",
        "xtick.major.width": 0.6,
        "ytick.major.width": 0.6,
        "axes.spines.top": False,
        "axes.spines.right": False,
        "pdf.fonttype": 42,
        "ps.fonttype": 42,
        "svg.fonttype": "none",
    })


def panel(ax, letter, title):
    """角标字母（NAVY 粗体）+ 标题（TEXT 粗体），置于面板左上角。"""
    ax.text(-0.04, 1.06, letter, transform=ax.transAxes, fontsize=15,
            fontweight="bold", color=NAVY, va="top", ha="right")
    ax.text(-0.02, 1.06, title, transform=ax.transAxes, fontsize=10.6,
            fontweight="bold", color=TEXT, va="top", ha="left")


def save(fig, path):
    fig.savefig(f"{path}.svg", bbox_inches="tight", facecolor=WHITE, pad_inches=0.02)
    fig.savefig(f"{path}.pdf", bbox_inches="tight", facecolor=WHITE, pad_inches=0.02)
    fig.savefig(f"{path}.png", dpi=600, bbox_inches="tight", facecolor=WHITE, pad_inches=0.02)
    plt.close(fig)
```
