## Supplementary material for "Conserved influenza A epitope candidate regions and a benchmark of ESM-2 sequence features": Li_et_al_Supplementary_Data: figure1_redesign.py

```
#!/usr/bin/env python3
# -*- coding: utf-8 -*-
"""
Figure 1 redesign: candidate screening workflow + multi-evidence landscape.

Usage:
    python figure1_redesign.py NC_main_figures_plotting_data.xlsx

Outputs:
    Figure1_redesign.svg
    Figure1_redesign.pdf
    Figure1_redesign.png

Expected candidate-level information (column names can vary; aliases are handled):
    candidate ID, protein source, H1N1/H3N2/H5N1 conservation status,
    z-PLL, contact density, composite/final score.

Conservation status should be one of: exact / fuzzy / absent
(or common variants such as conserved, partial, missing, 1/0.5/0).
"""

from __future__ import annotations

import sys
from pathlib import Path
import re
import numpy as np
import pandas as pd
import matplotlib as mpl
import matplotlib.pyplot as plt
from matplotlib import patches
from matplotlib.colors import ListedColormap, LinearSegmentedColormap, Normalize
from matplotlib.lines import Line2D


# ============================================================
# Global style
# ============================================================
mpl.rcParams.update({
    "font.family": ["Arial", "DejaVu Sans"],
    "font.size": 8.4,
    "axes.titlesize": 10.0,
    "axes.labelsize": 8.2,
    "xtick.labelsize": 7.2,
    "ytick.labelsize": 7.2,
    "axes.linewidth": 0.65,
    "pdf.fonttype": 42,
    "ps.fonttype": 42,
    "svg.fonttype": "none",
})

# Main palette
NAVY = "#102F57"
BLUE = "#245F9E"
BLUE_DARK = "#174C84"
BLUE_MID = "#4D86BE"
BLUE_LIGHT = "#AFCBE6"
RED = "#C63F36"
RED_DARK = "#9F2F29"
ORANGE = "#E9832E"
GRAY = "#8D939C"
GRAY_LIGHT = "#D8DDE3"
GRID = "#D9DDE3"
TEXT = "#142033"
WHITE = "#FFFFFF"

# Categorical conservation palette
CONS_COLORS = {
    "exact": BLUE_DARK,
    "fuzzy": BLUE_LIGHT,
    "absent": "#D5D9DE",
}

# Protein palette used in Panel E / protein-source strip
PROTEIN_COLORS = {
    "PB1": "#4A3FC0",
    "PB2": "#3B78C8",
    "NP": "#3C9270",
    "PA": "#5AA9A1",
    "HA": "#D04C4C",
    "NA": "#E98636",
    "NEP": "#909A70",
    "NS1": "#B58B50",
    "M": "#8D939C",
}
OTHER_PROTEIN_COLOR = "#777D87"

# Continuous maps
ZPLL_CMAP = LinearSegmentedColormap.from_list(
    "zpll", ["#1E5A94", "#B8D4E9", "#F7F7F7", "#F1B08D", "#C63F36"]
)
CONTACT_CMAP = LinearSegmentedColormap.from_list(
    "contact", ["#EAF2F9", "#96C0DF", "#1C5A96"]
)
SCORE_CMAP = LinearSegmentedColormap.from_list(
    "score", [BLUE_DARK, BLUE_LIGHT, "#F7F7F7", "#F3B48F", RED_DARK]
)

# Aliases: edit here only if your workbook uses unusual column names.
ALIASES = {
    "candidate": [
        "candidate", "candidate_id", "candidate id", "id", "sequence_id",
        "sequence id", "name", "candidate_name"
    ],
    "protein": [
        "protein", "protein_source", "protein source", "source_protein",
        "source protein", "gene", "gene_name", "gene name"
    ],
    "h1n1": [
        "h1n1", "h1n1_status", "h1n1 status", "h1_conservation",
        "h1n1_conservation", "h1n1 conservation"
    ],
    "h3n2": [
        "h3n2", "h3n2_status", "h3n2 status", "h3_conservation",
        "h3n2_conservation", "h3n2 conservation"
    ],
    "h5n1": [
        "h5n1", "h5n1_status", "h5n1 status", "h5_conservation",
        "h5n1_conservation", "h5n1 conservation"
    ],
    "zpll": [
        "z-pll", "z_pll", "zpll", "z pll", "pll_z", "pll z",
        "z-pll score", "z_pll_score"
    ],
    "contact": [
        "contact_density", "contact density", "contactdensity",
        "density", "mean_contact_density"
    ],
    "score": [
        "composite_score", "composite score", "final_score", "final score",
        "score", "priority_score", "priority score"
    ],
    "surface": [
        "surface", "surface_class", "surface class", "exposure",
        "protein_class", "protein class", "is_surface", "surface_exposure"
    ],
}


# ============================================================
# Data loading / harmonization
# ============================================================
def _norm_name(x: object) -> str:
    s = str(x).strip().lower()
    s = re.sub(r"[^a-z0-9]+", " ", s)
    return re.sub(r"\s+", " ", s).strip()


def _find_col(columns, aliases):
    norm_to_real = {_norm_name(c): c for c in columns}
    for a in aliases:
        key = _norm_name(a)
        if key in norm_to_real:
            return norm_to_real[key]
    return None


def _parse_conservation(v) -> str:
    if pd.isna(v):
        return "absent"

    if isinstance(v, (int, float, np.integer, np.floating)):
        x = float(v)
        if x >= 0.99:
            return "exact"
        if x > 0:
            return "fuzzy"
        return "absent"

    s = str(v).strip().lower()
    s_compact = re.sub(r"[^a-z0-9.+-]", "", s)

    exact_tokens = {
        "exact", "exactlyconserved", "conserved", "pass", "yes", "true",
        "identical", "match", "1", "1.0"
    }
    fuzzy_tokens = {
        "fuzzy", "partial", "near", "similar", "approx", "ambiguous",
        "0.5", "0.50"
    }
    absent_tokens = {
        "absent", "missing", "none", "no", "false", "fail", "0", "0.0", "-", "na", "n/a"
    }

    if s_compact in exact_tokens or "exact" in s or "identical" in s:
        return "exact"
    if s_compact in fuzzy_tokens or "fuzzy" in s or "partial" in s or "ambig" in s:
        return "fuzzy"
    if s_compact in absent_tokens or "absent" in s or "missing" in s:
        return "absent"

    try:
        x = float(s)
        if x >= 0.99:
            return "exact"
        if x > 0:
            return "fuzzy"
        return "absent"
    except Exception:
        raise ValueError(f"Unrecognized conservation value: {v!r}")


def _surface_class(value, protein) -> str:
    protein = str(protein).strip().upper()
    if value is None or pd.isna(value) or str(value).strip() == "":
        return "surface" if protein in {"HA", "NA"} else "internal"

    s = str(value).strip().lower()
    if s in {"1", "true", "yes", "surface", "ha/na", "external", "exposed"}:
        return "surface"
    if s in {"0", "false", "no", "internal", "other"}:
        return "internal"
    return "surface" if protein in {"HA", "NA"} else "internal"


def _candidate_table_from_sheet(df: pd.DataFrame, sheet_name: str):
    cols = list(df.columns)
    mapping = {k: _find_col(cols, v) for k, v in ALIASES.items()}

    required = ["protein", "h1n1", "h3n2", "h5n1", "zpll", "contact", "score"]
    missing = [k for k in required if mapping[k] is None]
    if missing:
        return None, mapping, missing

    out = pd.DataFrame(index=df.index)
    if mapping["candidate"] is not None:
        out["candidate"] = df[mapping["candidate"]].astype(str)
    else:
        out["candidate"] = [f"CAND_{i+1:03d}" for i in range(len(df))]

    out["protein"] = df[mapping["protein"]].astype(str).str.strip().str.upper()
    out["H1N1"] = df[mapping["h1n1"]].map(_parse_conservation)
    out["H3N2"] = df[mapping["h3n2"]].map(_parse_conservation)
    out["H5N1"] = df[mapping["h5n1"]].map(_parse_conservation)
    out["zPLL"] = pd.to_numeric(df[mapping["zpll"]], errors="coerce")
    out["contact_density"] = pd.to_numeric(df[mapping["contact"]], errors="coerce")
    out["composite_score"] = pd.to_numeric(df[mapping["score"]], errors="coerce")

    if mapping["surface"] is None:
        out["surface_class"] = [
            _surface_class(None, p) for p in out["protein"]
        ]
    else:
        out["surface_class"] = [
            _surface_class(v, p)
            for v, p in zip(df[mapping["surface"]], out["protein"])
        ]

    out = out.dropna(subset=["composite_score"]).copy()
    out["zPLL"] = out["zPLL"].fillna(out["zPLL"].median())
    out["contact_density"] = out["contact_density"].fillna(out["contact_density"].median())
    out["exact_count"] = (
        (out["H1N1"] == "exact").astype(int)
        + (out["H3N2"] == "exact").astype(int)
        + (out["H5N1"] == "exact").astype(int)
    )
    out = out.sort_values("composite_score", ascending=False).reset_index(drop=True)
    out["rank"] = np.arange(1, len(out) + 1)
    return out, mapping, []


def load_candidate_data(xlsx_path: str | Path) -> tuple[pd.DataFrame, str]:
    xlsx_path = Path(xlsx_path)
    if not xlsx_path.exists():
        raise FileNotFoundError(f"Input workbook not found: {xlsx_path}")

    book = pd.read_excel(xlsx_path, sheet_name=None, na_filter=False)
    attempts = []
    for sheet_name, df in book.items():
        if df is None or df.empty:
            continue
        result, mapping, missing = _candidate_table_from_sheet(df, sheet_name)
        if result is not None and len(result) >= 3:
            return result, sheet_name
        attempts.append((sheet_name, list(df.columns), missing))

    msg = [
        "Could not locate a candidate-level sheet with all required fields.",
        "Required logical fields: protein, H1N1, H3N2, H5N1, z-PLL, contact density, composite/final score.",
        "Workbook sheets and detected columns:",
    ]
    for sheet_name, cols, missing in attempts:
        msg.append(f"  - {sheet_name}: missing {missing}; columns={cols}")
    raise ValueError("\n".join(msg))


# ============================================================
# Utility drawing functions
# ============================================================
def clean_axis(ax, left=True, bottom=True):
    ax.spines["top"].set_visible(False)
    ax.spines["right"].set_visible(False)
    ax.spines["left"].set_visible(left)
    ax.spines["bottom"].set_visible(bottom)
    for sp in ax.spines.values():
        sp.set_linewidth(0.6)
        sp.set_color("#6F7680")
    ax.tick_params(length=3, width=0.55, color="#6F7680")


def panel_label(fig, x, y, letter, title):
    fig.text(x, y, letter, fontsize=15, fontweight="bold", color=NAVY, ha="left", va="top")
    fig.text(x + 0.026, y - 0.001, title, fontsize=10.6, fontweight="bold",
             color=TEXT, ha="left", va="top")


def minmax_norm(values):
    arr = np.asarray(values, dtype=float)
    lo, hi = np.nanmin(arr), np.nanmax(arr)
    if np.isclose(lo, hi):
        return np.full_like(arr, 0.5), lo, hi
    return (arr - lo) / (hi - lo), lo, hi


def draw_icon_evidence(ax, x, y, scale, color):
    # Three overlapping circles: multi-evidence integration
    for dx, dy in [(0, 0.012), (0.018, -0.004), (-0.018, -0.004)]:
        ax.add_patch(patches.Circle((x + dx * scale, y + dy * scale),
                                    0.018 * scale, fill=False, lw=1.6, ec=color))
    ax.add_patch(patches.Circle((x, y), 0.008 * scale, fc=color, ec="none"))


def draw_icon_mapping(ax, x, y, scale, color):
    ax.add_patch(patches.Circle((x, y), 0.028 * scale, fill=False, lw=1.7, ec=color))
    ax.plot([x + 0.020 * scale, x + 0.045 * scale],
            [y - 0.020 * scale, y - 0.045 * scale], color=color, lw=2.2, solid_capstyle="round")
    xs = np.linspace(x - 0.018 * scale, x + 0.015 * scale, 40)
    ys = y + 0.006 * scale * np.sin(np.linspace(0, 2.5 * np.pi, 40))
    ax.plot(xs, ys, color=color, lw=1.2)


def draw_icon_network(ax, x, y, scale, color):
    pts = np.array([[0, 0.026], [-0.025, 0], [0.025, 0], [0, -0.028]]) * scale
    pts[:, 0] += x
    pts[:, 1] += y
    for i in range(1, 4):
        ax.plot([pts[0, 0], pts[i, 0]], [pts[0, 1], pts[i, 1]], color=color, lw=1.3)
    ax.plot([pts[1, 0], pts[3, 0]], [pts[1, 1], pts[3, 1]], color=color, lw=1.0)
    ax.plot([pts[2, 0], pts[3, 0]], [pts[2, 1], pts[3, 1]], color=color, lw=1.0)
    for px, py in pts:
        ax.add_patch(patches.Circle((px, py), 0.007 * scale, fc=WHITE, ec=color, lw=1.4))


def draw_icon_target(ax, x, y, scale, color):
    for r in [0.030, 0.020, 0.010]:
        ax.add_patch(patches.Circle((x, y), r * scale, fill=False, lw=1.5, ec=color))
    ax.plot([x + 0.007 * scale, x + 0.038 * scale],
            [y + 0.007 * scale, y + 0.038 * scale], color=color, lw=1.6)
    ax.add_patch(patches.FancyArrowPatch(
        (x + 0.031 * scale, y + 0.031 * scale),
        (x + 0.046 * scale, y + 0.046 * scale),
        arrowstyle="-|>", mutation_scale=8, lw=1.2, color=color
    ))


def draw_workflow_panel(fig, rect):
    ax = fig.add_axes(rect)
    ax.set_xlim(0, 1)
    ax.set_ylim(0, 1)
    ax.axis("off")

    boxes = [
        {
            "title": "Evidence integration",
            "body": "Combine positional conservation\nwith B- and T-cell epitope evidence",
            "edge": "#6E9BD0", "face": "#F4F8FD", "icon": draw_icon_evidence, "icon_color": NAVY,
        },
        {
            "title": "Epitope mapping",
            "body": "Identify conserved epitope-containing\nregions on viral proteins",
            "edge": "#8D8AD0", "face": "#F7F6FC", "icon": draw_icon_mapping, "icon_color": NAVY,
        },
        {
            "title": "Candidate consolidation",
            "body": "Remove redundancy, integrate evidence\nand annotate candidate features",
            "edge": "#E79A54", "face": "#FFF8F1", "icon": draw_icon_network, "icon_color": "#A75622",
        },
        {
            "title": "Cross-subtype prioritization",
            "body": "Prioritize candidates conserved across\nH1N1, H3N2 and H5N1",
            "edge": "#D86B63", "face": "#FFF5F4", "icon": draw_icon_target, "icon_color": RED_DARK,
        },
    ]

    lefts = [0.02, 0.285, 0.55, 0.815]
    box_w = 0.205
    box_y = 0.18
    box_h = 0.64

    for i, (spec, x0) in enumerate(zip(boxes, lefts)):
        box = patches.FancyBboxPatch(
            (x0, box_y), box_w, box_h,
            boxstyle="round,pad=0.012,rounding_size=0.016",
            facecolor=spec["face"], edgecolor=spec["edge"], linewidth=0.85
        )
        ax.add_patch(box)
        spec["icon"](ax, x0 + 0.038, box_y + 0.40, 1.25, spec["icon_color"])
        ax.text(x0 + 0.075, box_y + 0.47, spec["title"],
                ha="left", va="center", fontsize=8.8, fontweight="bold", color=TEXT)
        ax.text(x0 + 0.075, box_y + 0.28, spec["body"],
                ha="left", va="center", fontsize=7.2, color=TEXT, linespacing=1.45)

        if i < len(boxes) - 1:
            x1 = x0 + box_w + 0.010
            x2 = lefts[i + 1] - 0.010
            ax.add_patch(patches.FancyArrowPatch(
                (x1, 0.50), (x2, 0.50),
                arrowstyle="-|>", mutation_scale=12, lw=1.1, color=NAVY
            ))

    return ax


def draw_row_grid(ax, n, linewidth=0.20):
    for y in np.arange(-0.5, n - 0.5, 1):
        ax.axhline(y, color=WHITE, lw=linewidth, zorder=3)


# ============================================================
# Panel B: integrated evidence landscape
# ============================================================
def draw_panel_b(fig, rect, df):
    x, y, w, h = rect
    gs = fig.add_gridspec(
        nrows=1, ncols=7,
        left=x, right=x + w, bottom=y, top=y + h,
        width_ratios=[0.22, 1.10, 0.30, 0.30, 0.34, 0.30, 0.78],
        wspace=0.16,
    )

    n = len(df)

    # Composite score color strip
    ax_score_strip = fig.add_subplot(gs[0, 0])
    score_normed, score_min, score_max = minmax_norm(df["composite_score"].values)
    score_matrix = (score_normed * 2 - 1).reshape(-1, 1)
    ax_score_strip.imshow(score_matrix, aspect="auto", cmap=SCORE_CMAP, vmin=-1, vmax=1, interpolation="nearest")
    ax_score_strip.set_title("Composite\nscore", fontsize=7.5, fontweight="bold", pad=6)
    ax_score_strip.set_xticks([])
    ax_score_strip.set_yticks([])
    for sp in ax_score_strip.spines.values():
        sp.set_visible(False)
    draw_row_grid(ax_score_strip, n)
    ax_score_strip.text(-0.62, -0.02, "High", transform=ax_score_strip.transAxes,
                        ha="left", va="top", color=RED_DARK, fontsize=7.0, fontweight="bold")
    ax_score_strip.text(-0.62, 1.02, "Low", transform=ax_score_strip.transAxes,
                        ha="left", va="bottom", color=BLUE_DARK, fontsize=7.0, fontweight="bold")

    # Conservation heatmap
    ax_cons = fig.add_subplot(gs[0, 1])
    cons_code = {"absent": 0, "fuzzy": 1, "exact": 2}
    cons = np.column_stack([
        df["H1N1"].map(cons_code),
        df["H3N2"].map(cons_code),
        df["H5N1"].map(cons_code),
    ])
    cons_cmap = ListedColormap([CONS_COLORS["absent"], CONS_COLORS["fuzzy"], CONS_COLORS["exact"]])
    ax_cons.imshow(cons, aspect="auto", cmap=cons_cmap, vmin=-0.5, vmax=2.5, interpolation="nearest")
    ax_cons.set_title("Conservation across subtypes", fontsize=7.8, fontweight="bold", pad=20)
    ax_cons.set_xticks([0, 1, 2], ["H1N1", "H3N2", "H5N1"])
    ax_cons.xaxis.tick_top()
    ax_cons.tick_params(axis="x", length=0, pad=5)
    ax_cons.set_yticks([])
    for sp in ax_cons.spines.values():
        sp.set_visible(False)
    draw_row_grid(ax_cons, n)
    ax_cons.axhline(-3.0, color="#9BA2AA", lw=0.6, clip_on=False)

    # Surface/internal strip
    ax_surface = fig.add_subplot(gs[0, 2])
    surface_code = df["surface_class"].map({"internal": 0, "surface": 1}).fillna(0).values.reshape(-1, 1)
    surface_cmap = ListedColormap([BLUE, RED])
    ax_surface.imshow(surface_code, aspect="auto", cmap=surface_cmap, vmin=-0.5, vmax=1.5, interpolation="nearest")
    ax_surface.set_title("Protein\nclass", fontsize=7.5, fontweight="bold", pad=6)
    ax_surface.set_xticks([])
    ax_surface.set_yticks([])
    for sp in ax_surface.spines.values():
        sp.set_visible(False)
    draw_row_grid(ax_surface, n)

    # z-PLL
    ax_z = fig.add_subplot(gs[0, 3])
    z = df["zPLL"].to_numpy(dtype=float)
    z_lim = max(abs(np.nanpercentile(z, 2)), abs(np.nanpercentile(z, 98)), 1e-9)
    ax_z.imshow(z.reshape(-1, 1), aspect="auto", cmap=ZPLL_CMAP,
                vmin=-z_lim, vmax=z_lim, interpolation="nearest")
    ax_z.set_title("z-PLL", fontsize=7.5, fontweight="bold", pad=6)
    ax_z.set_xticks([])
    ax_z.set_yticks([])
    for sp in ax_z.spines.values():
        sp.set_visible(False)
    draw_row_grid(ax_z, n)

    # Contact density
    ax_contact = fig.add_subplot(gs[0, 4])
    c = df["contact_density"].to_numpy(dtype=float)
    c_lo, c_hi = np.nanpercentile(c, [1, 99])
    if np.isclose(c_lo, c_hi):
        c_lo, c_hi = np.nanmin(c), np.nanmax(c) + 1e-9
    ax_contact.imshow(c.reshape(-1, 1), aspect="auto", cmap=CONTACT_CMAP,
                      vmin=c_lo, vmax=c_hi, interpolation="nearest")
    ax_contact.set_title("Contact\ndensity", fontsize=7.5, fontweight="bold", pad=6)
    ax_contact.set_xticks([])
    ax_contact.set_yticks([])
    for sp in ax_contact.spines.values():
        sp.set_visible(False)
    draw_row_grid(ax_contact, n)

    # Protein source strip
    ax_protein = fig.add_subplot(gs[0, 5])
    proteins = df["protein"].astype(str).str.upper().tolist()
    protein_order = list(dict.fromkeys(proteins))
    colors = [PROTEIN_COLORS.get(p, OTHER_PROTEIN_COLOR) for p in protein_order]
    p_code = {p: i for i, p in enumerate(protein_order)}
    p_vals = np.array([p_code[p] for p in proteins]).reshape(-1, 1)
    ax_protein.imshow(p_vals, aspect="auto", cmap=ListedColormap(colors),
                      vmin=-0.5, vmax=len(colors)-0.5, interpolation="nearest")
    ax_protein.set_title("Protein\nsource", fontsize=7.5, fontweight="bold", pad=6)
    ax_protein.set_xticks([])
    ax_protein.set_yticks([])
    for sp in ax_protein.spines.values():
        sp.set_visible(False)
    draw_row_grid(ax_protein, n)

    # Composite score bars aligned with rows
    ax_bar = fig.add_subplot(gs[0, 6])
    y_pos = np.arange(n)
    score = df["composite_score"].to_numpy(float)
    ax_bar.barh(y_pos, score, height=0.82, color=BLUE_DARK, edgecolor="none")
    ax_bar.set_ylim(n - 0.5, -0.5)
    ax_bar.set_xlim(0, max(1.0, float(np.nanmax(score) * 1.03)))
    ax_bar.set_title("Composite score", fontsize=7.5, fontweight="bold", pad=6)
    ax_bar.set_yticks([])
    ax_bar.set_xlabel("Score", labelpad=2)
    ax_bar.grid(axis="x", color=GRID, lw=0.45, alpha=0.8)
    ax_bar.set_axisbelow(True)
    clean_axis(ax_bar, left=False, bottom=True)

    # Legend row beneath Panel B
    legend_ax = fig.add_axes([x, y - 0.065, w, 0.052])
    legend_ax.axis("off")

    handles_cons = [
        patches.Patch(facecolor=CONS_COLORS["exact"], edgecolor="none", label="Exact"),
        patches.Patch(facecolor=CONS_COLORS["fuzzy"], edgecolor="none", label="Fuzzy"),
        patches.Patch(facecolor=CONS_COLORS["absent"], edgecolor="none", label="Absent"),
        patches.Patch(facecolor=RED, edgecolor="none", label="Surface (HA/NA)"),
        patches.Patch(facecolor=BLUE, edgecolor="none", label="Internal"),
    ]
    leg = legend_ax.legend(handles=handles_cons, ncol=5, loc="upper left",
                           frameon=False, handlelength=0.9, handleheight=0.9,
                           columnspacing=1.2, borderpad=0, fontsize=6.7)
    for t in leg.get_texts():
        t.set_color(TEXT)

    # Mini gradient legends
    g1 = fig.add_axes([x + w * 0.47, y - 0.053, w * 0.12, 0.012])
    g1.imshow(np.linspace(-1, 1, 256).reshape(1, -1), aspect="auto", cmap=ZPLL_CMAP)
    g1.set_xticks([0, 128, 255], ["Low", "0", "High"])
    g1.set_yticks([])
    g1.tick_params(axis="x", length=0, labelsize=6.0, pad=1)
    for sp in g1.spines.values(): sp.set_visible(False)
    fig.text(x + w * 0.53, y - 0.032, "z-PLL", ha="center", va="bottom", fontsize=6.4, color=TEXT)

    g2 = fig.add_axes([x + w * 0.63, y - 0.053, w * 0.12, 0.012])
    g2.imshow(np.linspace(0, 1, 256).reshape(1, -1), aspect="auto", cmap=CONTACT_CMAP)
    g2.set_xticks([0, 255], ["Low", "High"])
    g2.set_yticks([])
    g2.tick_params(axis="x", length=0, labelsize=6.0, pad=1)
    for sp in g2.spines.values(): sp.set_visible(False)
    fig.text(x + w * 0.69, y - 0.032, "Contact density", ha="center", va="bottom", fontsize=6.4, color=TEXT)

    return {
        "score_strip": ax_score_strip,
        "conservation": ax_cons,
        "surface": ax_surface,
        "zpll": ax_z,
        "contact": ax_contact,
        "protein": ax_protein,
        "bar": ax_bar,
    }


# ============================================================
# Panels C, D, E
# ============================================================
def draw_panel_c(fig, rect, df):
    ax = fig.add_axes(rect)
    counts = df["protein"].value_counts()
    proteins = counts.index.tolist()
    values = counts.values
    colors = [RED if p in {"HA", "NA"} else BLUE for p in proteins]

    y = np.arange(len(proteins))
    ax.barh(y, values, color=colors, height=0.62)
    ax.set_yticks(y, proteins)
    ax.invert_yaxis()
    ax.set_xlabel("Candidate count")
    ax.set_xlim(0, max(values) * 1.18)
    ax.grid(axis="x", color=GRID, linestyle="--", lw=0.5, alpha=0.8)
    ax.set_axisbelow(True)
    clean_axis(ax)

    for yi, v in zip(y, values):
        ax.text(v + max(values) * 0.015, yi, str(int(v)), va="center", ha="left", fontsize=7.2, color=TEXT)

    handles = [
        patches.Patch(facecolor=BLUE, label="Internal proteins"),
        patches.Patch(facecolor=RED, label="Surface proteins (HA/NA)"),
    ]
    ax.legend(handles=handles, frameon=False, fontsize=6.6, loc="lower right",
              handlelength=0.8, borderpad=0.2)
    return ax


def draw_panel_d(fig, rect, df):
    ax = fig.add_axes(rect)
    counts = df["exact_count"].value_counts().reindex([0, 1, 2, 3], fill_value=0)
    x = np.arange(4)
    colors = [GRAY, BLUE, ORANGE, RED]
    bars = ax.bar(x, counts.values, color=colors, width=0.52)
    ax.set_xticks(x, ["0", "1", "2", "3"])
    ax.set_xlabel("Subtypes with exact conservation")
    ax.set_ylabel("Candidates")
    ax.set_ylim(0, max(counts.values) * 1.28 + 1)
    clean_axis(ax)

    for bar, v in zip(bars, counts.values):
        ax.text(bar.get_x() + bar.get_width()/2, v + max(counts.values) * 0.035,
                str(int(v)), ha="center", va="bottom", fontsize=7.4, color=TEXT)

    # Data label is on the bar; callout text contains no count.
    ax.annotate(
        "Exactly conserved across\nH1N1, H3N2 and H5N1",
        xy=(3, counts.loc[3] * 0.88 if counts.loc[3] else 0),
        xytext=(3.55, max(counts.values) * 0.62),
        ha="left", va="center", fontsize=6.8, color=RED_DARK,
        bbox=dict(boxstyle="round,pad=0.35", fc="#FFF6F5", ec=RED, lw=0.7),
        arrowprops=dict(arrowstyle="-|>", color=RED, lw=0.8),
    )
    ax.set_xlim(-0.55, 4.65)
    return ax


def draw_panel_e(fig, rect, df, top_n=15):
    ax = fig.add_axes(rect)
    top = df.head(min(top_n, len(df))).copy()
    y = np.arange(1, len(top) + 1)

    # Thin guide lines from a common baseline to each point.
    baseline = max(0.0, float(top["composite_score"].min()) - 0.12)
    for yi, (_, row) in zip(y, top.iterrows()):
        p = str(row["protein"]).upper()
        c = PROTEIN_COLORS.get(p, OTHER_PROTEIN_COLOR)
        score = float(row["composite_score"])
        ax.plot([baseline, score], [yi, yi], color="#B9BEC5", lw=0.65, zorder=1)
        if row["surface_class"] == "surface":
            ax.scatter(score, yi, s=30, facecolor=c, edgecolor=c, linewidth=0.8, zorder=3)
        else:
            ax.scatter(score, yi, s=30, facecolor=WHITE, edgecolor=c, linewidth=1.1, zorder=3)

    ax.set_ylim(len(top) + 0.8, 0.2)
    ax.set_yticks(y)
    ax.set_ylabel("Candidate rank")
    xmin = max(0, baseline - 0.03)
    xmax = min(1.0, max(float(top["composite_score"].max()) + 0.06, 0.85))
    if xmax <= xmin:
        xmax = xmin + 0.5
    ax.set_xlim(xmin, xmax)
    ax.set_xlabel("Composite score")
    ax.grid(axis="y", color=GRID, lw=0.45, alpha=0.7)
    ax.set_axisbelow(True)
    clean_axis(ax)

    protein_present = list(dict.fromkeys(top["protein"].astype(str).str.upper()))
    protein_handles = [
        Line2D([0], [0], marker="o", linestyle="None", markersize=5,
               markerfacecolor=PROTEIN_COLORS.get(p, OTHER_PROTEIN_COLOR),
               markeredgecolor=PROTEIN_COLORS.get(p, OTHER_PROTEIN_COLOR), label=p)
        for p in protein_present
    ]
    exposure_handles = [
        Line2D([0], [0], marker="o", linestyle="None", markersize=5,
               markerfacecolor=TEXT, markeredgecolor=TEXT, label="Surface (HA/NA)"),
        Line2D([0], [0], marker="o", linestyle="None", markersize=5,
               markerfacecolor=WHITE, markeredgecolor=TEXT, label="Internal (other)"),
    ]

    leg1 = ax.legend(handles=protein_handles, title="Protein source", frameon=False,
                     fontsize=6.2, title_fontsize=6.5, loc="upper left",
                     bbox_to_anchor=(1.01, 1.03), borderaxespad=0,
                     labelspacing=0.25, handletextpad=0.35)
    ax.add_artist(leg1)
    ax.legend(handles=exposure_handles, title="Protein class", frameon=False,
              fontsize=6.2, title_fontsize=6.5, loc="lower left",
              bbox_to_anchor=(1.01, -0.02), borderaxespad=0,
              labelspacing=0.3, handletextpad=0.35)
    return ax


# ============================================================
# Figure assembly
# ============================================================
def make_figure(df: pd.DataFrame, out_prefix: str | Path = "Figure1_redesign"):
    out_prefix = Path(out_prefix)

    fig = plt.figure(figsize=(15.8, 10.2), facecolor=WHITE)

    # Panel labels/titles: intentionally no numeric counts in titles.
    panel_label(fig, 0.018, 0.975, "A", "Candidate screening workflow")
    draw_workflow_panel(fig, [0.040, 0.815, 0.935, 0.145])

    panel_label(fig, 0.018, 0.785, "B", "Multi-evidence landscape of candidates")
    draw_panel_b(fig, [0.045, 0.155, 0.595, 0.585], df)

    panel_label(fig, 0.655, 0.785, "C", "Candidates per protein")
    draw_panel_c(fig, [0.690, 0.585, 0.260, 0.155], df)

    panel_label(fig, 0.655, 0.535, "D", "Cross-subtype exact conservation")
    draw_panel_d(fig, [0.690, 0.355, 0.260, 0.145], df)

    panel_label(fig, 0.655, 0.310, "E", "Candidate prioritization by composite score")
    draw_panel_e(fig, [0.690, 0.105, 0.235, 0.165], df, top_n=15)

    # Thin separator under workflow for a cleaner hierarchy.
    fig.add_artist(Line2D([0.035, 0.965], [0.795, 0.795], transform=fig.transFigure,
                          color="#E4E7EB", lw=0.6))

    # Save vector first; PNG only for preview/manuscript convenience.
    fig.savefig(out_prefix.with_suffix(".svg"), bbox_inches="tight", facecolor=WHITE)
    fig.savefig(out_prefix.with_suffix(".pdf"), bbox_inches="tight", facecolor=WHITE)
    fig.savefig(out_prefix.with_suffix(".png"), dpi=600, bbox_inches="tight", facecolor=WHITE)
    plt.close(fig)


# ============================================================
# Command-line entry
# ============================================================
def main():
    if len(sys.argv) >= 2:
        input_path = Path(sys.argv[1])
    else:
        input_path = Path("NC_main_figures_plotting_data.xlsx")

    if len(sys.argv) >= 3:
        out_prefix = Path(sys.argv[2])
    else:
        out_prefix = Path("Figure1_redesign")

    df, sheet = load_candidate_data(input_path)
    print(f"Loaded candidate table from sheet: {sheet}")
    print(f"Candidates used: {len(df)}")
    print("Protein counts:")
    print(df["protein"].value_counts().to_string())
    print("Exact-conservation counts:")
    print(df["exact_count"].value_counts().sort_index().to_string())

    make_figure(df, out_prefix)
    print(f"Saved: {out_prefix.with_suffix('.svg')}")
    print(f"Saved: {out_prefix.with_suffix('.pdf')}")
    print(f"Saved: {out_prefix.with_suffix('.png')}")


if __name__ == "__main__":
    main()
```
