## Supplementary material for "Conserved influenza A epitope candidate regions and a benchmark of ESM-2 sequence features": Li_et_al_Supplementary_Data: lineage_conservation.py

```
#!/usr/bin/env python3
"""
Cross-lineage conservation of candidate epitopes using 2015-2025 H1N1 HA data
(with clade labels from the phylogenetic project). Forward-validation: are the
candidates (derived from 2002-2017 data) still conserved in recent lineages?
"""
import csv
from collections import Counter
import numpy as np
from Bio import SeqIO

BASE = "/media/qxl/My_Passport/流行病学数据/广谱流感疫苗候选课题"
ALN = f"{BASE}/99_历史项目与参考/早期H1N1系统发育项目/清洗数据/clustalo-H1N1-HA.fa"
META = f"{BASE}/99_历史项目与参考/早期H1N1系统发育项目/清洗数据/cleaned_metadata.csv"
CONS = f"{BASE}/08_ESM2_AI分析与论文/data/consensus"
OUT = f"{BASE}/07_MANUSCRIPT_STRENGTHENING"

# metadata: id -> clade, year
meta = {}
for r in csv.DictReader(open(META)):
    meta[r["Isolate_Id"]] = r

# load aligned nucleotide sequences, strip gaps, translate in frame
recs = list(SeqIO.parse(ALN, "fasta"))
print(f"加载 {len(recs)} 条比对序列")

def translate(seq):
    from Bio.Seq import Seq
    s = str(seq).upper().replace("-", "")
    for i in range(0, len(s)-2):
        if s[i:i+3] == "ATG":
            n = (len(s)-i)//3
            aa = []
            for c in range(n):
                codon = s[i+3*c:i+3*c+3]
                if len(codon) < 3 or "N" in codon:
                    aa.append("X")
                else:
                    aa.append(str(Seq(codon).translate()))
            return "".join(aa)
    return ""

# translate all, map to clade
seq_by_clade = Counter()
translated = {}
for r in recs:
    clade = meta.get(r.id, {}).get("Clade", "unassigned")
    aa = translate(r.seq)
    if aa and aa.count("X") <= 0.05*len(aa):
        seq_by_clade[clade] += 1
        translated.setdefault(clade, []).append(aa)

print(f"按谱系分组: {dict(seq_by_clade.most_common(10))}")

# candidates (HA)
cons_ha = str(list(SeqIO.parse(f"{CONS}/H1N1_HA.fasta", "fasta"))[0].seq)
cands = [r for r in csv.DictReader(open(f"{BASE}/08_ESM2_AI分析与论文/results/ai_ranking.csv")) if r["protein"]=="HA"]

# for each candidate, conservation fraction in each clade (>=90% identity)
def frac_in(seqs, epi):
    n=0; tot=0
    for s in seqs:
        if len(s) < len(epi): continue
        best=0.0
        for i in range(len(s)-len(epi)+1):
            best=max(best, sum(a==b for a,b in zip(epi,s[i:i+len(epi)]))/len(epi))
        tot+=1
        if best>=0.9: n+=1
    return n/tot if tot else np.nan

top_clades = [c for c,_ in seq_by_clade.most_common(6)]

print("\n=== 候选表位跨谱系保守 (2015-2025) ===")
print(f"{'epitope':24s} " + " ".join(f"{c[:12]:>14s}" for c in top_clades))
for r in cands:
    epi = r["epitope"]
    i = cons_ha.find(epi)
    if i < 0: continue
    vals = []
    for clade in top_clades:
        f = frac_in(translated.get(clade, []), epi)
        vals.append(f)
        print(f"{epi[:24]:24s} " + " ".join(f"{f:>14.2f}" if not np.isnan(f) else f"{'NA':>14s}" for f in [f]) + f"  [{clade[:12]}]")
    # 只在最后一行打印完整
# 汇总
print("\n=== 汇总: 候选跨谱系保守率 ===")
for r in cands:
    epi = r["epitope"]
    i = cons_ha.find(epi)
    if i < 0: continue
    all_seqs = [s for clade in top_clades for s in translated.get(clade, [])]
    f_all = frac_in(all_seqs, epi)
    f_clades = [frac_in(translated.get(c, []), epi) for c in top_clades]
    f_clades = [f for f in f_clades if not np.isnan(f)]
    print(f"  {epi[:22]:22s} 全谱系 {f_all:.2f} | 跨谱系最低 {min(f_clades):.2f} | 谱系间变异 {np.std(f_clades):.3f}")
```
