## Supplementary material for "Conserved influenza A epitope candidate regions and a benchmark of ESM-2 sequence features": Li_et_al_Supplementary_Data: make_fig1_overview.py

```
#!/usr/bin/env python3
"""
Figure 1 总览图：广谱候选的筛选结果与保守性全景（5 面板 A–E）。
A 筛选流程图；B 98候选综合证据热图；C 各蛋白候选数；D 跨亚型精确保守分布；E 前15候选棒棒糖。
数据源：tables/Table2_candidates_full_corrected.tsv
"""
import csv
from collections import Counter
import numpy as np
import matplotlib
matplotlib.use("Agg")
import matplotlib.pyplot as plt
from matplotlib.colors import ListedColormap
from matplotlib.patches import FancyBboxPatch
from matplotlib.gridspec import GridSpec

MS = "/media/qxl/My_Passport/流行病学数据/广谱流感疫苗候选课题/07_MANUSCRIPT_STRENGTHENING"
OUT = f"{MS}/figures/main/Fig1_overview.png"

BLUE = "#3A6EA5"; ORANGE = "#C77C3F"; PURPLE = "#7B5AA6"
SURFACE = "#1B9E8F"; INTERNAL = "#6B7B8C"; RED = "#B5453B"
GRAY = "#8A8F98"; INK = "#22262B"; LIGHT = "#C8CDD3"
plt.rcParams.update({
    "font.family": "Arial", "font.size": 8, "axes.spines.top": False,
    "axes.spines.right": False, "axes.linewidth": 0.6, "axes.edgecolor": INK,
    "axes.labelcolor": INK, "text.color": INK, "xtick.color": INK, "ytick.color": INK,
    "xtick.direction": "out", "ytick.direction": "out",
    "xtick.major.width": 0.6, "ytick.major.width": 0.6,
})

def panel(ax, label, title):
    ax.text(-0.08, 1.06, label, transform=ax.transAxes, fontsize=13, fontweight="bold",
            va="bottom", ha="left")
    ax.set_title(title, loc="left", fontsize=9, pad=10)

# ---------------------------------------------------------------- 数据
rows = list(csv.DictReader(open(f"{MS}/tables/Table2_candidates_full_corrected.tsv"), delimiter="\t"))
for r in rows:
    r["rank"] = int(r["rank"]); r["final_score"] = float(r["final_score"])
    r["zPLL"] = float(r["zPLL"]); r["contact_density"] = float(r["contact_density"])
    r["n_exact_subtypes"] = int(r["n_exact_subtypes"])
rows.sort(key=lambda r: r["rank"])   # rank 1..98 按 final_score 降序
N = len(rows)

surf = [1 if r["protein"] in ("HA", "NA") else 0 for r in rows]
def conv(v): return {"exact": 2, "fuzzy": 1, "-": 0}.get(v, 0)
cat = np.array([[conv(r["H1N1"]), conv(r["H3N2"]), conv(r["H5N1"])] for r in rows])
zpll = np.array([r["zPLL"] for r in rows]).reshape(-1, 1)
cd = np.array([r["contact_density"] for r in rows]).reshape(-1, 1)

prot_count = Counter(r["protein"] for r in rows)
prot_order = [p for p, _ in prot_count.most_common()]
prot_n = [prot_count[p] for p in prot_order]
n_exact_dist = Counter(r["n_exact_subtypes"] for r in rows)
top15 = rows[:15]

# ---------------------------------------------------------------- 图
fig = plt.figure(figsize=(13.5, 10.5))
gs = fig.add_gridspec(2, 2, height_ratios=[0.18, 1.0], width_ratios=[2.5, 1.0],
                      hspace=0.35, wspace=0.30)

# ===== A 流程图（顶部横跨）=====
axA = fig.add_subplot(gs[0, :]); axA.axis("off"); axA.set_xlim(0, 12); axA.set_ylim(0, 2.4)
steps = [
    ("Positional conservation\n+ B-cell + T-cell\n(16-mer / netMHCpan)", 0.3, 3.2, "#5B6672"),
    ("105 epitope-containing\nregions", 4.0, 2.1, "#7A8794"),
    ("98 unique candidates\n(after de-duplication)", 6.6, 2.5, ORANGE),
    ("38 exactly conserved\nin all 3 subtypes", 9.7, 2.1, RED),
]
for txt, x, w, c in steps:
    axA.add_patch(FancyBboxPatch((x, 0.9), w, 1.4, boxstyle="round,pad=0.06,rounding_size=0.16",
                                 fc=c, ec="none", alpha=0.9))
    axA.text(x + w/2, 1.6, txt, ha="center", va="center", fontsize=7.5, color="white", fontweight="bold")
for x1, x2 in [(3.5, 4.0), (6.1, 6.6), (9.1, 9.7)]:
    axA.annotate("", xy=(x2, 1.6), xytext=(x1, 1.6),
                 arrowprops=dict(arrowstyle="-|>", color=INK, lw=1.2))
panel(axA, "A", "Candidate screening workflow")

# ===== B 综合证据热图 =====
gsB = gs[1, 0].subgridspec(1, 5, width_ratios=[0.13, 1.6, 0.5, 0.5, 0.5], wspace=0.05)
ax_surf = fig.add_subplot(gsB[0])
ax_cat  = fig.add_subplot(gsB[1])
ax_z    = fig.add_subplot(gsB[2])
ax_cd   = fig.add_subplot(gsB[3])
ax_prot = fig.add_subplot(gsB[4])

ax_surf.imshow(np.array(surf).reshape(-1, 1), cmap=ListedColormap([INTERNAL, SURFACE]),
               aspect="auto", vmin=0, vmax=1)
ax_surf.set_xticks([]); ax_surf.set_yticks([])

ax_cat.imshow(cat, cmap=ListedColormap(["white", LIGHT, BLUE]), aspect="auto", vmin=0, vmax=2)
ax_cat.set_xticks([0, 1, 2]); ax_cat.set_xticklabels(["H1N1", "H3N2", "H5N1"], fontsize=7, rotation=45)
ax_cat.set_yticks([])

imz = ax_z.imshow(zpll, cmap="RdBu_r", aspect="auto", vmin=-2, vmax=2)
ax_z.set_xticks([0]); ax_z.set_xticklabels(["z-PLL"], fontsize=7, rotation=45); ax_z.set_yticks([])

imc = ax_cd.imshow(cd, cmap="Blues", aspect="auto", vmin=0, vmax=1)
ax_cd.set_xticks([0]); ax_cd.set_xticklabels(["contact\ndensity"], fontsize=7, rotation=45); ax_cd.set_yticks([])

ax_prot.set_xlim(0, 1); ax_prot.set_ylim(-0.5, N - 0.5); ax_prot.axis("off")
for i, r in enumerate(rows):
    ax_prot.text(0.03, N - 1 - i, r["protein"], fontsize=4.6, va="center",
                 color=SURFACE if r["protein"] in ("HA", "NA") else INTERNAL)

for ax in [ax_cat, ax_z, ax_cd, ax_surf]:
    for sp in ["top", "right", "bottom", "left"]:
        ax.spines[sp].set_visible(False)
    ax.tick_params(length=0)

# colorbars 放 B 面板下方
cbz = fig.colorbar(imz, ax=ax_z, fraction=0.5, pad=0.28, shrink=0.7)
cbz.set_label("z-PLL", fontsize=7); cbz.ax.tick_params(labelsize=6)
cbc = fig.colorbar(imc, ax=ax_cd, fraction=0.5, pad=0.28, shrink=0.7)
cbc.set_label("contact density", fontsize=7); cbc.ax.tick_params(labelsize=6)

ax_cat.set_title("B  98 candidates: multi-evidence heatmap (ranked by final_score)",
                 loc="left", fontsize=9, pad=10)
ax_cat.text(-0.10, 1.06, "B", transform=ax_cat.transAxes, fontsize=13, fontweight="bold",
            va="bottom", ha="left")

# 图例：surface/internal + 保守状态
from matplotlib.lines import Line2D
leg = ax_surf.legend(handles=[
    Line2D([0], [0], marker="s", color="w", markerfacecolor=SURFACE, markersize=8, label="surface (HA/NA)"),
    Line2D([0], [0], marker="s", color="w", markerfacecolor=INTERNAL, markersize=8, label="internal"),
    Line2D([0], [0], marker="s", color="w", markerfacecolor=BLUE, markersize=8, label="exact"),
    Line2D([0], [0], marker="s", color="w", markerfacecolor=LIGHT, markersize=8, label="fuzzy"),
    Line2D([0], [0], marker="s", color="w", markerfacecolor="white", markeredgecolor=GRAY, markersize=8, label="absent"),
], loc="lower left", frameon=False, fontsize=6, bbox_to_anchor=(-2.0, -0.15), ncol=5)

# ===== C / D / E（右列堆叠）=====
gsR = gs[1, 1].subgridspec(3, 1, hspace=0.55)
axC = fig.add_subplot(gsR[0])
cols = [SURFACE if p in ("HA", "NA") else INTERNAL for p in prot_order]
axC.barh(range(len(prot_order))[::-1], prot_n, color=cols, height=0.62)
axC.set_yticks(range(len(prot_order))[::-1]); axC.set_yticklabels(prot_order, fontsize=7)
axC.set_xlabel("Candidate count"); axC.set_xlim(0, max(prot_n) + 3)
for i, n in enumerate(prot_n):
    axC.text(n + 0.3, len(prot_order) - 1 - i, str(n), va="center", fontsize=6.5)
panel(axC, "C", "Candidates per protein")

axD = fig.add_subplot(gsR[1])
xs = [0, 1, 2, 3]
ys = [n_exact_dist.get(x, 0) for x in xs]
axD.bar(xs, ys, color=[GRAY, BLUE, ORANGE, RED], width=0.55)
axD.set_xticks(xs); axD.set_xticklabels(["0", "1", "2", "3"])
axD.set_xlabel("Exact subtypes conserved"); axD.set_ylabel("Candidates")
for x, y in zip(xs, ys):
    axD.text(x, y + 1.2, str(y), ha="center", fontsize=7)
panel(axD, "D", "Cross-subtype exact conservation")

axE = fig.add_subplot(gsR[2])
rank = [r["rank"] for r in top15]
score = [r["final_score"] for r in top15]
ecol = [SURFACE if r["protein"] in ("HA", "NA") else INTERNAL for r in top15]
axE.vlines(rank, 0, score, color=GRAY, lw=0.6, zorder=1)
axE.scatter(rank, score, s=30, color=ecol, zorder=2, edgecolor="white", linewidth=0.4)
axE.set_xticks(rank); axE.set_xticklabels(rank, fontsize=6)
axE.set_xlabel("Rank"); axE.set_ylabel("Composite score")
axE.set_xlim(0.5, 15.5)
panel(axE, "E", "Top 15 candidates by composite score")

fig.savefig(OUT, dpi=300, bbox_inches="tight", facecolor="white", pad_inches=0.03)
plt.close(fig)
print("saved", OUT)
```
