## Supplementary material for "Conserved influenza A epitope candidate regions and a benchmark of ESM-2 sequence features": Li_et_al_Supplementary_Data: make_fig2_stability.py

```
#!/usr/bin/env python3
"""
Figure 2：候选的时间与谱系稳定性（4 面板 A–D）。
A H1N1 代表候选 2006–2017 年度保守率折线（点大小∝序列数）；
B H1N1 六个 clade 保守率点图（附 n）；
C H3N2 十个表面候选×六 clade 热图（已核查：全零=亚型特异）；
D 候选跨时间/谱系/亚型稳定性总结。
"""
import csv, re
from collections import Counter, defaultdict
import numpy as np
import matplotlib
matplotlib.use("Agg")
import matplotlib.pyplot as plt
from Bio import SeqIO
from Bio.Seq import Seq

ROOT = "/media/qxl/My_Passport/流行病学数据/广谱流感疫苗候选课题"
RAW = f"{ROOT}/01_原始序列与分层数据/序列与地区分层/全球/序列/综合"
MS = f"{ROOT}/07_MANUSCRIPT_STRENGTHENING"
OUT = f"{MS}/figures/main/Fig2_stability.png"

BLUE = "#3A6EA5"; ORANGE = "#C77C3F"; PURPLE = "#7B5AA6"
SURFACE = "#1B9E8F"; RED = "#B5453B"; GRAY = "#8A8F98"; INK = "#22262B"
plt.rcParams.update({
    "font.family": "Arial", "font.size": 8, "axes.spines.top": False,
    "axes.spines.right": False, "axes.linewidth": 0.6, "axes.edgecolor": INK,
    "axes.labelcolor": INK, "text.color": INK, "xtick.color": INK, "ytick.color": INK,
    "xtick.direction": "out", "ytick.direction": "out",
    "xtick.major.width": 0.6, "ytick.major.width": 0.6,
})

def panel(ax, label, title):
    ax.text(-0.08, 1.06, label, transform=ax.transAxes, fontsize=13, fontweight="bold",
            va="bottom", ha="left")
    ax.set_title(title, loc="left", fontsize=9, pad=10)

def best_identity(seq, epi):
    if len(seq) < len(epi): return 0.0
    b = 0.0
    for i in range(len(seq) - len(epi) + 1):
        b = max(b, sum(a == c for a, c in zip(epi, seq[i:i+len(epi)])) / len(epi))
    return b

def frac(seqs, epi, thresh=0.9):
    if not seqs: return np.nan
    return np.mean([best_identity(s, epi) >= thresh for s in seqs])

def trans_atg(seq):
    s = str(seq).upper().replace("-", "")
    for i in range(len(s) - 2):
        if s[i:i+3] == "ATG":
            n = (len(s) - i) // 3; aa = []
            for c in range(n):
                cod = s[i+3*c:i+3*c+3]
                aa.append("X" if (len(cod) < 3 or "N" in cod) else str(Seq(cod).translate()))
            return "".join(aa)
    return ""

epi = "CKLRGVAPLHLGKCNIAGW"

# ---- A：逐年 ----
ys = defaultdict(list)
for r in SeqIO.parse(f"{RAW}/分类序列_H1N1/HA_sequences.fasta", "fasta"):
    mm = re.search(r"/20\d\d", r.description)
    if mm and 2006 <= int(mm.group(0)[1:]) <= 2017:
        ys[int(mm.group(0)[1:])].append(str(r.seq[:(len(r.seq)//3)*3].translate()))
years = sorted(ys); fracs = [frac(ys[y], epi) for y in years]; ns = [len(ys[y]) for y in years]

# ---- B：六 clade ----
meta = {}
for r in csv.DictReader(open(f"{ROOT}/99_历史项目与参考/早期H1N1系统发育项目/清洗数据/cleaned_metadata.csv")):
    meta[r["Isolate_Id"]] = r
aln = f"{ROOT}/99_历史项目与参考/早期H1N1系统发育项目/清洗数据/clustalo-H1N1-HA.fa"
by_clade = defaultdict(list)
for r in SeqIO.parse(aln, "fasta"):
    cl = meta.get(r.id, {}).get("Clade", "unassigned")
    aa = trans_atg(r.seq)
    if aa and aa.count("X") <= 0.05 * len(aa):
        by_clade[cl].append(aa)
clades = [c for c, _ in Counter({k: len(v) for k, v in by_clade.items()}).most_common(6)]
cf_clade = [frac(by_clade[c], epi) for c in clades]
n_clade = [len(by_clade[c]) for c in clades]

# ---- C：H3N2 ----
h3 = list(csv.DictReader(open(f"{MS}/09_H3N2_CROSS_LINEAGE/h3n2_cross_lineage.tsv"), delimiter="\t"))
ckeys = [k for k in h3[0].keys() if k.startswith("clade_")]
cnames = [k.replace("clade_", "") for k in ckeys]
h3mat = np.array([[float(r[k]) if r[k] not in ("", None) else 0.0 for k in ckeys] for r in h3])
h3lab = [f"{r['protein']} {r['epitope'][:10]}" for r in h3]

# ================================================================ 图
fig = plt.figure(figsize=(11, 7.5))
gs = fig.add_gridspec(2, 2, hspace=0.45, wspace=0.35)

# A 逐年（点大小∝n）
axA = fig.add_subplot(gs[0, 0])
size = 15 + 40 * (np.array(ns) / max(ns))   # 点大小映射序列数
axA.scatter(years, fracs, s=size, color=BLUE, alpha=0.85, edgecolor="white", linewidth=0.4, zorder=3)
axA.plot(years, fracs, color=BLUE, lw=1.0, alpha=0.5, zorder=2)
slope, inter, *_ = __import__("scipy.stats", fromlist=["linregress"]).linregress(years, fracs)
axA.plot(years, [slope*y+inter for y in years], "--", color=GRAY, lw=1)
axA.set_xlabel("Year"); axA.set_ylabel("Fraction ≥ 90% identity")
axA.set_ylim(0.4, 1.05)
axA.text(0.03, 0.05, "point size ~ n sequences", transform=axA.transAxes, fontsize=6.5, color=GRAY)
panel(axA, "A", f"H1N1 HA candidate {epi[:14]}… (2006–2017)")

# B 六 clade 点图
axB = fig.add_subplot(gs[0, 1])
yy = np.arange(6)[::-1]
axB.scatter(cf_clade, yy, s=60, color=BLUE, zorder=3)
axB.hlines(yy, 0, cf_clade, color=GRAY, lw=0.6, zorder=2)
axB.set_yticks(yy)
axB.set_yticklabels([f"{c[:16]}… (n={n})" for c, n in zip(clades, n_clade)], fontsize=6)
axB.set_xlabel("Conserved fraction"); axB.set_xlim(0, 1.05)
axB.axvline(0.9, color=GRAY, ls="--", lw=0.7)
for y, f in zip(yy, cf_clade):
    axB.text(f + 0.02, y, f"{f:.2f}", va="center", fontsize=6.5)
panel(axB, "B", "Conservation across six H1N1 clades (2015–2025)")

# C H3N2 热图（已核查标注）
axC = fig.add_subplot(gs[1, 0])
im = axC.imshow(h3mat, cmap="Blues", vmin=0, vmax=1, aspect="auto")
axC.set_xticks(range(len(cnames))); axC.set_xticklabels([c.replace(".", ".\n", 1) for c in cnames], fontsize=5)
axC.set_yticks(range(len(h3lab))); axC.set_yticklabels(h3lab, fontsize=6)
for i in range(h3mat.shape[0]):
    for j in range(h3mat.shape[1]):
        if h3mat[i, j] > 0.05:
            axC.text(j, i, f"{h3mat[i,j]:.2f}", ha="center", va="center", fontsize=5, color="white")
axC.tick_params(length=0)
cb = fig.colorbar(im, ax=axC, fraction=0.046, pad=0.03); cb.set_label("conserved fraction", fontsize=7); cb.ax.tick_params(labelsize=6)
axC.set_title("H3N2 clade conservation of 10 surface candidates\n(9/10 = 0 属亚型特异：7 个 H1N1/H5N1 来源，2 个映射问题)",
              loc="left", fontsize=7, pad=8, color=RED)
panel(axC, "C", "")

# D 稳定性总结
axD = fig.add_subplot(gs[1, 1])
dims = ["Time\n2006–2017\n(H1N1)", "Lineages\n2015–2025\n(6 clades)", "Subtype\nH3N2\n(cross)"]
vals = [np.mean(fracs), np.mean(cf_clade), 0.0]
ranges = [(np.min(fracs), np.max(fracs)), (np.min(cf_clade), np.max(cf_clade)), (0.0, 0.0)]
xs = np.arange(3)
axD.bar(xs, vals, color=[BLUE, BLUE, GRAY], width=0.5)
for x, v, r in zip(xs, vals, ranges):
    axD.plot([x, x], r, color=INK, lw=1.2)
    axD.text(x, v + 0.03, f"{v:.2f}", ha="center", fontsize=7)
axD.set_xticks(xs); axD.set_xticklabels(dims, fontsize=6.5)
axD.set_ylabel("Conserved fraction"); axD.set_ylim(0, 1.15)
axD.set_title("Candidate conserved within H1N1 (time & lineage)\nbut absent across subtype (H3N2)",
              loc="left", fontsize=7, pad=8, color=INK)
panel(axD, "D", "Stability summary")

fig.savefig(OUT, dpi=300, bbox_inches="tight", facecolor="white", pad_inches=0.03)
plt.close(fig)
print("saved", OUT)
print(f"A 逐年: 2006-2017 fraction 范围 {min(fracs):.2f}-{max(fracs):.2f}, 序列数 {min(ns)}-{max(ns)}")
print(f"B 六 clade: fraction {[round(f,3) for f in cf_clade]}")
```
