## Supplementary material for "Conserved influenza A epitope candidate regions and a benchmark of ESM-2 sequence features": Li_et_al_Supplementary_Data: make_fig2_v2.py

```
#!/usr/bin/env python3
"""Figure 2 — Temporal and lineage stability（3 panel，figure1_redesign 风格）。"""
import csv, re
from collections import Counter, defaultdict
import numpy as np
import matplotlib
matplotlib.use("Agg")
import matplotlib.pyplot as plt
from Bio import SeqIO
from Bio.Seq import Seq
from scipy import stats
import fig_style as FS

ROOT = "/media/qxl/My_Passport/流行病学数据/广谱流感疫苗候选课题"
RAW = f"{ROOT}/01_原始序列与分层数据/序列与地区分层/全球/序列/综合"
MS = f"{ROOT}/07_MANUSCRIPT_STRENGTHENING"
FIG = f"{MS}/figures/main"

FS.setup_style()

def best_identity(seq, epi):
    if len(seq) < len(epi): return 0.0
    return max(sum(a == c for a, c in zip(epi, seq[i:i+len(epi)])) / len(epi)
               for i in range(len(seq) - len(epi) + 1))
def frac(seqs, epi, thresh=0.9):
    if not seqs: return np.nan
    return np.mean([best_identity(s, epi) >= thresh for s in seqs])
def trans_atg(seq):
    s = str(seq).upper().replace("-", "")
    for i in range(len(s) - 2):
        if s[i:i+3] == "ATG":
            n = (len(s) - i) // 3; aa = []
            for c in range(n):
                cod = s[i+3*c:i+3*c+3]
                aa.append("X" if (len(cod) < 3 or "N" in cod) else str(Seq(cod).translate()))
            return "".join(aa)
    return ""

EPI = "CKLRGVAPLHLGKCNIAGW"

# ---- 数据 ----
ys = defaultdict(list)
for r in SeqIO.parse(f"{RAW}/分类序列_H1N1/HA_sequences.fasta", "fasta"):
    mm = re.search(r"/20\d\d", r.description)
    if mm and 2006 <= int(mm.group(0)[1:]) <= 2017:
        ys[int(mm.group(0)[1:])].append(str(r.seq[:(len(r.seq)//3)*3].translate()))
years = sorted(ys); ns = [len(ys[y]) for y in years]; fracs = [frac(ys[y], EPI) for y in years]

meta = {}
for r in csv.DictReader(open(f"{ROOT}/99_历史项目与参考/早期H1N1系统发育项目/清洗数据/cleaned_metadata.csv")):
    meta[r["Isolate_Id"]] = r
aln = f"{ROOT}/99_历史项目与参考/早期H1N1系统发育项目/清洗数据/clustalo-H1N1-HA.fa"
by_clade = defaultdict(list)
for r in SeqIO.parse(aln, "fasta"):
    cl = meta.get(r.id, {}).get("Clade", "unassigned")
    aa = trans_atg(r.seq)
    if aa and aa.count("X") <= 0.05 * len(aa):
        by_clade[cl].append(aa)
clades = [c for c, _ in Counter({k: len(v) for k, v in by_clade.items()}).most_common(6)]
n_clade = [len(by_clade[c]) for c in clades]
cf_clade = [frac(by_clade[c], EPI) for c in clades]

# ---- 图 ----
fig = plt.figure(figsize=(11, 7.5))
gs = fig.add_gridspec(2, 2, height_ratios=[1, 0.75], hspace=0.6, wspace=0.4)

# A 时间
ax = fig.add_subplot(gs[0, 0])
size = 14 + 42 * (np.array(ns) / max(ns))
ax.scatter(years, fracs, s=size, color=FS.BLUE, alpha=0.85, edgecolor="white", linewidth=0.4, zorder=3)
ax.plot(years, fracs, color=FS.BLUE_LIGHT, lw=1.0, alpha=0.7, zorder=2)
slope, inter, *_ = stats.linregress(years, fracs)
ax.plot(years, [slope*y+inter for y in years], "--", color=FS.GRAY, lw=1.0)
ax.axhline(0.9, color=FS.GRAY, lw=0.6, ls=":")
ax.set_xlabel("Year"); ax.set_ylabel("Conserved fraction"); ax.set_ylim(0, 1.1)
ax.set_title("point size reflects sequence count", fontsize=6.5, color=FS.GRAY, loc="right", pad=2)
FS.panel(ax, "A", "Temporal conservation of the selected HA candidate")

# B 谱系
ax = fig.add_subplot(gs[0, 1])
yy = np.arange(len(clades))[::-1]
ax.hlines(yy, 0, cf_clade, color=FS.GRAY_LIGHT, lw=1.2, zorder=1)
ax.scatter(cf_clade, yy, s=70, color=FS.BLUE, zorder=3)
ax.axvline(0.9, color=FS.GRAY, lw=0.6, ls=":")
ax.set_yticks(yy); ax.set_yticklabels([f"{c} (n={n})" for c, n in zip(clades, n_clade)], fontsize=6.5)
ax.set_xlabel("Conserved fraction"); ax.set_xlim(0, 1.08)
for y, f in zip(yy, cf_clade):
    ax.text(f + 0.015, y, f"{f:.2f}", va="center", fontsize=6.5)
FS.panel(ax, "B", "Conservation across circulating H1N1 lineages")

# C 总结（横跨）
ax = fig.add_subplot(gs[1, :])
dims = ["Temporal stability", "Lineage stability", "Cross-subtype support"]
means = [np.mean(fracs), np.mean(cf_clade), 0.0]
sds = [np.std(fracs), np.std(cf_clade), np.nan]
for i, (d, m, sd) in enumerate(zip(dims, means, sds)):
    ax.errorbar(m, i, xerr=sd if not np.isnan(sd) else 0, fmt="o", color=FS.BLUE, ms=7, lw=1.4, capsize=4)
    ax.text(m, i + 0.14, f"{m:.2f}" + (f" ± {sd:.2f}" if not np.isnan(sd) else ""), ha="left", fontsize=6.5)
ax.set_yticks(range(3)); ax.set_yticklabels(dims, fontsize=7)
ax.set_xlabel("Conserved fraction"); ax.set_xlim(-0.05, 1.15)
ax.axvline(0.9, color=FS.GRAY, lw=0.6, ls=":")
FS.panel(ax, "C", "Stability summary")

FS.save(fig, f"{FIG}/Figure2")
print("F2 逐年:", years, "frac", [round(f, 3) for f in fracs])
print("F2 六 clade:", clades, "frac", [round(f, 3) for f in cf_clade])
```
