## Supplementary material for "Conserved influenza A epitope candidate regions and a benchmark of ESM-2 sequence features": Li_et_al_Supplementary_Data: make_fig3_pll_conservation.py

```
#!/usr/bin/env python3
"""
Figure 3：ESM-2 PLL 与序列保守性的互补关系（5 面板 A–E）。
A–C：三亚型 HA 的 MSA conservation vs z-PLL 六边形密度散点图（标注 Spearman ρ）。
D：H1N1 HA 双轨图（上轨 MSA conservation，下轨 z-PLL）。
E：四象限图，标出「序列保守但 PLL 低」与「序列变化但 PLL 高」的位点。
数据：ESM .npy + 原始 HA 序列（按 make_figures_nc.py 的 cons_ident 算法）。
"""
import csv, os
from collections import Counter
import numpy as np
import matplotlib
matplotlib.use("Agg")
import matplotlib.pyplot as plt
from scipy import stats
from Bio import SeqIO

ROOT = "/media/qxl/My_Passport/流行病学数据/广谱流感疫苗候选课题"
ESM = f"{ROOT}/08_ESM2_AI分析与论文/results/esm2"
RAW = f"{ROOT}/01_原始序列与分层数据/序列与地区分层/全球/序列/综合"
OUT = f"{ROOT}/07_MANUSCRIPT_STRENGTHENING/figures/main/Fig3_pll_conservation.png"

H1N1 = "#3A6EA5"; H3N2 = "#C77C3F"; H5N1 = "#7B5AA6"
COL = {"H1N1": H1N1, "H3N2": H3N2, "H5N1": H5N1}
INK = "#22262B"; GRAY = "#8A8F98"
plt.rcParams.update({
    "font.family": "Arial", "font.size": 8, "axes.spines.top": False,
    "axes.spines.right": False, "axes.linewidth": 0.6, "axes.edgecolor": INK,
    "axes.labelcolor": INK, "text.color": INK, "xtick.color": INK, "ytick.color": INK,
    "xtick.direction": "out", "ytick.direction": "out",
    "xtick.major.width": 0.6, "ytick.major.width": 0.6,
})

def z(a): return (a - np.nanmean(a)) / np.nanstd(a)

def panel(ax, label, title):
    ax.text(-0.08, 1.06, label, transform=ax.transAxes, fontsize=13, fontweight="bold",
            va="bottom", ha="left")
    ax.set_title(title, loc="left", fontsize=9, pad=10)

def mode_seqs(sub, prot):
    recs = list(SeqIO.parse(f"{RAW}/分类序列_{sub}/{prot}_sequences.fasta", "fasta"))
    aa = [str(r.seq[:(len(r.seq)//3)*3].translate()) for r in recs]
    ln = Counter(len(s) for s in aa).most_common(1)[0][0]
    return [s for s in aa if len(s) == ln and s.count("X") <= 0.05 * len(s)]

def cons_ident(sub, prot):
    seqs = mode_seqs(sub, prot); L = len(seqs[0])
    c = [Counter(s[i] for s in seqs).most_common(1)[0][0] for i in range(L)]
    return np.array([sum(1 for s in seqs if s[i] == c[i]) / len(seqs) for i in range(L)])

def get_data(sub):
    pll = np.load(f"{ESM}/{sub}_HA_pll.npy")
    seq = open(f"{ESM}/{sub}_HA_seq.txt").read().strip()
    ci = cons_ident(sub, "HA")
    n = min(len(seq), len(ci))
    zz = z(pll[1:1+n])
    return ci[:n], zz, seq

# ================================================================ 图
fig = plt.figure(figsize=(11, 8.5))
gs = fig.add_gridspec(2, 3, height_ratios=[1, 1], hspace=0.45, wspace=0.35,
                      width_ratios=[1, 1, 1])

# A–C：六边形密度散点
for idx, sub in enumerate(["H1N1", "H3N2", "H5N1"]):
    ci, zz, seq = get_data(sub)
    rho, _ = stats.spearmanr(ci, zz)
    rp, _ = stats.pearsonr(ci, zz)
    ax = fig.add_subplot(gs[0, idx])
    hb = ax.hexbin(ci, zz, gridsize=40, cmap="Blues", mincnt=1, bins="log")
    ax.set_xlabel("MSA positional identity"); ax.set_ylabel("ESM-2 z-PLL")
    ax.text(0.03, 0.96, f"{sub}\nSpearman ρ = {rho:.3f}\nPearson r = {rp:.3f}",
            transform=ax.transAxes, fontsize=8, va="top", color=COL[sub], fontweight="bold")
    cb = fig.colorbar(hb, ax=ax, fraction=0.046, pad=0.03); cb.set_label("count", fontsize=7); cb.ax.tick_params(labelsize=6)
    panel(ax, "ABC"[idx], f"{sub} HA: conservation vs PLL")

# D：双轨图（H1N1 HA）
axD = fig.add_subplot(gs[1, :2])
ci, zz, seq = get_data("H1N1")
pos = np.arange(len(ci)) + 1
axD.plot(pos, ci, color=GRAY, lw=1.0, label="MSA conservation")
axD.set_xlabel("HA consensus position"); axD.set_ylabel("MSA identity", color=GRAY)
axD.set_ylim(0, 1.05)
axD.tick_params(axis="y", labelcolor=GRAY)
axD2 = axD.twinx()
axD2.plot(pos, zz, color=H1N1, lw=0.8, alpha=0.8, label="z-PLL")
axD2.set_ylabel("ESM-2 z-PLL", color=H1N1); axD2.tick_params(axis="y", labelcolor=H1N1)
axD2.spines["top"].set_visible(False)
lines = axD.get_lines() + axD2.get_lines()
axD.legend(lines, [l.get_label() for l in lines], frameon=False, fontsize=7, loc="upper right")
panel(axD, "D", "H1N1 HA: per-position conservation vs z-PLL")

# E：四象限图（H1N1 HA）
axE = fig.add_subplot(gs[1, 2])
ci, zz, seq = get_data("H1N1")
x_thresh = np.median(ci)   # 保守性中位数
y_thresh = 0.0             # z-PLL 自然分界
axE.axhline(y_thresh, color=INK, lw=0.7, ls="--")
axE.axvline(x_thresh, color=INK, lw=0.7, ls="--")
# 四个象限
Q = {"Q1": (ci >= x_thresh) & (zz >= y_thresh),   # 保守 + PLL 高
     "Q2": (ci <  x_thresh) & (zz >= y_thresh),   # 变化 + PLL 高（关注）
     "Q3": (ci <  x_thresh) & (zz <  y_thresh),   # 变化 + PLL 低
     "Q4": (ci >= x_thresh) & (zz <  y_thresh)}   # 保守 + PLL 低（关注）
axE.scatter(ci[Q["Q1"]], zz[Q["Q1"]], s=3, color="#C8CDD3", lw=0, alpha=0.4)
axE.scatter(ci[Q["Q3"]], zz[Q["Q3"]], s=3, color="#C8CDD3", lw=0, alpha=0.4)
axE.scatter(ci[Q["Q2"]], zz[Q["Q2"]], s=6, color=H5N1, lw=0, alpha=0.7, label="variable + high PLL")
axE.scatter(ci[Q["Q4"]], zz[Q["Q4"]], s=6, color=H3N2, lw=0, alpha=0.7, label="conserved + low PLL")
axE.set_xlabel("MSA positional identity"); axE.set_ylabel("ESM-2 z-PLL")
axE.legend(frameon=False, fontsize=6.5, loc="lower right")
axE.text(0.97, 0.03, f"n(Q2)={int(Q['Q2'].sum())}\nn(Q4)={int(Q['Q4'].sum())}",
         transform=axE.transAxes, ha="right", va="bottom", fontsize=6.5)
panel(axE, "E", "Quadrant view (H1N1 HA)")

fig.savefig(OUT, dpi=300, bbox_inches="tight", facecolor="white", pad_inches=0.03)
plt.close(fig)
print("saved", OUT)
print("相关系数（Spearman/Pearson）:")
for sub in ["H1N1", "H3N2", "H5N1"]:
    ci, zz, _ = get_data(sub)
    print(f"  {sub}: Spearman ρ={stats.spearmanr(ci,zz)[0]:.3f}, Pearson r={stats.pearsonr(ci,zz)[0]:.3f}")
```
