## Supplementary material for "Conserved influenza A epitope candidate regions and a benchmark of ESM-2 sequence features": Li_et_al_Supplementary_Data: make_fig3_v2.py

```
#!/usr/bin/env python3
"""Figure 3 — Conservation and PLL relationship（3 panel，figure1_redesign 风格）。"""
import csv
from collections import Counter
import numpy as np
import matplotlib
matplotlib.use("Agg")
import matplotlib.pyplot as plt
from Bio import SeqIO
from scipy import stats
import fig_style as FS

ROOT = "/media/qxl/My_Passport/流行病学数据/广谱流感疫苗候选课题"
ESM = f"{ROOT}/08_ESM2_AI分析与论文/results/esm2"
RAW = f"{ROOT}/01_原始序列与分层数据/序列与地区分层/全球/序列/综合"
MS = f"{ROOT}/07_MANUSCRIPT_STRENGTHENING"
FIG = f"{MS}/figures/main"

FS.setup_style()

def z(a): return (a - np.nanmean(a)) / np.nanstd(a)

def mode_seqs(sub, prot):
    recs = list(SeqIO.parse(f"{RAW}/分类序列_{sub}/{prot}_sequences.fasta", "fasta"))
    aa = [str(r.seq[:(len(r.seq)//3)*3].translate()) for r in recs]
    ln = Counter(len(s) for s in aa).most_common(1)[0][0]
    return [s for s in aa if len(s) == ln and s.count("X") <= 0.05 * len(s)]

def cons_ident(sub, prot):
    seqs = mode_seqs(sub, prot); L = len(seqs[0])
    c = [Counter(s[i] for s in seqs).most_common(1)[0][0] for i in range(L)]
    return np.array([sum(1 for s in seqs if s[i] == c[i]) / len(seqs) for i in range(L)])

def get_data(sub):
    pll = np.load(f"{ESM}/{sub}_HA_pll.npy")
    seq = open(f"{ESM}/{sub}_HA_seq.txt").read().strip()
    ci = cons_ident(sub, "HA"); n = min(len(seq), len(ci))
    return ci[:n], z(pll[1:1+n]), seq

data = {sub: get_data(sub) for sub in ["H1N1", "H3N2", "H5N1"]}
for sub, (ci, zz, seq) in data.items():
    print(f"{sub}: Spearman={stats.spearmanr(ci, zz)[0]:.3f}, Pearson={stats.pearsonr(ci, zz)[0]:.3f}")

fig = plt.figure(figsize=(11, 9))
gs = fig.add_gridspec(2, 2, height_ratios=[1, 1.1], hspace=0.5, wspace=0.35, width_ratios=[2.6, 1.2])

# A 三个 facet + shared colorbar
gsA = gs[0, :].subgridspec(1, 3, wspace=0.25)
axesA = [fig.add_subplot(gsA[i]) for i in range(3)]
for ax, sub in zip(axesA, ["H1N1", "H3N2", "H5N1"]):
    ci, zz, _ = data[sub]
    hb = ax.hexbin(ci, zz, gridsize=40, cmap="Blues", mincnt=1, bins="log")
    rho, _ = stats.spearmanr(ci, zz); rp, _ = stats.pearsonr(ci, zz)
    ax.text(0.03, 0.97, f"{sub}\nSpearman ρ = {rho:.2f}\nPearson r = {rp:.2f}",
            transform=ax.transAxes, fontsize=7.5, va="top", color=FS.TEXT)
    ax.set_xlabel("MSA positional identity"); ax.set_ylabel("ESM-2 z-PLL")
    ax.set_xlim(0.3, 1.02); ax.set_ylim(-3.2, 3.2)
cb = fig.colorbar(hb, ax=axesA, fraction=0.03, pad=0.02)
cb.set_label("Density (log count)", fontsize=7); cb.ax.tick_params(labelsize=6)
FS.panel(axesA[0], "A", "Conservation–PLL relationship across HA subtypes")

# B 双轨
gsB = gs[1, 0].subgridspec(2, 1, hspace=0.12)
axB1 = fig.add_subplot(gsB[0]); axB2 = fig.add_subplot(gsB[1], sharex=axB1)
ci, zz, seq = data["H1N1"]; pos = np.arange(len(ci)) + 1
axB1.plot(pos, ci, color=FS.BLUE, lw=1.0)
axB1.set_ylabel("MSA identity"); axB1.set_ylim(0, 1.05); axB1.tick_params(labelbottom=False)
axB2.plot(pos, zz, color=FS.BLUE_DARK, lw=0.8)
axB2.axhline(0, color=FS.GRAY, lw=0.6, ls=":")
axB2.set_ylabel("ESM-2 z-PLL"); axB2.set_xlabel("HA consensus position")
FS.panel(axB1, "B", "Position-resolved conservation and PLL landscape")

# C 四象限
axC = fig.add_subplot(gs[1, 1])
ci, zz, _ = data["H1N1"]
x_thresh = np.median(ci); y_thresh = 0.0
axC.axhline(y_thresh, color=FS.GRAY, lw=0.6, ls=":")
axC.axvline(x_thresh, color=FS.GRAY, lw=0.6, ls=":")
Q_hi_hi = (ci >= x_thresh) & (zz >= y_thresh)
Q_hi_lo = (ci >= x_thresh) & (zz < y_thresh)
Q_lo = ~(Q_hi_hi | Q_hi_lo)
axC.scatter(ci[Q_lo], zz[Q_lo], s=4, color=FS.GRAY_LIGHT, lw=0, alpha=0.6)
axC.scatter(ci[Q_hi_hi], zz[Q_hi_hi], s=8, color=FS.BLUE, lw=0, alpha=0.7, label="conserved + favorable PLL")
axC.scatter(ci[Q_hi_lo], zz[Q_hi_lo], s=8, color=FS.ORANGE, lw=0, alpha=0.8, label="conserved + low PLL")
axC.set_xlabel("MSA positional identity"); axC.set_ylabel("ESM-2 z-PLL")
axC.legend(frameon=False, fontsize=6, loc="lower left")
FS.panel(axC, "C", "High-conservation candidate space")

FS.save(fig, f"{FIG}/Figure2")
```
