## Supplementary material for "Conserved influenza A epitope candidate regions and a benchmark of ESM-2 sequence features": Li_et_al_Supplementary_Data: make_fig4_benchmark.py

```
#!/usr/bin/env python3
"""
Figure 4：T细胞与B细胞外部基准验证（5 面板 A–E）。
A T细胞 AUROC 森林图；B AUROC/AUPRC/Precision@10/20 点图；
C B细胞三类 mean z-PLL（箱线+抖动散点，显示所有原始点）；D 数据流；E 效应量。
"""
import csv
import numpy as np
import matplotlib
matplotlib.use("Agg")
import matplotlib.pyplot as plt
from Bio import SeqIO
from scipy import stats

ROOT = "/media/qxl/My_Passport/流行病学数据/广谱流感疫苗候选课题"
ESM = f"{ROOT}/08_ESM2_AI分析与论文/results/esm2"
CONS = f"{ROOT}/08_ESM2_AI分析与论文/data/consensus"
MS = f"{ROOT}/07_MANUSCRIPT_STRENGTHENING"
OUT = f"{MS}/figures/main/Fig4_benchmark.png"

BLUE = "#3A6EA5"; ORANGE = "#C77C3F"; PURPLE = "#7B5AA6"
GRAY = "#8A8F98"; INK = "#22262B"; BIND = "#E8A87C"; NEUT = "#B5453B"
plt.rcParams.update({
    "font.family": "Arial", "font.size": 8, "axes.spines.top": False,
    "axes.spines.right": False, "axes.linewidth": 0.6, "axes.edgecolor": INK,
    "axes.labelcolor": INK, "text.color": INK, "xtick.color": INK, "ytick.color": INK,
    "xtick.direction": "out", "ytick.direction": "out",
    "xtick.major.width": 0.6, "ytick.major.width": 0.6,
})

def z(a): return (a - np.nanmean(a)) / np.nanstd(a)

def panel(ax, label, title):
    ax.text(-0.08, 1.06, label, transform=ax.transAxes, fontsize=13, fontweight="bold",
            va="bottom", ha="left")
    ax.set_title(title, loc="left", fontsize=9, pad=10)

# ================================================================ T细胞 benchmark
lines = open(f"{MS}/01_ESM_INCREMENTAL_BENCHMARK/benchmark_results.tsv").read().splitlines()
m = {}
for l in lines[1:]:
    p = l.split("\t")
    if p[0].startswith("incremental"): continue
    m[p[0]] = dict(auc=float(p[1]), lo=float(p[2]), hi=float(p[3]),
                   auprc=float(p[4]), p10=float(p[5]), p20=float(p[6]))
models = ["A_conservation", "C_conservation+PLL", "D_conservation+PLL+contact"]
mlab = ["conservation", "+ PLL", "+ PLL\n+ contact"]

# ================================================================ B细胞 mean z-PLL
cons = {}
for sub in ["H1N1", "H3N2", "H5N1"]:
    for prot in ["HA", "NA"]:
        cons[(sub, prot)] = str(list(SeqIO.parse(f"{CONS}/{sub}_{prot}.fasta", "fasta"))[0].seq)
def region_pll(sub, prot, seq):
    s = cons[(sub, prot)]; i = s.find(seq)
    if i < 0:
        for j in range(len(s) - len(seq) + 1):
            if sum(a != b for a, b in zip(seq, s[j:j+len(seq)])) <= 1: i = j; break
    if i < 0: return np.nan
    pll = np.load(f"{ESM}/{sub}_{prot}_pll.npy"); zz = z(pll)
    return np.mean(zz[i+1:i+1+len(seq)])

c06 = list(csv.DictReader(open(f"{MS}/08_BCELL_EXTERNAL_BENCHMARK/06_BENCHMARK_READY_DATASET.tsv"), delimiter="\t"))
bg = list(csv.DictReader(open(f"{MS}/08_BCELL_EXTERNAL_BENCHMARK/07_MATCHED_BACKGROUND.tsv"), delimiter="\t"))
neut, bind, bgr = [], [], []
for c in c06:
    v = region_pll(c["subtype"], c["protein"], c["representative_seq"])
    if np.isnan(v): continue
    (neut if c["label_level"] == "L3_neutralization" else bind).append(v)
for r in bg:
    v = region_pll(r["subtype"], r["protein"], r["bg_sequence"])
    if not np.isnan(v): bgr.append(v)
neut = np.array(neut); bind = np.array(bind); bgr = np.array(bgr)

# ================================================================ 图
fig = plt.figure(figsize=(11.5, 8.5))
gs = fig.add_gridspec(2, 3, height_ratios=[1, 1.25], hspace=0.5, wspace=0.4)

# A 森林图
axA = fig.add_subplot(gs[0, 0])
ypos = [2, 1, 0]
for y, k in zip(ypos, models):
    r = m[k]
    axA.errorbar(r["auc"], y, xerr=[[r["auc"]-r["lo"]], [r["hi"]-r["auc"]]],
                 fmt="o", color=[GRAY, BLUE, ORANGE][ypos.index(y)], ms=6, lw=1.5, capsize=3)
axA.axvline(0.5, color=INK, ls="--", lw=0.7)
axA.set_yticks(ypos); axA.set_yticklabels(mlab, fontsize=7)
axA.set_xlabel("AUROC"); axA.set_xlim(0.35, 0.7)
axA.set_title("", loc="left")
panel(axA, "A", "T-cell benchmark (AUROC ± 95% CI)")

# B 点图
axB = fig.add_subplot(gs[0, 1])
metrics = ["auprc", "p10", "p20"]
x = np.arange(3)
for i, k in enumerate(models):
    r = m[k]
    axB.plot(x, [r["auprc"], r["p10"], r["p20"]], "o-", color=[GRAY, BLUE, ORANGE][i], lw=1.2, ms=5, label=mlab[i].replace("\n", " "))
axB.set_xticks(x); axB.set_xticklabels(["AUPRC", "Precision@10", "Precision@20"], fontsize=7)
axB.set_ylabel("Value"); axB.set_ylim(0, 1)
axB.legend(frameon=False, fontsize=6.5)
panel(axB, "B", "Auxiliary metrics by model")

# C 雨云（箱线 + 抖动散点，显示所有原始点）
axC = fig.add_subplot(gs[1, :2])
groups = [("Background", bgr, GRAY), ("Binding", bind, BIND), ("Neutralizing", neut, NEUT)]
for i, (name, vals, c) in enumerate(groups):
    bp = axC.boxplot(vals, positions=[i], widths=0.5, patch_artist=True,
                     boxprops=dict(facecolor=c, alpha=0.35, lw=0.8),
                     medianprops=dict(color=INK, lw=1.2),
                     whiskerprops=dict(lw=0.8), capprops=dict(lw=0.8),
                     flierprops=dict(markersize=0))
    jitter = np.random.default_rng(i).normal(0, 0.06, len(vals))
    axC.scatter(i + jitter, vals, s=12, color=c, alpha=0.75, edgecolor="white", linewidth=0.3, zorder=3)
    axC.text(i, np.mean(vals), f"  mean {np.mean(vals):+.2f}\n  n={len(vals)}",
             va="center", fontsize=7, color=INK)
axC.axhline(0, color=INK, ls="--", lw=0.6)
axC.set_xticks(range(3)); axC.set_xticklabels([g[0] for g in groups], fontsize=8)
axC.set_ylabel("ESM-2 mean z-PLL")
panel(axC, "C", "B-cell epitopes: mean z-PLL by class (all points shown)")

# D 数据流
axD = fig.add_subplot(gs[0, 2])
stages = [("IEDB\nB-cell", 122), ("HA/NA\nsurface", 70), ("Mapped\n(PASS+AMB)", 33), ("Clusters", 22)]
xs = np.arange(4)
axD.bar(xs, [s[1] for s in stages], color=["#5B6672", "#7A8794", ORANGE, NEUT], width=0.55)
axD.set_xticks(xs); axD.set_xticklabels([s[0] for s in stages], fontsize=6.5)
axD.set_ylabel("Count")
for x, (_, n) in zip(xs, stages):
    axD.text(x, n + 2, str(n), ha="center", fontsize=7)
panel(axD, "D", "B-cell data flow")

# E 效应量（Cohen's d vs Background）
axE = fig.add_subplot(gs[1, 2])
def cohens_d(a, b):
    na, nb = len(a), len(b)
    sp = np.sqrt(((na-1)*np.var(a, ddof=1) + (nb-1)*np.var(b, ddof=1)) / (na+nb-2))
    return (np.mean(a) - np.mean(b)) / sp
def boot_d_ci(a, b, n=2000, seed=0):
    rng = np.random.default_rng(seed)
    ds = []
    for _ in range(n):
        aa = rng.choice(a, len(a), replace=True); bb = rng.choice(b, len(b), replace=True)
        ds.append(cohens_d(aa, bb))
    return np.percentile(ds, [2.5, 97.5])
labels = ["Binding vs\nBackground", "Neutralizing vs\nBackground"]
d_bind = cohens_d(bind, bgr); ci_bind = boot_d_ci(bind, bgr)
d_neut = cohens_d(neut, bgr); ci_neut = boot_d_ci(neut, bgr)
ds = [d_bind, d_neut]; cis = [ci_bind, ci_neut]
for i, (d, ci) in enumerate(zip(ds, cis)):
    axE.errorbar(d, i, xerr=[[d-ci[0]], [ci[1]-d]], fmt="o", color=[BIND, NEUT][i], ms=7, lw=1.5, capsize=3)
    axE.text(d, i + 0.12, f"  d={d:.2f} [{ci[0]:.2f}, {ci[1]:.2f}]", fontsize=6.5, ha="left")
axE.axvline(0, color=INK, ls="--", lw=0.7)
axE.set_yticks(range(2)); axE.set_yticklabels(labels, fontsize=7)
axE.set_xlabel("Cohen's d (95% CI)")
panel(axE, "E", "Effect size vs matched background")

fig.savefig(OUT, dpi=300, bbox_inches="tight", facecolor="white", pad_inches=0.03)
plt.close(fig)
print("saved", OUT)
print(f"Background: n={len(bgr)}, mean={np.mean(bgr):+.3f}")
print(f"Binding: n={len(bind)}, mean={np.mean(bind):+.3f}, d={d_bind:.2f}")
print(f"Neutralizing: n={len(neut)}, mean={np.mean(neut):+.3f}, d={d_neut:.2f}")
```
