## Supplementary material for "Conserved influenza A epitope candidate regions and a benchmark of ESM-2 sequence features": Li_et_al_Supplementary_Data: make_fig4_v2.py

```
#!/usr/bin/env python3
"""Figure 4 — T-cell and B-cell benchmark（5 panel，figure1_redesign 风格）。"""
import csv
import numpy as np
import matplotlib
matplotlib.use("Agg")
import matplotlib.pyplot as plt
from Bio import SeqIO
from scipy import stats
from scipy.stats import gaussian_kde
import fig_style as FS

ROOT = "/media/qxl/My_Passport/流行病学数据/广谱流感疫苗候选课题"
ESM = f"{ROOT}/08_ESM2_AI分析与论文/results/esm2"
CONS = f"{ROOT}/08_ESM2_AI分析与论文/data/consensus"
MS = f"{ROOT}/07_MANUSCRIPT_STRENGTHENING"
FIG = f"{MS}/figures/main"

FS.setup_style()

def z(a): return (a - np.nanmean(a)) / np.nanstd(a)

lines = open(f"{MS}/01_ESM_INCREMENTAL_BENCHMARK/benchmark_results.tsv").read().splitlines()
m = {}
for l in lines[1:]:
    p = l.split("\t")
    if p[0].startswith("incremental"): continue
    m[p[0]] = dict(auc=float(p[1]), lo=float(p[2]), hi=float(p[3]),
                   auprc=float(p[4]), p10=float(p[5]), p20=float(p[6]))
models = ["A_conservation", "C_conservation+PLL", "D_conservation+PLL+contact"]
mlab = ["Conservation", "Conservation + PLL", "Conservation + PLL + contact"]
mcolor = [FS.GRAY, FS.BLUE, FS.ORANGE]

cons = {}
for sub in ["H1N1", "H3N2", "H5N1"]:
    for prot in ["HA", "NA"]:
        cons[(sub, prot)] = str(list(SeqIO.parse(f"{CONS}/{sub}_{prot}.fasta", "fasta"))[0].seq)
def region_pll(sub, prot, seq):
    s = cons[(sub, prot)]; i = s.find(seq)
    if i < 0:
        for j in range(len(s) - len(seq) + 1):
            if sum(a != b for a, b in zip(seq, s[j:j+len(seq)])) <= 1: i = j; break
    if i < 0: return np.nan
    pll = np.load(f"{ESM}/{sub}_{prot}_pll.npy"); zz = z(pll)
    return np.mean(zz[i+1:i+1+len(seq)])
c06 = list(csv.DictReader(open(f"{MS}/08_BCELL_EXTERNAL_BENCHMARK/06_BENCHMARK_READY_DATASET.tsv"), delimiter="\t"))
bg = list(csv.DictReader(open(f"{MS}/08_BCELL_EXTERNAL_BENCHMARK/07_MATCHED_BACKGROUND.tsv"), delimiter="\t"))
neut, bind, bgr = [], [], []
for c in c06:
    v = region_pll(c["subtype"], c["protein"], c["representative_seq"])
    if np.isnan(v): continue
    (neut if c["label_level"] == "L3_neutralization" else bind).append(v)
for r in bg:
    v = region_pll(r["subtype"], r["protein"], r["bg_sequence"])
    if not np.isnan(v): bgr.append(v)
neut, bind, bgr = np.array(neut), np.array(bind), np.array(bgr)

# cluster-level paired data（以 cluster 为统计单位，避免匹配背景伪重复）
from collections import defaultdict
pos_by_cluster = {}; _bg = defaultdict(list)
for c in c06:
    v = region_pll(c["subtype"], c["protein"], c["representative_seq"])
    if not np.isnan(v): pos_by_cluster[c["cluster_id"]] = (v, c["label_level"])
for r in bg:
    v = region_pll(r["subtype"], r["protein"], r["bg_sequence"])
    if not np.isnan(v): _bg[r["cluster_id"]].append(v)
bg_by_cluster = {k: np.mean(v) for k, v in _bg.items()}

def paired_d(cids):
    diffs = np.array([pos_by_cluster[c][0] - bg_by_cluster[c] for c in cids])
    return np.mean(diffs) / np.std(diffs)
def paired_d_ci(cids, n=2000, seed=0):
    rng = np.random.default_rng(seed)
    diffs = np.array([pos_by_cluster[c][0] - bg_by_cluster[c] for c in cids])
    ds = []
    for _ in range(n):
        sm = rng.choice(diffs, len(diffs), replace=True)
        if np.std(sm) > 0: ds.append(np.mean(sm) / np.std(sm))
    return np.percentile(ds, [2.5, 97.5])

fig = plt.figure(figsize=(11, 10))
gs = fig.add_gridspec(3, 2, height_ratios=[1, 1.2, 0.9], hspace=0.6, wspace=0.4)

# A T-cell forest
ax = fig.add_subplot(gs[0, 0])
for y, k in zip([2, 1, 0], models):
    r = m[k]
    ax.errorbar(r["auc"], y, xerr=[[r["auc"]-r["lo"]], [r["hi"]-r["auc"]]], fmt="o",
                color=mcolor[y], ms=7, lw=1.6, capsize=4)
ax.axvline(0.5, color=FS.GRAY, lw=0.7, ls="--")
ax.set_yticks([2, 1, 0]); ax.set_yticklabels(mlab, fontsize=7.5)
ax.set_xlabel("AUROC"); ax.set_xlim(0.4, 0.68)
FS.panel(ax, "A", "T-cell discrimination performance")

# B auxiliary dot plot
ax = fig.add_subplot(gs[0, 1])
metrics = ["AUPRC", "Precision@10", "Precision@20"]
x = np.arange(3)
for i, k in enumerate(models):
    r = m[k]
    ax.plot(x, [r["auprc"], r["p10"], r["p20"]], "o-", color=mcolor[i], lw=1.2, ms=5)
ax.set_xticks(x); ax.set_xticklabels(metrics, fontsize=7.5)
ax.set_ylabel("Value"); ax.set_ylim(0.4, 1.0)
ax.legend(mlab, frameon=False, fontsize=6.5, loc="lower right")
FS.panel(ax, "B", "Auxiliary ranking performance")

# C B-cell raincloud
ax = fig.add_subplot(gs[1, :])
groups = [("Background", bgr, FS.GRAY), ("Binding", bind, FS.ORANGE_LIGHT), ("Neutralizing", neut, FS.RED)]
for i, (name, vals, c) in enumerate(groups):
    if len(vals) > 1:
        kde = gaussian_kde(vals)
        xg = np.linspace(vals.min(), vals.max(), 100)
        d = kde(xg); d = d / d.max() * 0.38
        ax.fill_betweenx(xg, i - d, i, color=c, alpha=0.45, lw=0)
    ax.boxplot(vals, positions=[i], widths=0.16, patch_artist=True,
               boxprops=dict(facecolor="white", lw=0.8),
               medianprops=dict(color=FS.TEXT, lw=1.3), whiskerprops=dict(lw=0.8),
               capprops=dict(lw=0.8), flierprops=dict(markersize=0))
    jitter = np.random.default_rng(i).normal(i + 0.16, 0.028, len(vals))
    ax.scatter(jitter, vals, s=11, color=c, alpha=0.6, edgecolor="white", linewidth=0.3, zorder=3)
    ax.scatter([i + 0.06], [np.mean(vals)], marker="D", s=26, color=FS.TEXT, zorder=4)
ax.axhline(0, color=FS.GRAY, lw=0.7, ls="--")
ax.set_xticks(range(3))
ax.set_xticklabels([f"{g[0]}\nn = {len(g[1])}" for g in groups], fontsize=7.5)
ax.set_ylabel("ESM-2 mean z-PLL")
FS.panel(ax, "C", "B-cell epitope score distributions")

# D funnel
ax = fig.add_subplot(gs[2, 0])
stages = [("IEDB B-cell epitopes", 122), ("HA/NA-associated epitopes", 70),
          ("Mapped epitopes", 33), ("Non-redundant clusters", 22)]
maxn = stages[0][1]
for i, (name, n) in enumerate(stages):
    w = 0.25 + 0.55 * (n / maxn)
    y = 3 - i
    ax.add_patch(plt.Rectangle((0, y - 0.3), w, 0.6, color=[FS.GRAY_LIGHT, "#C8CDD3", FS.ORANGE_LIGHT, FS.ORANGE][i], ec="none"))
    ax.text(w / 2, y, f"{n}", ha="center", va="center", fontsize=8, fontweight="bold", color=FS.TEXT)
    ax.text(w + 0.03, y, name, ha="left", va="center", fontsize=6.8, color=FS.TEXT)
ax.set_xlim(0, 1.15); ax.set_ylim(-0.6, 3.6); ax.axis("off")
FS.panel(ax, "D", "B-cell evidence flow")

# E enrichment forest
ax = fig.add_subplot(gs[2, 1])
bind_ids = [c for c in pos_by_cluster if pos_by_cluster[c][1] != "L3_neutralization"]
neut_ids = [c for c in pos_by_cluster if pos_by_cluster[c][1] == "L3_neutralization"]
d_bind = paired_d(bind_ids); ci_bind = paired_d_ci(bind_ids)
d_neut = paired_d(neut_ids); ci_neut = paired_d_ci(neut_ids)
for i, (d, ci, c, lab) in enumerate(zip([d_bind, d_neut], [ci_bind, ci_neut],
                                        [FS.ORANGE, FS.RED], ["Binding vs background", "Neutralizing vs background"])):
    ax.errorbar(d, i, xerr=[[d-ci[0]], [ci[1]-d]], fmt="o", color=c, ms=8, lw=1.6, capsize=4)
    ax.text(d, i + 0.13, f"d = {d:.2f} [{ci[0]:.2f}, {ci[1]:.2f}]", ha="left", fontsize=6.5)
ax.axvline(0, color=FS.GRAY, lw=0.7, ls="--")
ax.set_yticks([0, 1]); ax.set_yticklabels(["Binding vs background", "Neutralizing vs background"], fontsize=7)
ax.set_xlabel("Paired Cohen's d"); ax.set_xlim(-0.5, 6.0)
FS.panel(ax, "E", "Enrichment relative to matched background")

FS.save(fig, f"{FIG}/Figure3")
print(f"B-cell: Background mean={np.mean(bgr):+.3f}, Binding mean={np.mean(bind):+.3f}, Neutralizing mean={np.mean(neut):+.3f}")
print(f"Cohen's d: Binding={d_bind:.2f}, Neutralizing={d_neut:.2f}")
```
