## Supplementary material for "Conserved influenza A epitope candidate regions and a benchmark of ESM-2 sequence features": Li_et_al_Supplementary_Data: make_fig5_structure.py

```
#!/usr/bin/env python3
"""
Figure 5：结构可及性、抗体足迹与选择压力（部分内容，A–E）。
A RSA vs contact-density 密度散点；B 8 抗体足迹轨道图（按 region 着色）；
C 各抗体接触残基数；D dN/dS 轨迹（原始点 + 滑动中位数）；E 候选/足迹/dN/dS 叠加。
注：RSA 表无残基位置（A 面板 position 未知），dN/dS 平滑列空（D 面板用滑动中位数现算）。
"""
import csv
import numpy as np
import pandas as pd
import matplotlib
matplotlib.use("Agg")
import matplotlib.pyplot as plt
from scipy import stats

ROOT = "/media/qxl/My_Passport/流行病学数据/广谱流感疫苗候选课题"
RES = f"{ROOT}/08_ESM2_AI分析与论文/results"
MS = f"{ROOT}/07_MANUSCRIPT_STRENGTHENING"
OUT = f"{MS}/figures/main/Fig5_structure.png"

BLUE = "#3A6EA5"; ORANGE = "#C77C3F"; PURPLE = "#7B5AA6"
STALK_C = "#6B7B8C"; RBD_C = ORANGE; NA_C = "#1B9E8F"
GRAY = "#8A8F98"; INK = "#22262B"; RED = "#B5453B"
plt.rcParams.update({
    "font.family": "Arial", "font.size": 8, "axes.spines.top": False,
    "axes.spines.right": False, "axes.linewidth": 0.6, "axes.edgecolor": INK,
    "axes.labelcolor": INK, "text.color": INK, "xtick.color": INK, "ytick.color": INK,
    "xtick.direction": "out", "ytick.direction": "out",
    "xtick.major.width": 0.6, "ytick.major.width": 0.6,
})

def panel(ax, label, title):
    ax.text(-0.08, 1.06, label, transform=ax.transAxes, fontsize=13, fontweight="bold",
            va="bottom", ha="left")
    ax.set_title(title, loc="left", fontsize=9, pad=10)

# ================================================================ 数据
# A：RSA vs contact-density
ha = np.genfromtxt(f"{RES}/rsa_validation.csv", delimiter=",", skip_header=1)
rsa, cd = ha[:, 0], ha[:, 1]
r_sp, _ = stats.spearmanr(rsa, cd); r_pe, _ = stats.pearsonr(rsa, cd)

# B/C：足迹
fps = list(csv.DictReader(open(
    f"{MS}/08_BCELL_EXTERNAL_BENCHMARK/structures/09_NEUTRALIZING_FOOTPRINTS_EXTENDED.tsv"), delimiter="\t"))
for f in fps:
    f["pos"] = [int(x) for x in f["consensus_positions"].split(";")]
    f["n"] = len(f["pos"])
region_color = {"stalk": STALK_C, "RBD (receptor-binding site)": RBD_C, "enzyme active site": NA_C}

# D：dN/dS
dnds = list(csv.DictReader(open(f"{MS}/dnds_per_codon.tsv"), delimiter="\t"))
codon = np.array([int(r["codon_index"]) for r in dnds])
dnds_val = np.array([np.nan if r["dN_dS"] in ("", "NA") else float(r["dN_dS"]) for r in dnds])
s = pd.Series(dnds_val)
smooth = s.rolling(window=15, center=True, min_periods=1).median().to_numpy()

# ================================================================ 图
fig = plt.figure(figsize=(11.5, 7.8))
gs = fig.add_gridspec(2, 3, height_ratios=[1, 1], hspace=0.5, wspace=0.4)

# A
axA = fig.add_subplot(gs[0, 0])
hb = axA.hexbin(rsa, cd, gridsize=30, cmap="Blues", mincnt=1, bins="log")
axA.set_xlabel("RSA (FreeSASA)"); axA.set_ylabel("Contact density")
axA.text(0.03, 0.96, f"Spearman ρ = {r_sp:+.3f}\nPearson r = {r_pe:+.3f}",
         transform=axA.transAxes, fontsize=8, va="top", fontweight="bold")
cb = fig.colorbar(hb, ax=axA, fraction=0.046, pad=0.03); cb.set_label("count", fontsize=7); cb.ax.tick_params(labelsize=6)
panel(axA, "A", "HA contact-density vs RSA (weak, not a proxy)")

# B 足迹轨道图
axB = fig.add_subplot(gs[0, 1:])
ha_ab = [f for f in fps if f["protein"] == "HA"]
ha_ab.sort(key=lambda f: (f["region"] != "stalk", f["antibody"]))
HA_LEN = 567
for i, f in enumerate(ha_ab):
    y = len(ha_ab) - 1 - i
    c = region_color.get(f["region"], GRAY)
    axB.hlines(y, 0, HA_LEN, color=GRAY, lw=0.4, alpha=0.5)
    axB.vlines(f["pos"], y - 0.25, y + 0.25, color=c, lw=1.2)
axB.set_yticks(range(len(ha_ab)))
axB.set_yticklabels([f"{f['antibody']} ({f['subtype']})" for f in ha_ab], fontsize=6.5)
axB.set_xlim(0, HA_LEN); axB.set_xlabel("HA consensus position")
axB.axvspan(18, 343, color=GRAY, alpha=0.04); axB.axvspan(344, HA_LEN, color=STALK_C, alpha=0.06)
axB.text(180, -0.8, "HA1 (head)", ha="center", fontsize=6, color=GRAY)
axB.text(440, -0.8, "HA2 (stalk)", ha="center", fontsize=6, color=STALK_C)
panel(axB, "B", "Neutralizing antibody footprints on HA consensus")

# C 抗体残基数
axC = fig.add_subplot(gs[1, 2])
all_ab = sorted(fps, key=lambda f: -f["n"])
names = [f['antibody'] for f in all_ab]
nn = [f['n'] for f in all_ab]
cc = [region_color.get(f['region'], GRAY) for f in all_ab]
axC.barh(range(len(all_ab))[::-1], nn, color=cc, height=0.6)
axC.set_yticks(range(len(all_ab))[::-1]); axC.set_yticklabels(names, fontsize=6.5)
axC.set_xlabel("Contact residues")
for i, n in enumerate(nn):
    axC.text(n + 0.3, len(nn) - 1 - i, str(n), va="center", fontsize=6)
panel(axC, "C", "Contact residues per antibody")

# D dN/dS 轨迹
axD = fig.add_subplot(gs[1, 0])
axD.scatter(codon, np.clip(dnds_val, 0, 5), s=4, color=GRAY, alpha=0.25, lw=0)
axD.plot(codon, np.clip(smooth, 0, 5), color=BLUE, lw=1.3)
axD.axhline(1, color=INK, ls="--", lw=0.7)
axD.set_xlabel("HA codon position"); axD.set_ylabel("dN/dS")
axD.set_ylim(0, 4)
panel(axD, "D", "Per-codon dN/dS (raw + sliding median, H1N1 HA)")

# E 叠加
axE = fig.add_subplot(gs[1, 1])
# dN/dS 轨迹（底部）
axE.plot(codon, np.clip(smooth, 0, 3) / 3, color=BLUE, lw=1.0, alpha=0.6, label="dN/dS (scaled)")
# 足迹（HA 抗体，H1N1）
for f in ha_ab:
    if f["subtype"] == "H1N1":
        axE.scatter(f["pos"], [0.75 + 0.02 * ha_ab.index(f)] * len(f["pos"]), s=8,
                    color=region_color.get(f["region"], GRAY), alpha=0.6)
# 候选区域
axE.axvspan(59, 77, color=RED, alpha=0.15, label="candidate CKLRG… (59–77)")
axE.set_xlabel("HA consensus position"); axE.set_ylim(-0.1, 1.1); axE.set_yticks([])
axE.legend(frameon=False, fontsize=6, loc="upper left")
panel(axE, "E", "Candidate, footprints, and dN/dS co-localization")

fig.savefig(OUT, dpi=300, bbox_inches="tight", facecolor="white", pad_inches=0.03)
plt.close(fig)
print("saved", OUT)
print(f"RSA vs contact-density: Spearman ρ={r_sp:+.3f}, Pearson r={r_pe:+.3f}")
```
