## Supplementary material for "Conserved influenza A epitope candidate regions and a benchmark of ESM-2 sequence features": Li_et_al_Supplementary_Data: make_fig5_v2.py

```
#!/usr/bin/env python3
"""Figure 5 — Structural, antibody and evolutionary context（5 panel，figure1_redesign 风格）。"""
import csv
import numpy as np
import pandas as pd
import matplotlib
matplotlib.use("Agg")
import matplotlib.pyplot as plt
from scipy import stats
import fig_style as FS

ROOT = "/media/qxl/My_Passport/流行病学数据/广谱流感疫苗候选课题"
RES = f"{ROOT}/08_ESM2_AI分析与论文/results"
MS = f"{ROOT}/07_MANUSCRIPT_STRENGTHENING"
FIG = f"{MS}/figures/main"

FS.setup_style()

rsa = np.genfromtxt(f"{RES}/rsa_validation.csv", delimiter=",", skip_header=1)
rsa_val, cd_val = rsa[:, 0], rsa[:, 1]
r_sp, _ = stats.spearmanr(rsa_val, cd_val); r_pe, _ = stats.pearsonr(rsa_val, cd_val)

fps = list(csv.DictReader(open(
    f"{MS}/08_BCELL_EXTERNAL_BENCHMARK/structures/09_NEUTRALIZING_FOOTPRINTS_EXTENDED.tsv"), delimiter="\t"))
ha_ab = [f for f in fps if f["protein"] == "HA"]
for f in ha_ab:
    f["pos"] = [int(x) for x in f["consensus_positions"].split(";")]
    f["n"] = len(f["pos"])
    f["region_short"] = "stalk" if f["region"] == "stalk" else "head"
ha_ab.sort(key=lambda f: (f["region_short"] == "stalk", f["antibody"]))
region_color = {"head": FS.ORANGE, "stalk": FS.BLUE}

dnds = list(csv.DictReader(open(f"{MS}/dnds_per_codon.tsv"), delimiter="\t"))
codon = np.array([int(r["codon_index"]) for r in dnds])
dnds_val = np.array([np.nan if r["dN_dS"] in ("", "NA") else float(r["dN_dS"]) for r in dnds])
smooth = pd.Series(dnds_val).rolling(window=15, center=True, min_periods=1).median().to_numpy()

CAND = "HA-34"; CAND_START, CAND_END = 59, 77

fig = plt.figure(figsize=(12, 9.5))
gs = fig.add_gridspec(2, 3, height_ratios=[1, 1.2], hspace=0.55, wspace=0.4)

# A RSA hexbin
ax = fig.add_subplot(gs[0, 0])
hb = ax.hexbin(rsa_val, cd_val, gridsize=30, cmap="Blues", mincnt=1, bins="log")
ax.text(0.03, 0.97, f"Spearman ρ = {r_sp:+.2f}\nPearson r = {r_pe:+.2f}",
        transform=ax.transAxes, fontsize=7.5, va="top", color=FS.TEXT)
ax.set_xlabel("Surface accessibility (RSA)"); ax.set_ylabel("Contact density")
cb = fig.colorbar(hb, ax=ax, fraction=0.046, pad=0.03); cb.set_label("count", fontsize=7); cb.ax.tick_params(labelsize=6)
FS.panel(ax, "A", "Surface accessibility and contact density")

# B 足迹 landscape
ax = fig.add_subplot(gs[0, 1:])
HA_LEN = 567
for i, f in enumerate(ha_ab):
    y = len(ha_ab) - 1 - i
    c = region_color[f["region_short"]]
    ax.hlines(y, 0, HA_LEN, color=FS.GRAY_LIGHT, lw=0.5)
    ax.vlines(f["pos"], y - 0.28, y + 0.28, color=c, lw=1.4)
ax.axvspan(18, 343, color=FS.GRAY, alpha=0.04)
ax.axvspan(344, HA_LEN, color=FS.BLUE, alpha=0.05)
ax.text(180, -0.9, "HA1 head", ha="center", fontsize=6.5, color=FS.GRAY)
ax.text(440, -0.9, "HA2 stalk", ha="center", fontsize=6.5, color=FS.BLUE)
ax.set_yticks(range(len(ha_ab)))
ax.set_yticklabels([f"{f['antibody']} ({f['subtype']})" for f in ha_ab], fontsize=6.5)
ax.set_xlim(0, HA_LEN); ax.set_xlabel("HA consensus position")
FS.panel(ax, "B", "Neutralizing-antibody footprints along HA")

# C contact burden
ax = fig.add_subplot(gs[1, 0])
all_ab = sorted(ha_ab, key=lambda f: -f["n"])
names = [f['antibody'] for f in all_ab]; nn = [f['n'] for f in all_ab]
cc = [region_color[f['region_short']] for f in all_ab]
ax.barh(range(len(all_ab))[::-1], nn, color=cc, height=0.6)
ax.set_yticks(range(len(all_ab))[::-1]); ax.set_yticklabels(names, fontsize=6.5)
ax.set_xlabel("Contact residues")
for i, n in enumerate(nn):
    ax.text(n + 0.3, len(nn) - 1 - i, str(n), va="center", fontsize=6)
FS.panel(ax, "C", "Antibody contact burden")

# D dN/dS
ax = fig.add_subplot(gs[1, 1])
ax.scatter(codon, np.clip(dnds_val, 0, 5), s=3, color=FS.GRAY_LIGHT, alpha=0.5, lw=0)
ax.plot(codon, np.clip(smooth, 0, 5), color=FS.BLUE_DARK, lw=1.3)
ax.axhline(1, color=FS.GRAY, lw=0.7, ls="--")
ax.set_xlabel("HA codon position"); ax.set_ylabel("dN/dS"); ax.set_ylim(0, 4)
FS.panel(ax, "D", "Evolutionary constraint along HA")

# E 集成位置证据
ax = fig.add_subplot(gs[1, 2])
ax.set_xlim(0, HA_LEN)
ax.axvspan(CAND_START, CAND_END, color=FS.ORANGE_LIGHT, alpha=0.5, zorder=1)
ax.text(CAND_START + 0.5, 2.4, CAND, fontsize=7, fontweight="bold", color=FS.TEXT, va="center", ha="left")
for i, f in enumerate(ha_ab):
    c = region_color[f["region_short"]]
    ax.scatter(f["pos"], [2.0 - i * 0.08] * len(f["pos"]), s=6, color=c, alpha=0.5, zorder=2)
ax.plot(codon, np.clip(smooth, 0, 3) / 3, color=FS.BLUE, lw=1.0, zorder=3)
ax.set_ylim(-0.1, 2.6); ax.set_yticks([])
ax.set_xlabel("HA consensus position")
ax.text(0.01, 0.88, "candidate", transform=ax.transAxes, fontsize=6, color=FS.ORANGE)
ax.text(0.01, 0.42, "contacts", transform=ax.transAxes, fontsize=6, color=FS.BLUE)
ax.text(0.01, 0.05, "dN/dS", transform=ax.transAxes, fontsize=6, color=FS.BLUE)
FS.panel(ax, "E", "Integrated positional evidence")

FS.save(fig, f"{FIG}/Figure4")
print(f"RSA: Spearman={r_sp:+.3f}, Pearson={r_pe:+.3f}")
print(f"足迹抗体: {[f['antibody'] for f in ha_ab]}, n={[f['n'] for f in ha_ab]}")
```
