## Supplementary material for "Conserved influenza A epitope candidate regions and a benchmark of ESM-2 sequence features": Li_et_al_Supplementary_Data: make_figures.py

```
#!/usr/bin/env python3
"""Generate the 8 figures for manuscript_v2 (consistent style)."""
import os, csv
from collections import Counter
import numpy as np
import matplotlib
matplotlib.use("Agg")
import matplotlib.pyplot as plt
from matplotlib.patches import FancyBboxPatch
from Bio import SeqIO
from scipy import stats

BASE = "/media/qxl/My_Passport/流行病学数据/广谱流感疫苗候选课题"
RES = f"{BASE}/08_ESM2_AI分析与论文/results"
ESM = f"{RES}/esm2"
CONS = f"{BASE}/08_ESM2_AI分析与论文/data/consensus"
RAW = f"{BASE}/01_原始序列与分层数据/序列与地区分层/全球/序列/综合"
FIG = f"{BASE}/07_MANUSCRIPT_STRENGTHENING/figures"
os.makedirs(FIG, exist_ok=True)

BLUE="#3A6EA5"; ORANGE="#C77C3F"; GREEN="#4E8D5C"; RED="#B5453B"; GRAY="#8A8F98"; INK="#2B2F36"
COL={"H1N1":BLUE,"H3N2":ORANGE,"H5N1":GREEN}
plt.rcParams.update({"font.family":"DejaVu Sans","font.size":9,"axes.spines.top":False,
    "axes.spines.right":False,"axes.edgecolor":INK,"axes.labelcolor":INK,"text.color":INK,
    "xtick.color":INK,"ytick.color":INK,"axes.linewidth":0.8})

def z(a): return (a-np.nanmean(a))/np.nanstd(a)

# ---- Fig 1: cross-subtype conservation heatmap ----
def fig1():
    rows=list(csv.DictReader(open(f"{RES}/ai_ranking.csv")))
    for r in rows: r["final_score"]=float(r["final_score"])
    rows.sort(key=lambda r:-r["final_score"]); top=rows[:16]
    cons={}
    for sub in ["H1N1","H3N2","H5N1"]:
        for prot in ["HA","NA","M1","NP","NEP","PA","PB1","PB2"]:
            p=f"{CONS}/{sub}_{prot}.fasta"
            if os.path.exists(p): cons[(sub,prot)]=str(list(SeqIO.parse(p,"fasta"))[0].seq)
    mat=[]; labels=[]
    for r in top:
        epi=r["epitope"]; prot=r["protein"]
        if prot=="M": prot="M1"
        row=[]
        for sub in ["H1N1","H3N2","H5N1"]:
            s=cons.get((sub,prot),"")
            best=0.0
            if s:
                for i in range(len(s)-len(epi)+1):
                    best=max(best,sum(a==b for a,b in zip(epi,s[i:i+len(epi)]))/len(epi))
            row.append(best)
        mat.append(row); labels.append(f'{prot}:{epi[:13]}')
    mat=np.array(mat)
    fig,ax=plt.subplots(figsize=(6,6))
    im=ax.imshow(mat,cmap="viridis",vmin=0.5,vmax=1.0,aspect="auto")
    ax.set_xticks(range(3)); ax.set_xticklabels(["H1N1","H3N2","H5N1"])
    ax.set_yticks(range(len(labels))); ax.set_yticklabels(labels,fontsize=7)
    for i in range(mat.shape[0]):
        for j in range(3):
            ax.text(j,i,f"{mat[i,j]:.2f}",ha="center",va="center",fontsize=6,color="white" if mat[i,j]<0.8 else INK)
    ax.set_title("Cross-subtype identity of top candidate regions",fontweight="bold")
    fig.colorbar(im,ax=ax,fraction=0.046,pad=0.03,label="identity")
    fig.tight_layout(); fig.savefig(f"{FIG}/Fig1_crosssubtype.png",dpi=200,bbox_inches="tight",facecolor="white"); plt.close(fig)
    print("Fig1 ok")

# ---- Fig 2: temporal stability ----
def fig2():
    import re
    from collections import defaultdict
    def year_seqs(sub,prot):
        p=f"{RAW}/分类序列_{sub}/{prot}_sequences.fasta"; ys=defaultdict(list)
        for r in SeqIO.parse(p,"fasta"):
            m=re.search(r"/20\d\d",r.description)
            if m and 2006<=int(m.group(0)[1:])<=2017:
                ys[int(m.group(0)[1:])].append(str(r.seq[:(len(r.seq)//3)*3].translate()))
        return ys
    def frac(seqs,epi):
        b=[]
        for s in seqs:
            x=0.0
            if len(s)>=len(epi):
                for i in range(len(s)-len(epi)+1): x=max(x,sum(a==c for a,c in zip(epi,s[i:i+len(epi)]))/len(epi))
            b.append(x)
        return (np.array(b)>=0.9).mean()
    cons_ha=str(list(SeqIO.parse(f"{CONS}/H1N1_HA.fasta","fasta"))[0].seq)
    epi="CKLRGVAPLHLGKCNIAGW"; i=cons_ha.find(epi)
    ys=year_seqs("H1N1","HA"); years=sorted(ys)
    fracs=[frac(ys[y],epi) for y in years]
    fig,ax=plt.subplots(figsize=(5.5,3.4))
    ax.plot(years,fracs,'o-',color=RED,lw=2,ms=5)
    slope,inter,_,p,_=stats.linregress(years,fracs)
    ax.plot(years,[slope*y+inter for y in years],'--',color=GRAY,lw=1.3)
    ax.set_xlabel("Year"); ax.set_ylabel("Fraction ≥90% identity")
    ax.set_title(f"Temporal stability of HA RBD candidate\nslope {slope:+.3f}/yr, p={p:.2f}",fontweight="bold")
    ax.set_ylim(0.4,1.05)
    fig.tight_layout(); fig.savefig(f"{FIG}/Fig2_temporal.png",dpi=200,bbox_inches="tight",facecolor="white"); plt.close(fig)
    print("Fig2 ok")

# ---- Fig 3: ranking + weight sensitivity ----
def fig3():
    rows=list(csv.DictReader(open(f"{RES}/ai_ranking.csv")))
    for r in rows:
        r["final_score"]=float(r["final_score"]); r["ai_conservation"]=float(r["ai_conservation"])
    rows=[r for r in rows if not np.isnan(r["ai_conservation"])]
    rows.sort(key=lambda r:-r["final_score"]); top=rows[:12]
    names=[f'{r["protein"]}:{r["epitope"][:12]}' for r in top]
    fig,ax=plt.subplots(figsize=(6,4.2))
    y=np.arange(len(top))[::-1]
    ax.barh(y,[r["final_score"] for r in top],color=[RED if r["protein"] in ("HA","NA") else BLUE for r in top],height=0.6)
    ax.set_yticks(y); ax.set_yticklabels(names,fontsize=7)
    ax.set_xlabel("Combined score")
    ax.set_title("Top-ranked candidate regions\n(surface glycoproteins in red)",fontweight="bold")
    fig.tight_layout(); fig.savefig(f"{FIG}/Fig3_ranking.png",dpi=200,bbox_inches="tight",facecolor="white"); plt.close(fig)
    print("Fig3 ok")

# ---- Fig 4: illustrative assemblies ----
def fig4():
    fig,ax=plt.subplots(figsize=(8,3.2)); ax.set_xlim(0,10); ax.set_ylim(0,3.2); ax.axis("off")
    rows=[("HA assembly",["RBD","antigenic 2"],"≤51 aa",RED),
          ("NA assembly",["A","B","C","D (active site)"],"≤119 aa",GREEN),
          ("NP / M1 / NEP",["NP core","M1","NEP"],"51-54 aa",BLUE)]
    y=2.6
    for name,segs,tot,c in rows:
        ax.text(0.1,y,name,fontsize=9,fontweight="bold",va="center",color=INK)
        x=1.8
        for s in segs:
            w=1.7; ax.add_patch(FancyBboxPatch((x,y-0.42),w,0.84,boxstyle="round,pad=0.05,rounding_size=0.14",fc=c,ec="none",alpha=0.85))
            ax.text(x+w/2,y,s,ha="center",va="center",fontsize=6.5,color="white"); x+=w+0.12
        ax.text(9.8,y,tot,fontsize=7,color=GRAY,ha="right",va="center"); y-=0.85
    ax.text(5,0.15,"Illustrative multi-epitope assemblies (not validated immunogens)",ha="center",fontsize=8,style="italic",color=GRAY)
    fig.savefig(f"{FIG}/Fig4_assemblies.png",dpi=200,bbox_inches="tight",facecolor="white"); plt.close(fig)
    print("Fig4 ok")

# ---- Fig 5: PLL vs conservation ----
def fig5():
    def mode_seqs(sub,prot):
        p=f"{RAW}/分类序列_{sub}/{prot}_sequences.fasta"; recs=list(SeqIO.parse(p,"fasta"))
        aa=[str(r.seq[:(len(r.seq)//3)*3].translate()) for r in recs]
        ln=Counter(len(s) for s in aa).most_common(1)[0][0]
        return [s for s in aa if len(s)==ln and s.count("X")<=0.05*len(s)]
    def cons_ident(sub,prot):
        seqs=mode_seqs(sub,prot); L=len(seqs[0])
        c=[Counter(s[i] for s in seqs).most_common(1)[0][0] for i in range(L)]
        return np.array([sum(1 for s in seqs if s[i]==c[i])/len(seqs) for i in range(L)])
    fig,ax=plt.subplots(figsize=(5,4))
    xs=[];ys=[]
    for sub in ["H1N1","H3N2","H5N1"]:
        pll=np.load(f"{ESM}/{sub}_HA_pll.npy"); seq=open(f"{ESM}/{sub}_HA_seq.txt").read().strip()
        ci=cons_ident(sub,"HA"); n=min(len(seq),len(ci)); zz=z(pll[1:1+n])
        ax.scatter(ci[:n],zz,s=5,color=COL[sub],alpha=0.3,lw=0)
        xs+=list(ci[:n]); ys+=list(zz)
    rho,p=stats.spearmanr(xs,ys)
    ax.text(0.03,0.95,f"Spearman ρ = {rho:.2f}",transform=ax.transAxes,fontsize=11,fontweight="bold",va="top")
    ax.set_xlabel("MSA positional identity"); ax.set_ylabel("ESM-2 z-PLL")
    ax.set_title("HA: PLL vs conservation (3 subtypes)",fontweight="bold")
    fig.tight_layout(); fig.savefig(f"{FIG}/Fig5_pll_conservation.png",dpi=200,bbox_inches="tight",facecolor="white"); plt.close(fig)
    print("Fig5 ok")

# ---- Fig 6: T-cell benchmark AUROC ----
def fig6():
    lines=open(f"{BASE}/07_MANUSCRIPT_STRENGTHENING/01_ESM_INCREMENTAL_BENCHMARK/benchmark_results.tsv").read().splitlines()
    rows={}
    for l in lines[1:]:
        p=l.split("\t")
        if p[0].startswith("incremental"): continue
        rows[p[0]]=dict(auc=float(p[1]),lo=float(p[2]),hi=float(p[3]))
    labels=['A: conservation','C: +PLL','D: +PLL+contact']
    keys=['A_conservation','C_conservation+PLL','D_conservation+PLL+contact']
    fig,ax=plt.subplots(figsize=(5,3.6))
    aucs=[rows[k]['auc'] for k in keys]; los=[rows[k]['lo'] for k in keys]; his=[rows[k]['hi'] for k in keys]
    x=np.arange(3)
    ax.bar(x,aucs,yerr=[[a-l for a,l in zip(aucs,los)],[h-a for a,h in zip(aucs,his)]],color=[BLUE,ORANGE,GREEN],capsize=4,width=0.5)
    ax.axhline(0.5,color=GRAY,ls='--',lw=0.8)
    ax.set_xticks(x); ax.set_xticklabels(labels,fontsize=8); ax.set_ylabel("AUROC"); ax.set_ylim(0.35,0.7)
    for i,a in enumerate(aucs): ax.text(i,a+0.012,f"{a:.3f}",ha="center",fontsize=8)
    ax.set_title("T-cell benchmark: ESM-2 no incremental value",fontweight="bold")
    fig.tight_layout(); fig.savefig(f"{FIG}/Fig6_benchmark_auroc.png",dpi=200,bbox_inches="tight",facecolor="white"); plt.close(fig)
    print("Fig6 ok")

# ---- Fig 7: contact-density vs RSA ----
def fig7():
    fig,ax=plt.subplots(figsize=(5,4))
    # HA
    ha=np.genfromtxt(f"{RES}/rsa_validation.csv",delimiter=',',skip_header=1)
    rsa_ha,cd_ha=ha[:,0],ha[:,1]
    r_ha,p_ha=stats.spearmanr(rsa_ha,cd_ha)
    ax.scatter(rsa_ha,cd_ha,s=5,color=RED,alpha=0.4,lw=0,label=f"HA (ρ={r_ha:+.2f})")
    # NA (recompute)
    import freesasa
    from Bio.PDB import PDBParser
    T3={'ALA':'A','ARG':'R','ASN':'N','ASP':'D','CYS':'C','GLN':'Q','GLU':'E','GLY':'G','HIS':'H','ILE':'I','LEU':'L','LYS':'K','MET':'M','PHE':'F','PRO':'P','SER':'S','THR':'T','TRP':'W','TYR':'Y','VAL':'V'}
    REF={'A':129.0,'R':274.0,'N':195.0,'D':193.0,'C':167.0,'Q':225.0,'E':223.0,'G':104.0,'H':224.0,'I':197.0,'L':201.0,'K':236.0,'M':224.0,'F':240.0,'P':159.0,'S':155.0,'T':172.0,'W':285.0,'Y':263.0,'V':174.0}
    res=freesasa.calc(freesasa.Structure(f"/tmp/2HTY.pdb"))
    ca=res.residueAreas().get("A",{})
    ch=PDBParser(QUIET=True).get_structure("2HTY",f"/tmp/2HTY.pdb")[0]["A"]
    na_seq=""; na_rsa=[]
    for r_ in ch:
        if r_.id[0]!=" ": continue
        aa=T3.get(r_.resname.strip())
        if aa is None: continue
        ra=ca.get(str(r_.id[1]))
        na_seq+=aa; na_rsa.append(ra.total/REF.get(aa,200.0) if ra else np.nan)
    from Bio.Align import PairwiseAligner
    cons_na=str(list(SeqIO.parse(f"{CONS}/H1N1_NA.fasta","fasta"))[0].seq)
    al=PairwiseAligner(); al.mode='local'; al.match_score=2; al.mismatch_score=-1; al.open_gap_score=-2; al.extend_gap_score=-1
    a=al.align(cons_na,na_seq)[0]; tb,qb=a.aligned; q2t={}
    for (ts,te),(qs,qe) in zip(tb,qb):
        for off in range(te-ts): q2t[qs+off]=ts+off
    acc_na=np.load(f"{ESM}/H1N1_NA_acc.npy")
    x=[];y=[]
    for qi,ci in q2t.items():
        if qi<len(na_rsa) and not np.isnan(na_rsa[qi]): x.append(na_rsa[qi]); y.append(acc_na[ci])
    r_na,p_na=stats.spearmanr(x,y)
    ax.scatter(x,y,s=5,color=BLUE,alpha=0.4,lw=0,label=f"NA (ρ={r_na:+.2f})")
    ax.set_xlabel("Relative solvent-accessible surface area (FreeSASA)")
    ax.set_ylabel("ESM-2 contact-density")
    ax.set_title("Contact-density is not an accessibility proxy",fontweight="bold")
    ax.legend(frameon=False)
    fig.tight_layout(); fig.savefig(f"{FIG}/Fig7_rsa.png",dpi=200,bbox_inches="tight",facecolor="white"); plt.close(fig)
    print("Fig7 ok")

# ---- Fig 8: neutralizing antibody footprints ----
def fig8():
    rows=list(csv.DictReader(open(f"{BASE}/07_MANUSCRIPT_STRENGTHENING/08_BCELL_EXTERNAL_BENCHMARK/structures/09_NEUTRALIZING_FOOTPRINTS.tsv"),delimiter="\t"))
    cons_ha=str(list(SeqIO.parse(f"{CONS}/H1N1_HA.fasta","fasta"))[0].seq)
    fig,ax=plt.subplots(figsize=(9,3.2))
    L=len(cons_ha)
    ax.plot(range(1,L+1),np.zeros(L),color=INK,lw=2)  # HA backbone
    ax.axvspan(344,L,color=GREEN,alpha=0.15)  # HA2
    ax.axvspan(18,343,color=GRAY,alpha=0.08)  # HA1
    ax.text(180,0.5,"HA1",ha="center",fontsize=8,color=GRAY); ax.text(430,0.5,"HA2 (stalk)",ha="center",fontsize=8,color=GREEN)
    colors={"CR6261":RED,"CR8020":ORANGE,"CR9114":BLUE}
    for k,r in enumerate(rows):
        pos=[int(x) for x in r["consensus_positions"].split(";")]
        ax.scatter(pos,[0.5+k*0.6]*len(pos),s=25,color=colors[r["antibody"]],label=f'{r["antibody"]} ({r["subtype"]})')
    ax.set_xlabel("HA consensus residue position"); ax.set_ylim(-0.3,2.6); ax.set_yticks([])
    ax.set_title("Broadly neutralizing stalk-antibody contact footprints",fontweight="bold")
    ax.legend(frameon=False,fontsize=8,loc="upper left")
    fig.tight_layout(); fig.savefig(f"{FIG}/Fig8_footprints.png",dpi=200,bbox_inches="tight",facecolor="white"); plt.close(fig)
    print("Fig8 ok")

if __name__=="__main__":
    fig1(); fig2(); fig3(); fig4(); fig5(); fig6(); fig7(); fig8()
    print("ALL FIGURES DONE")
```
