## Supplementary material for "Conserved influenza A epitope candidate regions and a benchmark of ESM-2 sequence features": Li_et_al_Supplementary_Data: make_figures_creative.py

```
#!/usr/bin/env python3
"""Creative figures:
  FigC1: 3D HA structure, C-alpha atoms colored by ESM-2 PLL (localization to head)
  FigC2: circular proteome map of conserved candidate regions (9 proteins)
  FigC3: ridge plot of PLL distributions (neutralizing vs binding vs background)
"""
import os, csv
import numpy as np
import matplotlib
matplotlib.use("Agg")
import matplotlib.pyplot as plt
from mpl_toolkits.mplot3d import Axes3D
from Bio import SeqIO
from Bio.PDB import PDBParser
from Bio.Align import PairwiseAligner
from scipy import stats

BASE = "/media/qxl/My_Passport/流行病学数据/广谱流感疫苗候选课题"
RES = f"{BASE}/08_ESM2_AI分析与论文/results"
ESM = f"{RES}/esm2"
CONS = f"{BASE}/08_ESM2_AI分析与论文/data/consensus"
FIG = f"{BASE}/07_MANUSCRIPT_STRENGTHENING/figures"

BLUE="#3A6EA5"; ORANGE="#C77C3F"; GREEN="#4E8D5C"; RED="#B5453B"; GRAY="#8A8F98"; INK="#2B2F36"
plt.rcParams.update({"font.family":"DejaVu Sans","font.size":9,"axes.edgecolor":INK,
    "axes.labelcolor":INK,"text.color":INK,"xtick.color":INK,"ytick.color":INK})

def z(a): return (a-np.nanmean(a))/np.nanstd(a)

T3={'ALA':'A','ARG':'R','ASN':'N','ASP':'D','CYS':'C','GLN':'Q','GLU':'E','GLY':'G','HIS':'H','ILE':'I','LEU':'L','LYS':'K','MET':'M','PHE':'F','PRO':'P','SER':'S','THR':'T','TRP':'W','TYR':'Y','VAL':'V'}

# ---------- FigC1: 3D HA colored by PLL ----------
def figC1():
    cons_ha = str(list(SeqIO.parse(f"{CONS}/H1N1_HA.fasta","fasta"))[0].seq)
    pll = np.load(f"{ESM}/H1N1_HA_pll.npy")
    zpll = z(pll)  # length = len(cons)+2
    aligner = PairwiseAligner(); aligner.mode='local'; aligner.match_score=2
    aligner.mismatch_score=-1; aligner.open_gap_score=-2; aligner.extend_gap_score=-1

    p = PDBParser(QUIET=True)
    s = p.get_structure("1RU7","/tmp/1RU7.pdb")[0]
    # collect CA coords + map to consensus for HA1 (A) + HA2 (B)
    pts=[]; cols=[]
    for chain_id in ["A","B"]:
        ch = s[chain_id]
        seq=""; resnums=[]
        for r in ch:
            if r.id[0]!=" ": continue
            aa=T3.get(r.resname.strip())
            if aa is None or "CA" not in r: continue
            seq+=aa; resnums.append(r)
        a = aligner.align(cons_ha, seq)[0]
        tb,qb = a.aligned; q2t={}
        for (ts,te),(qs,qe) in zip(tb,qb):
            for off in range(te-ts): q2t[qs+off]=ts+off
        for qi, r in enumerate(resnums):
            if qi in q2t:
                ci = q2t[qi]
                ca = r["CA"].coord
                pts.append(ca); cols.append(zpll[ci+1])
    pts=np.array(pts); cols=np.array(cols)
    fig=plt.figure(figsize=(7,6))
    ax=fig.add_subplot(111,projection="3d")
    sc=ax.scatter(pts[:,0],pts[:,1],pts[:,2],c=cols,cmap="viridis",s=22,vmin=-2,vmax=2,alpha=0.95)
    ax.set_axis_off()
    ax.view_init(elev=20,azim=-60)
    fig.colorbar(sc,ax=ax,fraction=0.02,pad=0.02,label="ESM-2 z-PLL")
    ax.set_title("HA structure colored by ESM-2 PLL (H1N1, PDB 1RU7)",fontweight="bold",pad=8)
    fig.tight_layout(); fig.savefig(f"{FIG}/FigC1_structure_pll.png",dpi=220,bbox_inches="tight",facecolor="white"); plt.close(fig)
    print("FigC1 (3D structure) ok")

# ---------- FigC2: circular proteome map ----------
def figC2():
    rows=list(csv.DictReader(open(f"{RES}/ai_ranking.csv")))
    prots=["HA","NA","M1","NEP","NP","NS1","PA","PB1","PB2"]
    fig,ax=plt.subplots(figsize=(8,8),subplot_kw={"projection":"polar"})
    ax.set_theta_zero_location("N"); ax.set_theta_direction(-1)
    n=len(prots)
    for i,p in enumerate(prots):
        theta=(2*np.pi*i/n)
        # protein arc
        ax.bar(theta, 1.0, width=2*np.pi/n*0.85, bottom=0.5, color=GRAY, alpha=0.15)
        ax.text(theta, 1.7, p, ha="center", va="center", fontsize=11, fontweight="bold", color=INK)
        # candidate regions as colored ticks
        cand=[r for r in rows if r["protein"]==p]
        for j,r in enumerate(cand[:8]):
            t2=theta + (j/(max(len(cand[:8]),1)-1)-0.5)*2*np.pi/n*0.6 if len(cand[:8])>1 else theta
            c=RED if p in ("HA","NA") else BLUE
            ax.bar(t2, 0.5, width=0.04, bottom=0.7, color=c, alpha=0.8)
    ax.set_ylim(0,1.9); ax.set_yticks([])
    ax.set_title("Conserved candidate regions across the influenza A proteome\n(red = surface glycoproteins, blue = internal)",fontweight="bold",pad=20)
    fig.tight_layout(); fig.savefig(f"{FIG}/FigC2_circular_map.png",dpi=200,bbox_inches="tight",facecolor="white"); plt.close(fig)
    print("FigC2 (circular map) ok")

# ---------- FigC3: ridge plot of PLL ----------
def figC3():
    # categories: neutralizing clusters, binding clusters, matched background
    c06=list(csv.DictReader(open(f"{BASE}/07_MANUSCRIPT_STRENGTHENING/08_BCELL_EXTERNAL_BENCHMARK/06_BENCHMARK_READY_DATASET.tsv"),delimiter="\t"))
    bg=list(csv.DictReader(open(f"{BASE}/07_MANUSCRIPT_STRENGTHENING/08_BCELL_EXTERNAL_BENCHMARK/07_MATCHED_BACKGROUND.tsv"),delimiter="\t"))
    cons={}
    for sub in ["H1N1","H3N2","H5N1"]:
        for prot in ["HA","NA"]:
            cons[(sub,prot)]=str(list(SeqIO.parse(f"{CONS}/{sub}_{prot}.fasta","fasta"))[0].seq)
    def region_pll(sub,prot,seq):
        s=cons[(sub,prot)]; i=s.find(seq)
        if i<0:
            for j in range(len(s)-len(seq)+1):
                if sum(a!=b for a,b in zip(seq,s[j:j+len(seq)]))<=1: i=j; break
        if i<0: return np.nan
        pll=np.load(f"{ESM}/{sub}_{prot}_pll.npy"); zz=z(pll)
        return np.mean(zz[i+1:i+1+len(seq)])
    neut=[]; bind=[]
    for c in c06:
        v=region_pll(c["subtype"],c["protein"],c["representative_seq"])
        if np.isnan(v): continue
        if c["label_level"]=="L3_neutralization": neut.append(v)
        else: bind.append(v)
    bgr=[]
    for r in bg:
        v=region_pll(r["subtype"],r["protein"],r["bg_sequence"])
        if not np.isnan(v): bgr.append(v)
    groups=[("Neutralization\n(n=%d)"%len(neut),neut,RED),
            ("Antibody-binding\n(n=%d)"%len(bind),bind,ORANGE),
            ("Matched background\n(n=%d)"%len(bgr),bgr,GRAY)]
    fig,axes=plt.subplots(len(groups),1,figsize=(6,5),sharex=True)
    for i,(name,vals,c) in enumerate(groups):
        ax=axes[i]
        vals=np.array(vals)
        ax.hist(vals,bins=25,color=c,alpha=0.8,density=True)
        ax.axvline(np.mean(vals),color=INK,lw=1.5)
        ax.text(0.98,0.7,f"mean {np.mean(vals):+.2f}",transform=ax.transAxes,ha="right",fontsize=8)
        ax.set_ylabel(name,fontsize=8); ax.set_yticks([])
        ax.spines["left"].set_visible(False)
    axes[-1].set_xlabel("ESM-2 mean z-PLL")
    fig.suptitle("PLL distributions: neutralizing vs binding vs background",fontweight="bold")
    fig.tight_layout(); fig.savefig(f"{FIG}/FigC3_ridge.png",dpi=200,bbox_inches="tight",facecolor="white"); plt.close(fig)
    print("FigC3 (ridge) ok")

if __name__=="__main__":
    figC1(); figC2(); figC3()
    print("CREATIVE FIGURES DONE")
```
