## Supplementary material for "Conserved influenza A epitope candidate regions and a benchmark of ESM-2 sequence features": Li_et_al_Supplementary_Data: make_figures_extra.py

```
#!/usr/bin/env python3
"""Additional figures (Fig 9-14) + fixed Fig 8 (no overlap)."""
import os, csv, re
from collections import Counter, defaultdict
import numpy as np
import matplotlib
matplotlib.use("Agg")
import matplotlib.pyplot as plt
from Bio import SeqIO
from scipy import stats

BASE = "/media/qxl/My_Passport/流行病学数据/广谱流感疫苗候选课题"
RES = f"{BASE}/08_ESM2_AI分析与论文/results"
ESM = f"{RES}/esm2"
CONS = f"{BASE}/08_ESM2_AI分析与论文/data/consensus"
RAW = f"{BASE}/01_原始序列与分层数据/序列与地区分层/全球/序列/综合"
FIG = f"{BASE}/07_MANUSCRIPT_STRENGTHENING/figures"

BLUE="#3A6EA5"; ORANGE="#C77C3F"; GREEN="#4E8D5C"; RED="#B5453B"; GRAY="#8A8F98"; INK="#2B2F36"
COL={"H1N1":BLUE,"H3N2":ORANGE,"H5N1":GREEN}
plt.rcParams.update({"font.family":"DejaVu Sans","font.size":9,"axes.spines.top":False,
    "axes.spines.right":False,"axes.edgecolor":INK,"axes.labelcolor":INK,"text.color":INK,
    "xtick.color":INK,"ytick.color":INK,"axes.linewidth":0.8})

def z(a): return (a-np.nanmean(a))/np.nanstd(a)
def smooth(a,k=15): return np.convolve(a,np.ones(k)/k,mode="same")

# ---- Fig 8 (fixed): footprints, better spacing ----
def fig8():
    rows=list(csv.DictReader(open(f"{BASE}/07_MANUSCRIPT_STRENGTHENING/08_BCELL_EXTERNAL_BENCHMARK/structures/09_NEUTRALIZING_FOOTPRINTS.tsv"),delimiter="\t"))
    cons_ha=str(list(SeqIO.parse(f"{CONS}/H1N1_HA.fasta","fasta"))[0].seq)
    L=len(cons_ha)
    fig,ax=plt.subplots(figsize=(9,3.8))
    # HA backbone at bottom
    ax.plot(range(1,L+1),np.zeros(L),color=INK,lw=2.5,alpha=0.7,zorder=1)
    ax.axvspan(344,L,color=GREEN,alpha=0.12)
    ax.axvspan(18,343,color=GRAY,alpha=0.06)
    # domain labels at bottom (below backbone)
    ax.text(180,-0.55,"HA1 (globular head)",ha="center",fontsize=8,color=GRAY)
    ax.text(440,-0.55,"HA2 (stalk / fusion)",ha="center",fontsize=8,color=GREEN)
    colors={"CR6261":RED,"CR8020":ORANGE,"CR9114":BLUE}
    for k,r in enumerate(rows):
        pos=[int(x) for x in r["consensus_positions"].split(";")]
        yy=1.0+k*1.1   # clear separation
        ax.scatter(pos,[yy]*len(pos),s=35,color=colors[r["antibody"]],zorder=2,
                   label=f'{r["antibody"]} ({r["subtype"]} HA stalk)')
    ax.set_xlabel("HA consensus residue position")
    ax.set_ylim(-1.0, 3.6); ax.set_yticks([])
    ax.set_title("Broadly neutralizing stalk-antibody contact footprints mapped to consensus",fontweight="bold")
    ax.legend(frameon=False,fontsize=8,loc="upper right")
    fig.tight_layout(); fig.savefig(f"{FIG}/Fig8_footprints.png",dpi=200,bbox_inches="tight",facecolor="white"); plt.close(fig)
    print("Fig8 (fixed) ok")

# ---- Fig 9: candidate landscape ----
def fig9():
    rows=list(csv.DictReader(open(f"{RES}/ai_ranking.csv")))
    cnt=Counter(r["protein"] for r in rows)
    prots=["HA","NA","M1","NEP","NP","NS1","PA","PB1","PB2"]
    vals=[cnt.get(p,0) for p in prots]
    fig,ax=plt.subplots(figsize=(6,3.4))
    ax.bar(prots,vals,color=[RED if p in ("HA","NA") else BLUE for p in prots],alpha=0.85)
    ax.set_xlabel("Viral protein"); ax.set_ylabel("Candidate regions")
    ax.set_title("Distribution of 98 candidate regions by protein",fontweight="bold")
    for i,v in enumerate(vals): ax.text(i,v+0.5,str(v),ha="center",fontsize=8)
    fig.tight_layout(); fig.savefig(f"{FIG}/Fig9_candidate_landscape.png",dpi=200,bbox_inches="tight",facecolor="white"); plt.close(fig)
    print("Fig9 ok")

# ---- Fig 10: B-cell benchmark data flow ----
def fig10():
    fig,ax=plt.subplots(figsize=(8,3.6)); ax.set_xlim(0,10); ax.set_ylim(0,4); ax.axis("off")
    steps=[("IEDB B-cell\n(122 epitopes)",0.3,"#5B6672"),
           ("HA/NA surface\n(70)",2.6,"#5B6672"),
           ("Mapped to consensus\n(33 PASS/AMBIG)",4.9,"#7A8794"),
           ("Non-redundant\nclusters (22)",7.2,RED)]
    for i,(txt,x,c) in enumerate(steps):
        ax.add_patch(plt.Rectangle((x,1.4),1.9,1.1,fc=c,ec="none",alpha=0.9))
        ax.text(x+0.95,1.95,txt,ha="center",va="center",fontsize=8,color="white",fontweight="bold")
        if i<3:
            ax.annotate("",xy=(x+1.95,1.95),xytext=(x+2.55,1.95),
                        arrowprops=dict(arrowstyle="-|>",color=GRAY,lw=1.4))
    # exclusion notes
    ax.text(1.25,0.7,"exclude 52 non-HA/NA",ha="center",fontsize=6.5,color=GRAY)
    ax.text(3.75,0.7,"exclude 37 (12 conformational,\n15 no-consensus, 10 drift)",ha="center",fontsize=6.5,color=GRAY)
    ax.text(6.05,0.7,"merge 11 overlapping",ha="center",fontsize=6.5,color=GRAY)
    ax.text(5,3.5,"B-cell benchmark data flow",ha="center",fontsize=10,fontweight="bold",color=INK)
    fig.tight_layout(); fig.savefig(f"{FIG}/Fig10_dataflow.png",dpi=200,bbox_inches="tight",facecolor="white"); plt.close(fig)
    print("Fig10 ok")

# ---- Fig 11: raw sequence heterogeneity ----
def fig11():
    fig,axes=plt.subplots(1,3,figsize=(9,3.2))
    for ax,sub in zip(axes,["H1N1","H3N2","H5N1"]):
        p=f"{RAW}/分类序列_{sub}/HA_sequences.fasta"
        lens=[len(r.seq) for r in SeqIO.parse(p,"fasta")]
        ax.hist(lens,bins=40,color=COL[sub],alpha=0.8)
        ax.axvline(1701,color=RED,ls='--',lw=1.2,label="modal 1701 nt")
        ax.set_xlabel("HA nucleotide length"); ax.set_ylabel("Count")
        ax.set_title(f"{sub} HA",fontweight="bold")
        if sub=="H1N1": ax.legend(frameon=False,fontsize=6)
    fig.suptitle("Raw HA sequence length heterogeneity",fontweight="bold",y=1.03)
    fig.tight_layout(); fig.savefig(f"{FIG}/Fig11_heterogeneity.png",dpi=200,bbox_inches="tight",facecolor="white"); plt.close(fig)
    print("Fig11 ok")

# ---- Fig 12: HA PLL profile ----
def fig12():
    fig,ax=plt.subplots(figsize=(8,3.2))
    for sub in ["H1N1","H3N2","H5N1"]:
        pll=np.load(f"{ESM}/{sub}_HA_pll.npy"); seq=open(f"{ESM}/{sub}_HA_seq.txt").read().strip()
        zz=smooth(z(pll[1:1+len(seq)]))
        ax.plot(np.arange(len(seq)),zz,lw=1.5,label=sub,color=COL[sub])
    ax.axvspan(18,343,color=GRAY,alpha=0.1); ax.axvspan(344,566,color=GREEN,alpha=0.08)
    ax.text(180,ax.get_ylim()[1]*0.85,"HA1 head",ha="center",fontsize=7,color=GRAY)
    ax.text(440,ax.get_ylim()[1]*0.85,"HA2",ha="center",fontsize=7,color=GREEN)
    ax.set_xlabel("HA residue position"); ax.set_ylabel("ESM-2 z-PLL")
    ax.set_title("PLL localizes to the HA1 globular head (functional region)",fontweight="bold")
    ax.legend(frameon=False,fontsize=8)
    fig.tight_layout(); fig.savefig(f"{FIG}/Fig12_ha_pll.png",dpi=200,bbox_inches="tight",facecolor="white"); plt.close(fig)
    print("Fig12 ok")

# ---- Fig 13: NA PLL profile ----
def fig13():
    fig,ax=plt.subplots(figsize=(8,3.2))
    for sub in ["H1N1","H3N2","H5N1"]:
        pll=np.load(f"{ESM}/{sub}_NA_pll.npy"); seq=open(f"{ESM}/{sub}_NA_seq.txt").read().strip()
        zz=smooth(z(pll[1:1+len(seq)]))
        ax.plot(np.arange(len(seq)),zz,lw=1.5,label=sub,color=COL[sub])
    n=len(seq)
    ax.axvspan(n//3,2*n//3,color=GREEN,alpha=0.1)
    ax.text(n//2,ax.get_ylim()[1]*0.85,"enzyme-active site",ha="center",fontsize=7,color=GREEN)
    ax.set_xlabel("NA residue position"); ax.set_ylabel("ESM-2 z-PLL")
    ax.set_title("PLL localizes to the NA enzyme-active site",fontweight="bold")
    ax.legend(frameon=False,fontsize=8)
    fig.tight_layout(); fig.savefig(f"{FIG}/Fig13_na_pll.png",dpi=200,bbox_inches="tight",facecolor="white"); plt.close(fig)
    print("Fig13 ok")

# ---- Fig 14: HLA population coverage ----
def fig14():
    fig,(ax1,ax2)=plt.subplots(1,2,figsize=(7,3.4))
    ax1.pie([94.4,5.6],colors=[GREEN,"#E0E4E8"],startangle=90,counterclock=False,
            wedgeprops=dict(width=0.35,edgecolor="white"))
    ax1.text(0,0.03,"94.4%",ha="center",va="center",fontsize=16,fontweight="bold")
    ax1.text(0,-0.32,"14 HLA-I alleles",ha="center",fontsize=7,color=INK)
    ax1.set_title("HLA-I coverage",fontweight="bold",fontsize=9)
    ax2.pie([73.0,27.0],colors=[BLUE,"#E0E4E8"],startangle=90,counterclock=False,
            wedgeprops=dict(width=0.35,edgecolor="white"))
    ax2.text(0,0.03,"73.0%",ha="center",va="center",fontsize=16,fontweight="bold")
    ax2.text(0,-0.32,"6 HLA-DRB1 alleles",ha="center",fontsize=7,color=INK)
    ax2.set_title("HLA-II coverage",fontweight="bold",fontsize=9)
    fig.suptitle("Population coverage of screening alleles",fontweight="bold",y=1.02)
    fig.tight_layout(); fig.savefig(f"{FIG}/Fig14_hlacoverage.png",dpi=200,bbox_inches="tight",facecolor="white"); plt.close(fig)
    print("Fig14 ok")

if __name__=="__main__":
    fig8(); fig9(); fig10(); fig11(); fig12(); fig13(); fig14()
    print("EXTRA FIGURES DONE")
```
