## Supplementary material for "Conserved influenza A epitope candidate regions and a benchmark of ESM-2 sequence features": Li_et_al_Supplementary_Data: make_figures_final.py

```
#!/usr/bin/env python3
"""
生成 Fig 17（H3N2 扩展：表面候选亚型特异）与 Fig 18（扩展中和抗体足迹）。
风格沿用 make_figures_extra.py 的配色与 rcParams。
"""
import csv
import matplotlib
matplotlib.use("Agg")
import matplotlib.pyplot as plt
import numpy as np

ROOT = "/media/qxl/My_Passport/流行病学数据/广谱流感疫苗候选课题"
BASE = f"{ROOT}/07_MANUSCRIPT_STRENGTHENING"
FIG = f"{BASE}/figures"

BLUE="#3A6EA5"; ORANGE="#C77C3F"; GREEN="#4E8D5C"; RED="#B5453B"; GRAY="#8A8F98"; INK="#2B2F36"
COL={"H1N1":BLUE,"H3N2":ORANGE,"H5N1":GREEN}
plt.rcParams.update({"font.family":"DejaVu Sans","font.size":9,"axes.spines.top":False,
    "axes.spines.right":False,"axes.edgecolor":INK,"axes.labelcolor":INK,"text.color":INK,
    "xtick.color":INK,"ytick.color":INK,"axes.linewidth":0.8})

# ------------------------------------------------------------- 数据载入
def load_surface():
    """表面候选（HA/NA），含真实跨亚型与 H3N2 保守率。"""
    cst = {}
    for r in csv.DictReader(open(f"{BASE}/05_MANUSCRIPT_CONSISTENCY/cross_subtype_true.csv")):
        cst[(r["protein"], r["epitope"])] = r
    h3 = {}
    for r in csv.DictReader(open(f"{BASE}/09_H3N2_CROSS_LINEAGE/h3n2_cross_lineage.tsv"), delimiter="\t"):
        h3[(r["protein"], r["epitope"])] = r
    out = []
    for r in csv.DictReader(open(f"{ROOT}/08_ESM2_AI分析与论文/results/ai_ranking.csv")):
        if r["protein"] not in ("HA", "NA"):
            continue
        cs = cst.get((r["protein"], r["epitope"]), {})
        hr = h3.get((r["protein"], r["epitope"]), {})
        out.append(dict(
            protein=r["protein"], epitope=r["epitope"],
            H1N1=cs.get("H1N1", "-"), H3N2=cs.get("H3N2", "-"), H5N1=cs.get("H5N1", "-"),
            frac_all=float(hr["frac_all_H3N2"]) if "frac_all_H3N2" in hr else np.nan))
    # 按蛋白、长度排序
    out.sort(key=lambda x: (x["protein"] != "HA", x["epitope"]))
    return out

def load_footprints():
    rows = list(csv.DictReader(open(
        f"{BASE}/08_BCELL_EXTERNAL_BENCHMARK/structures/09_NEUTRALIZING_FOOTPRINTS_EXTENDED.tsv"), delimiter="\t"))
    for r in rows:
        r["pos"] = [int(x) for x in r["consensus_positions"].split(";") if x]
    return rows

# ------------------------------------------------------------- Fig 17
def fig17():
    surf = load_surface()
    labs = [f"{r['protein']} {r['epitope'][:22]}" for r in surf]
    # Panel A: consensus presence (exact=1, fuzzy=0.5, absent=0)
    val = {"exact": 1.0, "fuzzy": 0.5, "-": 0.0}
    A = np.array([[val[r[s]] for s in ("H1N1", "H3N2", "H5N1")] for r in surf])
    # Panel B: H3N2 clade conservation (candidate × 6 clades)
    h3 = {}
    for r in csv.DictReader(open(f"{BASE}/09_H3N2_CROSS_LINEAGE/h3n2_cross_lineage.tsv"), delimiter="\t"):
        h3[(r["protein"], r["epitope"])] = r
    clade_keys = [k for k in h3[list(h3.keys())[0]].keys() if k.startswith("clade_")]
    clade_names = [k.replace("clade_", "") for k in clade_keys]
    B = np.zeros((len(surf), len(clade_keys)))
    for i, r in enumerate(surf):
        hr = h3.get((r["protein"], r["epitope"]), {})
        for j, k in enumerate(clade_keys):
            v = hr.get(k, "")
            B[i, j] = float(v) if v not in ("", None) else 0.0

    fig, axes = plt.subplots(1, 2, figsize=(9, 3.6), gridspec_kw={"width_ratios": [1, 2.2]})

    # Panel A
    ax = axes[0]
    im = ax.imshow(A, cmap="Blues", vmin=0, vmax=1, aspect="auto")
    ax.set_xticks(range(3)); ax.set_xticklabels(["H1N1", "H3N2", "H5N1"])
    ax.set_yticks(range(len(labs))); ax.set_yticklabels(labs, fontsize=7.5)
    ax.set_title("A. Presence in subtype consensus", fontsize=9, loc="left", pad=8)
    for i in range(A.shape[0]):
        for j in range(A.shape[1]):
            t = "exact" if A[i, j] == 1.0 else ("fuzzy" if A[i, j] == 0.5 else "—")
            ax.text(j, i, t, ha="center", va="center", fontsize=6,
                    color="white" if A[i, j] > 0.4 else INK)
    ax.tick_params(length=0)

    # Panel B
    ax = axes[1]
    im2 = ax.imshow(B, cmap="Blues", vmin=0, vmax=1, aspect="auto")
    ax.set_xticks(range(len(clade_names))); ax.set_xticklabels(clade_names, rotation=45, ha="right", fontsize=7)
    ax.set_yticks(range(len(labs))); ax.set_yticklabels([])
    ax.set_title("B. Conservation in H3N2 clades (2010–2025, ≥90% identity)", fontsize=9, loc="left", pad=8)
    for i in range(B.shape[0]):
        for j in range(B.shape[1]):
            if B[i, j] > 0.05:
                ax.text(j, i, f"{B[i, j]:.2f}", ha="center", va="center", fontsize=6, color="white")
    ax.tick_params(length=0)
    fig.colorbar(im2, ax=axes[1], fraction=0.03, pad=0.02, label="conserved fraction")

    fig.tight_layout()
    fig.savefig(f"{FIG}/Fig17_h3n2_subtype_specificity.png", dpi=200, bbox_inches="tight", facecolor="white")
    plt.close(fig)
    print("saved Fig17_h3n2_subtype_specificity.png")

# ------------------------------------------------------------- Fig 18
def fig18():
    fps = load_footprints()
    ha = [f for f in fps if f["protein"] == "HA"]
    na = [f for f in fps if f["protein"] == "NA"]
    # 排序：stalk 在前，RBD 在后（按 region）
    ha.sort(key=lambda f: (f["region"] != "stalk", f["antibody"]))

    fig, axes = plt.subplots(2, 1, figsize=(9, 4.2), gridspec_kw={"height_ratios": [len(ha), len(na)]})

    # HA 面板
    ax = axes[0]
    HA_LEN = 567
    for i, f in enumerate(ha):
        c = COL[f["subtype"]]
        y = len(ha) - 1 - i
        ax.hlines(y, 0, HA_LEN, color=GRAY, lw=0.4, alpha=0.5)
        ax.vlines(f["pos"], y - 0.25, y + 0.25, color=c, lw=1.2)
    ax.set_yticks(range(len(ha)))
    ax.set_yticklabels([f"{f['antibody']} ({f['subtype']}, {f['region'].split(' ')[0]})" for f in ha], fontsize=7.5)
    ax.set_xlim(0, HA_LEN); ax.set_xlabel("HA consensus position")
    ax.set_title("Neutralizing-antibody contact footprints on HA (H3 numbering ≈ position)", fontsize=9, loc="left")

    # NA 面板
    ax = axes[1]
    NA_LEN = 470
    for i, f in enumerate(na):
        c = COL[f["subtype"]]
        y = len(na) - 1 - i
        ax.hlines(y, 0, NA_LEN, color=GRAY, lw=0.4, alpha=0.5)
        ax.vlines(f["pos"], y - 0.3, y + 0.3, color=c, lw=1.4)
    ax.set_yticks(range(len(na)))
    ax.set_yticklabels([f"{f['antibody']} ({f['subtype']}, {f['region']})" for f in na], fontsize=7.5)
    ax.set_xlim(0, NA_LEN); ax.set_xlabel("NA consensus position")
    ax.set_title("Footprints on NA", fontsize=9, loc="left")

    fig.tight_layout()
    fig.savefig(f"{FIG}/Fig18_extended_footprints.png", dpi=200, bbox_inches="tight", facecolor="white")
    plt.close(fig)
    print("saved Fig18_extended_footprints.png")

if __name__ == "__main__":
    fig17()
    fig18()
```
