## Supplementary material for "Conserved influenza A epitope candidate regions and a benchmark of ESM-2 sequence features": Li_et_al_Supplementary_Data: make_figures_nc.py

```
#!/usr/bin/env python3
"""
NC (Nature Communications) 风格图重做：
- Arial 字体，无内嵌标题（图注放正文）
- 面板角标 a/b/c/d：小写加粗，左上角，无背景框，位置统一对称
- 克制配色：分类色 BLUE/ORANGE/GREEN/RED，热图用单 hue 'Blues'，3D 用 'RdBu_r'
- 细线、刻度朝外、充足留白

直接绘制 7 张主文多面板图 + 5 张附录图（不拼接 PNG）。
"""
import os, csv, re
from collections import Counter, defaultdict
import numpy as np
import matplotlib
matplotlib.use("Agg")
import matplotlib.pyplot as plt
import matplotlib.image as mpimg
from matplotlib.patches import FancyBboxPatch
from Bio import SeqIO
from scipy import stats

ROOT = "/media/qxl/My_Passport/流行病学数据/广谱流感疫苗候选课题"
RES = f"{ROOT}/08_ESM2_AI分析与论文/results"
ESM = f"{RES}/esm2"
CONS = f"{ROOT}/08_ESM2_AI分析与论文/data/consensus"
RAW = f"{ROOT}/01_原始序列与分层数据/序列与地区分层/全球/序列/综合"
MS = f"{ROOT}/07_MANUSCRIPT_STRENGTHENING"
FIG = f"{MS}/figures"
MAIN = f"{FIG}/main"
SUPP = f"{FIG}/supplementary"
os.makedirs(MAIN, exist_ok=True); os.makedirs(SUPP, exist_ok=True)

BLUE="#3A6EA5"; ORANGE="#C77C3F"; GREEN="#4E8D5C"; RED="#B5453B"; GRAY="#8A8F98"; INK="#22262B"
COL={"H1N1":BLUE,"H3N2":ORANGE,"H5N1":GREEN}

# ---- NC 风格 rcParams ----
plt.rcParams.update({
    "font.family": "Arial",
    "font.size": 8,
    "axes.spines.top": False,
    "axes.spines.right": False,
    "axes.linewidth": 0.6,
    "axes.edgecolor": INK, "axes.labelcolor": INK, "text.color": INK,
    "xtick.color": INK, "ytick.color": INK,
    "xtick.direction": "out", "ytick.direction": "out",
    "xtick.major.width": 0.6, "ytick.major.width": 0.6,
    "xtick.major.size": 3, "ytick.major.size": 3,
    "axes.axisbelow": True,
})

def z(a): return (a - np.nanmean(a)) / np.nanstd(a)

def panel(ax, label, title=None):
    """角标 a/b/c/d（小写加粗，左上角，无背景）；可选面板标题 title。"""
    kw = dict(fontsize=12, fontweight="bold", va="bottom", ha="left")
    if hasattr(ax, "get_zlim"):          # 3D 轴用 text2D
        ax.text2D(-0.06, 1.08, label, transform=ax.transAxes, **kw)
    else:
        ax.text(-0.06, 1.08, label, transform=ax.transAxes, **kw)
    if title is not None:
        ax.set_title(title, loc="left", fontsize=8.5, pad=10)

def save(fig, name, dpi=300):
    fig.savefig(name, dpi=dpi, bbox_inches="tight", facecolor="white", pad_inches=0.02)
    plt.close(fig)
    print("saved", os.path.basename(name))

# ================================================================ 数据助手
def load_cons():
    cons = {}
    for sub in ["H1N1","H3N2","H5N1"]:
        for prot in ["HA","NA","M1","NP","NEP","NS1","PA","PB1","PB2"]:
            p = f"{CONS}/{sub}_{prot}.fasta"
            if os.path.exists(p):
                cons[(sub, prot)] = str(list(SeqIO.parse(p, "fasta"))[0].seq)
    return cons

def best_identity(seq, epi):
    if len(seq) < len(epi): return 0.0
    b = 0.0
    for i in range(len(seq) - len(epi) + 1):
        b = max(b, sum(a == c for a, c in zip(epi, seq[i:i+len(epi)])) / len(epi))
    return b

def frac(seqs, epi, thresh=0.9):
    if not seqs: return np.nan
    return np.mean([best_identity(s, epi) >= thresh for s in seqs])

# ================================================================ Fig 1 (2×2)
def fig1():
    cons = load_cons()
    rows = list(csv.DictReader(open(f"{RES}/ai_ranking.csv")))
    for r in rows: r["final_score"] = float(r["final_score"])
    rows.sort(key=lambda r: -r["final_score"]); top = rows[:16]

    fig, axes = plt.subplots(2, 2, figsize=(7.2, 6.6))

    # --- a: cross-subtype identity heatmap ---
    ax = axes[0, 0]
    mat, labels = [], []
    for r in top:
        epi, prot = r["epitope"], ("M1" if r["protein"] == "M" else r["protein"])
        row = [best_identity(cons.get((s, prot), ""), epi) for s in ("H1N1", "H3N2", "H5N1")]
        mat.append(row); labels.append(f"{prot} {epi[:12]}")
    mat = np.array(mat)
    im = ax.imshow(mat, cmap="Blues", vmin=0.5, vmax=1.0, aspect="auto")
    ax.set_xticks(range(3)); ax.set_xticklabels(["H1N1","H3N2","H5N1"])
    ax.set_yticks(range(len(labels))); ax.set_yticklabels(labels, fontsize=6.5)
    for i in range(mat.shape[0]):
        for j in range(3):
            ax.text(j, i, f"{mat[i,j]:.2f}", ha="center", va="center", fontsize=5.5,
                    color="white" if mat[i, j] < 0.8 else INK)
    ax.tick_params(length=0)
    cb = fig.colorbar(im, ax=ax, fraction=0.046, pad=0.03); cb.set_label("identity", fontsize=7); cb.ax.tick_params(labelsize=6)
    panel(ax, "a", "Cross-subtype conservation")

    # --- b: temporal stability ---
    ax = axes[0, 1]
    def year_seqs(sub, prot):
        ys = defaultdict(list)
        for r in SeqIO.parse(f"{RAW}/分类序列_{sub}/{prot}_sequences.fasta", "fasta"):
            m = re.search(r"/20\d\d", r.description)
            if m and 2006 <= int(m.group(0)[1:]) <= 2017:
                ys[int(m.group(0)[1:])].append(str(r.seq[:(len(r.seq)//3)*3].translate()))
        return ys
    epi = "CKLRGVAPLHLGKCNIAGW"
    ys = year_seqs("H1N1", "HA"); years = sorted(ys)
    fracs = [frac(ys[y], epi) for y in years]
    ax.plot(years, fracs, "o-", color=RED, lw=1.4, ms=4)
    slope, inter, *_ = stats.linregress(years, fracs)
    ax.plot(years, [slope*y+inter for y in years], "--", color=GRAY, lw=1)
    ax.set_xlabel("Year"); ax.set_ylabel("Fraction ≥ 90% identity")
    ax.set_ylim(0.4, 1.05); ax.set_yticks([0.5, 0.7, 0.9])
    panel(ax, "b", "Temporal stability (H1N1 HA)")

    # --- c: H1N1 cross-lineage (2015-2025, 6 clades) ---
    ax = axes[1, 0]
    meta = {}
    for r in csv.DictReader(open(f"{ROOT}/99_历史项目与参考/早期H1N1系统发育项目/清洗数据/cleaned_metadata.csv")):
        meta[r["Isolate_Id"]] = r
    def trans_atg(seq):
        from Bio.Seq import Seq
        s = str(seq).upper().replace("-", "")
        for i in range(len(s)-2):
            if s[i:i+3] == "ATG":
                n = (len(s)-i)//3; aa = []
                for c in range(n):
                    cod = s[i+3*c:i+3*c+3]
                    aa.append("X" if (len(cod) < 3 or "N" in cod) else str(Seq(cod).translate()))
                return "".join(aa)
        return ""
    aln = f"{ROOT}/99_历史项目与参考/早期H1N1系统发育项目/清洗数据/clustalo-H1N1-HA.fa"
    by_clade = defaultdict(list)
    for r in SeqIO.parse(aln, "fasta"):
        cl = meta.get(r.id, {}).get("Clade", "unassigned")
        aa = trans_atg(r.seq)
        if aa and aa.count("X") <= 0.05*len(aa):
            by_clade[cl].append(aa)
    top_clades = [c for c, _ in Counter({k: len(v) for k, v in by_clade.items()}).most_common(6)]
    f_clades = [frac(by_clade[c], epi) for c in top_clades]
    xs = np.arange(6)
    ax.bar(xs, f_clades, color=BLUE, width=0.55)
    ax.set_xticks(xs); ax.set_xticklabels([c.replace(".", ".\n", 1) for c in top_clades], fontsize=5.5)
    ax.set_ylabel("Conserved fraction"); ax.set_ylim(0, 1.1)
    ax.axhline(0.9, color=GRAY, ls="--", lw=0.8)
    panel(ax, "c", "Cross-lineage conservation (H1N1)")

    # --- d: H3N2 subtype-specificity (surface candidates × clades) ---
    ax = axes[1, 1]
    h3 = {}
    for r in csv.DictReader(open(f"{MS}/09_H3N2_CROSS_LINEAGE/h3n2_cross_lineage.tsv"), delimiter="\t"):
        h3[(r["protein"], r["epitope"])] = r
    surf = [r for r in rows if r["protein"] in ("HA", "NA")]
    ckeys = [k for k in h3[list(h3.keys())[0]].keys() if k.startswith("clade_")]
    B = np.zeros((len(surf), len(ckeys)))
    slab = []
    for i, r in enumerate(surf):
        hr = h3.get((r["protein"], r["epitope"]), {})
        for j, k in enumerate(ckeys):
            v = hr.get(k, ""); B[i, j] = float(v) if v not in ("", None) else 0.0
        slab.append(f"{r['protein']} {r['epitope'][:10]}")
    im = ax.imshow(B, cmap="Blues", vmin=0, vmax=1, aspect="auto")
    ax.set_xticks(range(len(ckeys)))
    ax.set_xticklabels([c.replace("clade_", "").replace(".", ".\n", 1) for c in ckeys], fontsize=5.5)
    ax.set_yticks(range(len(slab))); ax.set_yticklabels(slab, fontsize=6)
    for i in range(B.shape[0]):
        for j in range(B.shape[1]):
            if B[i, j] > 0.05:
                ax.text(j, i, f"{B[i,j]:.2f}", ha="center", va="center", fontsize=5.5, color="white")
    ax.tick_params(length=0)
    cb = fig.colorbar(im, ax=ax, fraction=0.046, pad=0.03); cb.set_label("conserved fraction", fontsize=7); cb.ax.tick_params(labelsize=6)
    panel(ax, "d", "H3N2 clade conservation")

    save(fig, f"{MAIN}/Fig1_candidate_conservation.png")

# ================================================================ Fig 2 (1×2)
def fig2():
    fig, axes = plt.subplots(1, 2, figsize=(7.2, 3.4))
    def mode_seqs(sub, prot):
        recs = list(SeqIO.parse(f"{RAW}/分类序列_{sub}/{prot}_sequences.fasta", "fasta"))
        aa = [str(r.seq[:(len(r.seq)//3)*3].translate()) for r in recs]
        ln = Counter(len(s) for s in aa).most_common(1)[0][0]
        return [s for s in aa if len(s) == ln and s.count("X") <= 0.05*len(s)]
    def cons_ident(sub, prot):
        seqs = mode_seqs(sub, prot); L = len(seqs[0])
        c = [Counter(s[i] for s in seqs).most_common(1)[0][0] for i in range(L)]
        return np.array([sum(1 for s in seqs if s[i] == c[i]) / len(seqs) for i in range(L)])

    ax = axes[0]
    xs, ys = [], []
    for sub in ["H1N1", "H3N2", "H5N1"]:
        pll = np.load(f"{ESM}/{sub}_HA_pll.npy"); seq = open(f"{ESM}/{sub}_HA_seq.txt").read().strip()
        ci = cons_ident(sub, "HA"); n = min(len(seq), len(ci)); zz = z(pll[1:1+n])
        ax.scatter(ci[:n], zz, s=4, color=COL[sub], alpha=0.35, lw=0)
        xs += list(ci[:n]); ys += list(zz)
    rho, _ = stats.spearmanr(xs, ys)
    ax.text(0.03, 0.95, f"Spearman ρ = {rho:.2f}", transform=ax.transAxes, fontsize=10, fontweight="bold", va="top")
    ax.set_xlabel("MSA positional identity"); ax.set_ylabel("ESM-2 z-PLL")
    panel(ax, "a", "PLL vs. MSA conservation")

    ax = axes[1]
    from Bio.PDB import PDBParser
    from Bio.Align import PairwiseAligner
    T3 = {'ALA':'A','ARG':'R','ASN':'N','ASP':'D','CYS':'C','GLN':'Q','GLU':'E','GLY':'G','HIS':'H','ILE':'I','LEU':'L','LYS':'K','MET':'M','PHE':'F','PRO':'P','SER':'S','THR':'T','TRP':'W','TYR':'Y','VAL':'V'}
    cons_ha = str(list(SeqIO.parse(f"{CONS}/H1N1_HA.fasta", "fasta"))[0].seq)
    pll = np.load(f"{ESM}/H1N1_HA_pll.npy"); zpll = z(pll)
    al = PairwiseAligner(); al.mode = "local"; al.match_score = 2; al.mismatch_score = -1; al.open_gap_score = -2; al.extend_gap_score = -1
    s = PDBParser(QUIET=True).get_structure("1RU7", "/tmp/1RU7.pdb")[0]
    pts, cols = [], []
    for cid in ["A", "B"]:
        ch = s[cid]; seq = ""; resnums = []
        for r in ch:
            if r.id[0] != " ": continue
            aa = T3.get(r.resname.strip())
            if aa is None or "CA" not in r: continue
            seq += aa; resnums.append(r)
        a = al.align(cons_ha, seq)[0]; tb, qb = a.aligned; q2t = {}
        for (ts, te), (qs, qe) in zip(tb, qb):
            for off in range(te-ts): q2t[qs+off] = ts+off
        for qi, r in enumerate(resnums):
            if qi in q2t:
                ca = r["CA"].coord; pts.append(ca); cols.append(zpll[q2t[qi]+1])
    pts = np.array(pts); cols = np.array(cols)
    from mpl_toolkits.mplot3d import Axes3D
    ax3 = fig.add_subplot(122, projection="3d")
    sc = ax3.scatter(pts[:,0], pts[:,1], pts[:,2], c=cols, cmap="RdBu_r", s=14, vmin=-2, vmax=2)
    ax3.set_axis_off(); ax3.view_init(elev=20, azim=-60)
    fig.colorbar(sc, ax=ax3, fraction=0.02, pad=0.02, label="ESM-2 z-PLL")
    panel(ax3, "b", "HA structure colored by PLL")
    axes[1].set_axis_off()

    save(fig, f"{MAIN}/Fig2_esm_pll.png")

# ================================================================ Fig 3 (单面板)
def fig3():
    lines = open(f"{MS}/01_ESM_INCREMENTAL_BENCHMARK/benchmark_results.tsv").read().splitlines()
    rows = {}
    for l in lines[1:]:
        p = l.split("\t")
        if p[0].startswith("incremental"): continue
        rows[p[0]] = dict(auc=float(p[1]), lo=float(p[2]), hi=float(p[3]))
    keys = ['A_conservation', 'C_conservation+PLL', 'D_conservation+PLL+contact']
    labels = ['conservation', '+ PLL', '+ PLL\n+ contact']
    aucs = [rows[k]['auc'] for k in keys]; los = [rows[k]['lo'] for k in keys]; his = [rows[k]['hi'] for k in keys]
    fig, ax = plt.subplots(figsize=(3.6, 3.2))
    x = np.arange(3)
    ax.bar(x, aucs, yerr=[[a-l for a, l in zip(aucs, los)], [h-a for a, h in zip(aucs, his)]],
           color=[GRAY, BLUE, ORANGE], capsize=3, width=0.5, error_kw=dict(lw=0.8, ecolor=INK))
    ax.set_title("T-cell benchmark: ESM-2 adds no incremental value", loc="left", fontsize=9, pad=8)
    ax.axhline(0.5, color=INK, ls="--", lw=0.7)
    ax.set_xticks(x); ax.set_xticklabels(labels, fontsize=7); ax.set_ylabel("AUROC"); ax.set_ylim(0.35, 0.7)
    for i, a in enumerate(aucs): ax.text(i, a+0.012, f"{a:.3f}", ha="center", fontsize=7)
    save(fig, f"{MAIN}/Fig3_tcell_benchmark.png")

# ================================================================ Fig 4 (单面板)
def fig4():
    fig, ax = plt.subplots(figsize=(3.8, 3.4))
    ha = np.genfromtxt(f"{RES}/rsa_validation.csv", delimiter=",", skip_header=1)
    rsa_ha, cd_ha = ha[:,0], ha[:,1]
    r_ha, _ = stats.spearmanr(rsa_ha, cd_ha)
    ax.scatter(rsa_ha, cd_ha, s=4, color=RED, alpha=0.4, lw=0, label=f"HA (ρ = {r_ha:+.2f})")
    # NA
    import freesasa
    from Bio.PDB import PDBParser
    T3 = {'ALA':'A','ARG':'R','ASN':'N','ASP':'D','CYS':'C','GLN':'Q','GLU':'E','GLY':'G','HIS':'H','ILE':'I','LEU':'L','LYS':'K','MET':'M','PHE':'F','PRO':'P','SER':'S','THR':'T','TRP':'W','TYR':'Y','VAL':'V'}
    REF = {'A':129.0,'R':274.0,'N':195.0,'D':193.0,'C':167.0,'Q':225.0,'E':223.0,'G':104.0,'H':224.0,'I':197.0,'L':201.0,'K':236.0,'M':224.0,'F':240.0,'P':159.0,'S':155.0,'T':172.0,'W':285.0,'Y':263.0,'V':174.0}
    res = freesasa.calc(freesasa.Structure("/tmp/2HTY.pdb"))
    ca = res.residueAreas().get("A", {})
    ch = PDBParser(QUIET=True).get_structure("2HTY", "/tmp/2HTY.pdb")[0]["A"]
    na_seq, na_rsa = "", []
    for r_ in ch:
        if r_.id[0] != " ": continue
        aa = T3.get(r_.resname.strip())
        if aa is None: continue
        ra = ca.get(str(r_.id[1]))
        na_seq += aa; na_rsa.append(ra.total/REF.get(aa, 200.0) if ra else np.nan)
    from Bio.Align import PairwiseAligner
    cons_na = str(list(SeqIO.parse(f"{CONS}/H1N1_NA.fasta", "fasta"))[0].seq)
    al = PairwiseAligner(); al.mode = "local"; al.match_score = 2; al.mismatch_score = -1; al.open_gap_score = -2; al.extend_gap_score = -1
    a = al.align(cons_na, na_seq)[0]; tb, qb = a.aligned; q2t = {}
    for (ts, te), (qs, qe) in zip(tb, qb):
        for off in range(te-ts): q2t[qs+off] = ts+off
    acc_na = np.load(f"{ESM}/H1N1_NA_acc.npy")
    x, y = [], []
    for qi, ci in q2t.items():
        if qi < len(na_rsa) and not np.isnan(na_rsa[qi]): x.append(na_rsa[qi]); y.append(acc_na[ci])
    r_na, _ = stats.spearmanr(x, y)
    ax.scatter(x, y, s=4, color=BLUE, alpha=0.4, lw=0, label=f"NA (ρ = {r_na:+.2f})")
    ax.set_title("Contact-density vs. solvent accessibility", loc="left", fontsize=9, pad=8)
    ax.set_xlabel("Relative solvent accessibility (FreeSASA)"); ax.set_ylabel("ESM-2 contact-density")
    ax.legend(frameon=False, fontsize=7)
    save(fig, f"{MAIN}/Fig4_contact_rsa.png")

# ================================================================ Fig 5 (1×2)
def fig5():
    fig, axes = plt.subplots(1, 2, figsize=(7.2, 3.4))
    # a: ridge
    c06 = list(csv.DictReader(open(f"{MS}/08_BCELL_EXTERNAL_BENCHMARK/06_BENCHMARK_READY_DATASET.tsv"), delimiter="\t"))
    bg = list(csv.DictReader(open(f"{MS}/08_BCELL_EXTERNAL_BENCHMARK/07_MATCHED_BACKGROUND.tsv"), delimiter="\t"))
    cons = {}
    for sub in ["H1N1","H3N2","H5N1"]:
        for prot in ["HA","NA"]:
            cons[(sub, prot)] = str(list(SeqIO.parse(f"{CONS}/{sub}_{prot}.fasta", "fasta"))[0].seq)
    def region_pll(sub, prot, seq):
        s = cons[(sub, prot)]; i = s.find(seq)
        if i < 0:
            for j in range(len(s)-len(seq)+1):
                if sum(a != b for a, b in zip(seq, s[j:j+len(seq)])) <= 1: i = j; break
        if i < 0: return np.nan
        pll = np.load(f"{ESM}/{sub}_{prot}_pll.npy"); zz = z(pll)
        return np.mean(zz[i+1:i+1+len(seq)])
    neut, bind = [], []
    for c in c06:
        v = region_pll(c["subtype"], c["protein"], c["representative_seq"])
        if np.isnan(v): continue
        (neut if c["label_level"] == "L3_neutralization" else bind).append(v)
    bgr = [region_pll(r["subtype"], r["protein"], r["bg_sequence"]) for r in bg]
    bgr = [v for v in bgr if not np.isnan(v)]
    groups = [("Neutralizing", neut, RED), ("Binding", bind, ORANGE), ("Background", bgr, GRAY)]
    ax = axes[0]
    for i, (name, vals, c) in enumerate(groups):
        vals = np.array(vals)
        ax.hist(vals, bins=22, color=c, alpha=0.75, density=True, histtype="stepfilled")
        ax.axvline(np.mean(vals), color=INK, lw=1)
        ax.text(0.98, 0.72 - i*0.0, f"{name} (n={len(vals)})  mean {np.mean(vals):+.2f}",
                transform=ax.transAxes, ha="right", fontsize=6.5, color=c)
    ax.set_xlabel("ESM-2 mean z-PLL"); ax.set_yticks([])
    for sp in ["left", "top", "right"]: ax.spines[sp].set_visible(False)
    panel(ax, "a", "PLL by epitope class")

    # b: dataflow
    ax = axes[1]; ax.set_xlim(0, 10); ax.set_ylim(0, 4); ax.axis("off")
    steps = [("IEDB B-cell\n(122)", 0.3, "#5B6672"),
             ("HA/NA\n(70)", 2.6, "#5B6672"),
             ("Mapped\n(33)", 4.9, "#7A8794"),
             ("Clusters\n(22)", 7.2, RED)]
    for i, (txt, x, c) in enumerate(steps):
        ax.add_patch(plt.Rectangle((x, 1.5), 1.7, 1.0, fc=c, ec="none", alpha=0.9))
        ax.text(x+0.85, 2.0, txt, ha="center", va="center", fontsize=6.5, color="white", fontweight="bold")
        if i < 3:
            ax.annotate("", xy=(x+1.75, 2.0), xytext=(x+2.5, 2.0),
                        arrowprops=dict(arrowstyle="-|>", color=GRAY, lw=1))
    panel(ax, "b", "B-cell data flow")
    save(fig, f"{MAIN}/Fig5_bcell_benchmark.png")

# ================================================================ Fig 6 (1×2)
def fig6():
    fig, axes = plt.subplots(1, 2, figsize=(7.2, 3.6))
    # a: stalk footprints on HA backbone
    ax = axes[0]
    rows = list(csv.DictReader(open(f"{MS}/08_BCELL_EXTERNAL_BENCHMARK/structures/09_NEUTRALIZING_FOOTPRINTS.tsv"), delimiter="\t"))
    cons_ha = str(list(SeqIO.parse(f"{CONS}/H1N1_HA.fasta", "fasta"))[0].seq); L = len(cons_ha)
    ax.plot(range(1, L+1), np.zeros(L), color=INK, lw=1.5, alpha=0.7)
    ax.axvspan(344, L, color=GREEN, alpha=0.12); ax.axvspan(18, 343, color=GRAY, alpha=0.05)
    ax.text(180, -0.5, "HA1", ha="center", fontsize=6.5, color=GRAY)
    ax.text(440, -0.5, "HA2", ha="center", fontsize=6.5, color=GREEN)
    colors = {"CR6261": RED, "CR8020": ORANGE, "CR9114": BLUE}
    for k, r in enumerate(rows):
        pos = [int(x) for x in r["consensus_positions"].split(";")]
        ax.scatter(pos, [1.0 + k*1.0]*len(pos), s=18, color=colors[r["antibody"]], zorder=2,
                   label=f'{r["antibody"]} ({r["subtype"]})')
    ax.set_xlabel("HA consensus position"); ax.set_ylim(-1, 3.4); ax.set_yticks([])
    ax.legend(frameon=False, fontsize=6.5, loc="upper right")
    panel(ax, "a", "Stalk antibody footprints")

    # b: extended footprints (all 8)
    ax = axes[1]
    fps = list(csv.DictReader(open(f"{MS}/08_BCELL_EXTERNAL_BENCHMARK/structures/09_NEUTRALIZING_FOOTPRINTS_EXTENDED.tsv"), delimiter="\t"))
    ha = [f for f in fps if f["protein"] == "HA"]; na = [f for f in fps if f["protein"] == "NA"]
    ha.sort(key=lambda f: (f["region"] != "stalk", f["antibody"]))
    HA_LEN = 567
    for i, f in enumerate(ha):
        y = len(ha) - 1 - i
        ax.hlines(y, 0, HA_LEN, color=GRAY, lw=0.4, alpha=0.5)
        ax.vlines([int(x) for x in f["consensus_positions"].split(";")], y-0.25, y+0.25, color=COL[f["subtype"]], lw=1)
    ax.set_yticks(range(len(ha)))
    ax.set_yticklabels([f"{f['antibody']} ({f['subtype']})" for f in ha], fontsize=6)
    ax.set_xlim(0, HA_LEN); ax.set_xlabel("HA consensus position")
    ax2 = ax.twinx()
    NA_LEN = 470
    for i, f in enumerate(na):
        ax2.hlines(2 + i, 0, NA_LEN, color=GRAY, lw=0.4, alpha=0.5)
        ax2.vlines([int(x) for x in f["consensus_positions"].split(";")], 2+i-0.3, 2+i+0.3, color=COL[f["subtype"]], lw=1.3)
    ax2.set_yticks([2]); ax2.set_yticklabels([f"{na[0]['antibody']} ({na[0]['subtype']}) NA"], fontsize=6)
    ax2.spines["top"].set_visible(False); ax2.spines["right"].set_visible(False)
    panel(ax, "b", "Extended footprints (stalk + RBD + NA)")
    save(fig, f"{MAIN}/Fig6_antibody_footprints.png")

# ================================================================ Fig 7 (单面板)
def fig7():
    d = np.genfromtxt(f"{MS}/dnds_per_codon.tsv", delimiter="\t", skip_header=1)
    idx, dn_ds = d[:, 0], d[:, 3]
    dn_ds = np.clip(dn_ds, 0, 5)
    sm = np.convolve(dn_ds, np.ones(15)/15, mode="same")
    fig, ax = plt.subplots(figsize=(6.5, 2.6))
    ax.plot(idx, sm, color=BLUE, lw=1.2)
    ax.axhline(1, color=INK, ls="--", lw=0.7)
    ax.set_title("Per-codon dN/dS along H1N1 HA", loc="left", fontsize=9, pad=8)
    ax.set_xlabel("HA codon position"); ax.set_ylabel("dN/dS (smoothed)")
    ax.set_ylim(0, 3)
    save(fig, f"{MAIN}/Fig7_dnds.png")

# ================================================================ 附录图（NC 风格重画）
def supp_figs():
    rows = list(csv.DictReader(open(f"{RES}/ai_ranking.csv")))
    for r in rows:
        r["final_score"] = float(r["final_score"]); r["ai_conservation"] = float(r["ai_conservation"])
    # S1 ranking
    rows2 = [r for r in rows if not np.isnan(r["ai_conservation"])]
    rows2.sort(key=lambda r: -r["final_score"]); top = rows2[:12]
    names = [f'{r["protein"]} {r["epitope"][:11]}' for r in top]
    fig, ax = plt.subplots(figsize=(4, 3.6))
    y = np.arange(len(top))[::-1]
    ax.barh(y, [r["final_score"] for r in top], color=[RED if r["protein"] in ("HA","NA") else BLUE for r in top], height=0.6)
    ax.set_yticks(y); ax.set_yticklabels(names, fontsize=6.5); ax.set_xlabel("Combined score")
    save(fig, f"{SUPP}/S1_ranking_sensitivity.png")

    # S2 assemblies
    fig, ax = plt.subplots(figsize=(6, 2.8)); ax.set_xlim(0, 10); ax.set_ylim(0, 3.2); ax.axis("off")
    rows3 = [("HA", ["RBD", "antigenic 2"], RED), ("NA", ["A", "B", "C", "D"], GREEN), ("NP / M1 / NEP", ["NP core", "M1", "NEP"], BLUE)]
    y = 2.6
    for name, segs, c in rows3:
        ax.text(0.1, y, name, fontsize=8, fontweight="bold", va="center")
        x = 1.8
        for s in segs:
            w = 1.6; ax.add_patch(FancyBboxPatch((x, y-0.4), w, 0.8, boxstyle="round,pad=0.05,rounding_size=0.14", fc=c, ec="none", alpha=0.85))
            ax.text(x+w/2, y, s, ha="center", va="center", fontsize=6, color="white"); x += w+0.12
        y -= 0.85
    save(fig, f"{SUPP}/S2_assemblies.png")

    # S3 circular map
    prots = ["HA","NA","M1","NEP","NP","NS1","PA","PB1","PB2"]
    fig, ax = plt.subplots(figsize=(6, 6), subplot_kw={"projection": "polar"})
    ax.set_theta_zero_location("N"); ax.set_theta_direction(-1)
    n = len(prots)
    for i, p in enumerate(prots):
        theta = 2*np.pi*i/n
        ax.bar(theta, 1.0, width=2*np.pi/n*0.85, bottom=0.5, color=GRAY, alpha=0.15)
        ax.text(theta, 1.7, p, ha="center", va="center", fontsize=10, fontweight="bold", color=INK)
        cand = [r for r in rows if r["protein"] == p]
        for j, r in enumerate(cand[:8]):
            t2 = theta + (j/(max(len(cand[:8]), 1)-1)-0.5)*2*np.pi/n*0.6 if len(cand[:8]) > 1 else theta
            ax.bar(t2, 0.5, width=0.04, bottom=0.7, color=RED if p in ("HA","NA") else BLUE, alpha=0.8)
    ax.set_ylim(0, 1.9); ax.set_yticks([])
    save(fig, f"{SUPP}/S3_circular_map.png")

    # S4 heterogeneity
    fig, axes = plt.subplots(1, 3, figsize=(7, 2.6))
    for ax, sub in zip(axes, ["H1N1", "H3N2", "H5N1"]):
        lens = [len(r.seq) for r in SeqIO.parse(f"{RAW}/分类序列_{sub}/HA_sequences.fasta", "fasta")]
        ax.hist(lens, bins=40, color=COL[sub], alpha=0.8)
        ax.axvline(1701, color=RED, ls="--", lw=1)
        ax.set_xlabel("HA nt length"); ax.set_ylabel("Count")
    save(fig, f"{SUPP}/S4_length_heterogeneity.png")

    # S5 HLA coverage
    fig, (ax1, ax2) = plt.subplots(1, 2, figsize=(5, 2.6))
    ax1.pie([94.4, 5.6], colors=[GREEN, "#E0E4E8"], startangle=90, counterclock=False, wedgeprops=dict(width=0.35))
    ax1.text(0, 0.02, "94.4%", ha="center", va="center", fontsize=14, fontweight="bold")
    ax1.text(0, -0.3, "14 HLA-I", ha="center", fontsize=6.5)
    ax2.pie([73.0, 27.0], colors=[BLUE, "#E0E4E8"], startangle=90, counterclock=False, wedgeprops=dict(width=0.35))
    ax2.text(0, 0.02, "73.0%", ha="center", va="center", fontsize=14, fontweight="bold")
    ax2.text(0, -0.3, "6 HLA-DRB1", ha="center", fontsize=6.5)
    save(fig, f"{SUPP}/S5_hla_coverage.png")

if __name__ == "__main__":
    fig1(); fig2(); fig3(); fig4(); fig5(); fig6(); fig7(); supp_figs()
    print("ALL NC FIGURES DONE")
```
