## Supplementary material for "Conserved influenza A epitope candidate regions and a benchmark of ESM-2 sequence features": Li_et_al_Supplementary_Data: make_figures_nc_optimized.py

```
#!/usr/bin/env python3
"""
NC (Nature Communications) 风格图重做：
- Arial 字体，无内嵌标题（图注放正文）
- 面板角标 a/b/c/d：小写加粗，左上角，无背景框，位置统一对称
- 克制配色：分类色 BLUE/ORANGE/GREEN/RED，热图用单 hue 'Blues'，3D 用 'RdBu_r'
- 细线、刻度朝外、充足留白

直接绘制 7 张主文多面板图 + 5 张附录图（不拼接 PNG）。
"""
import os, csv, re, io, shutil, subprocess, tempfile, warnings
from collections import Counter, defaultdict
import numpy as np
import matplotlib
matplotlib.use("Agg")
import matplotlib.pyplot as plt
from matplotlib import font_manager
import matplotlib.image as mpimg
from matplotlib.patches import FancyBboxPatch
from Bio import SeqIO
from scipy import stats

ROOT = os.environ.get("FLU_PROJECT_ROOT", "/media/qxl/My_Passport/流行病学数据/广谱流感疫苗候选课题")
RES = f"{ROOT}/08_ESM2_AI分析与论文/results"
ESM = f"{RES}/esm2"
CONS = f"{ROOT}/08_ESM2_AI分析与论文/data/consensus"
RAW = f"{ROOT}/01_原始序列与分层数据/序列与地区分层/全球/序列/综合"
MS = f"{ROOT}/07_MANUSCRIPT_STRENGTHENING"
FIG = f"{MS}/figures"
MAIN = f"{FIG}/main"
SUPP = f"{FIG}/supplementary"
PDB_1RU7 = os.environ.get("PDB_1RU7", "/tmp/1RU7.pdb")
PDB_2HTY = os.environ.get("PDB_2HTY", "/tmp/2HTY.pdb")

# 期刊图建议同时保留位图和矢量版本；如只需要 PNG，可设为 False。
EXPORT_PDF = True
SHOW_PANEL_TITLES = False
MAX_HEATMAP_ROWS = 20

os.makedirs(MAIN, exist_ok=True)
os.makedirs(SUPP, exist_ok=True)

BLUE="#3A6EA5"; ORANGE="#C77C3F"; GREEN="#4E8D5C"; RED="#B5453B"; GRAY="#8A8F98"; INK="#22262B"
COL={"H1N1":BLUE,"H3N2":ORANGE,"H5N1":GREEN}

# ---- NC 风格 rcParams ----
# Linux 环境经常没有 Arial。优先 Arial，否则使用 Liberation Sans/DejaVu Sans，避免反复 font warning。
available_fonts = {f.name for f in font_manager.fontManager.ttflist}
FONT = "Arial" if "Arial" in available_fonts else ("Liberation Sans" if "Liberation Sans" in available_fonts else "DejaVu Sans")
if FONT != "Arial":
    warnings.warn(f"Arial not found; using {FONT}. For final submission, install Arial if the journal requires it.")

plt.rcParams.update({
    "font.family": "sans-serif",
    "font.sans-serif": [FONT, "Arial", "Liberation Sans", "DejaVu Sans"],
    "font.size": 8,
    "axes.spines.top": False,
    "axes.spines.right": False,
    "axes.linewidth": 0.6,
    "axes.edgecolor": INK, "axes.labelcolor": INK, "text.color": INK,
    "xtick.color": INK, "ytick.color": INK,
    "xtick.direction": "out", "ytick.direction": "out",
    "xtick.major.width": 0.6, "ytick.major.width": 0.6,
    "xtick.major.size": 3, "ytick.major.size": 3,
    "axes.axisbelow": True,
})

def z(a):
    """NaN-safe z-score；常量数组返回 0，避免除以 0。"""
    a = np.asarray(a, dtype=float)
    mu = np.nanmean(a)
    sd = np.nanstd(a)
    if not np.isfinite(sd) or sd == 0:
        return np.zeros_like(a, dtype=float)
    return (a - mu) / sd


def require_file(path):
    if not os.path.exists(path):
        raise FileNotFoundError(f"Required input not found: {path}")
    return path


def panel(ax, label, title=None):
    """Nature 风格面板角标；默认不在图内重复写面板标题。"""
    kw = dict(fontsize=12, fontweight="bold", va="bottom", ha="left")
    if hasattr(ax, "get_zlim"):          # 3D 轴用 text2D
        ax.text2D(-0.06, 1.08, label, transform=ax.transAxes, **kw)
    else:
        ax.text(-0.06, 1.08, label, transform=ax.transAxes, **kw)
    if SHOW_PANEL_TITLES and title is not None:
        ax.set_title(title, loc="left", fontsize=8.5, pad=10)


def save(fig, name, dpi=300):
    """保存 PNG；默认同时保存 PDF 矢量版本。"""
    fig.savefig(name, dpi=dpi, bbox_inches="tight", facecolor="white", pad_inches=0.02)
    if EXPORT_PDF:
        pdf = os.path.splitext(name)[0] + ".pdf"
        fig.savefig(pdf, bbox_inches="tight", facecolor="white", pad_inches=0.02)
    plt.close(fig)
    print("saved", os.path.basename(name))


def extract_year(meta_row=None, fallback_text=""):
    """从常见 metadata 字段或文本中提取 4 位年份。找不到则返回 None。"""
    texts = []
    if meta_row:
        priority = ["Year", "year", "Collection_Year", "collection_year", "Collection_Date", "collection_date", "Date", "date"]
        texts.extend(str(meta_row.get(k, "")) for k in priority)
        texts.extend(str(v) for v in meta_row.values())
    texts.append(str(fallback_text))
    for text in texts:
        m = re.search(r"(?<!\d)(19\d{2}|20\d{2})(?!\d)", text)
        if m:
            return int(m.group(1))
    return None


def translate_cds_frame0(seq):
    """按已知 CDS 的 frame 0 翻译；不再通过“第一个 ATG”猜起始位点。"""
    from Bio.Seq import Seq
    s = str(seq).upper().replace("-", "").replace(".", "")
    s = re.sub(r"[^ACGTN]", "N", s)
    s = s[:(len(s)//3)*3]
    if not s:
        return ""
    aa = str(Seq(s).translate())
    return aa.rstrip("*")


def rolling_nanmean(a, window=15):
    """边缘不补 0 的滑动均值，同时忽略 NaN/inf。"""
    a = np.asarray(a, dtype=float)
    finite = np.isfinite(a)
    vals = np.where(finite, a, 0.0)
    ker = np.ones(int(window), dtype=float)
    num = np.convolve(vals, ker, mode="same")
    den = np.convolve(finite.astype(float), ker, mode="same")
    out = np.full_like(a, np.nan, dtype=float)
    np.divide(num, den, out=out, where=den > 0)
    return out

# ================================================================ 数据助手
def load_cons():
    cons = {}
    for sub in ["H1N1","H3N2","H5N1"]:
        for prot in ["HA","NA","M1","NP","NEP","NS1","PA","PB1","PB2"]:
            p = f"{CONS}/{sub}_{prot}.fasta"
            if os.path.exists(p):
                cons[(sub, prot)] = str(list(SeqIO.parse(p, "fasta"))[0].seq)
    return cons

def best_identity(seq, epi):
    if not epi or len(seq) < len(epi): return 0.0
    b = 0.0
    for i in range(len(seq) - len(epi) + 1):
        b = max(b, sum(a == c for a, c in zip(epi, seq[i:i+len(epi)])) / len(epi))
    return b

def frac(seqs, epi, thresh=0.9):
    if not seqs: return np.nan
    return np.mean([best_identity(s, epi) >= thresh for s in seqs])

# ================================================================ Fig 1 (2×2)
def fig1():
    cons = load_cons()
    ranking_file = require_file(f"{RES}/ai_ranking.csv")
    rows = list(csv.DictReader(open(ranking_file, encoding="utf-8")))
    for r in rows:
        r["final_score"] = float(r["final_score"])
    rows.sort(key=lambda r: -r["final_score"])
    top = rows[:16]

    fig, axes = plt.subplots(2, 2, figsize=(7.2, 6.6))

    # --- a: cross-subtype identity heatmap ---
    ax = axes[0, 0]
    mat, labels = [], []
    for r in top:
        epi, prot = r["epitope"], ("M1" if r["protein"] == "M" else r["protein"])
        row = [best_identity(cons.get((sub, prot), ""), epi) for sub in ("H1N1", "H3N2", "H5N1")]
        mat.append(row)
        labels.append(f"{prot} {epi[:12]}")
    mat = np.asarray(mat, dtype=float)
    im = ax.imshow(mat, cmap="Blues", vmin=0.5, vmax=1.0, aspect="auto")
    ax.set_xticks(range(3)); ax.set_xticklabels(["H1N1", "H3N2", "H5N1"])
    ax.set_yticks(range(len(labels))); ax.set_yticklabels(labels, fontsize=6.5)
    for i in range(mat.shape[0]):
        for j in range(mat.shape[1]):
            # Blues: 高值背景更深，因此高值用白字、低值用深色字。
            ax.text(j, i, f"{mat[i, j]:.2f}", ha="center", va="center", fontsize=5.5,
                    color="white" if mat[i, j] >= 0.78 else INK)
    ax.tick_params(length=0)
    cb = fig.colorbar(im, ax=ax, fraction=0.046, pad=0.03)
    cb.set_label("Identity", fontsize=7); cb.ax.tick_params(labelsize=6)
    panel(ax, "a", "Cross-subtype conservation")

    # --- b: temporal stability ---
    ax = axes[0, 1]

    def year_seqs(sub, prot):
        ys = defaultdict(list)
        fasta = require_file(f"{RAW}/分类序列_{sub}/{prot}_sequences.fasta")
        for rec in SeqIO.parse(fasta, "fasta"):
            year = extract_year(fallback_text=rec.description)
            if year is None or not (2006 <= year <= 2017):
                continue
            aa = translate_cds_frame0(rec.seq)
            if aa and aa.count("X") <= 0.05 * len(aa):
                ys[year].append(aa)
        return ys

    epi = "CKLRGVAPLHLGKCNIAGW"
    ys = year_seqs("H1N1", "HA")
    years = sorted(ys)
    fracs = [frac(ys[y], epi) for y in years]
    if years:
        ax.plot(years, fracs, "o-", color=RED, lw=1.4, ms=4)
        finite = np.isfinite(fracs)
        if np.sum(finite) >= 2:
            xfit = np.asarray(years)[finite]
            yfit = np.asarray(fracs)[finite]
            slope, inter, *_ = stats.linregress(xfit, yfit)
            ax.plot(xfit, slope*xfit + inter, "--", color=GRAY, lw=1)
    else:
        ax.text(0.5, 0.5, "No valid 2006–2017 records", transform=ax.transAxes,
                ha="center", va="center", fontsize=7, color=GRAY)
    ax.set_xlabel("Year"); ax.set_ylabel("Fraction ≥ 90% identity")
    ax.set_ylim(0.4, 1.05); ax.set_yticks([0.5, 0.7, 0.9])
    panel(ax, "b", "Temporal stability (H1N1 HA)")

    # --- c: H1N1 cross-lineage, strictly 2015–2025 ---
    ax = axes[1, 0]
    meta_file = require_file(f"{ROOT}/99_历史项目与参考/早期H1N1系统发育项目/清洗数据/cleaned_metadata.csv")
    meta = {}
    for rr in csv.DictReader(open(meta_file, encoding="utf-8")):
        if "Isolate_Id" in rr:
            meta[rr["Isolate_Id"]] = rr

    aln = require_file(f"{ROOT}/99_历史项目与参考/早期H1N1系统发育项目/清洗数据/clustalo-H1N1-HA.fa")
    by_clade = defaultdict(list)
    skipped_no_year = 0
    for rec in SeqIO.parse(aln, "fasta"):
        mr = meta.get(rec.id, {})
        year = extract_year(mr, rec.description)
        if year is None:
            skipped_no_year += 1
            continue
        if not (2015 <= year <= 2025):
            continue
        cl = mr.get("Clade", "unassigned") or "unassigned"
        aa = translate_cds_frame0(rec.seq)
        if aa and aa.count("X") <= 0.05 * len(aa):
            by_clade[cl].append(aa)

    top_clades = [c for c, _ in Counter({k: len(v) for k, v in by_clade.items()}).most_common(6)]
    if top_clades:
        f_clades = [frac(by_clade[c], epi) for c in top_clades]
        xs = np.arange(len(top_clades))
        ax.bar(xs, f_clades, color=BLUE, width=0.55)
        ax.set_xticks(xs)
        ax.set_xticklabels([c.replace(".", ".\n", 1) for c in top_clades], fontsize=5.5)
    else:
        ax.text(0.5, 0.5, "No valid 2015–2025 clade records", transform=ax.transAxes,
                ha="center", va="center", fontsize=7, color=GRAY)
    if skipped_no_year:
        print(f"Fig1c: skipped {skipped_no_year} records with no parseable year")
    ax.set_ylabel("Conserved fraction"); ax.set_ylim(0, 1.1)
    ax.axhline(0.9, color=GRAY, ls="--", lw=0.8)
    panel(ax, "c", "Cross-lineage conservation (H1N1)")

    # --- d: H3N2 subtype-specificity (surface candidates × clades) ---
    ax = axes[1, 1]
    h3_file = require_file(f"{MS}/09_H3N2_CROSS_LINEAGE/h3n2_cross_lineage.tsv")
    h3 = {}
    for rr in csv.DictReader(open(h3_file, encoding="utf-8"), delimiter="	"):
        h3[(rr["protein"], rr["epitope"])] = rr
    ckeys = sorted({k for rr in h3.values() for k in rr if k.startswith("clade_")})

    # rows 已按 final_score 排序，只绘制有 cross-lineage 数据的前 N 个表面候选，避免热图不可读。
    surf = [r for r in rows if r["protein"] in ("HA", "NA") and (r["protein"], r["epitope"]) in h3]
    if len(surf) > MAX_HEATMAP_ROWS:
        print(f"Fig1d: {len(surf)} surface candidates available; plotting top {MAX_HEATMAP_ROWS} by final_score")
        surf = surf[:MAX_HEATMAP_ROWS]

    B = np.full((len(surf), len(ckeys)), np.nan, dtype=float)
    slab = []
    for i, r in enumerate(surf):
        hr = h3.get((r["protein"], r["epitope"]), {})
        for j, k in enumerate(ckeys):
            v = hr.get(k, "")
            if v not in ("", None, "NA", "NaN", "nan"):
                try:
                    B[i, j] = float(v)
                except ValueError:
                    pass
        slab.append(f"{r['protein']} {r['epitope'][:10]}")

    cmap = plt.get_cmap("Blues").copy()
    cmap.set_bad("#ECEFF1")
    im = ax.imshow(np.ma.masked_invalid(B), cmap=cmap, vmin=0, vmax=1, aspect="auto")
    ax.set_xticks(range(len(ckeys)))
    ax.set_xticklabels([c.replace("clade_", "").replace(".", ".\n", 1) for c in ckeys], fontsize=5.5)
    ax.set_yticks(range(len(slab))); ax.set_yticklabels(slab, fontsize=6)
    for i in range(B.shape[0]):
        for j in range(B.shape[1]):
            if np.isfinite(B[i, j]):
                ax.text(j, i, f"{B[i, j]:.2f}", ha="center", va="center", fontsize=5.2,
                        color="white" if B[i, j] >= 0.58 else INK)
    ax.tick_params(length=0)
    cb = fig.colorbar(im, ax=ax, fraction=0.046, pad=0.03)
    cb.set_label("Conserved fraction", fontsize=7); cb.ax.tick_params(labelsize=6)
    panel(ax, "d", "H3N2 clade conservation")

    save(fig, f"{MAIN}/Fig1_candidate_conservation.png")

# ================================================================ Fig 2 (1×2)
def fig2():
    """ESM-2 PLL 与序列保守性、结构位置的关系。"""
    fig = plt.figure(figsize=(7.2, 3.4))
    gs = fig.add_gridspec(1, 2, width_ratios=[1.0, 1.05], wspace=0.32)
    ax = fig.add_subplot(gs[0, 0])
    ax3 = fig.add_subplot(gs[0, 1], projection="3d")

    def protein_sequences(sub, prot):
        fasta = require_file(f"{RAW}/分类序列_{sub}/{prot}_sequences.fasta")
        aa = []
        for rec in SeqIO.parse(fasta, "fasta"):
            seq = translate_cds_frame0(rec.seq)
            if seq and seq.count("X") <= 0.05 * len(seq):
                aa.append(seq)
        return aa

    def positional_identity(sub, prot):
        """优先使用 MAFFT 做 MSA；没有 MAFFT 时回退到等长序列的逐位统计，并明确 warning。"""
        seqs = protein_sequences(sub, prot)
        if not seqs:
            return np.array([], dtype=float)
        cons_file = require_file(f"{CONS}/{sub}_{prot}.fasta")
        consensus = str(list(SeqIO.parse(cons_file, "fasta"))[0].seq)

        mafft = shutil.which("mafft")
        if mafft:
            with tempfile.NamedTemporaryFile("w", suffix=".faa", delete=False) as fh:
                tmp = fh.name
                fh.write(">__CONSENSUS__\n" + consensus + "\n")
                for i, seq in enumerate(seqs):
                    fh.write(f">seq_{i}\n{seq}\n")
            try:
                cp = subprocess.run([mafft, "--auto", "--quiet", tmp], check=True,
                                    stdout=subprocess.PIPE, stderr=subprocess.PIPE, text=True)
                aln = list(SeqIO.parse(io.StringIO(cp.stdout), "fasta"))
            finally:
                try:
                    os.unlink(tmp)
                except OSError:
                    pass
            if not aln:
                raise RuntimeError(f"MAFFT returned an empty alignment for {sub} {prot}")
            ref = next((str(r.seq) for r in aln if r.id == "__CONSENSUS__"), None)
            if ref is None:
                raise RuntimeError("Consensus sequence missing from MAFFT output")
            aln_seqs = [str(r.seq) for r in aln if r.id != "__CONSENSUS__"]
            out = []
            for col, refaa in enumerate(ref):
                if refaa == "-":
                    continue
                chars = [x[col] for x in aln_seqs if x[col] not in ("-", "X")]
                if not chars:
                    out.append(np.nan)
                else:
                    out.append(Counter(chars).most_common(1)[0][1] / len(chars))
            return np.asarray(out, dtype=float)

        warnings.warn(
            f"MAFFT not found: {sub} {prot} conservation falls back to equal-length sequences. "
            "Install MAFFT for publication-grade MSA positional identity."
        )
        ln = Counter(len(x) for x in seqs).most_common(1)[0][0]
        seqs = [x for x in seqs if len(x) == ln]
        L = min(ln, len(consensus))
        return np.asarray([
            Counter(x[i] for x in seqs if x[i] != "X").most_common(1)[0][1] /
            max(1, sum(x[i] != "X" for x in seqs))
            for i in range(L)
        ], dtype=float)

    xs, ys = [], []
    for sub in ["H1N1", "H3N2", "H5N1"]:
        pll = np.load(require_file(f"{ESM}/{sub}_HA_pll.npy"))
        seq = open(require_file(f"{ESM}/{sub}_HA_seq.txt"), encoding="utf-8").read().strip()
        ci = positional_identity(sub, "HA")
        n = min(len(seq), len(ci), max(0, len(pll) - 1))
        if n == 0:
            continue
        zz = z(pll[1:1+n])
        valid = np.isfinite(ci[:n]) & np.isfinite(zz)
        ax.scatter(ci[:n][valid], zz[valid], s=4, color=COL[sub], alpha=0.35, lw=0, label=sub)
        xs.extend(ci[:n][valid]); ys.extend(zz[valid])

    if len(xs) >= 3:
        rho, _ = stats.spearmanr(xs, ys)
        ax.text(0.03, 0.95, f"Spearman ρ = {rho:.2f}", transform=ax.transAxes,
                fontsize=9, fontweight="bold", va="top")
    ax.set_xlabel("Positional identity")
    ax.set_ylabel("ESM-2 z-PLL")
    ax.legend(frameon=False, fontsize=6.5, markerscale=2, loc="lower left")
    panel(ax, "a", "PLL vs. sequence conservation")

    from Bio.PDB import PDBParser
    from Bio.Align import PairwiseAligner
    T3 = {'ALA':'A','ARG':'R','ASN':'N','ASP':'D','CYS':'C','GLN':'Q','GLU':'E','GLY':'G','HIS':'H','ILE':'I','LEU':'L','LYS':'K','MET':'M','PHE':'F','PRO':'P','SER':'S','THR':'T','TRP':'W','TYR':'Y','VAL':'V'}
    cons_ha = str(list(SeqIO.parse(require_file(f"{CONS}/H1N1_HA.fasta"), "fasta"))[0].seq)
    pll = np.load(require_file(f"{ESM}/H1N1_HA_pll.npy")); zpll = z(pll)
    al = PairwiseAligner(); al.mode = "local"; al.match_score = 2; al.mismatch_score = -1; al.open_gap_score = -2; al.extend_gap_score = -1
    s3d = PDBParser(QUIET=True).get_structure("1RU7", require_file(PDB_1RU7))[0]
    pts, cols = [], []
    for cid in ["A", "B"]:
        if cid not in s3d:
            continue
        ch = s3d[cid]; seq3d = ""; residues = []
        for res in ch:
            if res.id[0] != " ":
                continue
            aa = T3.get(res.resname.strip())
            if aa is None or "CA" not in res:
                continue
            seq3d += aa; residues.append(res)
        if not seq3d:
            continue
        a = al.align(cons_ha, seq3d)[0]
        tb, qb = a.aligned
        q2t = {}
        for (ts, te), (qs, qe) in zip(tb, qb):
            for off in range(min(te-ts, qe-qs)):
                q2t[qs+off] = ts+off
        for qi, res in enumerate(residues):
            ci = q2t.get(qi)
            if ci is None:
                continue
            pll_idx = ci + 1
            if pll_idx >= len(zpll):
                continue
            pts.append(res["CA"].coord)
            cols.append(zpll[pll_idx])

    pts = np.asarray(pts); cols = np.asarray(cols)
    if len(pts):
        sc = ax3.scatter(pts[:, 0], pts[:, 1], pts[:, 2], c=cols, cmap="RdBu_r", s=14, vmin=-2, vmax=2)
        cb = fig.colorbar(sc, ax=ax3, fraction=0.035, pad=0.02)
        cb.set_label("ESM-2 z-PLL", fontsize=7)
        cb.ax.tick_params(labelsize=6)
    ax3.set_axis_off(); ax3.view_init(elev=20, azim=-60)
    panel(ax3, "b", "HA structure colored by PLL")

    save(fig, f"{MAIN}/Fig2_esm_pll.png")

# ================================================================ Fig 3 (单面板)
def fig3():
    lines = open(f"{MS}/01_ESM_INCREMENTAL_BENCHMARK/benchmark_results.tsv").read().splitlines()
    rows = {}
    for l in lines[1:]:
        p = l.split("\t")
        if p[0].startswith("incremental"): continue
        rows[p[0]] = dict(auc=float(p[1]), lo=float(p[2]), hi=float(p[3]))
    keys = ['A_conservation', 'C_conservation+PLL', 'D_conservation+PLL+contact']
    labels = ['conservation', '+ PLL', '+ PLL\n+ contact']
    aucs = [rows[k]['auc'] for k in keys]; los = [rows[k]['lo'] for k in keys]; his = [rows[k]['hi'] for k in keys]
    fig, ax = plt.subplots(figsize=(3.6, 3.2))
    x = np.arange(3)
    ax.bar(x, aucs, yerr=[[a-l for a, l in zip(aucs, los)], [h-a for a, h in zip(aucs, his)]],
           color=[GRAY, BLUE, ORANGE], capsize=3, width=0.5, error_kw=dict(lw=0.8, ecolor=INK))
    ax.axhline(0.5, color=INK, ls="--", lw=0.7)
    ax.set_xticks(x); ax.set_xticklabels(labels, fontsize=7); ax.set_ylabel("AUROC")
    ymin = min(0.45, min(los) - 0.03); ymax = max(0.65, max(his) + 0.04)
    ax.set_ylim(ymin, ymax)
    for i, a in enumerate(aucs): ax.text(i, a+0.012, f"{a:.3f}", ha="center", fontsize=7)
    save(fig, f"{MAIN}/Fig3_tcell_benchmark.png")

# ================================================================ Fig 4 (单面板)
def fig4():
    fig, ax = plt.subplots(figsize=(3.8, 3.4))
    ha = np.genfromtxt(f"{RES}/rsa_validation.csv", delimiter=",", skip_header=1)
    rsa_ha, cd_ha = ha[:,0], ha[:,1]
    r_ha, _ = stats.spearmanr(rsa_ha, cd_ha)
    ax.scatter(rsa_ha, cd_ha, s=4, color=RED, alpha=0.4, lw=0, label=f"HA (ρ = {r_ha:+.2f})")
    # NA
    import freesasa
    from Bio.PDB import PDBParser
    T3 = {'ALA':'A','ARG':'R','ASN':'N','ASP':'D','CYS':'C','GLN':'Q','GLU':'E','GLY':'G','HIS':'H','ILE':'I','LEU':'L','LYS':'K','MET':'M','PHE':'F','PRO':'P','SER':'S','THR':'T','TRP':'W','TYR':'Y','VAL':'V'}
    REF = {'A':129.0,'R':274.0,'N':195.0,'D':193.0,'C':167.0,'Q':225.0,'E':223.0,'G':104.0,'H':224.0,'I':197.0,'L':201.0,'K':236.0,'M':224.0,'F':240.0,'P':159.0,'S':155.0,'T':172.0,'W':285.0,'Y':263.0,'V':174.0}
    res = freesasa.calc(freesasa.Structure(require_file(PDB_2HTY)))
    ca = res.residueAreas().get("A", {})
    ch = PDBParser(QUIET=True).get_structure("2HTY", require_file(PDB_2HTY))[0]["A"]
    na_seq, na_rsa = "", []
    for r_ in ch:
        if r_.id[0] != " ": continue
        aa = T3.get(r_.resname.strip())
        if aa is None: continue
        ra = ca.get(str(r_.id[1]))
        na_seq += aa; na_rsa.append(ra.total/REF.get(aa, 200.0) if ra else np.nan)
    from Bio.Align import PairwiseAligner
    cons_na = str(list(SeqIO.parse(f"{CONS}/H1N1_NA.fasta", "fasta"))[0].seq)
    al = PairwiseAligner(); al.mode = "local"; al.match_score = 2; al.mismatch_score = -1; al.open_gap_score = -2; al.extend_gap_score = -1
    a = al.align(cons_na, na_seq)[0]; tb, qb = a.aligned; q2t = {}
    for (ts, te), (qs, qe) in zip(tb, qb):
        for off in range(te-ts): q2t[qs+off] = ts+off
    acc_na = np.load(f"{ESM}/H1N1_NA_acc.npy")
    x, y = [], []
    for qi, ci in q2t.items():
        if qi < len(na_rsa) and not np.isnan(na_rsa[qi]): x.append(na_rsa[qi]); y.append(acc_na[ci])
    r_na, _ = stats.spearmanr(x, y)
    ax.scatter(x, y, s=4, color=BLUE, alpha=0.4, lw=0, label=f"NA (ρ = {r_na:+.2f})")
    ax.set_xlabel("Relative solvent accessibility (FreeSASA)"); ax.set_ylabel("ESM-2 contact-density")
    ax.legend(frameon=False, fontsize=7)
    save(fig, f"{MAIN}/Fig4_contact_rsa.png")

# ================================================================ Fig 5 (1×2)
def fig5():
    fig, axes = plt.subplots(1, 2, figsize=(7.2, 3.4))
    # a: distribution comparison
    c06 = list(csv.DictReader(open(require_file(f"{MS}/08_BCELL_EXTERNAL_BENCHMARK/06_BENCHMARK_READY_DATASET.tsv"), encoding="utf-8"), delimiter="	"))
    bg = list(csv.DictReader(open(require_file(f"{MS}/08_BCELL_EXTERNAL_BENCHMARK/07_MATCHED_BACKGROUND.tsv"), encoding="utf-8"), delimiter="	"))
    cons = {}
    for sub in ["H1N1", "H3N2", "H5N1"]:
        for prot in ["HA", "NA"]:
            cons[(sub, prot)] = str(list(SeqIO.parse(require_file(f"{CONS}/{sub}_{prot}.fasta"), "fasta"))[0].seq)

    def region_pll(sub, prot, seq):
        s0 = cons[(sub, prot)]; i = s0.find(seq)
        if i < 0:
            for j in range(len(s0)-len(seq)+1):
                if sum(a != b for a, b in zip(seq, s0[j:j+len(seq)])) <= 1:
                    i = j; break
        if i < 0:
            return np.nan
        pll = np.load(require_file(f"{ESM}/{sub}_{prot}_pll.npy")); zz = z(pll)
        vals = zz[i+1:i+1+len(seq)]
        return np.nanmean(vals) if len(vals) else np.nan

    neut, bind = [], []
    for c in c06:
        v = region_pll(c["subtype"], c["protein"], c["representative_seq"])
        if np.isnan(v):
            continue
        (neut if c["label_level"] == "L3_neutralization" else bind).append(v)
    bgr = [region_pll(r["subtype"], r["protein"], r["bg_sequence"]) for r in bg]
    bgr = [v for v in bgr if np.isfinite(v)]
    groups = [("Neutralizing", neut, RED), ("Binding", bind, ORANGE), ("Background", bgr, GRAY)]

    ax = axes[0]
    for i, (name, vals, c) in enumerate(groups):
        vals = np.asarray(vals, dtype=float)
        if len(vals) == 0:
            continue
        ax.hist(vals, bins=22, color=c, alpha=0.45, density=True, histtype="stepfilled")
        ax.axvline(np.mean(vals), color=c, lw=1.1)
        ax.text(0.98, 0.92 - i*0.10, f"{name} (n={len(vals)})  mean {np.mean(vals):+.2f}",
                transform=ax.transAxes, ha="right", va="top", fontsize=6.5, color=c)
    ax.set_xlabel("ESM-2 mean z-PLL"); ax.set_yticks([])
    for sp in ["left", "top", "right"]:
        ax.spines[sp].set_visible(False)
    panel(ax, "a", "PLL by epitope class")

    # b: data flow
    ax = axes[1]; ax.set_xlim(0, 10); ax.set_ylim(0, 4); ax.axis("off")
    steps = [("IEDB B-cell\n(122)", 0.3, "#5B6672"),
             ("HA/NA\n(70)", 2.6, "#5B6672"),
             ("Mapped\n(33)", 4.9, "#7A8794"),
             ("Clusters\n(22)", 7.2, RED)]
    box_w = 1.7
    for i, (txt, x0, c) in enumerate(steps):
        ax.add_patch(plt.Rectangle((x0, 1.5), box_w, 1.0, fc=c, ec="none", alpha=0.9))
        ax.text(x0+box_w/2, 2.0, txt, ha="center", va="center", fontsize=6.5, color="white", fontweight="bold")
        if i < len(steps)-1:
            x_next = steps[i+1][1]
            ax.annotate("", xy=(x_next-0.08, 2.0), xytext=(x0+box_w+0.08, 2.0),
                        arrowprops=dict(arrowstyle="-|>", color=GRAY, lw=1))
    panel(ax, "b", "B-cell data flow")
    save(fig, f"{MAIN}/Fig5_bcell_benchmark.png")

# ================================================================ Fig 6 (1×2)
def fig6():
    # 左：3 个 stalk 抗体 footprint；右：HA/NA extended footprints 分开坐标轴，避免 twin-axis 尺度混淆。
    fig = plt.figure(figsize=(7.2, 3.8))
    outer = fig.add_gridspec(1, 2, width_ratios=[1.0, 1.15], wspace=0.35)
    ax = fig.add_subplot(outer[0, 0])

    # a: stalk footprints on HA backbone
    rows = list(csv.DictReader(open(require_file(f"{MS}/08_BCELL_EXTERNAL_BENCHMARK/structures/09_NEUTRALIZING_FOOTPRINTS.tsv"), encoding="utf-8"), delimiter="	"))
    cons_ha = str(list(SeqIO.parse(require_file(f"{CONS}/H1N1_HA.fasta"), "fasta"))[0].seq); L = len(cons_ha)
    ax.plot(range(1, L+1), np.zeros(L), color=INK, lw=1.5, alpha=0.7)
    ax.axvspan(344, L, color=GREEN, alpha=0.12); ax.axvspan(18, 343, color=GRAY, alpha=0.05)
    ax.text(180, -0.5, "HA1", ha="center", fontsize=6.5, color=GRAY)
    ax.text(440, -0.5, "HA2", ha="center", fontsize=6.5, color=GREEN)
    colors = {"CR6261": RED, "CR8020": ORANGE, "CR9114": BLUE}
    for k, r in enumerate(rows):
        pos = [int(x) for x in r["consensus_positions"].split(";") if x]
        ax.scatter(pos, [1.0 + k*1.0]*len(pos), s=18, color=colors.get(r["antibody"], GRAY), zorder=2,
                   label=f'{r["antibody"]} ({r["subtype"]})')
    ax.set_xlabel("HA consensus position"); ax.set_ylim(-1, max(3.4, len(rows)+0.4)); ax.set_yticks([])
    ax.legend(frameon=False, fontsize=6.5, loc="upper right")
    panel(ax, "a", "Stalk antibody footprints")

    # b: extended footprints, HA 与 NA 分开画，标签与数据使用同一顺序。
    fps = list(csv.DictReader(open(require_file(f"{MS}/08_BCELL_EXTERNAL_BENCHMARK/structures/09_NEUTRALIZING_FOOTPRINTS_EXTENDED.tsv"), encoding="utf-8"), delimiter="	"))
    ha = [f for f in fps if f["protein"] == "HA"]
    na = [f for f in fps if f["protein"] == "NA"]
    ha.sort(key=lambda f: (f.get("region", "") != "stalk", f["antibody"]))
    na.sort(key=lambda f: f["antibody"])

    right = outer[0, 1].subgridspec(2, 1, height_ratios=[max(len(ha), 1), max(len(na), 1)], hspace=0.55)
    ax_ha = fig.add_subplot(right[0, 0])
    ax_na = fig.add_subplot(right[1, 0])

    HA_LEN = 567
    for i, f in enumerate(ha):
        y = i
        ax_ha.hlines(y, 0, HA_LEN, color=GRAY, lw=0.4, alpha=0.5)
        pos = [int(x) for x in f["consensus_positions"].split(";") if x]
        ax_ha.vlines(pos, y-0.25, y+0.25, color=COL.get(f["subtype"], GRAY), lw=1)
    ax_ha.set_yticks(range(len(ha)))
    ax_ha.set_yticklabels([f"{f['antibody']} ({f['subtype']})" for f in ha], fontsize=6)
    ax_ha.set_xlim(0, HA_LEN); ax_ha.set_xlabel("HA consensus position", fontsize=7)
    ax_ha.tick_params(axis="x", labelsize=6)
    if len(ha) > 1:
        ax_ha.set_ylim(-0.6, len(ha)-0.4)
    panel(ax_ha, "b", "Extended footprints")

    NA_LEN = 470
    for i, f in enumerate(na):
        y = i
        ax_na.hlines(y, 0, NA_LEN, color=GRAY, lw=0.4, alpha=0.5)
        pos = [int(x) for x in f["consensus_positions"].split(";") if x]
        ax_na.vlines(pos, y-0.25, y+0.25, color=COL.get(f["subtype"], GRAY), lw=1.2)
    ax_na.set_yticks(range(len(na)))
    ax_na.set_yticklabels([f"{f['antibody']} ({f['subtype']})" for f in na], fontsize=6)
    ax_na.set_xlim(0, NA_LEN); ax_na.set_xlabel("NA consensus position", fontsize=7)
    ax_na.tick_params(axis="x", labelsize=6)
    if len(na) > 1:
        ax_na.set_ylim(-0.6, len(na)-0.4)

    save(fig, f"{MAIN}/Fig6_antibody_footprints.png")

# ================================================================ Fig 7 (单面板)
def fig7():
    d = np.genfromtxt(require_file(f"{MS}/dnds_per_codon.tsv"), delimiter="	", skip_header=1)
    idx, dn_ds = d[:, 0], d[:, 3]
    # 仅为可视化限制极端值；先把非有限值设为 NaN，再用边缘修正的 rolling mean。
    dn_ds = np.where(np.isfinite(dn_ds), np.clip(dn_ds, 0, 5), np.nan)
    sm = rolling_nanmean(dn_ds, window=15)
    fig, ax = plt.subplots(figsize=(6.5, 2.6))
    ax.plot(idx, sm, color=BLUE, lw=1.2)
    ax.axhline(1, color=INK, ls="--", lw=0.7)
    ax.set_xlabel("HA codon position"); ax.set_ylabel("dN/dS (15-codon mean)")
    finite = sm[np.isfinite(sm)]
    ymax = min(5.0, max(1.5, (np.nanpercentile(finite, 99) + 0.2) if len(finite) else 3.0))
    ax.set_ylim(0, ymax)
    save(fig, f"{MAIN}/Fig7_dnds.png")

# ================================================================ 附录图（NC 风格重画）
def supp_figs():
    rows = list(csv.DictReader(open(f"{RES}/ai_ranking.csv")))
    for r in rows:
        r["final_score"] = float(r["final_score"]); r["ai_conservation"] = float(r["ai_conservation"])
    # S1 ranking
    rows2 = [r for r in rows if not np.isnan(r["ai_conservation"])]
    rows2.sort(key=lambda r: -r["final_score"]); top = rows2[:12]
    names = [f'{r["protein"]} {r["epitope"][:11]}' for r in top]
    fig, ax = plt.subplots(figsize=(4, 3.6))
    y = np.arange(len(top))[::-1]
    ax.barh(y, [r["final_score"] for r in top], color=[RED if r["protein"] in ("HA","NA") else BLUE for r in top], height=0.6)
    ax.set_yticks(y); ax.set_yticklabels(names, fontsize=6.5); ax.set_xlabel("Combined score")
    save(fig, f"{SUPP}/S1_ranking_sensitivity.png")

    # S2 assemblies
    fig, ax = plt.subplots(figsize=(6, 2.8)); ax.set_xlim(0, 10); ax.set_ylim(0, 3.2); ax.axis("off")
    rows3 = [("HA", ["RBD", "antigenic 2"], RED), ("NA", ["A", "B", "C", "D"], GREEN), ("NP / M1 / NEP", ["NP core", "M1", "NEP"], BLUE)]
    y = 2.6
    for name, segs, c in rows3:
        ax.text(0.1, y, name, fontsize=8, fontweight="bold", va="center")
        x = 1.8
        for s in segs:
            w = 1.6; ax.add_patch(FancyBboxPatch((x, y-0.4), w, 0.8, boxstyle="round,pad=0.05,rounding_size=0.14", fc=c, ec="none", alpha=0.85))
            ax.text(x+w/2, y, s, ha="center", va="center", fontsize=6, color="white"); x += w+0.12
        y -= 0.85
    save(fig, f"{SUPP}/S2_assemblies.png")

    # S3 circular map
    prots = ["HA","NA","M1","NEP","NP","NS1","PA","PB1","PB2"]
    fig, ax = plt.subplots(figsize=(6, 6), subplot_kw={"projection": "polar"})
    ax.set_theta_zero_location("N"); ax.set_theta_direction(-1)
    n = len(prots)
    for i, p in enumerate(prots):
        theta = 2*np.pi*i/n
        ax.bar(theta, 1.0, width=2*np.pi/n*0.85, bottom=0.5, color=GRAY, alpha=0.15)
        ax.text(theta, 1.7, p, ha="center", va="center", fontsize=10, fontweight="bold", color=INK)
        cand = [r for r in rows if r["protein"] == p]
        for j, r in enumerate(cand[:8]):
            t2 = theta + (j/(max(len(cand[:8]), 1)-1)-0.5)*2*np.pi/n*0.6 if len(cand[:8]) > 1 else theta
            ax.bar(t2, 0.5, width=0.04, bottom=0.7, color=RED if p in ("HA","NA") else BLUE, alpha=0.8)
    ax.set_ylim(0, 1.9); ax.set_yticks([])
    save(fig, f"{SUPP}/S3_circular_map.png")

    # S4 heterogeneity
    fig, axes = plt.subplots(1, 3, figsize=(7, 2.6))
    for ax, sub in zip(axes, ["H1N1", "H3N2", "H5N1"]):
        lens = [len(r.seq) for r in SeqIO.parse(f"{RAW}/分类序列_{sub}/HA_sequences.fasta", "fasta")]
        ax.hist(lens, bins=40, color=COL[sub], alpha=0.8)
        ax.axvline(1701, color=RED, ls="--", lw=1)
        ax.set_xlabel("HA nt length"); ax.set_ylabel("Count")
    save(fig, f"{SUPP}/S4_length_heterogeneity.png")

    # S5 HLA coverage
    fig, (ax1, ax2) = plt.subplots(1, 2, figsize=(5, 2.6))
    ax1.pie([94.4, 5.6], colors=[GREEN, "#E0E4E8"], startangle=90, counterclock=False, wedgeprops=dict(width=0.35))
    ax1.text(0, 0.02, "94.4%", ha="center", va="center", fontsize=14, fontweight="bold")
    ax1.text(0, -0.3, "14 HLA-I", ha="center", fontsize=6.5)
    ax2.pie([73.0, 27.0], colors=[BLUE, "#E0E4E8"], startangle=90, counterclock=False, wedgeprops=dict(width=0.35))
    ax2.text(0, 0.02, "73.0%", ha="center", va="center", fontsize=14, fontweight="bold")
    ax2.text(0, -0.3, "6 HLA-DRB1", ha="center", fontsize=6.5)
    save(fig, f"{SUPP}/S5_hla_coverage.png")

def run_all():
    jobs = [fig1, fig2, fig3, fig4, fig5, fig6, fig7, supp_figs]
    errors = []
    for fn in jobs:
        try:
            print(f"[RUN] {fn.__name__}")
            fn()
        except Exception as exc:
            errors.append((fn.__name__, exc))
            print(f"[FAILED] {fn.__name__}: {type(exc).__name__}: {exc}")
    if errors:
        msg = "\n".join(f"- {name}: {type(exc).__name__}: {exc}" for name, exc in errors)
        raise RuntimeError(f"Some figures failed:\n{msg}")
    print("ALL NC FIGURES DONE")


if __name__ == "__main__":
    run_all()
```
