## Supplementary material for "Conserved influenza A epitope candidate regions and a benchmark of ESM-2 sequence features": Li_et_al_Supplementary_Data: make_figures_v2.py

```
#!/usr/bin/env python3
"""
主图 Figure 2–5 正式重设计（SCI 论文主图水平）。
原则：不改统计数值；标题无数字/年份/缩写/省略号；完整显示名称；
candidate 用规范 ID；蓝主/灰背景/橙辅助/红强调；SVG 矢量文字。
"""
import csv, re
from collections import Counter, defaultdict
import numpy as np
import matplotlib
matplotlib.use("Agg")
import matplotlib.pyplot as plt
from matplotlib.colors import ListedColormap
from Bio import SeqIO
from Bio.Seq import Seq
from scipy import stats
from scipy.stats import gaussian_kde

ROOT = "/media/qxl/My_Passport/流行病学数据/广谱流感疫苗候选课题"
RES = f"{ROOT}/08_ESM2_AI分析与论文/results"
ESM = f"{RES}/esm2"
CONS = f"{ROOT}/08_ESM2_AI分析与论文/data/consensus"
RAW = f"{ROOT}/01_原始序列与分层数据/序列与地区分层/全球/序列/综合"
MS = f"{ROOT}/07_MANUSCRIPT_STRENGTHENING"
FIG = f"{MS}/figures/main"

# ---- 颜色（蓝主/灰背景/橙辅助/红强调）----
BLUE = "#245F9E"; BLUE_DARK = "#174C84"; BLUE_LIGHT = "#AFCBE6"
GRAY = "#8D939C"; GRAY_LIGHT = "#D8DDE3"; GRID = "#D9DDE3"
ORANGE = "#E9832E"; ORANGE_LIGHT = "#F1C8A0"
RED = "#C63F36"; INK = "#142033"

plt.rcParams.update({
    "font.family": "Arial", "font.size": 8.4, "axes.titlesize": 9.5,
    "axes.labelsize": 8.2, "xtick.labelsize": 7.2, "ytick.labelsize": 7.2,
    "axes.linewidth": 0.65, "axes.edgecolor": "#6F7680", "text.color": INK,
    "xtick.color": INK, "ytick.color": INK,
    "xtick.direction": "out", "ytick.direction": "out",
    "xtick.major.width": 0.6, "ytick.major.width": 0.6,
    "axes.spines.top": False, "axes.spines.right": False,
    "pdf.fonttype": 42, "ps.fonttype": 42, "svg.fonttype": "none",
})

def panel(ax, letter, title):
    ax.text(-0.06, 1.04, letter, transform=ax.transAxes, fontsize=14, fontweight="bold",
            va="bottom", ha="right", color=BLUE_DARK)
    ax.set_title(title, loc="left", fontsize=9.5, pad=12, fontweight="bold")

def save(fig, name):
    for ext in ("svg", "pdf", "png"):
        kw = dict(bbox_inches="tight", facecolor="white", pad_inches=0.02)
        if ext == "png":
            kw["dpi"] = 600
        fig.savefig(f"{FIG}/{name}.{ext}", **kw)
    plt.close(fig)
    print(f"saved {name}.svg/.pdf/.png")

def z(a): return (a - np.nanmean(a)) / np.nanstd(a)
def best_identity(seq, epi):
    if len(seq) < len(epi): return 0.0
    return max(sum(a == c for a, c in zip(epi, seq[i:i+len(epi)])) / len(epi)
               for i in range(len(seq) - len(epi) + 1))
def frac(seqs, epi, thresh=0.9):
    if not seqs: return np.nan
    return np.mean([best_identity(s, epi) >= thresh for s in seqs])
def trans_atg(seq):
    s = str(seq).upper().replace("-", "")
    for i in range(len(s) - 2):
        if s[i:i+3] == "ATG":
            n = (len(s) - i) // 3; aa = []
            for c in range(n):
                cod = s[i+3*c:i+3*c+3]
                aa.append("X" if (len(cod) < 3 or "N" in cod) else str(Seq(cod).translate()))
            return "".join(aa)
    return ""

# ============================================================ 数据加载（一次算，供各图复用）
EPI = "CKLRGVAPLHLGKCNIAGW"

def load_temporal():
    ys = defaultdict(list)
    for r in SeqIO.parse(f"{RAW}/分类序列_H1N1/HA_sequences.fasta", "fasta"):
        mm = re.search(r"/20\d\d", r.description)
        if mm and 2006 <= int(mm.group(0)[1:]) <= 2017:
            ys[int(mm.group(0)[1:])].append(str(r.seq[:(len(r.seq)//3)*3].translate()))
    years = sorted(ys)
    return years, [len(ys[y]) for y in years], [frac(ys[y], EPI) for y in years]

def load_clades():
    meta = {}
    for r in csv.DictReader(open(f"{ROOT}/99_历史项目与参考/早期H1N1系统发育项目/清洗数据/cleaned_metadata.csv")):
        meta[r["Isolate_Id"]] = r
    aln = f"{ROOT}/99_历史项目与参考/早期H1N1系统发育项目/清洗数据/clustalo-H1N1-HA.fa"
    by_clade = defaultdict(list)
    for r in SeqIO.parse(aln, "fasta"):
        cl = meta.get(r.id, {}).get("Clade", "unassigned")
        aa = trans_atg(r.seq)
        if aa and aa.count("X") <= 0.05 * len(aa):
            by_clade[cl].append(aa)
    clades = [c for c, _ in Counter({k: len(v) for k, v in by_clade.items()}).most_common(6)]
    return clades, [len(by_clade[c]) for c in clades], [frac(by_clade[c], EPI) for c in clades]

def load_h3n2():
    h3 = list(csv.DictReader(open(f"{MS}/09_H3N2_CROSS_LINEAGE/h3n2_cross_lineage.tsv"), delimiter="\t"))
    ckeys = [k for k in h3[0].keys() if k.startswith("clade_")]
    cnames = [k.replace("clade_", "") for k in ckeys]
    mat = np.array([[float(r[k]) if r[k] not in ("", None) else 0.0 for k in ckeys] for r in h3])
    ids = [f"{r['protein']}-{r['length']}" for r in h3]  # 占位，稍后覆盖
    return cnames, mat, h3

def load_pll(sub):
    pll = np.load(f"{ESM}/{sub}_HA_pll.npy")
    seq = open(f"{ESM}/{sub}_HA_seq.txt").read().strip()
    return pll, seq

def mode_seqs(sub, prot):
    recs = list(SeqIO.parse(f"{RAW}/分类序列_{sub}/{prot}_sequences.fasta", "fasta"))
    aa = [str(r.seq[:(len(r.seq)//3)*3].translate()) for r in recs]
    ln = Counter(len(s) for s in aa).most_common(1)[0][0]
    return [s for s in aa if len(s) == ln and s.count("X") <= 0.05 * len(s)]

def cons_ident(sub, prot):
    seqs = mode_seqs(sub, prot); L = len(seqs[0])
    c = [Counter(s[i] for s in seqs).most_common(1)[0][0] for i in range(L)]
    return np.array([sum(1 for s in seqs if s[i] == c[i]) / len(seqs) for i in range(L)])

# ============================================================ Figure 2
def fig2():
    years, ns, fracs = load_temporal()
    clades, n_clade, cf_clade = load_clades()
    cnames, h3mat, h3 = load_h3n2()

    # 表面候选 ID（protein-rank）
    surf_rows = [r for r in csv.DictReader(open(f"{MS}/tables/Table2_candidates_full_corrected.tsv"), delimiter="\t")
                 if r["protein"] in ("HA", "NA")]
    h3lab = [f"{r['protein']}-{int(r['rank']):02d}" for r in surf_rows]

    fig = plt.figure(figsize=(12.5, 9.5))
    gs = fig.add_gridspec(2, 2, height_ratios=[1, 1.35], hspace=0.55, wspace=0.35)

    # ---- A 时间 ----
    ax = fig.add_subplot(gs[0, 0])
    size = 14 + 42 * (np.array(ns) / max(ns))
    ax.scatter(years, fracs, s=size, color=BLUE, alpha=0.85, edgecolor="white", linewidth=0.4, zorder=3)
    ax.plot(years, fracs, color=BLUE_LIGHT, lw=1.0, alpha=0.7, zorder=2)
    slope, inter, *_ = stats.linregress(years, fracs)
    ax.plot(years, [slope*y+inter for y in years], "--", color=GRAY, lw=1.0)
    ax.axhline(0.9, color=GRAY, lw=0.6, ls=":")
    ax.set_xlabel("Year"); ax.set_ylabel("Conserved fraction")
    ax.set_ylim(0, 1.1)
    ax.set_title("point size reflects sequence count", fontsize=6.5, color=GRAY, loc="right", pad=2)
    panel(ax, "A", "Temporal conservation of the selected HA candidate")

    # ---- B 谱系 lollipop ----
    ax = fig.add_subplot(gs[0, 1])
    yy = np.arange(len(clades))[::-1]
    ax.hlines(yy, 0, cf_clade, color=GRAY_LIGHT, lw=1.2, zorder=1)
    ax.scatter(cf_clade, yy, s=70, color=BLUE, zorder=3)
    ax.axvline(0.9, color=GRAY, lw=0.6, ls=":")
    ax.set_yticks(yy)
    ax.set_yticklabels([f"{c} (n={n})" for c, n in zip(clades, n_clade)], fontsize=6.5)
    ax.set_xlabel("Conserved fraction"); ax.set_xlim(0, 1.08)
    for y, f in zip(yy, cf_clade):
        ax.text(f + 0.015, y, f"{f:.2f}", va="center", fontsize=6.5)
    panel(ax, "B", "Conservation across circulating H1N1 lineages")

    # ---- C landscape heatmap ----
    ax = fig.add_subplot(gs[1, :])
    im = ax.imshow(h3mat, cmap=ListedColormap(["#F0F2F4", BLUE_LIGHT, BLUE_DARK, BLUE]),
                   aspect="auto", vmin=0, vmax=1, interpolation="nearest")
    ax.set_xticks(range(len(cnames)))
    ax.set_xticklabels(cnames, fontsize=6.5, rotation=30, ha="right")
    ax.set_yticks(range(len(h3lab)))
    ax.set_yticklabels(h3lab, fontsize=6.5)
    for i in range(h3mat.shape[0]):
        for j in range(h3mat.shape[1]):
            if h3mat[i, j] > 0.05:
                ax.text(j, i, f"{h3mat[i,j]:.2f}", ha="center", va="center",
                        fontsize=5.5, color="white" if h3mat[i, j] > 0.4 else INK)
    ax.tick_params(length=0)
    # HA/NA 小色条（行标签旁）
    for i, r in enumerate(surf_rows):
        c = ORANGE if r["protein"] == "HA" else BLUE
        ax.add_patch(plt.Rectangle((-0.55, i - 0.5), 0.28, 1.0, color=c, clip_on=False, lw=0))
    cb = fig.colorbar(im, ax=ax, fraction=0.025, pad=0.02)
    cb.set_label("Conserved fraction", fontsize=7); cb.ax.tick_params(labelsize=6)
    panel(ax, "C", "Candidate conservation across H3N2 lineages")

    # ---- D summary ----
    ax = fig.add_subplot(gs[1, 1])  # 占位，稍后调整
    fig.delaxes(ax)
    # 重新布局 D 放右下
    # （C 已占整行，D 单独一行下方）

    fig.suptitle("", fontsize=1)
    save(fig, "Figure2")
    # 打印校验
    print("F2 校验: A years", years, "n", ns)
    print("F2 校验: B clade frac", [round(f, 3) for f in cf_clade])

# ============================================================ 主入口
if __name__ == "__main__":
    fig2()
```
