## Supplementary material for "Conserved influenza A epitope candidate regions and a benchmark of ESM-2 sequence features": Li_et_al_Supplementary_Data: make_supplementary_figures.py

```
#!/usr/bin/env python3
"""
补充图 S1–S3：
S1 98候选完整证据热图；S2 时间与谱系验证；S3 模型与外部基准。
数据源：tables/Table2_candidates_full_corrected.tsv + 各分析结果文件。
"""
import csv, re
from collections import Counter, defaultdict
import numpy as np
import matplotlib
matplotlib.use("Agg")
import matplotlib.pyplot as plt
from matplotlib.colors import ListedColormap
from Bio import SeqIO
from Bio.Seq import Seq
from scipy import stats

ROOT = "/media/qxl/My_Passport/流行病学数据/广谱流感疫苗候选课题"
RES = f"{ROOT}/08_ESM2_AI分析与论文/results"
ESM = f"{RES}/esm2"
CONS = f"{ROOT}/08_ESM2_AI分析与论文/data/consensus"
RAW = f"{ROOT}/01_原始序列与分层数据/序列与地区分层/全球/序列/综合"
MS = f"{ROOT}/07_MANUSCRIPT_STRENGTHENING"
SUPP = f"{MS}/figures/supplementary"

BLUE = "#245F9E"; BLUE_DARK = "#174C84"; BLUE_LIGHT = "#AFCBE6"
ORANGE = "#C77C3F"; PURPLE = "#7B5AA6"; RED = "#C63F36"
SURFACE = "#1B9E8F"; INTERNAL = "#6B7B8C"; GRAY = "#8A8F98"; INK = "#22262B"
LIGHT = "#C8CDD3"
plt.rcParams.update({
    "font.family": "Arial", "font.size": 8, "axes.spines.top": False,
    "axes.spines.right": False, "axes.linewidth": 0.6, "axes.edgecolor": INK,
    "axes.labelcolor": INK, "text.color": INK, "xtick.color": INK, "ytick.color": INK,
    "xtick.direction": "out", "ytick.direction": "out",
    "xtick.major.width": 0.6, "ytick.major.width": 0.6,
})

def panel(ax, label, title):
    ax.text(-0.08, 1.06, label, transform=ax.transAxes, fontsize=13, fontweight="bold",
            va="bottom", ha="left")
    ax.set_title(title, loc="left", fontsize=9, pad=10)

def z(a): return (a - np.nanmean(a)) / np.nanstd(a)
def best_identity(seq, epi):
    if len(seq) < len(epi): return 0.0
    return max(sum(a == c for a, c in zip(epi, seq[i:i+len(epi)])) / len(epi)
               for i in range(len(seq) - len(epi) + 1))
def frac(seqs, epi, thresh=0.9):
    if not seqs: return np.nan
    return np.mean([best_identity(s, epi) >= thresh for s in seqs])
def trans_atg(seq):
    s = str(seq).upper().replace("-", "")
    for i in range(len(s) - 2):
        if s[i:i+3] == "ATG":
            n = (len(s) - i) // 3; aa = []
            for c in range(n):
                cod = s[i+3*c:i+3*c+3]
                aa.append("X" if (len(cod) < 3 or "N" in cod) else str(Seq(cod).translate()))
            return "".join(aa)
    return ""

rows = list(csv.DictReader(open(f"{MS}/tables/Table2_candidates_full_corrected.tsv"), delimiter="\t"))
for r in rows:
    r["rank"] = int(r["rank"]); r["final_score"] = float(r["final_score"])
    r["combined_BT"] = float(r["combined_BT"]); r["zPLL"] = float(r["zPLL"])
    r["contact_density"] = float(r["contact_density"])
rows.sort(key=lambda r: r["rank"]); N = len(rows)

def conv(v): return {"exact": 2, "fuzzy": 1, "-": 0}.get(v, 0)
cat = np.array([[conv(r["H1N1"]), conv(r["H3N2"]), conv(r["H5N1"])] for r in rows])
score = np.array([r["final_score"] for r in rows])
bt = np.array([r["combined_BT"] for r in rows])
zpll = np.array([r["zPLL"] for r in rows])
cd = np.array([r["contact_density"] for r in rows])
surf = np.array([1 if r["protein"] in ("HA", "NA") else 0 for r in rows])

# ================================================================ S1
def figS1():
    fig = plt.figure(figsize=(11, 9))
    gs = fig.add_gridspec(1, 8, width_ratios=[0.16, 0.5, 0.5, 1.1, 0.45, 0.45, 0.12, 0.55], wspace=0.05)
    axes = [fig.add_subplot(gs[0, i]) for i in range(8)]
    ax_rank, ax_score, ax_bt, ax_cat, ax_z, ax_cd, ax_surf, ax_prot = axes

    # rank 标注
    ax_rank.set_xlim(0, 1); ax_rank.set_ylim(-0.5, N - 0.5); ax_rank.axis("off")
    for i, r in enumerate(rows):
        ax_rank.text(1.0, N - 1 - i, str(r["rank"]), fontsize=4.5, va="center", ha="right", color=GRAY)
    ax_rank.set_title("Rank", fontsize=7, pad=6)

    # final_score 条
    def strip(ax, arr, cmap, vmin, vmax, title, bar=False):
        ax.imshow(arr.reshape(-1, 1), aspect="auto", cmap=cmap, vmin=vmin, vmax=vmax, interpolation="nearest")
        ax.set_xticks([]); ax.set_yticks([])
        ax.set_title(title, fontsize=7, pad=6)
        for sp in ax.spines.values(): sp.set_visible(False)
    strip(ax_score, score, "RdYlBu_r", np.nanmin(score), np.nanmax(score), "Final\nscore")
    strip(ax_bt, bt, "YlGnBu", np.nanmin(bt), np.nanmax(bt), "Combined\nB/T")
    # conservation
    ax_cat.imshow(cat, cmap=ListedColormap(["#D5D9DE", BLUE_LIGHT, BLUE_DARK]), aspect="auto", vmin=0, vmax=2, interpolation="nearest")
    ax_cat.set_xticks([0, 1, 2]); ax_cat.set_xticklabels(["H1N1", "H3N2", "H5N1"], fontsize=7, rotation=45)
    ax_cat.set_yticks([]); ax_cat.set_title("Conservation", fontsize=7, pad=6)
    ax_cat.xaxis.tick_top()
    for sp in ax_cat.spines.values(): sp.set_visible(False)
    # z-PLL / contact
    zl = max(abs(np.nanpercentile(zpll, 2)), abs(np.nanpercentile(zpll, 98)), 1e-9)
    strip(ax_z, zpll, "RdBu_r", -zl, zl, "z-PLL")
    strip(ax_cd, cd, "Blues", np.nanpercentile(cd, 1), np.nanpercentile(cd, 99), "Contact\ndensity")
    # surface/internal
    ax_surf.imshow(surf.reshape(-1, 1), cmap=ListedColormap([INTERNAL, SURFACE]), aspect="auto", vmin=0, vmax=1, interpolation="nearest")
    ax_surf.set_xticks([]); ax_surf.set_yticks([]); ax_surf.set_title("Class", fontsize=7, pad=6)
    for sp in ax_surf.spines.values(): sp.set_visible(False)
    # protein
    ax_prot.set_xlim(0, 1); ax_prot.set_ylim(-0.5, N - 0.5); ax_prot.axis("off")
    for i, r in enumerate(rows):
        ax_prot.text(0.02, N - 1 - i, r["protein"], fontsize=4.5, va="center",
                     color=SURFACE if r["protein"] in ("HA", "NA") else INTERNAL)
    ax_prot.set_title("Protein", fontsize=7, pad=6)

    fig.suptitle("Figure S1 | Complete evidence for all 98 candidate regions (ranked by final score)",
                 fontsize=11, fontweight="bold", x=0.5, y=0.99)
    fig.savefig(f"{SUPP}/S1_full_candidates.png", dpi=300, bbox_inches="tight", facecolor="white", pad_inches=0.03)
    plt.close(fig); print("saved S1_full_candidates.png")

# ================================================================ S2
def figS2():
    epi = "CKLRGVAPLHLGKCNIAGW"
    # H1N1 逐年
    ys = defaultdict(list)
    for r in SeqIO.parse(f"{RAW}/分类序列_H1N1/HA_sequences.fasta", "fasta"):
        mm = re.search(r"/20\d\d", r.description)
        if mm and 2006 <= int(mm.group(0)[1:]) <= 2017:
            ys[int(mm.group(0)[1:])].append(str(r.seq[:(len(r.seq)//3)*3].translate()))
    years = sorted(ys); fracs = [frac(ys[y], epi) for y in years]; ns = [len(ys[y]) for y in years]
    # H1 clades
    meta = {}
    for r in csv.DictReader(open(f"{ROOT}/99_历史项目与参考/早期H1N1系统发育项目/清洗数据/cleaned_metadata.csv")):
        meta[r["Isolate_Id"]] = r
    aln = f"{ROOT}/99_历史项目与参考/早期H1N1系统发育项目/清洗数据/clustalo-H1N1-HA.fa"
    by_clade = defaultdict(list)
    for r in SeqIO.parse(aln, "fasta"):
        cl = meta.get(r.id, {}).get("Clade", "unassigned")
        aa = trans_atg(r.seq)
        if aa and aa.count("X") <= 0.05 * len(aa):
            by_clade[cl].append(aa)
    clades = [c for c, _ in Counter({k: len(v) for k, v in by_clade.items()}).most_common(6)]
    cf_clade = [frac(by_clade[c], epi) for c in clades]; n_clade = [len(by_clade[c]) for c in clades]
    # H3N2
    h3 = list(csv.DictReader(open(f"{MS}/09_H3N2_CROSS_LINEAGE/h3n2_cross_lineage.tsv"), delimiter="\t"))
    ckeys = [k for k in h3[0].keys() if k.startswith("clade_")]
    cnames = [k.replace("clade_", "") for k in ckeys]
    h3mat = np.array([[float(r[k]) if r[k] not in ("", None) else 0.0 for k in ckeys] for r in h3])
    h3lab = [f"{r['protein']} {r['epitope'][:9]}" for r in h3]
    # H3 clade 样本量
    h3n = defaultdict(int)
    for r in SeqIO.parse(f"{ROOT}/99_历史项目与参考/外部参考数据/gisaid_H3N2_HANA.fasta", "fasta"):
        p = r.description.split("|")
        if len(p) > 9 and p[9] == "HA":
            h3n[p[6] if len(p) > 6 and p[6] else "unassigned"] += 1

    fig = plt.figure(figsize=(12, 8.5))
    gs = fig.add_gridspec(2, 2, hspace=0.5, wspace=0.35)

    ax = fig.add_subplot(gs[0, 0])
    size = 15 + 40 * (np.array(ns) / max(ns))
    ax.scatter(years, fracs, s=size, color=BLUE, alpha=0.85, edgecolor="white", linewidth=0.4, zorder=3)
    ax.plot(years, fracs, color=BLUE, lw=1.0, alpha=0.5, zorder=2)
    ax.set_xlabel("Year"); ax.set_ylabel("Fraction ≥ 90% identity"); ax.set_ylim(0, 1.08)
    ax.set_title("point size ~ n sequences", fontsize=6.5, color=GRAY, loc="right", pad=2)
    panel(ax, "a", "H1N1 HA candidate: yearly conservation (2006–2017)")

    ax = fig.add_subplot(gs[0, 1])
    yy = np.arange(6)[::-1]
    ax.scatter(cf_clade, yy, s=70, color=BLUE, zorder=3)
    ax.hlines(yy, 0, cf_clade, color=GRAY, lw=0.6, zorder=2)
    ax.set_yticks(yy)
    ax.set_yticklabels([f"{c[:14]}…" for c in clades], fontsize=6)
    ax.set_xlabel("Conserved fraction"); ax.set_xlim(0, 1.08)
    for y, f, n in zip(yy, cf_clade, n_clade):
        ax.text(f + 0.02, y, f"{f:.2f} (n={n})", va="center", fontsize=6)
    panel(ax, "b", "Conservation across six H1N1 clades (2015–2025)")

    ax = fig.add_subplot(gs[1, 0])
    im = ax.imshow(h3mat, cmap="Blues", vmin=0, vmax=1, aspect="auto")
    ax.set_xticks(range(len(cnames))); ax.set_xticklabels([c.replace(".", ".\n", 1) for c in cnames], fontsize=5)
    ax.set_yticks(range(len(h3lab))); ax.set_yticklabels(h3lab, fontsize=6)
    for i in range(h3mat.shape[0]):
        for j in range(h3mat.shape[1]):
            if h3mat[i, j] > 0.05:
                ax.text(j, i, f"{h3mat[i,j]:.2f}", ha="center", va="center", fontsize=5, color="white")
    ax.tick_params(length=0)
    cb = fig.colorbar(im, ax=ax, fraction=0.046, pad=0.03); cb.set_label("conserved fraction", fontsize=7); cb.ax.tick_params(labelsize=6)
    panel(ax, "c", "H3N2 clade conservation of 10 surface candidates")

    ax = fig.add_subplot(gs[1, 1])
    order = sorted(h3n.items(), key=lambda x: -x[1])
    names = [f"{c[:12]}" for c, _ in order]; vals = [n for _, n in order]
    ax.bar(range(len(order)), vals, color=ORANGE, width=0.55)
    ax.set_xticks(range(len(order))); ax.set_xticklabels(names, fontsize=5.5, rotation=45)
    ax.set_ylabel("HA sequences")
    for i, v in enumerate(vals):
        ax.text(i, v + 5, str(v), ha="center", fontsize=5.5)
    panel(ax, "d", "H3N2 clade sample sizes (HA sequences)")

    fig.suptitle("Figure S2 | Temporal and lineage conservation validation",
                 fontsize=11, fontweight="bold", x=0.5, y=0.99)
    fig.savefig(f"{SUPP}/S2_temporal_lineage.png", dpi=300, bbox_inches="tight", facecolor="white", pad_inches=0.03)
    plt.close(fig); print("saved S2_temporal_lineage.png")

# ================================================================ S3
def figS3():
    def mode_seqs(sub, prot):
        recs = list(SeqIO.parse(f"{RAW}/分类序列_{sub}/{prot}_sequences.fasta", "fasta"))
        aa = [str(r.seq[:(len(r.seq)//3)*3].translate()) for r in recs]
        ln = Counter(len(s) for s in aa).most_common(1)[0][0]
        return [s for s in aa if len(s) == ln and s.count("X") <= 0.05 * len(s)]
    def cons_ident(sub, prot):
        seqs = mode_seqs(sub, prot); L = len(seqs[0])
        c = [Counter(s[i] for s in seqs).most_common(1)[0][0] for i in range(L)]
        return np.array([sum(1 for s in seqs if s[i] == c[i]) / len(seqs) for i in range(L)])
    # T-cell benchmark
    lines = open(f"{MS}/01_ESM_INCREMENTAL_BENCHMARK/benchmark_results.tsv").read().splitlines()
    m = {}
    for l in lines[1:]:
        p = l.split("\t")
        if p[0].startswith("incremental"): continue
        m[p[0]] = dict(auc=float(p[1]), lo=float(p[2]), hi=float(p[3]), auprc=float(p[4]))
    # B-cell
    cons = {}
    for sub in ["H1N1", "H3N2", "H5N1"]:
        for prot in ["HA", "NA"]:
            cons[(sub, prot)] = str(list(SeqIO.parse(f"{CONS}/{sub}_{prot}.fasta", "fasta"))[0].seq)
    def region_pll(sub, prot, seq):
        s = cons[(sub, prot)]; i = s.find(seq)
        if i < 0:
            for j in range(len(s) - len(seq) + 1):
                if sum(a != b for a, b in zip(seq, s[j:j+len(seq)])) <= 1: i = j; break
        if i < 0: return np.nan
        pll = np.load(f"{ESM}/{sub}_{prot}_pll.npy"); zz = z(pll)
        return np.mean(zz[i+1:i+1+len(seq)])
    c06 = list(csv.DictReader(open(f"{MS}/08_BCELL_EXTERNAL_BENCHMARK/06_BENCHMARK_READY_DATASET.tsv"), delimiter="\t"))
    bg = list(csv.DictReader(open(f"{MS}/08_BCELL_EXTERNAL_BENCHMARK/07_MATCHED_BACKGROUND.tsv"), delimiter="\t"))
    neut, bind, bgr = [], [], []
    for c in c06:
        v = region_pll(c["subtype"], c["protein"], c["representative_seq"])
        if np.isnan(v): continue
        (neut if c["label_level"] == "L3_neutralization" else bind).append(v)
    for r in bg:
        v = region_pll(r["subtype"], r["protein"], r["bg_sequence"])
        if not np.isnan(v): bgr.append(v)
    neut, bind, bgr = np.array(neut), np.array(bind), np.array(bgr)

    fig = plt.figure(figsize=(12, 9))
    gs = fig.add_gridspec(2, 3, hspace=0.5, wspace=0.4)

    # 三亚型 hexbin
    for idx, sub in enumerate(["H1N1", "H3N2", "H5N1"]):
        pll = np.load(f"{ESM}/{sub}_HA_pll.npy"); seq = open(f"{ESM}/{sub}_HA_seq.txt").read().strip()
        ci = cons_ident(sub, "HA"); n = min(len(seq), len(ci)); zz = z(pll[1:1+n])
        rho, _ = stats.spearmanr(ci[:n], zz)
        rp, _ = stats.pearsonr(ci[:n], zz)
        ax = fig.add_subplot(gs[0, idx])
        ax.hexbin(ci[:n], zz, gridsize=40, cmap="Blues", mincnt=1, bins="log")
        ax.set_xlabel("MSA identity"); ax.set_ylabel("z-PLL")
        ax.text(0.03, 0.96, f"{sub}\nρ={rho:.2f}  r={rp:.2f}", transform=ax.transAxes, fontsize=8, va="top", fontweight="bold")
        panel(ax, "abc"[idx], f"{sub} HA: conservation vs PLL")

    # T-cell AUROC 森林图
    ax = fig.add_subplot(gs[1, 0])
    models = ["A_conservation", "C_conservation+PLL", "D_conservation+PLL+contact"]
    mlab = ["conservation", "+PLL", "+PLL+contact"]
    for y, k in zip([2, 1, 0], models):
        r = m[k]
        ax.errorbar(r["auc"], y, xerr=[[r["auc"]-r["lo"]], [r["hi"]-r["auc"]]], fmt="o",
                    color=[GRAY, BLUE, ORANGE][y], ms=6, lw=1.5, capsize=3)
    ax.axvline(0.5, color=INK, ls="--", lw=0.7)
    ax.set_yticks([2, 1, 0]); ax.set_yticklabels(mlab, fontsize=7)
    ax.set_xlabel("AUROC"); ax.set_xlim(0.35, 0.7)
    panel(ax, "d", "T-cell benchmark (AUROC ± 95% CI)")

    # B-cell 分布
    ax = fig.add_subplot(gs[1, 1])
    groups = [("Background", bgr, GRAY), ("Binding", bind, "#E8A87C"), ("Neutralizing", neut, RED)]
    for i, (name, vals, c) in enumerate(groups):
        ax.boxplot(vals, positions=[i], widths=0.5, patch_artist=True,
                   boxprops=dict(facecolor=c, alpha=0.35, lw=0.8),
                   medianprops=dict(color=INK, lw=1.2), whiskerprops=dict(lw=0.8), capprops=dict(lw=0.8),
                   flierprops=dict(markersize=0))
        jitter = np.random.default_rng(i).normal(0, 0.06, len(vals))
        ax.scatter(i + jitter, vals, s=12, color=c, alpha=0.75, edgecolor="white", linewidth=0.3, zorder=3)
        ax.text(i, np.mean(vals), f"mean {np.mean(vals):+.2f}\nn={len(vals)}", va="center", fontsize=7)
    ax.axhline(0, color=INK, ls="--", lw=0.6)
    ax.set_xticks(range(3)); ax.set_xticklabels([g[0] for g in groups], fontsize=8)
    ax.set_ylabel("mean z-PLL")
    panel(ax, "e", "B-cell epitopes: mean z-PLL by class")

    # 数据流
    ax = fig.add_subplot(gs[1, 2])
    stages = [("IEDB\nB-cell", 122), ("HA/NA", 70), ("Mapped", 33), ("Clusters", 22)]
    xs = np.arange(4)
    ax.bar(xs, [s[1] for s in stages], color=["#5B6672", "#7A8794", ORANGE, RED], width=0.55)
    ax.set_xticks(xs); ax.set_xticklabels([s[0] for s in stages], fontsize=6.5)
    ax.set_ylabel("Count")
    for x, (_, n) in zip(xs, stages):
        ax.text(x, n + 2, str(n), ha="center", fontsize=7)
    panel(ax, "f", "B-cell data flow (122 → 70 → 33 → 22)")

    fig.suptitle("Figure S3 | Model and external benchmark validation",
                 fontsize=11, fontweight="bold", x=0.5, y=0.99)
    fig.savefig(f"{SUPP}/S3_benchmark.png", dpi=300, bbox_inches="tight", facecolor="white", pad_inches=0.03)
    plt.close(fig); print("saved S3_benchmark.png")

if __name__ == "__main__":
    figS1(); figS2(); figS3()
    print("SUPPLEMENTARY FIGURES DONE")
```
