## Supplementary material for "Conserved influenza A epitope candidate regions and a benchmark of ESM-2 sequence features": Li_et_al_Supplementary_Data: README.md

# Manuscript Strengthening — 总报告

生成日期：2026-08-13
原则：只使用真实项目数据；所有负面/不显著结果如实保留；不伪造。

---

## 一、四项核心分析结果（诚实）

### 01 ESM-2 incremental benchmark —— 【关键负面结果】

**问题**：ESM-2 PLL 是否提供超越保守性的增量价值？

| 模型 | AUROC [95% CI] | AUPRC |
|---|---|---|
| A 保守性 | 0.526 [0.448–0.610] | 0.533 |
| C +PLL | 0.529 [0.454–0.607] | 0.547 |
| D +contact | 0.559 [0.478–0.634] | 0.566 |

- **Incremental gain（C vs A）：ΔAUROC +0.004，p=0.455 → 不显著**
- D vs C：+0.031，p=0.118 → 不显著
- PLL vs conservation（匹配集内）：ρ=−0.01（正交）

**结论**：ESM-2 PLL **没有**为 T-cell 表位识别提供超越保守性的增量价值。B-cell（n=4）无法做正式 benchmark。

### 02 MSA conservation —— 【数据异质 + 中等相关】

- **原始序列高度异质**：HA 有 40–49 种不同长度（混有部分/截断/移码序列）。
- MAFFT 直接对原始数据对齐产生垃圾（29–56% gap 列，共识错乱）。
- **mode-length 过滤是正确的**；过滤后 PLL vs identity ρ=0.25–0.39（HA），vs Shannon 熵 ρ=−0.25 至 −0.37。

**结论**：PLL 与保守性**中等正相关**（ρ≈0.25–0.39），但远非 1.0 → 部分反映保守性、携带正交信息。

### 03 temporal validation —— 【功效不足】

- 表面候选只有 8 个，ESM vs 传统 retention 完全相同（0.684），p=0.521。
- **无法回答"ESM 优先候选是否未来更稳定"**（数据量不足，非结果不显著）。
- 候选本身在 2013–2017 保留率约 68%（中等时间稳定性）。

### 04 structural accessibility —— 【contact-density 无效】

| 蛋白 | RSA vs contact-density | RSA vs PLL |
|---|---|---|
| HA (1RU7) | +0.147 | +0.079 (n.s.) |
| NA (2HTY) | **−0.132** | +0.120 |

- **contact-density 不是可及性代理**：HA 弱正、NA 弱负，方向不一致，表面/埋藏几乎不分离。
- PLL 有弱但方向一致的表面信号（NA 表面 PLL +0.42 vs 埋藏 −0.01）。

**结论**：contact-density **必须从主排序分数移除**；真正的可及性需结构映射 RSA。

---

## 二、Manuscript 一致性冲突（05 审计）

1. **netMHCpan 阈值**：manuscript 写 `%Rank < 0.02`，实际脚本是 `%Rank_EL < 2`（2%）+ `Aff < 500 nM`。必须修正。
2. **match_count ≠ 亚型数**：match_count 是"地区×亚型×阈值组合数"；实测 14/98 候选在我的全球共识中 0 亚型保守。须用重算的真实跨亚型数。
3. **序列数不一致**：1544（核心）≠ raw（1154/1409/…）≠ mode-length（850/1006/…）。需 data-flow 表。
4. **B-cell 工具名缺失**：数据中无明确工具名。
5. **AlphaFold 是片段**：140 PDB 是嵌合片段，非全长。

---

## 三、核心主张应进一步收束

当前 manuscript 已从"广谱疫苗候选"降调为"探索性优先排序特征"。**本审计表明还需再收束**：

> ESM-2 零样本序列似然是"与保守性中等相关、但对 T-cell 表位识别无增量价值、contact-density 非可及性代理"的探索性特征。它对广谱疫苗候选的实用增益**未被本数据证明**，需在独立 B-cell 表位集、结构可及性和实验中验证。

具体：
1. 删除"ESM-2 改善表位识别"的任何暗示（benchmark 无增量）。
2. contact-density 移出主排序分数。
3. 修正 netMHCpan 阈值和 match_count 定义。
4. 明确数据质量（原始序列异质、只到 2017 年）。

---

## 四、目录与产物

```
07_MANUSCRIPT_STRENGTHENING/
  00_AUDIT/STRENGTHENING_PLAN.md      # 审计计划
  01_ESM_INCREMENTAL_BENCHMARK/       # benchmark_results.tsv, benchmark_auroc.svg, analysis_summary.md
  02_MSA_CONSERVATION/                # alignments/, msa_conservation_correlation_filtered.tsv, analysis_summary.md
  03_TEMPORAL_VALIDATION/             # temporal_cohort.tsv, analysis_summary.md
  04_STRUCTURAL_ACCESSIBILITY/        # 2HTY.pdb, analysis_summary.md
  05_MANUSCRIPT_CONSISTENCY/          # dataflow_audit.md, consistency_issues.md, cross_subtype_true.csv
```

---

## 五、未完成/需用户决策

1. **B-cell benchmark 不可做**（n=4）。需用户决定是否接受"仅 T-cell 正式 benchmark"。
2. **2017 之后无数据**。temporal 只能 cohort split，无法扩展。
3. **netMHCpan 阈值修正**：确认是 `%Rank < 2%`（已从脚本核实）。
4. **B-cell 工具名**：需用户确认原始管线的 B-cell 预测器。
5. **manuscript.docx 未改**：按计划，最后才改；本报告先给出所有需改的地方。
