## Supplementary material for "Conserved influenza A epitope candidate regions and a benchmark of ESM-2 sequence features": Li_et_al_Supplementary_Data: recalc_table2.py

```
#!/usr/bin/env python3
"""
重算 Table 2 的跨亚型保守性：用 cross_subtype_true.csv 的真实跨亚型数
替换 ai_ranking.csv 的 match_count（= 地区×亚型×阈值组合数，语义错误）。

输出：
- tables/Table2_candidates_full_corrected.tsv（98 候选 + 真实跨亚型列）
- tables/Table2_top15.md（top15 摘要，含 n_exact_subtypes）
"""
import csv
from collections import Counter

BASE = "/media/qxl/My_Passport/流行病学数据/广谱流感疫苗候选课题/07_MANUSCRIPT_STRENGTHENING"

# 1. 真实跨亚型数据（protein 归一化 M -> M1）
cst = {}
for r in csv.DictReader(open(f"{BASE}/05_MANUSCRIPT_CONSISTENCY/cross_subtype_true.csv")):
    prot = "M1" if r["protein"] == "M" else r["protein"]
    cst[(prot, r["epitope"])] = r

# 2. 原 Table2
rows = list(csv.DictReader(open(f"{BASE}/tables/Table2_candidates_full.tsv"), delimiter="\t"))

def n_exact(r):
    return sum(1 for s in ("H1N1", "H3N2", "H5N1") if r[s] == "exact")

# 3. join + 写纠正版
out_rows = []
for r in rows:
    key = ("M1" if r["protein"] == "M" else r["protein"], r["epitope"])
    cs = cst.get(key, {})
    h1 = cs.get("H1N1", "-"); h3 = cs.get("H3N2", "-"); h5 = cs.get("H5N1", "-")
    ne = sum(1 for s in (h1, h3, h5) if s == "exact")
    out_rows.append(dict(
        rank=r["rank"], protein=r["protein"], epitope=r["epitope"], aa_len=r["aa_len"],
        original_match_count=r["match_count"],  # 语义澄清：原管线匹配次数
        n_exact_subtypes=ne, H1N1=h1, H3N2=h3, H5N1=h5,
        combined_BT=r["combined_BT"], zPLL=r["zPLL"],
        contact_density=r["contact_density"], mapped_subtype=r["mapped_subtype"],
        final_score=r["final_score"]))

with open(f"{BASE}/tables/Table2_candidates_full_corrected.tsv", "w", newline="") as f:
    w = csv.DictWriter(f, fieldnames=list(out_rows[0].keys()), delimiter="\t")
    w.writeheader(); w.writerows(out_rows)

# 4. top15 markdown（含跨亚型数）
surf = [r for r in out_rows if r["protein"] in ("HA", "NA")]
internal = [r for r in out_rows if r["protein"] not in ("HA", "NA")]
print("=== 真实跨亚型精确保守分布 ===")
print("全体 98 候选 n_exact:", dict(sorted(Counter(r["n_exact_subtypes"] for r in out_rows).items())))
print("内部蛋白 n_exact:", dict(sorted(Counter(r["n_exact_subtypes"] for r in internal).items())))
print("表面(HA/NA) n_exact:", dict(sorted(Counter(r["n_exact_subtypes"] for r in surf).items())))
print("\n表面候选详表:")
for r in surf:
    print(f"  {r['protein']:3s} {r['epitope'][:38]:38s} n_exact={r['n_exact_subtypes']}  H1N1={r['H1N1']:6s} H3N2={r['H3N2']:6s} H5N1={r['H5N1']:6s}")

with open(f"{BASE}/tables/Table2_top15.md", "w") as f:
    f.write("| Rank | Protein | Epitope | n_exact_subtypes | Combined B/T | z-PLL | Contact-density |\n")
    f.write("|---|---|---|---|---|---|---|\n")
    for r in out_rows[:15]:
        f.write(f"| {r['rank']} | {r['protein']} | {r['epitope'][:20]} | {r['n_exact_subtypes']} | "
                f"{float(r['combined_BT']):.2f} | {float(r['zPLL']):.2f} | {float(r['contact_density']):.2f} |\n")

print(f"\n已写 tables/Table2_candidates_full_corrected.tsv ({len(out_rows)} 行) 与 tables/Table2_top15.md")
```
