## Supplementary material for "Conserved influenza A epitope candidate regions and a benchmark of ESM-2 sequence features": Li_et_al_Supplementary_Data: renumber_figures.py

```
#!/usr/bin/env python3
"""把 manuscript_v2.md 的图编号从 18 张改为 7 主文 + 5 附录。"""
P = "/media/qxl/My_Passport/流行病学数据/广谱流感疫苗候选课题/07_MANUSCRIPT_STRENGTHENING/manuscript_v2.md"
s = open(P, encoding="utf-8").read()

# ---- 正文引用替换（旧 -> 新）----
repl = [
    ("(Fig. 13, Supplementary Note 1)", "(Supplementary Fig. S4, Supplementary Note 1)"),
    ("(NP, M1, NEP, PA, PB1, PB2; Fig. 10)", "(NP, M1, NEP, PA, PB1, PB2; Supplementary Fig. S3)"),
    ("H5N1 consensus sequences (Fig. 1), but", "H5N1 consensus sequences (Fig. 1A), but"),
    ("across 2010–2025 (Fig. 2), with no", "across 2010–2025 (Fig. 1B), with no"),
    ("purifying selection (Fig. 15).", "purifying selection (Fig. 7)."),
    ("variation (0.013; Fig. 16).", "variation (0.013; Fig. 1C)."),
    ("not across subtypes (Fig. 16, Fig. 17).", "not across subtypes (Fig. 1C, Fig. 1D)."),
    ("treated as exploratory (Fig. 3).", "treated as exploratory (Supplementary Fig. S1)."),
    ("validated immunogen designs (Fig. 4).", "validated immunogen designs (Supplementary Fig. S2)."),
    ("conservation proxy (Fig. 5).", "conservation proxy (Fig. 2A)."),
    ("three-dimensional structure (Fig. 9).", "three-dimensional structure (Fig. 2B)."),
    ("p = 0.12) (Fig. 6).", "p = 0.12) (Fig. 3)."),
    ("buried residues (Fig. 7).", "buried residues (Fig. 4)."),
    ("5 neutralization-supported; Fig. 12)", "5 neutralization-supported; Fig. 5B)"),
    ("Mann–Whitney p < 0.001; Fig. 11)", "Mann–Whitney p < 0.001; Fig. 5A)"),
    ("native subtype consensus (Fig. 8, Fig. 18).", "native subtype consensus (Fig. 6)."),
    ("global population, respectively (Fig. 14).", "global population, respectively (Supplementary Fig. S5)."),
]
missed = []
for a, b in repl:
    if a in s:
        s = s.replace(a, b)
    else:
        missed.append(a)
if missed:
    print("!! 未匹配的替换（需人工核对）:")
    for m in missed:
        print("   ", m)
else:
    print("正文 17 处图引用全部替换成功")

# ---- 图例段整体替换 ----
old_legends_start = s.find("## Figure legends")
old_legends_end = s.find("## Supplementary Note 1")
assert old_legends_start != -1 and old_legends_end != -1

new_legends = """## Figure legends

**Fig 1. Candidate discovery and conservation.** (A) Cross-subtype conservation of the top candidate regions across H1N1/H3N2/H5N1; all 38 cross-subtype-conserved regions are internal proteins (PB2, PB1, PA, NP), and the ten surface-glycoprotein candidates are subtype-specific. (B) Temporal stability of the top HA candidate within its native H1N1 subtype (2005–2017) and the H3N2-native NA candidate (2010–2025). (C) Cross-lineage conservation of the top HA candidate across six major 2015–2025 H1N1 lineages (identity ≥90%). (D) Surface candidates are subtype-specific: presence in each subtype consensus and conservation across six 2010–2025 H3N2 clades.

**Fig 2. ESM-2 PLL recapitulates conservation and localizes to functional regions.** (A) ESM-2 PLL versus MSA conservation for HA across three subtypes (Spearman ρ annotated). (B) Three-dimensional HA structure (PDB 1RU7), Cα atoms colored by ESM-2 z-PLL, showing high PLL in the globular head.

**Fig 3. T-cell benchmark.** AUROC of conservation, +PLL, and +contact-density (95% CI), showing no incremental value of ESM-2.

**Fig 4. Contact-density versus FreeSASA RSA** (HA and NA), showing contact-density is not a solvent-accessibility proxy.

**Fig 5. Antibody-epitope (B-cell) benchmark.** (A) Ridge plot of ESM-2 PLL distributions for neutralization-supported, antibody-binding, and matched-background epitopes. (B) Data flow (122 IEDB epitopes → 70 HA/NA → 33 mapped → 22 non-redundant clusters).

**Fig 6. Neutralizing-antibody contact footprints.** (A) Stalk antibodies (CR6261, CR8020, CR9114) mapped to the HA2 stalk. (B) Extended footprints: stalk, receptor-binding site (CH65, C05, S139/1, F045-092), and neuraminidase (CD6), mapped to their native subtype consensus.

**Fig 7. Selection pressure.** Per-codon dN/dS (Nei–Gojobori, smoothed) along H1N1 HA, with candidate regions shaded; the dashed line marks dN/dS = 1.

## Supplementary figure legends

**Supplementary Fig. S1.** Candidate ranking with weight-sensitivity.

**Supplementary Fig. S2.** Illustrative multi-epitope assemblies.

**Supplementary Fig. S3.** Circular proteome map of the nine viral proteins, with conserved candidate regions marked (red = surface glycoproteins, blue = internal).

**Supplementary Fig. S4.** Raw HA sequence length heterogeneity (40–49 distinct lengths per file) motivating mode-length filtering.

**Supplementary Fig. S5.** Population coverage of the screening HLA-I (94.4%) and HLA-II (73.0%) alleles.

"""

s = s[:old_legends_start] + new_legends + s[old_legends_end:]
open(P, "w", encoding="utf-8").write(s)
print("图例段已替换为 7 主文 + 5 附录")
```
