## Supplementary material for "Conserved influenza A epitope candidate regions and a benchmark of ESM-2 sequence features": Li_et_al_Supplementary_Data: analysis_summary.md

# H3N2 跨谱系扩展 + NA 漂移排查 —— 结果摘要（诚实）

生成日期：2026-08-13

## 一、做了什么

1. **H3N2 跨谱系扩展（任务 1）**：用 `gisaid_H3N2_HANA.fasta`（HA 4265 + NA 4264，2010–2025，带 HA clade 标签），对全部 10 个表面候选（HA 4 + NA 6）计算 ≥90% 一致度的保守率，按整体 / 6 个主要 clade / 16 个年份三维度。
2. **NA 漂移排查（任务 2）**：用早期H1N1系统发育项目的 NA 数据（1911 条，2015–2025，带 Clade），对 6 个 NA 候选做同样的跨谱系 + 逐年分析，量化 2017→2025 变化。

翻译方法：ATG-scan（与既有 `lineage_conservation.py` 一致）。已核对：H3N2 NA 活性位点 `RYPGVRCVC` 99.4% 命中、HA2 茎部 `QAAIDQINGKLNR` 79.4% 命中，翻译可靠。

## 二、核心结果

### H3N2 中的表面候选保守率（2010–2025）

| 候选 | 来源亚型 | 全 H3N2 | 结论 |
|---|---|---|---|
| HA `CKLRGVAPLHLGKCNIAGW`（RBD，19aa） | H1N1 | **0.00** | 在 H3N2 完全不存在 |
| HA `KHLLSRINHFEKI`（跨膜，13aa） | H5N1 | 0.00 | 不存在 |
| HA `IMYRLSCEQFNR`（RBD，12aa） | — | 0.00 | 不存在 |
| HA `YTCPRHTGK`（RBD，9aa） | — | 0.00 | 不存在 |
| NA `RYPGVRCVC`（活性位点，9aa） | **H3N2** | **0.99** | 唯一 H3N2 来源候选，稳定 |
| NA 其余 5 个 | H5N1 | 0.00 | 不存在 |

### 候选在 H1N1/H3N2/H5N1 共识中的精确存在性

| 蛋白 | 候选 | H1N1 | H3N2 | H5N1 |
|---|---|---|---|---|
| HA | CKLRGVAPLHLGKCNIAGW | √ | · | · |
| HA | KHLLSRINHFEKI | · | · | √ |
| HA | IMYRLSCEQFNR / YTCPRHTGK | · | · | · |
| NA | 5 个（NFLTEK…/RSPHR…/HLECR…/RIGSKG…/MVIGIV…） | · | · | √ |
| NA | RYPGVRCVC | · | √ | · |

### NA 漂移（任务 2）

- **H3N2 中**：`RYPGVRCVC` 逐年 0.98–1.00，slope ≈ 0.000/yr，**无漂移**。
- **H1N1（2015–2025）中**：5 个 NA 候选 0.00（不存在）；长 NA 活性区 56-mer `RIGSKG…` 仅 0.06，slope −0.002/yr（平台，非漂移）。

## 三、关键诚实结论（修正交接中的说法）

1. **表面糖蛋白候选是「亚型特异」的，不是「跨亚型保守」的。**
   - 明星 HA RBD 候选 `CKLRGVAPLHLGKCNIAGW` 只在 H1N1 精确存在，在 H3N2/H5N1 完全不存在（H3N2 2010–2025 全 0.00）。
   - 6 个 NA 候选里 5 个只在 H5N1、1 个（RYPGVRCVC）只在 H3N2。
   - 交接里说的「38 跨亚型精确保守」候选，经 cross_subtype_true.csv 核实**全是内部蛋白**（PB2/PB1/NP/PA/M1/NS1），**没有一个 HA/NA 表面候选是三亚型精确保守的**。

2. **「NA 候选 2017→2025 漂移」这个说法不成立，应改为「亚型特异、无法映射」。**
   - 交接里"没映射成功"才是关键：NA 候选映射不到 H3N2/H1N1，是因为它们本就是 H5N1 来源，而不是在时间上漂移。
   - 唯一能检验时间稳定性的 H3N2 来源 NA 候选（RYPGVRCVC）在 2010–2025 **稳定不漂移**（99–100%）。
   - 因此"HA 候选跨谱系保守、NA 候选漂移"的差异化叙事是**伪差异**——真正差异是"HA 候选是 H1N1 特异、NA 候选是 H5N1/H3N2 特异"。

3. **对 manuscript 的含义（严重）**：
   - 结果 §2 和 Fig 16 的表述「HA RBD 候选 CKLRGVAPLHLGKCNIAGW 和 NA 活性区 RIGSKG… 是跨亚型最保守的候选」**事实错误**，必须改。
   - "cross-subtype conserved" 只能用于内部蛋白候选；表面候选应改述为"within-subtype / 亚型特异保守"。
   - 摘要里 "38 exactly conserved in all three subtypes" 若指表面候选，需删；若指全部 98 候选，需加注"38 均为内部蛋白"。

## 四、产物

- `h3n2_cross_lineage.tsv` — HA/NA 候选在 H3N2 各 clade 的保守率
- `h3n2_temporal.tsv` — HA/NA 候选在 H3N2 2010–2025 逐年保守率
- `h1n1_na_temporal.tsv` — NA 候选在 H1N1 2015–2025 逐年保守率
- `h3n2_cross_lineage.py` / `na_cross_lineage.py` — 复现脚本
