## Supplementary material for "Conserved influenza A epitope candidate regions and a benchmark of ESM-2 sequence features": Li_et_al_Supplementary_Data: h3n2_cross_lineage.py

```
#!/usr/bin/env python3
"""
H3N2 跨谱系/跨时间保守性验证（扩展自 lineage_conservation.py 的 H1N1 版本）。

问题：2002-2017 发现期得到的 HA/NA 候选表位，在 2010-2025 的 H3N2 中
（4265 HA + 4264 NA，带 HA clade 标签）是否仍然保守？

诚实设计：
- 用 ATG-scan 翻译核苷酸序列（与既有 lineage_conservation.py 一致，已核对 NA
  活性位点 RYPGVRCVC 99.4% 命中、HA2 茎部 QAAIDQINGKLNR 79.4% 命中）。
- 对每个候选，计算在 H3N2 中 >=90% 一致度的序列比例（滑动窗口），
  按整体 / 各主要 HA clade / 各年份三个维度。
- 同时报告每个候选在 H1N1/H3N2/H5N1 共识中的精确存在性（用于揭示
  "跨亚型精确保守" 主张与表面糖蛋白候选的真实关系）。
"""
import csv
from collections import Counter, defaultdict
import numpy as np
from Bio import SeqIO
from Bio.Seq import Seq

BASE = "/media/qxl/My_Passport/流行病学数据/广谱流感疫苗候选课题"
FA = f"{BASE}/99_历史项目与参考/外部参考数据/gisaid_H3N2_HANA.fasta"
CONS = f"{BASE}/08_ESM2_AI分析与论文/data/consensus"
RANK = f"{BASE}/08_ESM2_AI分析与论文/results/ai_ranking.csv"
OUT = f"{BASE}/07_MANUSCRIPT_STRENGTHENING/09_H3N2_CROSS_LINEAGE"

# ---------------------------------------------------------------- translation
def trans_atg(seq):
    s = str(seq).upper().replace("-", "")
    for i in range(0, len(s) - 2):
        if s[i:i+3] == "ATG":
            n = (len(s) - i) // 3
            aa = []
            for c in range(n):
                codon = s[i+3*c:i+3*c+3]
                aa.append("X" if (len(codon) < 3 or "N" in codon) else str(Seq(codon).translate()))
            return "".join(aa)
    return ""

def best_identity(seq, epi):
    """sliding-window best identity of epi against seq (like lineage_conservation.py)"""
    if len(seq) < len(epi):
        return 0.0
    best = 0.0
    for i in range(len(seq) - len(epi) + 1):
        best = max(best, sum(a == b for a, b in zip(epi, seq[i:i+len(epi)])) / len(epi))
    return best

def frac_conserved(seqs, epi, thresh=0.9):
    if not seqs:
        return np.nan
    hits = sum(1 for s in seqs if best_identity(s, epi) >= thresh)
    return hits / len(seqs)

# ------------------------------------------------------------- load H3N2 seqs
def load_h3n2():
    by_gene = {"HA": defaultdict(list), "NA": defaultdict(list)}   # year -> [aa]
    clade_by_gene = {"HA": defaultdict(list), "NA": defaultdict(list)}  # clade -> [aa]
    for r in SeqIO.parse(FA, "fasta"):
        p = r.description.split("|")
        if len(p) <= 9:
            continue
        gene = p[9]
        if gene not in ("HA", "NA"):
            continue
        clade = p[6] if len(p) > 6 and p[6] else "unassigned"
        year = p[7][:4] if len(p) > 7 and p[7][:4].isdigit() else "unknown"
        aa = trans_atg(r.seq)
        if not aa or aa.count("X") > 0.05 * len(aa):
            continue
        by_gene[gene][year].append(aa)
        clade_by_gene[gene][clade].append(aa)
    return by_gene, clade_by_gene

# ------------------------------------------------------------ load candidates
def load_candidates():
    rows = [r for r in csv.DictReader(open(RANK)) if r["protein"] in ("HA", "NA")]
    return rows

# ------------------------------------------- cross-subtype presence in cons
def cross_subtype_presence():
    cons = {}
    for sub in ["H1N1", "H3N2", "H5N1"]:
        cons[sub] = {
            "HA": str(list(SeqIO.parse(f"{CONS}/{sub}_HA.fasta", "fasta"))[0].seq),
            "NA": str(list(SeqIO.parse(f"{CONS}/{sub}_NA.fasta", "fasta"))[0].seq),
        }
    return cons

def main():
    by_gene, clade_by_gene = load_h3n2()
    cands = load_candidates()
    cons = cross_subtype_presence()

    for g in ("HA", "NA"):
        yrs = sorted([y for y in by_gene[g] if y.isdigit()])
        n = sum(len(v) for v in by_gene[g].values())
        print(f"\n{'='*90}\n{g}: {n} 条翻译序列, 年份 {yrs[0]}–{yrs[-1]}")

    # top clades (HA clade labels carried on both HA and NA)
    top_clades = [c for c, _ in Counter({k: len(v) for k, v in clade_by_gene["HA"].items()}).most_common(6)]

    results = []
    for g in ("HA", "NA"):
        print(f"\n--- {g} 候选在 H3N2 中的保守性 ---")
        print(f"{'epitope':30s} {'len':>3s} {'全H3N2':>7s} " +
              " ".join(f"{c[:14]:>14s}" for c in top_clades))
        for r in [c for c in cands if c["protein"] == g]:
            epi = r["epitope"]
            seqs_all = [s for v in by_gene[g].values() for s in v]
            f_all = frac_conserved(seqs_all, epi)
            f_clades = [frac_conserved(clade_by_gene[g].get(c, []), epi) for c in top_clades]
            row = [f_all] + f_clades
            print(f"{epi[:30]:30s} {len(epi):3d} " +
                  f"{f_all:7.2f} " +
                  " ".join(f"{f:14.2f}" if not np.isnan(f) else f"{'NA':>14s}" for f in f_clades))
            results.append(dict(protein=g, epitope=epi, length=len(epi),
                                frac_all=f_all, frac_clades=f_clades))

    # ------------------------------------------------ temporal (per-year) detail
    print("\n\n=== 逐年份保守率 (>=90% identity) ===")
    all_years = sorted({y for g in ("HA", "NA") for y in by_gene[g] if y.isdigit()})
    with open(f"{OUT}/h3n2_temporal.tsv", "w") as f:
        f.write("protein\tepitope\t" + "\t".join(all_years) + "\n")
        for r in cands:
            g = r["protein"]; epi = r["epitope"]
            fracs = [frac_conserved(by_gene[g].get(y, []), epi) for y in all_years]
            f.write(f"{g}\t{epi}\t" +
                    "\t".join("" if np.isnan(x) else f"{x:.3f}" for x in fracs) + "\n")
    print(f"已写 h3n2_temporal.tsv（{len(all_years)} 个年份: {all_years[0]}–{all_years[-1]}）")

    # ------------------------------------------------ NA drift quantification
    print("\n\n=== NA 候选时间漂移量化 (H3N2, 2017 vs 2025) ===")
    na_years = [y for y in by_gene["NA"] if y.isdigit()]
    dr = []
    for r in [c for c in cands if c["protein"] == "NA"]:
        epi = r["epitope"]
        # 早期 (<=2017) vs 后期 (>=2018)，及逐年斜率
        early = [s for y in na_years if y <= "2017" for s in by_gene["NA"][y]]
        late = [s for y in na_years if y >= "2018" for s in by_gene["NA"][y]]
        f_early = frac_conserved(early, epi)
        f_late = frac_conserved(late, epi)
        # per-year slope (only over years with data)
        yseq = [(int(y), by_gene["NA"][y]) for y in na_years]
        fracs = [(int(y), frac_conserved(v, epi)) for y, v in yseq]
        fracs = [(y, f) for y, f in fracs if not np.isnan(f)]
        slope = np.polyfit([x for x, _ in fracs], [f for _, f in fracs], 1)[0] if len(fracs) >= 4 else np.nan
        delta = (f_late - f_early) if not (np.isnan(f_late) or np.isnan(f_early)) else np.nan
        dr.append(dict(epitope=epi, f_early=f_early, f_late=f_late, delta=delta, slope=slope))
        print(f"  {epi[:34]:34s} 早期(≤2017)={f_early:.2f}  后期(≥2018)={f_late:.2f}  Δ={delta:+.2f}  slope={slope:+.3f}/yr")

    # ------------------------------------------------ cross-subtype presence
    print("\n\n=== 候选在 H1N1/H3N2/H5N1 共识中的精确存在性 ===")
    print(f"{'protein':7s} {'epitope':40s} {'H1N1':>6s} {'H3N2':>6s} {'H5N1':>6s}")
    for r in cands:
        g = r["protein"]; epi = r["epitope"]
        marks = [("√" if epi in cons[s][g] else "·") for s in ("H1N1", "H3N2", "H5N1")]
        print(f"{g:7s} {epi[:40]:40s} {marks[0]:>6s} {marks[1]:>6s} {marks[2]:>6s}")

    # save summary
    with open(f"{OUT}/h3n2_cross_lineage.tsv", "w", newline="") as f:
        w = csv.writer(f, delimiter="\t")
        w.writerow(["protein", "epitope", "length", "frac_all_H3N2"] +
                   [f"clade_{c}" for c in top_clades])
        for r in results:
            w.writerow([r["protein"], r["epitope"], r["length"],
                        f"{r['frac_all']:.3f}"] +
                       [f"{x:.3f}" if not np.isnan(x) else "" for x in r["frac_clades"]])
    print(f"\n已写 h3n2_cross_lineage.tsv 与 h3n2_temporal.tsv 到 {OUT}")

if __name__ == "__main__":
    main()
```
