## Supplementary material for "Conserved influenza A epitope candidate regions and a benchmark of ESM-2 sequence features": Li_et_al_Supplementary_Data: na_cross_lineage.py

```
#!/usr/bin/env python3
"""
NA 候选跨谱系/时间保守性（H1N1 2015-2025 数据）——排查"NA 候选 2017→2025 漂移"。

用早期H1N1系统发育项目的 NA 数据（1911 条，2015-2025，带 Clade 标签），
镜像 lineage_conservation.py 对 HA 的做法，检查 6 个 NA 候选表位在近期
H1N1 NA 谱系/年份中是否保守或漂移。

诚实设计：
- NA 序列以 EPI_ISL id 关联 cleaned_metadata.csv 的 Clade / Collection_Date。
- 用 clustalo-H1N1-NA.fa（已比对，含 gap），去掉 gap 后 ATG-scan 翻译。
- 同时报告每个 NA 候选在 H1N1/H3N2/H5N1 共识中的精确存在性，
  以澄清"候选是哪个亚型来源"这一关键前提。
"""
import csv
from collections import Counter, defaultdict
import numpy as np
from Bio import SeqIO
from Bio.Seq import Seq

BASE = "/media/qxl/My_Passport/流行病学数据/广谱流感疫苗候选课题"
ALN = f"{BASE}/99_历史项目与参考/早期H1N1系统发育项目/清洗数据/clustalo-H1N1-NA.fa"
META = f"{BASE}/99_历史项目与参考/早期H1N1系统发育项目/清洗数据/cleaned_metadata.csv"
CONS = f"{BASE}/08_ESM2_AI分析与论文/data/consensus"
RANK = f"{BASE}/08_ESM2_AI分析与论文/results/ai_ranking.csv"
OUT = f"{BASE}/07_MANUSCRIPT_STRENGTHENING/09_H3N2_CROSS_LINEAGE"

def trans_atg(seq):
    s = str(seq).upper().replace("-", "")
    for i in range(0, len(s) - 2):
        if s[i:i+3] == "ATG":
            n = (len(s) - i) // 3
            aa = []
            for c in range(n):
                codon = s[i+3*c:i+3*c+3]
                aa.append("X" if (len(codon) < 3 or "N" in codon) else str(Seq(codon).translate()))
            return "".join(aa)
    return ""

def best_identity(seq, epi):
    if len(seq) < len(epi):
        return 0.0
    best = 0.0
    for i in range(len(seq) - len(epi) + 1):
        best = max(best, sum(a == b for a, b in zip(epi, seq[i:i+len(epi)])) / len(epi))
    return best

def frac_conserved(seqs, epi, thresh=0.9):
    if not seqs:
        return np.nan
    return sum(1 for s in seqs if best_identity(s, epi) >= thresh) / len(seqs)

# ------------------------------------------------- load metadata & sequences
meta = {}
for r in csv.DictReader(open(META)):
    meta[r["Isolate_Id"]] = r

by_clade = defaultdict(list)
by_year = defaultdict(list)
n_loaded = 0
for r in SeqIO.parse(ALN, "fasta"):
    clade = meta.get(r.id, {}).get("Clade", "unassigned")
    date = meta.get(r.id, {}).get("Collection_Date", "")
    year = date[:4] if date[:4].isdigit() else "unknown"
    aa = trans_atg(r.seq)
    if not aa or aa.count("X") > 0.05 * len(aa):
        continue
    n_loaded += 1
    by_clade[clade].append(aa)
    by_year[year].append(aa)

print(f"加载 H1N1 NA {n_loaded} 条（2015-2025）")
years = sorted([y for y in by_year if y.isdigit()])
print(f"年份范围: {years[0]}–{years[-1]}；谱系 top6: {[c for c,_ in Counter({k:len(v) for k,v in by_clade.items()}).most_common(6)]}")

# ------------------------------------------------- candidates
cands = [r for r in csv.DictReader(open(RANK)) if r["protein"] == "NA"]

# ------------------------------------------------- consensus presence
cons = {s: str(list(SeqIO.parse(f"{CONS}/{s}_NA.fasta", "fasta"))[0].seq)
        for s in ("H1N1", "H3N2", "H5N1")}

top_clades = [c for c, _ in Counter({k: len(v) for k, v in by_clade.items()}).most_common(6)]

print("\n=== NA 候选在 H1N1 (2015-2025) 的保守性 ===")
print(f"{'epitope':36s} {'len':>3s} {'全H1N1':>7s} " +
      " ".join(f"{c[:12]:>12s}" for c in top_clades) + "   {'H1N1':>5s} {'H3N2':>5s} {'H5N1':>5s}")
rows = []
for r in cands:
    epi = r["epitope"]
    seqs_all = [s for v in by_clade.values() for s in v]
    f_all = frac_conserved(seqs_all, epi)
    f_clades = [frac_conserved(by_clade.get(c, []), epi) for c in top_clades]
    marks = ["√" if epi in cons[s] else "·" for s in ("H1N1", "H3N2", "H5N1")]
    print(f"{epi[:36]:36s} {len(epi):3d} {f_all:7.2f} " +
          " ".join(f"{f:12.2f}" if not np.isnan(f) else f"{'NA':>12s}" for f in f_clades) +
          f"   {marks[0]:>5s} {marks[1]:>5s} {marks[2]:>5s}")
    rows.append((epi, f_all, f_clades, marks))

# --------------------------------------------- temporal drift (H1N1 NA)
print("\n=== H1N1 NA 候选逐年保守率（>=90% identity）===")
with open(f"{OUT}/h1n1_na_temporal.tsv", "w") as f:
    f.write("epitope\t" + "\t".join(years) + "\n")
    for r in cands:
        epi = r["epitope"]
        fracs = [frac_conserved(by_year.get(y, []), epi) for y in years]
        f.write(f"{epi}\t" + "\t".join("" if np.isnan(x) else f"{x:.3f}" for x in fracs) + "\n")

print("\n=== NA 候选 2017→2025 漂移量化（H1N1）===")
for r in cands:
    epi = r["epitope"]
    early = [s for y in years if y <= "2017" for s in by_year[y]]
    late = [s for y in years if y >= "2018" for s in by_year[y]]
    f_early = frac_conserved(early, epi)
    f_late = frac_conserved(late, epi)
    fracs = [(int(y), frac_conserved(by_year[y], epi)) for y in years]
    fracs = [(y, f) for y, f in fracs if not np.isnan(f)]
    slope = np.polyfit([x for x, _ in fracs], [f for _, f in fracs], 1)[0] if len(fracs) >= 4 else np.nan
    delta = (f_late - f_early) if not (np.isnan(f_late) or np.isnan(f_early)) else np.nan
    print(f"  {epi[:34]:34s} 早期(≤2017)={f_early:.2f}  后期(≥2018)={f_late:.2f}  Δ={delta:+.2f}  slope={slope:+.3f}/yr")

print(f"\n已写 h1n1_na_temporal.tsv 到 {OUT}")
```
