## Supplementary material for "Conserved influenza A epitope candidate regions and a benchmark of ESM-2 sequence features": Li_et_al_Supplementary_Data: analysis_summary.md

# ESM-2 incremental-value benchmark — 结果摘要

生成日期：2026-08-13
固定随机种子：42

## 设计

- **Positives**：IEDB T-cell 表位，n=101（103 个 T-cell 中 101 个成功定位到共识）。
- **Negatives**：matched non-epitope windows（同蛋白、同长度、保守度 ±0.05 匹配、不与任何已知表位重叠），n=101。
- **模型**：
  - A：conservation only
  - C：conservation + PLL
  - D：conservation + PLL + contact-density
  - （B = conservation + conventional B/T score 未做，因外部 IEDB 表位无 netMHCpan 分数）

## 结果

| 模型 | AUROC [95% CI] | AUPRC | P@10 | P@20 |
|---|---|---|---|---|
| A 保守性 | 0.526 [0.448–0.610] | 0.533 | 0.90 | 0.60 |
| C +PLL | 0.529 [0.454–0.607] | 0.547 | 0.70 | 0.70 |
| D +contact | 0.559 [0.478–0.634] | 0.566 | 0.70 | 0.65 |

**Incremental gain（bootstrap 配对检验）**：
- C vs A：ΔAUROC = +0.004 [CI −0.073–+0.076]，p(Δ≤0) = 0.455 → **不显著**
- D vs C：ΔAUROC = +0.031 [CI −0.018–+0.084]，p(Δ≤0) = 0.118 → **不显著**

**PLL vs conservation（匹配集内）**：Spearman ρ = −0.010（p=0.89）→ **几乎完全正交**

## 结论（诚实、负面）

1. 仅用保守性无法区分 IEDB T-cell 表位与匹配负样本（AUROC≈0.53），符合"T-cell 表位不由保守性驱动"。
2. **ESM-2 PLL 未提供超越保守性的增量价值**（ΔAUROC +0.004，p=0.455）。
3. PLL 在此匹配集内与保守性正交，说明 PLL 是"保守性类"特征，但并非独立的 T-cell 表位信号。

## 含义（对 manuscript）

- 这**不是**"ESM-2 无用"，而是明确边界：**ESM-2 PLL ≈ 保守性补充特征，对 T-cell 表位识别无增量**。
- 对 B-cell/抗体靶点的增量价值无法评估（n=4），需更大 B-cell 验证集。
- manuscript 的核心主张应进一步收束：ESM-2 是"探索性优先排序特征"，对 T-cell 表位的增量增益未被证明。

## 复现

`python3 run_benchmark.py` → `benchmark_results.tsv`

## 注意

- P@10 对 Model A 偏高（0.90）可能是 matched-negative 设计的尾部伪影（部分高保守正样本缺少足够匹配负样本）；AUROC/AUPRC 为主要指标。
- 未伪造数据；负面结果如实保留。
