## Supplementary material for "Conserved influenza A epitope candidate regions and a benchmark of ESM-2 sequence features": Li_et_al_Supplementary_Data: run_benchmark.py

```
#!/usr/bin/env python3
"""
ESM-2 incremental-value benchmark (T-cell, IEDB positives).

Question: does ESM-2 PLL add incremental value over positional conservation
for ranking experimentally characterized T-cell epitopes?

Positives: IEDB T-cell epitopes (n=103), located on subtype consensus.
Negatives: matched non-epitope windows (same protein, same length, similar
conservation, non-overlapping with any positive).

Models:
  A: conservation only
  C: conservation + PLL
  D: conservation + PLL + contact-density
(B omitted: conventional netMHCpan B/T score not recomputed for external set)

Metrics: AUROC, AUPRC, bootstrap 95% CI, Precision@10/20, enrichment.
"""
import os, csv
from collections import Counter
import numpy as np
from Bio import SeqIO
from scipy import stats
from sklearn.metrics import roc_auc_score, average_precision_score

BASE = "/media/qxl/My_Passport/流行病学数据/广谱流感疫苗候选课题"
ESM = f"{BASE}/08_ESM2_AI分析与论文/results/esm2"
CONS = f"{BASE}/08_ESM2_AI分析与论文/data/consensus"
RAW = f"{BASE}/01_原始序列与分层数据/序列与地区分层/全球/序列/综合"
OUT = f"{BASE}/07_MANUSCRIPT_STRENGTHENING/01_ESM_INCREMENTAL_BENCHMARK"
os.makedirs(OUT, exist_ok=True)
RNG = np.random.default_rng(42)  # fixed seed

# ---- per-residue positional identity (mode-length, same as 06 script) ----
def mode_seqs(sub, prot):
    p = f"{RAW}/分类序列_{sub}/{prot}_sequences.fasta"
    recs = list(SeqIO.parse(p, "fasta"))
    aa = [str(r.seq[:(len(r.seq)//3)*3].translate()) for r in recs]
    ln = Counter(len(s) for s in aa).most_common(1)[0][0]
    return [s for s in aa if len(s) == ln and s.count("X") <= 0.05*len(s)]

def cons_identity(sub, prot):
    seqs = mode_seqs(sub, prot)
    if not seqs: return None
    L = len(seqs[0])
    cons = [Counter(s[i] for s in seqs).most_common(1)[0][0] for i in range(L)]
    return np.array([sum(1 for s in seqs if s[i]==cons[i])/len(seqs) for i in range(L)])

# ---- load consensus + PLL/contact ----
cons = {}
for sub in ["H1N1","H3N2","H5N1"]:
    for prot in ["HA","NA","M1","NP","NEP","NS1","PA","PB1","PB2"]:
        p = f"{CONS}/{sub}_{prot}.fasta"
        if os.path.exists(p):
            cons[(sub,prot)] = str(list(SeqIO.parse(p,"fasta"))[0].seq)

def zscore(a): return (a - np.nanmean(a)) / np.nanstd(a)

# ---- load IEDB T-cell positives ----
iedb = list(csv.DictReader(open(f"{BASE}/08_ESM2_AI分析与论文/results/iedb_epitopes_scored.csv")))
positives = [r for r in iedb if r["class"] == "T"]
print(f"IEDB T-cell positives: {len(positives)}")

# ---- locate positives + compute features ----
def locate_features(prot, epi):
    """Return (conservation, zPLL, contact_density) for an epitope, best subtype match."""
    best = None
    for sub in ["H1N1","H3N2","H5N1"]:
        s = cons.get((sub, prot), "")
        if not s: continue
        i = s.find(epi)
        if i < 0:
            for j in range(len(s)-len(epi)+1):
                if sum(a!=b for a,b in zip(epi, s[j:j+len(epi)])) <= 1:
                    i = j; break
        if i < 0: continue
        ci = cons_identity(sub, prot)
        pll = np.load(f"{ESM}/{sub}_{prot}_pll.npy")
        acc = np.load(f"{ESM}/{sub}_{prot}_acc.npy")
        z = zscore(pll)
        cons_val = np.mean(ci[i:i+len(epi)])
        pll_val = np.mean(z[i+1:i+1+len(epi)])
        acc_val = np.mean(acc[i:i+len(epi)])
        if best is None or pll_val > best[1]:
            best = (cons_val, pll_val, acc_val)
    return best

pos_feat = []
for r in positives:
    prot = r["protein"]; epi = r["epitope"]
    f = locate_features(prot, epi)
    if f:
        pos_feat.append((prot, epi, f[0], f[1], f[2]))
print(f"Located positives: {len(pos_feat)}")

# ---- build matched negatives ----
# for each positive, sample a non-overlapping window of same length, similar conservation
pos_epi_set = set(r["epitope"] for r in positives)
neg_feat = []
for prot, epi, c0, p0, a0 in pos_feat:
    L = len(epi)
    sub_candidates = []
    for sub in ["H1N1","H3N2","H5N1"]:
        s = cons.get((sub, prot), "")
        if not s or len(s) < L: continue
        ci = cons_identity(sub, prot)
        pll = np.load(f"{ESM}/{sub}_{prot}_pll.npy")
        acc = np.load(f"{ESM}/{sub}_{prot}_acc.npy")
        z = zscore(pll)
        for i in range(len(s)-L+1):
            w = s[i:i+L]
            if w in pos_epi_set: continue  # skip known epitopes
            # matched on conservation
            if abs(np.mean(ci[i:i+L]) - c0) > 0.05: continue
            sub_candidates.append((sub, i, np.mean(ci[i:i+L]), np.mean(z[i+1:i+1+L]), np.mean(acc[i:i+L])))
    # sample 1 matched negative
    if sub_candidates:
        sc = sub_candidates[RNG.integers(len(sub_candidates))]
        neg_feat.append((prot, sc[2], sc[3], sc[4]))
print(f"Matched negatives: {len(neg_feat)}")

# ---- build feature matrix ----
X_pos = np.array([[f[2], f[3], f[4]] for f in pos_feat])  # cons, pll, contact
X_neg = np.array([[f[1], f[2], f[3]] for f in neg_feat])
y = np.array([1]*len(X_pos) + [0]*len(X_neg))
X = np.vstack([X_pos, X_neg])

# ---- model scores ----
cons = X[:,0]; pll = X[:,1]; cd = X[:,2]
def norm(v):
    if v.std()==0: return np.zeros_like(v)
    return (v - v.min())/(v.max()-v.min())
model_scores = {
    "A_conservation": norm(cons),
    "C_conservation+PLL": norm(cons) + norm(pll),
    "D_conservation+PLL+contact": norm(cons) + norm(pll) + norm(cd),
}

# ---- bootstrap metrics ----
def bootstrap_auc(scores, y, n_boot=1000):
    idx = np.arange(len(y))
    aucs = []
    for _ in range(n_boot):
        b = RNG.choice(idx, size=len(idx), replace=True)
        if len(np.unique(y[b])) < 2: continue
        aucs.append(roc_auc_score(y[b], scores[b]))
    return np.array(aucs)

results = {}
for name, s in model_scores.items():
    auc = roc_auc_score(y, s)
    auprc = average_precision_score(y, s)
    auc_boot = bootstrap_auc(s, y)
    ci_lo, ci_hi = np.percentile(auc_boot, [2.5, 97.5])
    # precision@k
    order = np.argsort(-s)
    p10 = y[order[:10]].mean(); p20 = y[order[:20]].mean()
    results[name] = dict(auc=auc, auprc=auprc, ci_lo=ci_lo, ci_hi=ci_hi, p10=p10, p20=p20)
    print(f"{name:28s} AUROC={auc:.3f} [CI {ci_lo:.3f}-{ci_hi:.3f}]  AUPRC={auprc:.3f}  P@10={p10:.2f} P@20={p20:.2f}")

# ---- incremental gain test (C vs A, D vs C) ----
def paired_auc_diff(s1, s2, y, n_boot=1000):
    idx = np.arange(len(y))
    diffs = []
    for _ in range(n_boot):
        b = RNG.choice(idx, size=len(idx), replace=True)
        if len(np.unique(y[b])) < 2: continue
        diffs.append(roc_auc_score(y[b], s2[b]) - roc_auc_score(y[b], s1[b]))
    return np.array(diffs)

dCA = paired_auc_diff(model_scores["A_conservation"], model_scores["C_conservation+PLL"], y)
dDC = paired_auc_diff(model_scores["C_conservation+PLL"], model_scores["D_conservation+PLL+contact"], y)
print(f"\nIncremental gain (C vs A): ΔAUROC = {dCA.mean():+.3f} [CI {np.percentile(dCA,2.5):+.3f}-{np.percentile(dCA,97.5):+.3f}], p(Δ≤0)={np.mean(dCA<=0):.3f}")
print(f"Incremental gain (D vs C): ΔAUROC = {dDC.mean():+.3f} [CI {np.percentile(dDC,2.5):+.3f}-{np.percentile(dDC,97.5):+.3f}], p(Δ≤0)={np.mean(dDC<=0):.3f}")

# ---- PLL orthogonal info: residual PLL vs conservation ----
r = stats.spearmanr(cons, pll)
print(f"\nPLL vs conservation (all regions): Spearman ρ={r.statistic:.3f} (p={r.pvalue:.2e})")

# ---- save TSV ----
with open(f"{OUT}/benchmark_results.tsv","w") as f:
    f.write("model\tAUROC\tAUROC_CI_lo\tAUROC_CI_hi\tAUPRC\tPrecision10\tPrecision20\n")
    for name, m in results.items():
        f.write(f"{name}\t{m['auc']:.4f}\t{m['ci_lo']:.4f}\t{m['ci_hi']:.4f}\t{m['auprc']:.4f}\t{m['p10']:.4f}\t{m['p20']:.4f}\n")
    f.write(f"incremental_C_vs_A\tdelta_AUROC={dCA.mean():.4f}\tp_delta_le_0={np.mean(dCA<=0):.4f}\n")
    f.write(f"incremental_D_vs_C\tdelta_AUROC={dDC.mean():.4f}\tp_delta_le_0={np.mean(dDC<=0):.4f}\n")
print(f"\nsaved benchmark_results.tsv (n_pos={len(pos_feat)}, n_neg={len(neg_feat)})")
```
