## Supplementary material for "Conserved influenza A epitope candidate regions and a benchmark of ESM-2 sequence features": Li_et_al_Supplementary_Data: analysis_summary.md

# Alignment-aware conservation（MSA）— 结果摘要

生成日期：2026-08-13

## 关键发现 1：原始序列高度异质（数据质量）

直接检查原始核苷酸序列的长度分布：

| 蛋白 | 总序列数 | 不同长度种类 | 模态长度 |
|---|---|---|---|
| H1N1 HA | 1154 | **40 种** | 1701 nt (567 aa, 850 条) |
| H3N2 HA | 591 | **29 种** | 1701 nt (567 aa, 335 条) |
| H5N1 HA | 406 | **49 种** | 1704 nt (568 aa, 103 条) |

- 每个蛋白都有数十种不同长度（40–49 种），说明原始数据混有部分序列、截断、移码或长度变异。
- 只有模态长度（1701 nt ≈ 567 aa，H5N1 因多碱基切割位点为 568 aa）对应真正的全长 HA。

## 关键发现 2：MAFFT 直接对原始数据对齐是错的

- 对未过滤的原始数据跑 MAFFT：
  - H1N1 HA：893 列，36.6% 是 gap 列。
  - H3N2 HA：784 列，29.0% gap；MAFFT 共识与 mode-length 共识**只有 8.4% 一致度**。
  - H5N1 HA：1263 列，55.7% gap；一致度 8.4%。
- 结论：因为原始数据混有大量非全长序列，直接 MAFFT 产生垃圾对齐。

## 关键发现 3：mode-length 过滤是正确做法

- 我的 mode-length 共识（先过滤到模态长度、再逐位多数投票）与"过滤后的 MAFFT"应一致。
- H1N1 HA 已验证：MAFFT 共识无 gap 长度 566 vs mode 共识 567，**一致度 = 1.0000**。
- 即：对全长序列，"mode-length positional identity" = "对齐后 positional identity"，我的原始做法**是站得住的**。

## 修正后的相关性（过滤到模态长度 ±2 aa 再 MAFFT）

见 `msa_conservation_correlation_filtered.tsv`（重跑中）。
核心指标是 **PLL vs Shannon entropy**（熵比一致度更全面，能捕捉氨基酸多样性）——用于判断 PLL 是否只是重复传统保守性、还是提供正交信息。

## 对 manuscript 的含义

1. manuscript 应补一句数据质量说明：原始序列长度异质（40–49 种），保守性基于模态长度过滤后的全长序列。
2. 不能用"MSA column-wise conservation"——应写"positional identity over mode-length (full-length) sequences"，或明确"MAFFT 对齐全长序列后的 positional identity"。
3. PLL vs 保守性的相关性需用过滤后的干净数据重报（此前 mode-length 的 ρ=0.32/0.23/0.13 是基于干净全长序列，是可辩护的）。

## 复现

- 数据异质：`检查长度分布`（见上方脚本）。
- 干净 MSA：`run_msa.py`（先 mode-length 过滤，再 MAFFT，算 identity/entropy/gapf/consf，映射到共识，与 PLL 相关）。

## 修正后相关性（过滤到模态长度 ±2 aa，再 MAFFT）— 已完成

| 亚型 | 蛋白 | PLL vs identity ρ | PLL vs entropy ρ |
|---|---|---|---|
| H1N1 | HA | +0.366 | −0.367 |
| H3N2 | HA | +0.247 | −0.245 |
| H5N1 | HA | +0.385 | −0.354 |
| H1N1 | NA | +0.157 | −0.107 |
| H3N2 | NA | +0.107 | −0.107 |
| H5N1 | NA | +0.122 | −0.124 |

**结论**：
- PLL 与 MSA 位置一致度**中等正相关**（HA ρ≈0.25–0.39，NA ρ≈0.11–0.16）。
- PLL 与 Shannon 熵**中等负相关**（符合"熵高=保守低"的预期）。
- 相关系数远非 1.0，说明 PLL **部分反映保守性、但携带大量正交信息**。
- 这与 benchmark 结果一致：PLL 是"保守性类"特征，但对 T-cell 表位无增量。

数据：`msa_conservation_correlation_filtered.tsv`
