## Supplementary material for "Conserved influenza A epitope candidate regions and a benchmark of ESM-2 sequence features": Li_et_al_Supplementary_Data: run_msa.py

```
#!/usr/bin/env python3
"""
Alignment-aware conservation via MAFFT.

For each subtype × protein, translate raw nucleotide sequences and align with
MAFFT, then compute per-column: positional identity, Shannon entropy,
gap frequency, consensus frequency. Correlate ESM-2 PLL with each.

Does NOT overwrite the existing mode-length conservation results.
"""
import os, subprocess, csv
from collections import Counter
import numpy as np
from Bio import SeqIO
from scipy import stats

BASE = "/media/qxl/My_Passport/流行病学数据/广谱流感疫苗候选课题"
RAW = f"{BASE}/01_原始序列与分层数据/序列与地区分层/全球/序列/综合"
ESM = f"{BASE}/08_ESM2_AI分析与论文/results/esm2"
CONS = f"{BASE}/08_ESM2_AI分析与论文/data/consensus"
OUT = f"{BASE}/07_MANUSCRIPT_STRENGTHENING/02_MSA_CONSERVATION"
os.makedirs(f"{OUT}/alignments", exist_ok=True)

SUBS = ["H1N1", "H3N2", "H5N1"]
PROTS = ["HA", "NA", "M1", "NP", "NEP", "NS1", "PA", "PB1", "PB2"]

def translate_aa(nt):
    n = (len(nt)//3)*3
    return str(nt[:n].translate())

def run_mafft(fasta_in, fasta_out):
    subprocess.run(["mafft", "--auto", "--quiet", fasta_in],
                   stdout=open(fasta_out, "w"), check=True)

def column_metrics(seqs):
    """per-column: identity, Shannon entropy, gap fraction, consensus fraction."""
    L = max(len(s) for s in seqs)
    identity, entropy, gapf, consf = [], [], [], []
    for i in range(L):
        col = [s[i] for s in seqs if i < len(s)]
        if not col:
            continue
        n = len(col)
        counts = Counter(col)
        cons_aa = counts.most_common(1)[0][0]
        consf.append(counts[cons_aa]/n)
        gapf.append(col.count('-')/n)
        # Shannon entropy over amino acids (treat gap as a symbol)
        ps = [c/n for c in counts.values()]
        entropy.append(-sum(p*np.log2(p) for p in ps))
        # identity to consensus (excluding gaps)
        non_gap = [c for c in col if c != '-']
        identity.append(sum(1 for c in non_gap if c==cons_aa)/len(non_gap) if non_gap else np.nan)
    return dict(identity=np.array(identity), entropy=np.array(entropy),
                gapf=np.array(gapf), consf=np.array(consf))

def zscore(a): return (a - np.nanmean(a)) / np.nanstd(a)

summary = []
for sub in SUBS:
    for prot in PROTS:
        rawp = f"{RAW}/分类序列_{sub}/{prot}_sequences.fasta"
        if not os.path.exists(rawp):
            continue
        # translate + write
        recs = list(SeqIO.parse(rawp, "fasta"))
        aa = [translate_aa(r.seq) for r in recs]
        aa = [s for s in aa if s.count('X') <= 0.05*len(s)]
        tmp_in = f"{OUT}/alignments/{sub}_{prot}_in.fa"
        tmp_out = f"{OUT}/alignments/{sub}_{prot}_mafft.fa"
        with open(tmp_in, "w") as f:
            for k, s in enumerate(aa):
                f.write(f">seq{k}\n{s}\n")
        run_mafft(tmp_in, tmp_out)
        aln = [str(r.seq) for r in SeqIO.parse(tmp_out, "fasta")]
        m = column_metrics(aln)
        # load ESM-2 consensus + PLL
        cp = f"{CONS}/{sub}_{prot}.fasta"
        if not os.path.exists(cp):
            continue
        cons = str(list(SeqIO.parse(cp, "fasta"))[0].seq)
        pll = np.load(f"{ESM}/{sub}_{prot}_pll.npy")
        z = zscore(pll)
        # compute MSA consensus (majority with gaps)
        L = len(aln[0])
        msa_cons = []
        for i in range(L):
            col = [s[i] for s in aln if i < len(s)]
            msa_cons.append(Counter(col).most_common(1)[0][0])
        # map: walk msa_cons; each non-gap residue = next consensus position
        pos_to_col = [j for j, aa in enumerate(msa_cons) if aa != '-']
        n = min(len(cons), len(pos_to_col))
        # correlation PLL vs each metric (mapping consensus pos -> MSA col)
        for metric in ["identity", "entropy", "gapf", "consf"]:
            vals = m[metric]
            zz = z[1:1+n]
            cols = np.array(pos_to_col[:n])
            v = vals[cols] if len(vals) > 0 else np.array([])
            mask = ~np.isnan(v)
            if mask.sum() < 30: continue
            rho, p = stats.spearmanr(v[mask], zz[mask])
            summary.append((sub, prot, metric, round(rho,3), round(float(p),3)))

with open(f"{OUT}/msa_conservation_correlation.tsv","w") as f:
    f.write("subtype\tprotein\tmetric\tspearman_rho\tp_value\n")
    for s in summary:
        f.write("\t".join(map(str,s))+"\n")

# print key comparisons
print(f"{'subtype':8s} {'prot':5s} {'metric':10s} {'rho':>6s} {'p':>6s}")
for s in summary:
    print(f"{s[0]:8s} {s[1]:5s} {s[2]:10s} {s[3]:6.3f} {s[4]:<6}")

# summary: PLL vs identity vs entropy (HA, pooled)
print("\n=== PLL vs MSA metrics (HA, per subtype) ===")
for s in summary:
    if s[1] == "HA" and s[2] in ("identity","entropy"):
        print(f"  {s[0]} {s[2]:8s} rho={s[3]} p={s[4]}")
print("\nsaved msa_conservation_correlation.tsv")
```
