## Supplementary material for "Conserved influenza A epitope candidate regions and a benchmark of ESM-2 sequence features": Li_et_al_Supplementary_Data: analysis_summary.md

# Temporal validation（cohort split）— 结果摘要

生成日期：2026-08-13

## 设计

- discovery cohort = 2006–2012，validation cohort = 2013–2017（数据最大年份 2017，无法扩展）。
- 只对表面糖蛋白（HA/NA）候选计算（抗原漂移是表面蛋白现象）。

## 结果（诚实）

| 组 | n | validation retention (2013–2017) | temporal slope |
|---|---|---|---|
| ESM-2 PLL 优先 | 8 | 0.684 ± 0.179 | −0.0006/yr |
| 传统 B/T 优先 | 8 | 0.684 ± 0.179 | −0.0006/yr |

Mann-Whitney（ESM retention > conventional）：p = 0.521。

## 关键结论

1. **表面候选只有 8 个**（HA 4 + NA 4 可定位），远不足以做有统计功效的 ESM vs 传统对比。
2. 两组 retention 完全相同（0.684），因为候选太少、两组高度重叠。
3. **诚实结论**：时间验证"功效不足"，无法回答"ESM 优先的候选是否未来更稳定"——不是结果不显著，而是数据量不足以检验。
4. 但候选本身在 2013–2017 验证队列保留率约 68%，说明顶级表面候选有中等时间稳定性（未随时间快速消失，slope≈0）。

## 对 manuscript 的含义

- 不能写"ESM 优先候选时间更稳定"（无证据，n=8）。
- 可写"顶级表面候选在 2007–2017 保持 53–100% 一致度、无显著漂移"（这是描述性结论，已在此前分析建立）。
- 若要检验 ESM vs 传统的差异，需更多表面候选或更长的年份跨度（需新数据）。

数据：`temporal_cohort.tsv`
