## Supplementary material for "Conserved influenza A epitope candidate regions and a benchmark of ESM-2 sequence features": Li_et_al_Supplementary_Data: run_temporal.py

```
#!/usr/bin/env python3
"""
Temporal validation via cohort split (discovery 2006-2012 vs validation 2013-2017).

Question: do candidates selected from early sequences stay conserved in later years?
Compare ESM-2-PLL-prioritized vs conventional-B/T-prioritized candidates.
"""
import os, csv, re
from collections import defaultdict
import numpy as np
from Bio import SeqIO
from scipy import stats

BASE = "/media/qxl/My_Passport/流行病学数据/广谱流感疫苗候选课题"
RAW = f"{BASE}/01_原始序列与分层数据/序列与地区分层/全球/序列/综合"
OUT = f"{BASE}/07_MANUSCRIPT_STRENGTHENING/03_TEMPORAL_VALIDATION"
os.makedirs(OUT, exist_ok=True)

DISCOVERY = (2006, 2012)   # early cohort
VALIDATION = (2013, 2017)  # later cohort

def parse_year(h):
    m = re.search(r"/20\d\d", h); return int(m.group(0)[1:]) if m else None

def year_seqs(sub, prot):
    p = f"{RAW}/分类序列_{sub}/{prot}_sequences.fasta"
    ys = defaultdict(list)
    for r in SeqIO.parse(p, "fasta"):
        y = parse_year(r.description)
        if y and DISCOVERY[0] <= y <= VALIDATION[1]:
            ys[y].append(str(r.seq[:(len(r.seq)//3)*3].translate()))
    return ys

def epitope_frac(seqs, epi, thresh=0.9):
    bests = []
    for s in seqs:
        b = 0.0
        if len(s) >= len(epi):
            for i in range(len(s)-len(epi)+1):
                b = max(b, sum(a==c for a,c in zip(epi, s[i:i+len(epi)]))/len(epi))
        bests.append(b)
    return (np.array(bests) >= thresh).mean() if bests else np.nan

# load candidates
rows = list(csv.DictReader(open(f"{BASE}/08_ESM2_AI分析与论文/results/ai_ranking.csv")))
for r in rows:
    r["final_score"] = float(r["final_score"])
    r["combined"] = float(r["combined"]) if r["combined"] not in ("",None) else np.nan
    r["ai_conservation"] = float(r["ai_conservation"]) if r["ai_conservation"] not in ("",None) else np.nan
rows = [r for r in rows if not np.isnan(r["ai_conservation"])]

# for each candidate, compute discovery vs validation retention (using H1N1 HA/NA as substrate)
results = []
for r in rows:
    prot = r["protein"]
    if prot not in ("HA", "NA"):  # focus surface proteins where temporal drift matters most
        continue
    epi = r["epitope"]
    # try each subtype; use best (most conserved across years)
    best_retention = None; best_slope = None
    for sub in ["H1N1", "H3N2", "H5N1"]:
        ys = year_seqs(sub, prot)
        disc = [s for y in sorted(ys) for s in ys[y] if DISCOVERY[0] <= y <= DISCOVERY[1]]
        val = [s for y in sorted(ys) for s in ys[y] if VALIDATION[0] <= y <= VALIDATION[1]]
        if not disc or not val: continue
        fdisc = epitope_frac(disc, epi)
        fval = epitope_frac(val, epi)
        if np.isnan(fdisc) or np.isnan(fval): continue
        retention = fval
        # yearly slope
        years = sorted(ys); fracs = [epitope_frac(ys[y], epi) for y in years]
        fracs = [f for f in fracs if not np.isnan(f)]
        slope = np.polyfit(range(len(fracs)), fracs, 1)[0] if len(fracs) >= 4 else np.nan
        if best_retention is None or retention > best_retention:
            best_retention = retention; best_slope = slope
    if best_retention is not None:
        results.append(dict(protein=prot, epitope=epi, pll=r["ai_conservation"],
                            combined=r["combined"], retention=best_retention, slope=best_slope))

# split into ESM-2-prioritized (top PLL) vs conventional-prioritized (top B/T)
results.sort(key=lambda x: -x["pll"])
esm_top = results[:15]
results.sort(key=lambda x: -(x["combined"] if not np.isnan(x["combined"]) else 0))
conv_top = results[:15]

def summarize(g, name):
    ret = np.array([r["retention"] for r in g])
    slopes = np.array([r["slope"] for r in g if not np.isnan(r["slope"])])
    print(f"{name}: n={len(g)}, validation retention = {ret.mean():.3f}±{ret.std():.3f}, "
          f"slope = {slopes.mean():+.4f}/yr" if len(slopes) else f"{name}: n={len(g)}")
    return ret

print("=== 时间验证: ESM-2 优先 vs 传统分优先 ===")
ret_esm = summarize(esm_top, "ESM-2 PLL 优先")
ret_conv = summarize(conv_top, "传统 B/T 优先")
if len(ret_esm) and len(ret_conv):
    U, p = stats.mannwhitneyu(ret_esm, ret_conv, alternative="greater")
    print(f"Mann-Whitney (ESM retention > conventional): p={p:.3f}")

with open(f"{OUT}/temporal_cohort.tsv","w") as f:
    f.write("protein\tepitope\tPLL\tB_T_score\tvalidation_retention\ttemporal_slope\n")
    for r in results:
        f.write(f"{r['protein']}\t{r['epitope']}\t{r['pll']:.3f}\t{r['combined']}\t{r['retention']:.3f}\t{r['slope']}\n")
print(f"\nsaved temporal_cohort.tsv ({len(results)} surface candidates)")
```
